# Bacterial retrons enable precise gene editing in human cells

**DOI:** 10.1101/2021.03.29.437260

**Authors:** Bin Zhao, Shi-An A. Chen, Jiwoo Lee, Hunter B. Fraser

**Author notes:** Corresponding Author **Correspondence:** Hunter B. Fraser. These authors contributed equally to this work (Bin Zhao, Shi-An A. Chen, Jiwoo Lee).

## Abstract

Retrons are bacterial genetic elements involved in anti-phage defense. They have the unique ability to reverse transcribe RNA into multicopy single-stranded DNA (msDNA) that remains covalently linked to their template RNA. Retrons coupled with CRISPR-Cas9 in yeast have been shown to improve editing efficiency of precise genome editing via homology-directed repair (HDR). HDR editing efficiency has been limited by challenges associated with delivering extracellular donor DNA encoding the desired mutation. In this study, we tested the ability of retrons to produce msDNA as donor DNA and facilitate HDR by tethering msDNA to guide RNA in HEK293T and K562 cells. Through heterologous reconstitution of retrons from multiple bacterial species with the CRISPR-Cas9 system, we demonstrated HDR rates of up to 11.3%. Overall, our findings represent the first step in extending retron-based precise gene editing to human cells.

## Introduction

Precise genome editing is a promising tool for identifying causal genetic variants and their function, generating disease models, performing gene therapy, among other applications ^1, 2^. Traditionally, precise genome editing with scarless replacement of alleles or insertion of synthetic sequences requires *in vitro* delivery of DNA donors. However, it has proven challenging to induce cells to utilize donor DNA to conduct homology-directed repair (HDR), resulting in non-homologous end joining (NHEJ) repair, which is error-prone ^3^. To date, the most efficient donor delivery systems are *in vitro* synthetic DNA and viral vectors ^4, 5^. Synthetic DNA donors are delivered to cells directly via electroporation or by packaging them into particles without specifically targeting the nucleus, while viral vectors such as adeno-associated virus (AAV) are transduced to enter the nucleus ^6-8^. In both *in vitro* synthetic DNA and viral vector donor delivery, the donors are non-renewable after delivery and are depleted over time, decreasing editing after cell division in mitotic progeny. In addition, neither method scales well for multiplexed editing, which requires specific guide and donor combinations that can only happen by chance with bulk delivery. Finally, synthetic DNA and viral vector donor delivery is limited by cost and labor when scaling up for screening through tens of thousands of individual variants. Therefore, a biological solution enabling *in nucleo* donor generation would fundamentally improve the scalability and multiplexing capabilities for genomic knock-ins.

Retrons have been studied since the 1970s as bacterial genetic elements that encode unique features ^9, 10^, one of which is the production of multicopy single-stranded DNA (msDNA), which has been biochemically purified from retron-expressing cells ^11^. The minimal retron element consists of a contiguous cassette that encodes an RNA (msr-msd) and a reverse transcriptase (RT) which can reverse transcribe the msd into msDNA that is covalently tethered to its template RNA. Retron sequences are diverse among bacterial species but share similar RNA secondary structures ^9^. The RT recognizes the secondary structure of retron RNA hairpin loops in the msr region and subsequently initiates reverse transcription branching off of the guanosine residue flanking the self-annealed double-stranded DNA priming region ^12, 13^. This process has two properties that differentiate retrons from typical viral reverse transcriptases commonly used in biotechnology ^9^. First, the RT targets only the msr-msd from the same retron as its RNA template, providing specificity that may be useful for avoiding off-target reverse transcription ^12^. Second, the RNA template self-anneals intramolecularly *in cis* rather than requiring primers *in trans* to increase efficiency. Combining these two features allows cell-autonomous production of specific single-stranded donor DNA in the nucleus, circumventing the need for external donor delivery through chemical methods or viral vectors. Therefore, when coupled with targeted nucleases, retrons are promising biological sources for generating DNA donors for template-mediated precise genome editing.

We have previously demonstrated that retron-derived msDNA can facilitate template-mediated precise genome editing in yeast ^14^. Cas9-Retron precISe Parallel Editing via homologY (CRISPEY) utilizes retrons to produce donor DNA molecules and vastly improves HDR editing efficiency to ∼96%, allowing for the characterization of the fitness effects of over 16,000 natural genetic variants at single-base resolution. A similar strategy utilizing Cas9, retrons, and single-stranded DNA binding proteins has been demonstrated in bacteria ^15^, in which msDNA can be incorporated into the bacterial genome without the aid of targeted nucleases ^16^. Importantly, a previous study in mouse 3T3 cells provided evidence that msDNA can be produced in mammalian cells at very low amounts ^17^. Hence, we envisioned that retron-generated msDNA could be harnessed for precise gene editing in human cells (Figure 1A).

**Figure 1.**
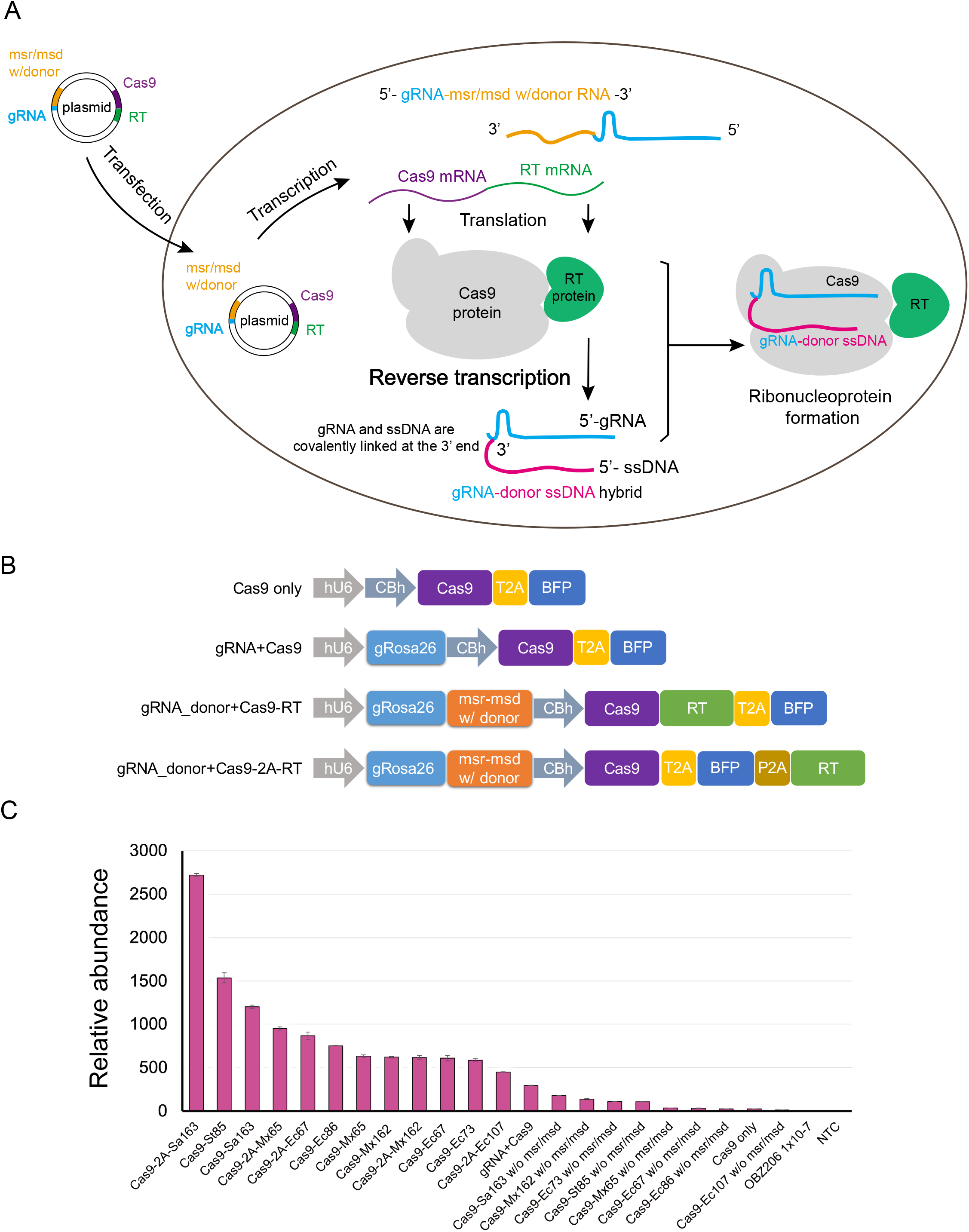
Retrons produce msDNA in human cells. (A) Schematic of our strategy to deploy retrons to generate gRNA-msDNA hybrids as intracellular donors. (B) Schematic showing construct design for the qPCR assay: guide RNA and msr-msd expression was driven by the human U6 promoter; human codon–optimized RT and SpCas9 expression was driven by the CBh promoter; donor templates were inserted into the replaceable regions of msd. T2A, P2A: self-cleaving peptides. (C) Relative abundance of msDNA produced by different retrons. DNA amplified from the same volume of OBZ206 ssDNA at 1 × 10^−7^ ng/μL was set as one-fold to calculate the relative abundance. Data presented as mean ± s.d. (n=2 experiments). NTC: non-transfected control. Cas9 only: cells transfected with the plasmid without gRNA or any retron. gRNA+Cas9: cells transfected with the plasmid expressing both gRNA and Cas9. w/o msr/msd: cells transfected with plasmid that co-express gRNA, Cas9 and RT, but no msr/msd was inserted. Other labels in the X axis indicate cells transfected with plasmids that carry different retron sequences.

## Materials and Methods

All plasmid, retron, and oligo sequences are provided in the Supplemental Information.

### Retron Plasmids

DNA sequences for retrons, primers, and plasmids used in this study are listed in the Supplemental Information. Genes encoding SpCas9 and BFP were obtained from previously reported plasmids (Addgene plasmid # 64323, #64216, #64322, Ralf Kühn lab ^18^; mCherry was amplified from Addgene plasmid # 60954, Jonathan Weissman lab ^19^). Retron genes were synthesized as gBlocks Gene Fragments (Integrated DNA Technologies) or clonal genes (Twist Bioscience). GFP donor genes were synthesized as gBlocks Gene Fragments (Integrated DNA Technologies). Primers and gRNA were synthesized as oligos (Integrated DNA Technologies). The parental vector (Addgene plasmid # 64323, Ralf Kühn’s lab) was digested by restriction endonucleases (New England Biolabs). The digested vector backbone was purified using Monarch DNA Gel Extraction Kit (New England Biolabs) or NucleoSpin Gel and PCR Clean-up Kit (Macherey-Nagel). gRNA targeting BFP (gBFP) was inserted with Golden Gate cloning. PCR was performed using Q5 High-Fidelity DNA Polymerase or Q5 High-Fidelity 2X Master mix (New England Biolabs). PCR products were purified using Monarch PCR & DNA Cleanup Kit (New England Biolabs). RT, msr-msd, and donors were inserted into the digested vector backbone with Gibson Assembly using NEBuilder HiFi DNA Assembly Master Mix (New England Biolabs). Donors were replaced via double digestion by SpeI and AvrII (New England Biolabs). Plasmids were amplified using Stbl3 competent cells prepared with The Mix & Go! E. coli Transformation Kit and Buffer Set (Zymo Research) and extracted by the Plasmid Plus Midi Kit (Qiagen) following the manufacturer’s protocol. Extracted plasmids were measured by Nanodrop (Thermo Fisher Scientific), normalized to the same concentration, and subsequently validated by Sanger sequencing.

### Cell lines and culture

HEK293T BFP and K562 BFP reporter cells were provided by Dr. Jacob Corn (ETH Zürich) and Dr. Christopher D Richardson (UCSB). K562 wildtype cells were provided by Dr. Stanley Qi’s group (Stanford). HEK293T wildtype and HEK293T BFP reporter cells were maintained in Dulbecco’s Modified Eagle’s Medium (DMEM) with GlutaMax (Thermo Fisher Scientific) supplemented with 10% v/v fetal bovine serum (FBS) (Gibco) and 10% penicillin-streptomycin (Thermo Fisher Scientific). K562 wildtype and K562 BFP reporter cells were maintained in PRMI 1640 (Thermo Fisher Scientific) supplemented with 10% v/v FBS (Gibco) and 10% penicillin-streptomycin (Thermo Fisher Scientific). All cells were maintained at 37°C with 5% CO2.

### Transfection

All dilutions and complex formations used Opti-MEM Reduced Serum Medium (Thermo Fisher Scientific). For transfection of one well in a 48-well Poly-D-Lysine coated plate (Corning), 1 μg plasmids were mixed with transfection reagents using Lipofectamine 3000 (Life Technologies) according to the manufacturer’s protocol with the following modifications: after incubating for 30 minutes at room temperature, the DNA-reagent complex was added to each well; then 200,000 HEK293T BFP reporter cells in 200 μL DMEM media were added into each well and mixed gently with the DNA-transfection reagents.

The Neon™ Transfection System 10 μL kit (Thermo Fisher Scientific) was used to transfect K562 and K562 BFP reporter cells according to the manufacturer’s protocol. Cells were washed in Dulbecco’s phosphate-buffered saline (DPBS) (Thermo Fisher Scientific), transfected with 1 μg plasmids at 1050v/20ms/2 pulses and cultured in a 24-well Nunc cell culture plate (Thermo Fisher Scientific) at the density of 200,000/well in PRMI 1640 (Thermo Fisher Scientific) supplemented with 10% FBS (Gibco).

### qPCR

qPCR assay was performed 72 hours post-transfection. K562 cells were spun down at 1,000 rpm for 5 mins. Then, cell pellets were washed in DPBS. Cell pellets were harvested after being spun down again at 1,000 rpm for 5 mins. The gRNA-msDNA hybrid was extracted with the QuickExtract RNA Extraction Solution (Lucigen) according to the manufacturer’s protocol. The extract was digested using double-stranded DNase (Thermo Fisher Scientific). The digested product was purified by using the ssDNA/RNA Clean & Concentrator Kit (Zymo Research). The purified product was then used as the qPCR template. qPCR primers are listed in Supplemental Note 1. The qPCR assay was carried out using iQ SYBR Green Supermix (Bio-Rad). qPCR data was collected on the CFX384 Touch™ Real-Time PCR Detection System (Bio-Rad). We performed a sequential 10x dilution and used this ssDNA as a measurement standard to generate a series of positive signals, which reflected the slope of log-linear regions in a qPCR assay (Supplemental Figure 1).

### Flow cytometry

293T BFP and K562 BFP reporter cells were washed in DPBS (Thermo Fisher Scientific), resuspended in DPBS (Thermo Fisher Scientific), supplemented with 5% v/v FBS (Gibco) at a concentration of 1,000,000 cells/mL, and measured via flow cytometry. Transfected cells were isolated by using a red fluorescence protein mCherry, which was co-expressed with Cas9 and RT. Editing outcomes were recorded 72 hours, 96 hours and 7 days post-transfection or electroporation on an Attune NxT flow cytometer (Invitrogen). For 293T BFP reporter cells, data at 72 hours are presented in the figures. For K562 BFP reporter cells, data at 96 hours are presented in the figures. All plots were analyzed using FlowJo v 10.7.1.

## Results

### Retrons produce msDNA in human cells

To implement the CRISPEY strategy in human cells, we first tested whether msDNA can be produced in human cells. To maximize the chance of msDNA production, we estimated the expression of multiple candidate retrons. To date, hundreds of putative retrons have been characterized by computational analyses, among which 16 were experimentally validated to produce msDNA via *in vitro* assays ^9^. We codon-optimized eight fully annotated RTs with publically available protein sequences and synthesized the corresponding retron RNA to enable heterologous expression in human cells (Table 1, Supplemental Table 1 and Supplemental Note 1). Since the biosynthesis of msDNA requires both msr-msd and RT, we combined the sequences of msr-msd and RT with SpCas9/sgRNA into a single multicistronic vector. We drove the transcription of the chimeric sgRNA-msr-msd by the RNA polymerase III promoter U6 and embedded the donor DNA templates in the replaceable regions within msd sequences (Figure 1B). To measure the total free gRNA-msDNA product, we utilized a gRNA targeting mouse gene Rosa26 and re-constructed the CRISPR-Cas9 vectors that were established in Chu et al. ^18^ (See Materials and Methods). We drove the co-expression of RT and SpCas9 by the CBh promoter, an RNA polymerase II promoter that provides robust and long-term expression ^20^. We engineered Cas9-RT fusion proteins to test whether spatial colocalization of these proteins improved their editing efficiency. In parallel, we employed the 2A self-cleaving peptide-based multi-gene expression system to co-express Cas9 and RT ^21^ in case the functions of Cas9 and/or RT were impeded by steric hindrance.

**Table 1.** Summary of retrons used in this study. Adapted from Simon et al. ^9^. Sequences are shown in Supplemental Note 1.

| Retron | Species | Bacterial taxon | Reference | Source |
| --- | --- | --- | --- | --- |
| Sa163 | Stigmatella aurantiaca | Deltaproteobacteria | Yee et al. 1984 | NCBI Reference Sequence: ZP_01463804.1 |
| Ec86 | Escherichia coli | Gammaproteobacteria | Lim & Maas 1989 | Uniprot: P23070 |
| Mx65 | Myxococcus xanthus | Deltaproteobacteria | Inouye et al. 1990 | Uniprot: P23071 |
| Mx162 | Myxococcus xanthus | Deltaproteobacteria | Yee et al. 1984 | Uniprot: P23072 |
| Ec73 | Escherichia coli | Gammaproteobacteria | Sun et al. 1991 | Uniprot: Q7M2A9 |
| Ec107 | Escherichia coli | Gammaproteobacteria | Herzer et al. 1992 | Uniprot: Q05804 |
| St85 | Salmonella enterica Typhimurium | Gammaproteobacteria | Ahmed & Shimamoto 2003 | GenBank: ACY91017.1 |
| Ec67 | Escherichia coli | Gammaproteobacteria | Lampson et al. 1989 | Uniprot: P21325 |

To test whether retrons can generate msDNA in human cells, we transfected the multicistronic plasmids into human K562 cells and measured msDNA by qPCR. Since bacterial retron products are normally absent in human cells, we synthesized a single-stranded DNA (ssDNA) as the standard template, which is the same as the DNA donor template inserted into the plasmids (Supplemental Note 1). A summary of retrons that generated msDNA in K562 cells are summarized in Table S1. Collectively, of the eight retrons, Sa163 showed the highest msDNA production activity under the conditions tested (Figure 1C). While several other retrons had higher msDNA production than retron Ec86, we previously validated retron Ec86-mediated CRISPEY in yeast ^14^. These studies and our results suggested that both Sa163 and Ec86 possessed the potential to be explored as tools for precise gene editing and were selected for further study.

### Retron Ec86 and Sa163 enable HDR in both suspension and adherent human cell lines

To test if retrons can promote HDR, we used the reporter cell lines previously described in Richardson et al. ^22^ that used BFP-to-GFP conversions as editing readout. When HDR occurs, a three-nucleotide substitution converts the integrated BFP reporter into GFP (Figure 2A). We co-expressed the red fluorescence protein mCherry with Cas9 and RT in the reporter line and used the multicistronic retron plasmid to generate donors to convert BFP to GFP (Figure 2B). After inducing edits for each retron, we isolated transfected cells by flow cytometry to evaluate HDR. We used the BFP-GFP donor template to convert the protein expression from BFP to GFP. In K562 BFP reporter cells, controls without retron components had the baseline level of HDR at only 0.054% HDR events, whereas Ec86 had the highest level of HDR at 11.3% HDR events (Figure 2C). Sa163, the retron with the highest level of msDNA production (Figure 1C), had lower levels of HDR at 4.28% HDR events (Figure 2C).

**Figure 2.**
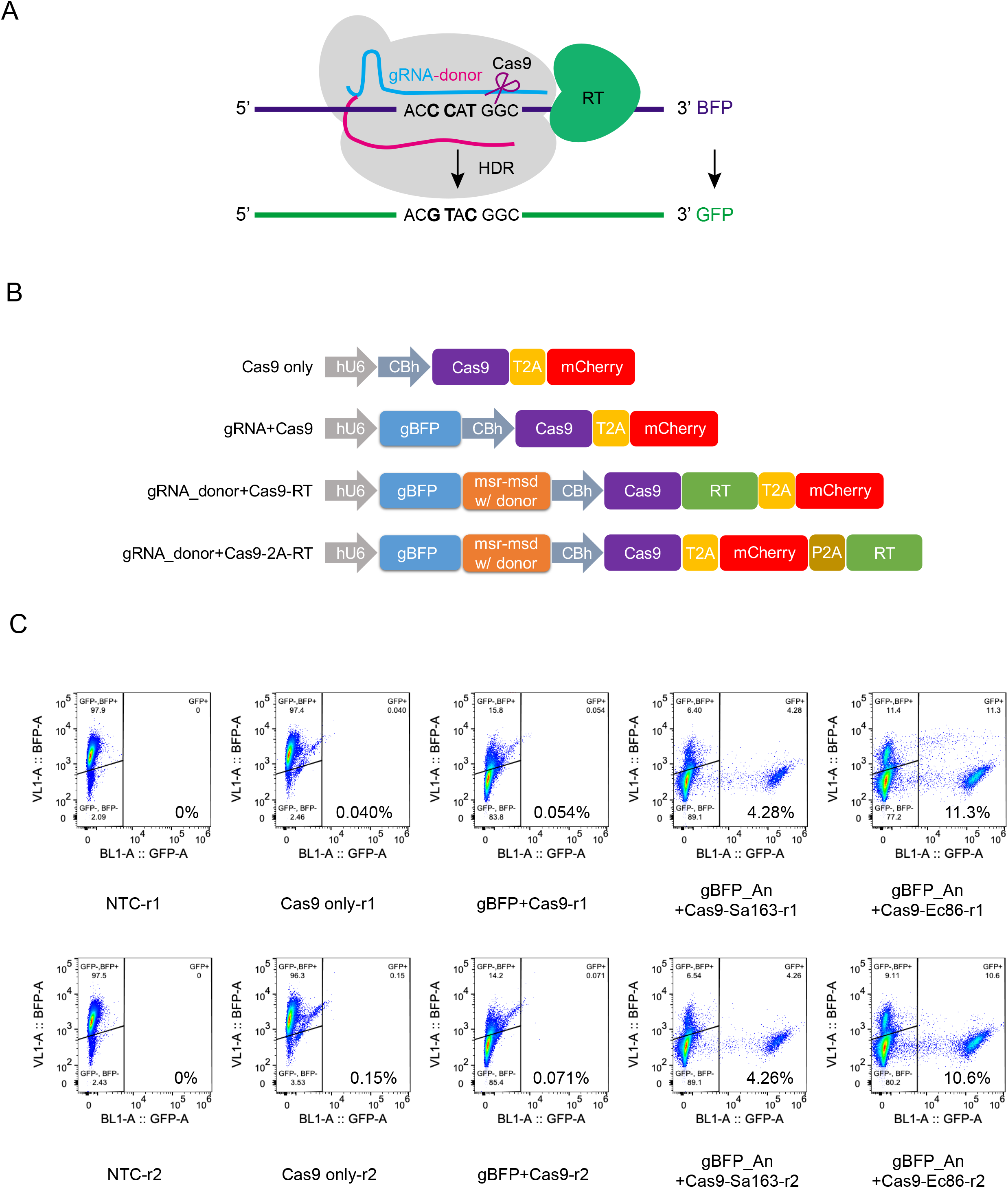
Retrons enable HDR in K562 and 293T BFP reporter cells. (A) Schematic showing the principle of BFP reporter cell line, adapted from Richardson et al. ^22^ (Figure 3a). (B) Schematic of plasmid design. (C) BFP-GFP scatter plots showing retron-enabled HDR rate detected in K562 BFP reporter cells by flow cytometry. Two replicate experiments are shown, r1: replicate 1; r2: replicate 2.

Since previous studies have shown that DNA donor length and strand type (e.g., target vs. non-target) influence editing ^22^, we tested three pairs of gRNA target- and non-target-strand donor templates (Supplemental Figure 2B). In general, higher HDR rates were detected when using donors that were complementary to the non-target strand (Supplemental Figure 2C-2E), which was consistent with previous studies showing that Cas9 first releases the non-target strand after cleavage ^22^. Although free Sa163 seemed to produce more than twofold msDNA than Cas9-Sa163 RT fusion (Cas9-2A-Sa163 vs. Cas9-Sa163, Figure 1C), HDR editing by Cas9-Sa163 fusion was 3.3-fold higher (gBFP-An+Cas9-Sa163 vs. gBFP-An+Cas9-2A-Sa163 = 4.28% : 1.29%, Supplemental Figure 2D). Having Sa163 RT fused to Cas9 may increase HDR by producing msDNA in the vicinity of the double-strand breaks (DSBs), even though the free Sa163 RT can produce more msDNAs that are globally distributed in the nucleus.

Notably, regardless of which retron was co-expressed with gRNA-Cas9 and which DNA template was used, CRISPR-Cas9 cutting efficiency in human cells was comparable to gRNA-Cas9 alone (Figure 2C, Supplemental Figure 2D and 2E). This finding suggests that our gRNA-msDNA hybrid did not severely impact the gRNA recognition and that our Cas9-RT fusion protein did not suppress Cas9 activity of DNA cleavage. Taken together, our findings indicated that retrons Ec86 and Sa163 can facilitate HDR in both K562 and HEK293T cells (Supplemental Figure 2C-2E).

## Discussion

Retrons are unique bacterial DNA elements that are capable of generating msDNA *in vivo* through reverse transcription. Recently, two independent groups have reported the role of retrons in antiphage defense in prokaryotes ^23, 24^. We hypothesized that retron-generated msDNA could be utilized to generate repair templates for precise genome editing in human cells. Here, we showed that: (1) retrons from different bacterial species have a wide range of RT activity in human cells and (2) simultaneous expression of retron RT with a hybrid retron RNA/sgRNA transcript can facilitate precise editing in HEK293 and K562 cells. Building on our previous study of retron Ec86 in yeast ^14^, our results suggest that both retron Ec86 and Sa163 may enable precise gene editing in human cells.

Increasing the supply of donor template at the editing site has been shown to improve the efficiency of HDR repair in yeast and mammalian cells ^25, 26^. The CRISPEY gRNA-retron design allows the sgRNA and msDNA to be covalently linked, which is intended to make the donor template immediately available for HDR repair at Cas9-induced DSBs. Further improvement of the retron RT processivity may generate more gRNA/msDNA hybrids available for recruitment or simply increase the donor template concentration in the nucleus to increase the probability of HDR over NHEJ. Similarly, the retron RNA scaffold can also be engineered to provide increased affinity for RT binding or activity. Both the retron RT and retron RNA can be engineered through directed evolution or knowledge-based enzyme variant design, such as that seen with group II intron RTs ^27^.

We have explored several experimentally validated retrons for msDNA production. Notably, Sa163 generated the most msDNA in K562 cells, as measured by qPCR (Figure 1C). However, Sa163 had a lower HDR rate when compared to Ec86 in the BFP-GFP reporter (Supplemental Figure 2C). This discrepancy between donor abundance and HDR rate raises the question of whether higher donor concentration increases HDR editing. One possible explanation for this discrepancy is post-transfection cell toxicity that may occur with high retron activity, as suggested by decreasing expression of mCherry in the cell population over time (not shown). Although Sa163 produced more msDNA, the increased toxicity may be contributing to reduced cellular viability in cells with higher RT and msDNA expression, resulting in fewer HDR-edited cells. Therefore, future work may survey retron species for efficient generation of msDNA with lower toxicity, which may also improve CRISPEY editing efficiency without the necessity of engineering of RT and retron RNA.

Many potential avenues exist for further optimization of CRISPEY in humans or other species. For example, single-strand annealing proteins (SSAPs) have been shown as effector proteins to improve retron-mediated editing in bacteria ^15, 16, 28^. More recently, there is evidence that SSAPs also improve efficiency of Cas9-mediated knock-ins in human cells ^29^. It is thus tempting to speculate that co-expression of SSAP may further improve CRISPEY editing efficiency.

Recent advances in base editors ^30-32^ and prime editors ^33^ have led to highly efficient editing rates. However, these methods are limited to short genetic alterations. In contrast, CRISPEY can efficiently insert gene-length fragments (e.g., GFP) in yeast ^14^. This approach may expand the length of potential knock-ins in human cells by circumventing the need to deliver long donor DNA molecules.

Beyond precise gene editing, there are other promising applications of retron activity in human cells for generating ssDNA. Generation of ssDNA is of great interest due to its use in biotechnology (e.g., DNA origami) ^34^, genome modification (e.g., intrachromosomal recombination) ^35^, and generation single stranded oligonucleotides that can fold into 3D structures that bind target molecules (i.e., aptamers) ^36, 37^.

## Conclusion

In this study, we harnessed the capability of retrons to generate intracellular gRNA-msDNA hybrid molecules and repurposed the products for CRISPR-Cas9-mediated, homology-directed repair in human cells. We presented a precise genome editing method that provides unique advantages in donor delivery, especially for repair templates that are difficult to deliver via conventional approaches. With further optimization, the msDNA generated by retrons has vast potential in precise gene editing and other biotechnology applications.

## Acknowledgments

We thank Dr. Jacob Corn (ETH Zürich) and Dr. Christopher D. Richardson (UCSB) for providing the HEK293T BFP and K562 BFP reporter cells as published in Richardson et al. (2016); Dr. Ralf Kühn’s group for providing the plasmids as published in Chu et al. (2015); Jing Bian, Yanyu Zhu, Dehua Zhao in Dr. Stanley Qi’s lab (Stanford) for providing access to the Neon transfection system (Invitrogen) and K562 wildtype cells; the Stanford Biology Department for providing access to the Attune NxT flow cytometer (Invitrogen). We thank all Fraser lab members, Dr. Michael Bassik, Josh Tycko, Cameron Lee, Gaelen Hess (Bassik lab, Stanford) and Xiaoshu Xu (Qi lab, Stanford) for helpful discussions.

---

**Supplemental Figure 1.**
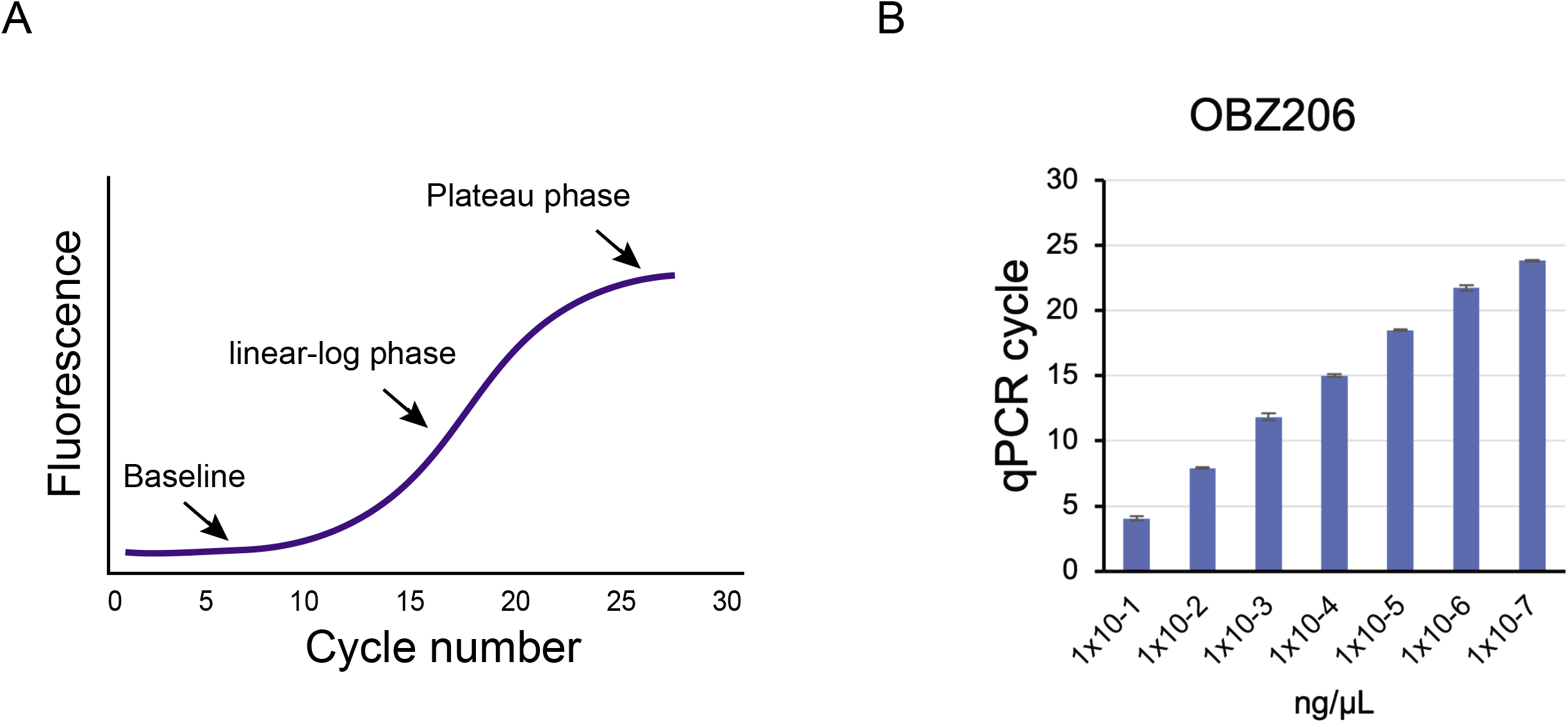

**Supplemental Figure 2.**
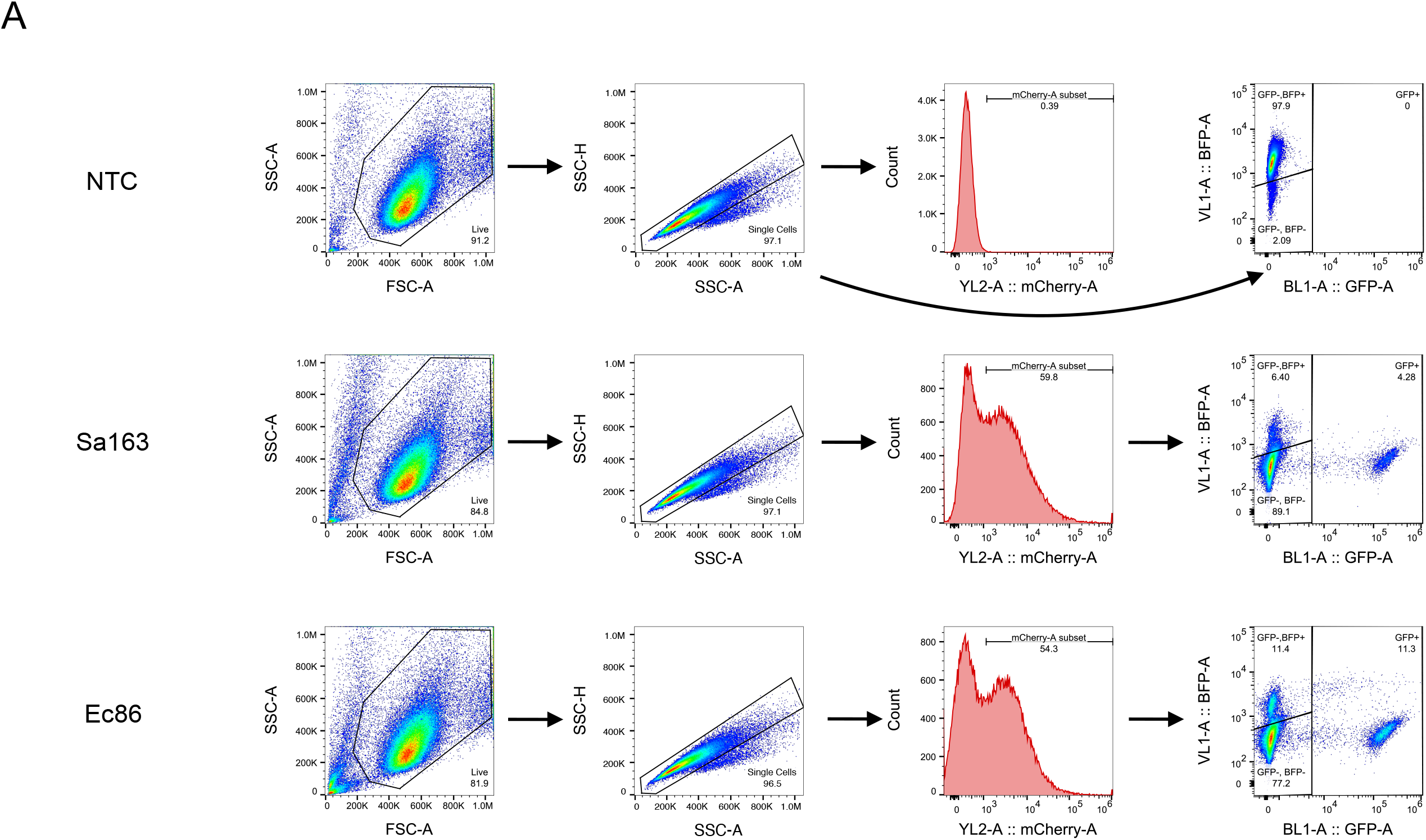
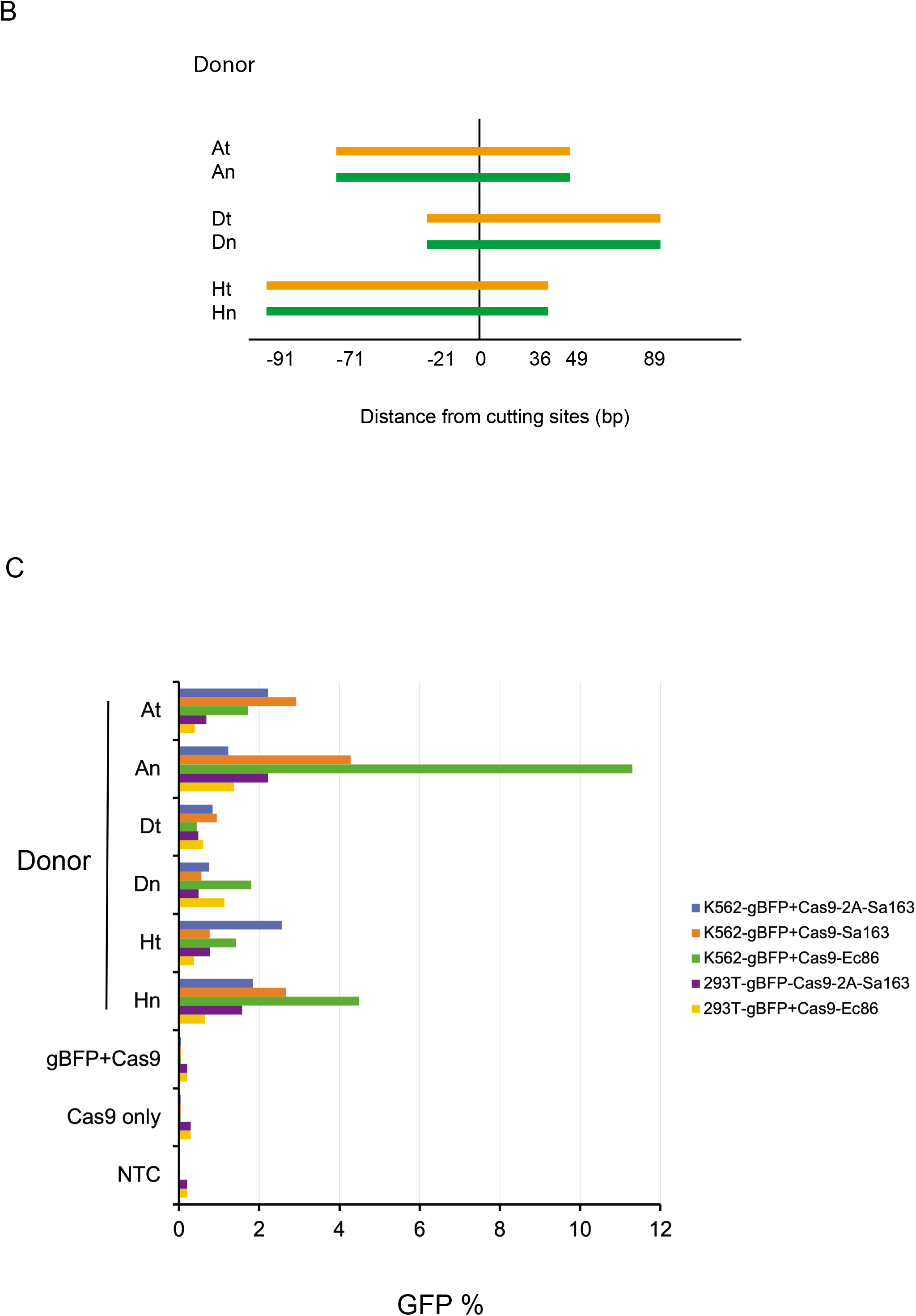
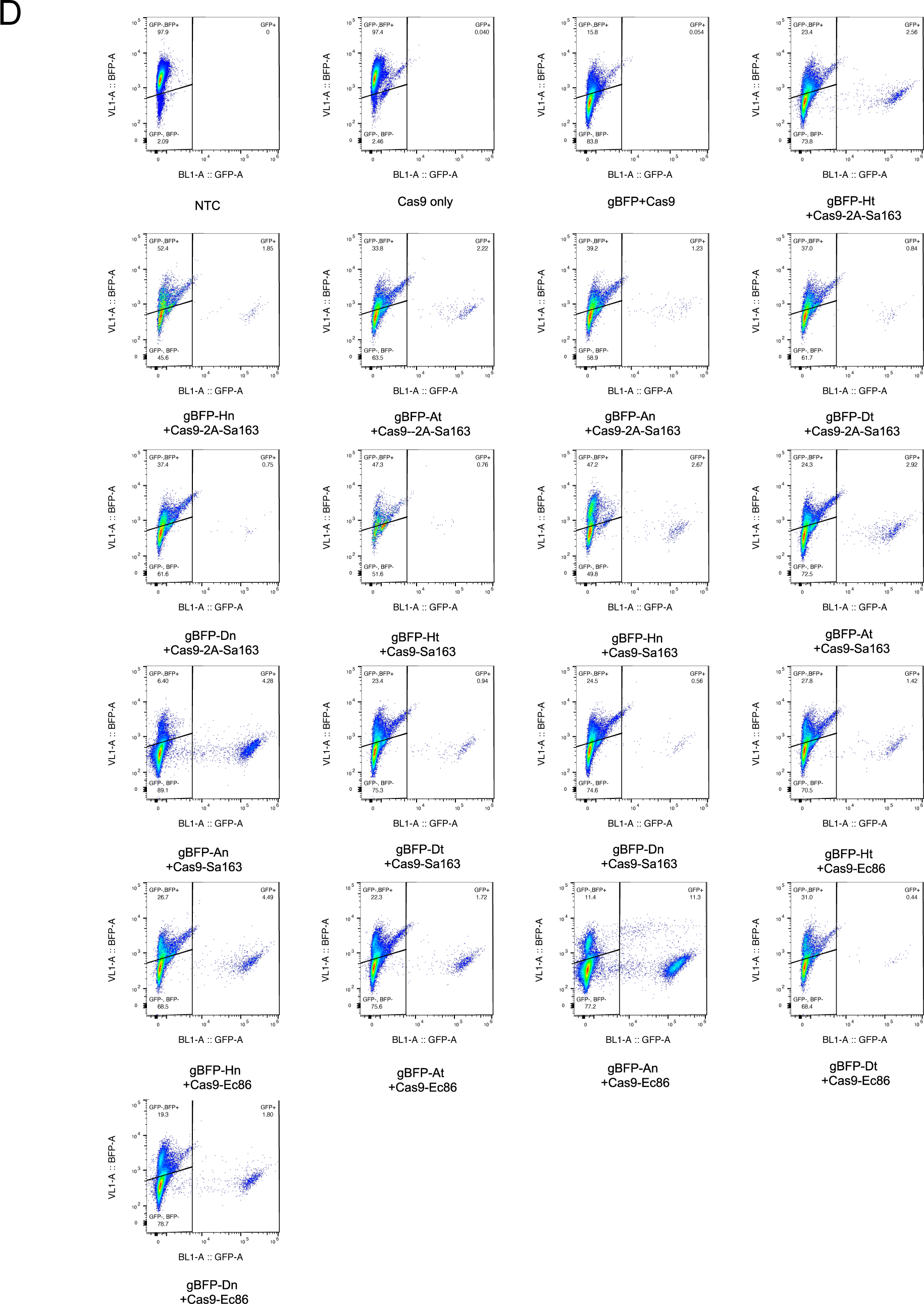
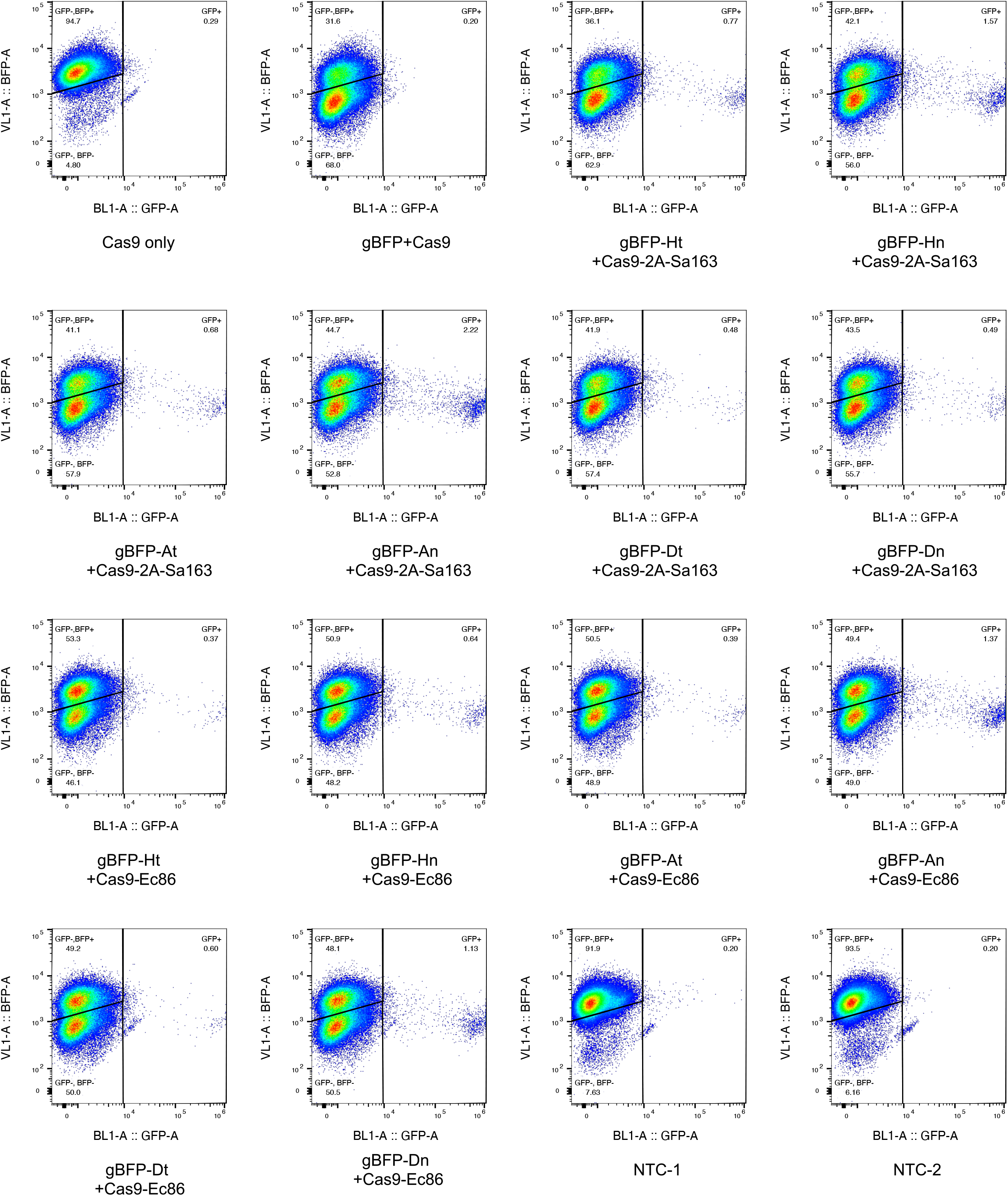

## References

1. Hsu PD, Lander ES, Zhang F. Development and applications of CRISPR-Cas9 for genome engineering. Cell. Jun 2014;157(6):1262–1278. doi:10.1016/j.cell.2014.05.010

2. Xiong X, Chen M, Lim WA, Zhao D, Qi LS. CRISPR/Cas9 for Human Genome Engineering and Disease Research. Annu Rev Genomics Hum Genet. 08 2016;17:131–54. doi:10.1146/annurev-genom-083115-022258

3. Certo MT, Ryu BY, Annis JE, et al. Tracking genome engineering outcome at individual DNA breakpoints. Nat Methods. Jul 2011;8(8):671–6. doi:10.1038/nmeth.1648

4. Yip BH. Recent Advances in CRISPR/Cas9 Delivery Strategies. Biomolecules. 05 2020;10(6)doi:10.3390/biom10060839

5. Lino CA, Harper JC, Carney JP, Timlin JA. Delivering CRISPR: a review of the challenges and approaches. Drug Deliv. Nov 2018;25(1):1234–1257. doi:10.1080/10717544.2018.1474964

6. Dever DP, Bak RO, Reinisch A, et al. CRISPR/Cas9 β-globin gene targeting in human haematopoietic stem cells. Nature. 11 2016;539(7629):384–389. doi:10.1038/nature20134

7. Sather BD, Romano Ibarra GS, Sommer K, et al. Efficient modification of CCR5 in primary human hematopoietic cells using a megaTAL nuclease and AAV donor template. Sci Transl Med. Sep 2015;7(307):307ra156. doi:10.1126/scitranslmed.aac5530

8. Wang J, Exline CM, DeClercq JJ, et al. Homology-driven genome editing in hematopoietic stem and progenitor cells using ZFN mRNA and AAV6 donors. Nat Biotechnol. Dec 2015;33(12):1256–1263. doi:10.1038/nbt.3408

9. Simon AJ, Ellington AD, Finkelstein IJ. Retrons and their applications in genome engineering. Nucleic Acids Res. 12 2019;47(21):11007–11019. doi:10.1093/nar/gkz865

10. Inouye S, Hsu MY, Eagle S, Inouye M. Reverse transcriptase associated with the biosynthesis of the branched RNA-linked msDNA in Myxococcus xanthus. Cell. Feb 1989;56(4):709–17. doi:10.1016/0092-8674(89)90593-x

11. Yee T, Furuichi T, Inouye S, Inouye M. Multicopy single-stranded DNA isolated from a gram-negative bacterium, Myxococcus xanthus. Cell. Aug 1984;38(1):203–9. doi:10.1016/0092-8674(84)90541-5

12. Shimamoto T, Hsu MY, Inouye S, Inouye M. Reverse transcriptases from bacterial retrons require specific secondary structures at the 5’-end of the template for the cDNA priming reaction. J Biol Chem. Feb 1993;268(4):2684–92.

13. Shimamoto T, Inouye M, Inouye S. The formation of the 2’,5’-phosphodiester linkage in the cDNA priming reaction by bacterial reverse transcriptase in a cell-free system. J Biol Chem. Jan 1995;270(2):581–8. doi:10.1074/jbc.270.2.581

14. Sharon E, Chen SA, Khosla NM, Smith JD, Pritchard JK, Fraser HB. Functional Genetic Variants Revealed by Massively Parallel Precise Genome Editing. Cell. 10 2018;175(2):544-557.e16. doi:10.1016/j.cell.2018.08.057

15. Schubert MG, Goodman DB, Wannier TM, et al. High throughput functional variant screens via in-vivo production of single-stranded DNA. bioRxiv. 2020:2020.03.05.975441. doi:10.1101/2020.03.05.975441

16. Farzadfard F, Lu TK. Synthetic biology. Genomically encoded analog memory with precise in vivo DNA writing in living cell populations. Science. Nov 2014;346(6211):1256272. doi:10.1126/science.1256272

17. Mirochnitchenko O, Inouye S, Inouye M. Production of single-stranded DNA in mammalian cells by means of a bacterial retron. J Biol Chem. Jan 1994;269(4):2380–3.

18. Chu VT, Weber T, Wefers B, et al. Increasing the efficiency of homology-directed repair for CRISPR-Cas9-induced precise gene editing in mammalian cells. Nat Biotechnol. May 2015;33(5):543–8. doi:10.1038/nbt.3198

19. Gilbert LA, Horlbeck MA, Adamson B, et al. Genome-Scale CRISPR-Mediated Control of Gene Repression and Activation. Cell. Oct 2014;159(3):647–61. doi:10.1016/j.cell.2014.09.029

20. Gray SJ, Foti SB, Schwartz JW, et al. Optimizing promoters for recombinant adeno-associated virus-mediated gene expression in the peripheral and central nervous system using self-complementary vectors. Hum Gene Ther. Sep 2011;22(9):1143–53. doi:10.1089/hum.2010.245

21. Ibrahimi A, Vande Velde G, Reumers V, et al. Highly efficient multicistronic lentiviral vectors with peptide 2A sequences. Hum Gene Ther. Aug 2009;20(8):845–60. doi:10.1089/hum.2008.188

22. Richardson CD, Ray GJ, DeWitt MA, Curie GL, Corn JE. Enhancing homology-directed genome editing by catalytically active and inactive CRISPR-Cas9 using asymmetric donor DNA. Nat Biotechnol. Mar 2016;34(3):339–44. doi:10.1038/nbt.3481

23. Millman A, Bernheim A, Stokar-Avihail A, et al. Bacterial Retrons Function In Anti-Phage Defense. Cell. Dec 2020;183(6):1551-1561.e12. doi:10.1016/j.cell.2020.09.065

24. Gao L, Altae-Tran H, Böhning F, et al. Diverse enzymatic activities mediate antiviral immunity in prokaryotes. Science. 08 2020;369(6507):1077–1084. doi:10.1126/science.aba0372

25. Lee K, Mackley VA, Rao A, et al. Synthetically modified guide RNA and donor DNA are a versatile platform for CRISPR-Cas9 engineering. Elife. 05 2017;6doi:10.7554/eLife.25312

26. Roy KR, Smith JD, Vonesch SC, et al. Multiplexed precision genome editing with trackable genomic barcodes in yeast. Nat Biotechnol. 07 2018;36(6):512–520. doi:10.1038/nbt.4137

27. Zhao C, Liu F, Pyle AM. An ultraprocessive, accurate reverse transcriptase encoded by a metazoan group II intron. RNA. 02 2018;24(2):183–195. doi:10.1261/rna.063479.117

28. Wannier TM, Ciaccia PN, Ellington AD, et al. Recombineering and MAGE. Nature Reviews Methods Primers. 2021/01/14 2021;1(1):7. doi:10.1038/s43586-020-00006-x

29. Wang C, Cheng JKW, Zhang Q, et al. Microbial single-strand annealing proteins enable CRISPR gene-editing tools with improved knock-in efficiencies and reduced off-target effects. Nucleic Acids Res. Feb 2021;doi:10.1093/nar/gkaa1264

30. Gaudelli NM, Komor AC, Rees HA, et al. Programmable base editing of A•T to G•C in genomic DNA without DNA cleavage. Nature. 11 2017;551(7681):464–471. doi:10.1038/nature24644

31. Rees HA, Liu DR. Base editing: precision chemistry on the genome and transcriptome of living cells. Nat Rev Genet. 12 2018;19(12):770–788. doi:10.1038/s41576-018-0059-1

32. Komor AC, Kim YB, Packer MS, Zuris JA, Liu DR. Programmable editing of a target base in genomic DNA without double-stranded DNA cleavage. Nature. 05 2016;533(7603):420–4. doi:10.1038/nature17946

33. Anzalone AV, Randolph PB, Davis JR, et al. Search-and-replace genome editing without double-strand breaks or donor DNA. Nature. 12 2019;576(7785):149–157. doi:10.1038/s41586-019-1711-4

34. Rothemund PW. Folding DNA to create nanoscale shapes and patterns. Nature. Mar 2006;440(7082):297–302. doi:10.1038/nature04586

35. Datta HJ, Glazer PM. Intracellular generation of single-stranded DNA for chromosomal triplex formation and induced recombination. Nucleic Acids Res. Dec 2001;29(24):5140–7. doi:10.1093/nar/29.24.5140

36. Keefe AD, Pai S, Ellington A. Aptamers as therapeutics. Nat Rev Drug Discov. 07 2010;9(7):537–50. doi:10.1038/nrd3141

37. Lee JF, Stovall GM, Ellington AD. Aptamer therapeutics advance. Curr Opin Chem Biol. Jun 2006;10(3):282–9. doi:10.1016/j.cbpa.2006.03.015

