## Supplemental Information for "Bacterial retrons enable precise gene editing in human cells"

[2] Corresponding Author

This PDF file includes:

Supplemental Figure Legends

Supplemental Note 1

Supplemental Table 1

Supplemental Note 2

Supplemental References

### Supplemental Figure Legends

**Supplemental Figure 1**. (A) A typical amplification curve generated in qPCR assay. During the linear-log phase, PCR products of the target gene approximately double in each cycle. Amplification stops at the plateau phase. (B) The synthesized DNA oligo OBZ206 was used as a "ladder" to indicate the qPCR linear-log phase in this test. Data presented as mean ± s.d. (n=3).

**Supplemental Figure 2**. All FACS plots related to Figure 2. (A) Gating strategy to detect the HDR rate as shown in Figure 2C. From left to right, all cells were first gated for size by forward scatter area (FSC-A) and side scatter area (SSC-A); single cells were further selected by side scatter area (SSC-A) and side scatter height (SSC-H); then cells transfected with retron-CRISPR plasmids were determined by YL2-A (mCherry-A); subsequently, BFP to GFP conversion rates were measured by VL-A(BFP-A) and BL-A(GFP-A). Since the mCherry gene was carried by a plasmid, for non-transfected control (NTC) cells, the BFP to GFP conversion rate was measured among all single cell populations. (B) Schematic of the three pairs of target strand (At, Dt, Ht) and non-target strand (An, Dn, Hn) donor templates tested, adapted from Richardson et al. (Figure 3c) [1]. (C) The graph summarizes the HDR percentages achieved by Ec86 and Sa163 among variable donor templates. (D) All FACS plots tested in K562 BFP reporter cells. (E) All FACS plots tested in HEK293T BFP reporter cells.

### Supplemental Note 1

**qPCR standard template--- the single-stranded oligodeoxynucleotides (ssODNs) sequence**

OBZ206:

GACGACGGCAACTACAAGACCCGCGCCGAGGTGAAGTTCGAGGGCGACACCCTG
GTGAACCGCATCGAGCTGAAGGGCATCGACTTCAAGGAGGACGGCAACATCCTG
GGGCACAAGCTGGAGTACAACTACAACAGCCACAACGTCTATATCACCGCCGAC
AAGCAGAAGAACGGCATCAAGGCCAACTTCAAGATCCG

**qPCR primers:**

Forward primer: OBZ152 AGCTGAAGGGCATCGACTT

Reverse primer: OBZ153 GGATCTTGAAGTTGGCCTTG

**Donor sequence**:

TCTTCAAGGACGACGGCAACTACAAGACCCGCGCCGAGGTGAAGTTCGAGGGCG
ACACCCTGGTGAACCGCATCGAGCTGAAGGGCATCGACTTCAAGGAGGACGGCA
ACATCCTGGGGCACAAGCTGGAGTACAACTACAACAGCCACAACGTCTATATCA
CCGCCGACAAGCAGAAGAACGGCATCAAGGCCAACTTCAAGATCCG

Donor sequence used in qPCR assay was amplified from the TLR repair vector in Chu et al. [2].

**qPCR conditions**

| **Component** | **Volume (μl)** |
|---|---|

| | |
|---|---|
| Total | 10 |
| H2O | 3.4 |
| iQTM SYBR® Green supermix (2x) | 5 |
| OBZ152+OBZ153 (5µM each) | 0.6 |
| extracted ssDNA | 1 |

**qPCR Cycling steps**

| Temperature (°C) | Time | Cycles |
|---|---|---|
| 95 | 3 min | 1 |
| 95 | 10 s | 40 |
| 55 | 30 s | |

**Supplemental Table 1. Retrons used in qPCR assay**

| Name | msr-msd sequence | RT (amino acid sequence) |
|---|---|---|
| Ec86 | ATGCGCACCCTTAGCG AGAGGTTTATCATTAA GGTCAACCTCTGGATG TTGTTTCGGCATCCTGC ATTGAATCTGAGTTACT GTCTGTTTcCCTAGGGA ACCCGTTTCTTCTGACG TAAGGGTGCGCA | MKSAEYLNTFRLRNLGLPVMNNLHDMSKAT RISVETLRLLIYTADFRYRIYTVEKKGPEKRMR TIYQPSRELKALQGWVLRNILDKLSSSPFSIGFE KHQSILNNATPHIGANFILNIDLEDFFPSLTANK VFGVFHSLGYNRLISSVLTKICCYKNLLPQGAP SSPKLANLICSKLDYRIQGYAGSRGLIYTRYAD DLTLSAQSMKKVVKARDFLFSIIPSEGLVINSK KTCISGPRSQRKVTGLVISQEKVGIGREKYKEI RAKIHHIFCGKSSEIEHVRGWLSFILSVDSKSH RRLITYISKLEKKYGKNPLNKAKT |
| Ec107 | CGCCAGCAGTGGCAAT AGCGTTTCCGGCCTTTT GTGCCGGGAGGGTCGG CGAGTCGCTGACTTAA CGCCAGTAGTATGTCC ATATACCCAAAGTCGC TTCATTGTACCTGAGTA CGCTTCGCGTACGTCG CGCTGACGCGCTCAGT ACAGTTACGCGCCTTC GGGATGGTTTAATGGT ATTGCCGCTGTTGGCG | MDATRTTLLALDLFGSPGWSADKEIQRLHALS NHAGRHYRRIILSKRHGGQRLVLAPDYLLKTV QRNILKNVLSQFPLSPFATAYRPGCPIVSNAQP HCQQPQILKLDIENFFDSISWLQVWRVFRQAQ LPRNVVTMLTWICCYNDALPQGAPTSPAISNL VMRRFDERIGEWCQARGITYTRYCDDMTFSG HFNARQVKNKVCGLLAELGLSLNKRKGCLIA ACKRQQVTGIVVNHKPQLAREARRALRQEVH LCQKYGVISHLSHRGELDPSGDLHAQATAYL YALQGRINWLLQINPEDEAFQQARESVKRML VAW |

| | | |
|---|---|---|
| Mx65 | ggatccGGGCTCGCAGAT<br>GAGCCATGAGTACCGC<br>GGTGTTTCGCCGCGGG<br>GGTGTTCTGTCCCCATC<br>TCTTCGCCAGGGTCCC<br>AGCGTACGCAACGCAG<br>GGAGCCCCGGGTCCAA<br>CGCCTCGCAGGTCGTC<br>CCCTGGCCT | MSWFDTTLSRLKGLFSRPVTRSTTGLDVPLDA<br>HGRPQDVVTETVSTSGPLKPGHLRQVRRDAR<br>LLPKGVRRYTPGRKKWMEAAEARRLFSATLR<br>TRNRNLRDLLPDEAQLARYGLPVWRTEEDVA<br>AALGVSVGVLRHYSIHRPRERVRHYVTFAVP<br>KRSGGVRLLHAPKRRLKALQRRMLALLVSKL<br>PVSPQAHGFVPGRSIKTGAAPHVGRRVVLKLD<br>LKDFFPSVTFARVRGLLIALGYGYPVAATLAV<br>LMTESERQPVELEGILFHVPVGPRVCVQGAPT<br>SPALCNAVLLRLDRRLAGLARRYGYTYTRYA<br>DDLTFSGDDVTALERVRALAARYVQEEGFEV<br>NREKTRVQRRGGAQRVTGVTVNTTLGLSREE<br>RPRLRAMLHQEARSEDVEAHRAHLDGLLAYV<br>KMLNPEQAERLARRRKPRGT |
| Mx162 | GCGCGAGCAGCCGAGA<br>GAGGTCCGGAGTGCAT<br>CAGCCTGAGCGCCTCG<br>AGCGGCGGAGCGGCGT<br>TGCGCCGCTCCGGTTG<br>GAATGCAGGACACTCT<br>CCGCAAGGTAGCCTGT<br>TCTTGGCTCTCTCCCTC<br>CTAGGCACTACGGCCA<br>GGGTGGGTAGCGGAGC<br>CAACGACGCGACCGCC<br>GTTTACCCACCCCGGC<br>CGTAGTGCCTAGGAGG<br>GGAGAGCCGGTGAGGC<br>TACCGTGCCCC | MTARLDPFVPAASPQAVPTPELTAPSSDAAAK<br>REARRLAHEALLVRAKAIDEAGGADDWVQA<br>QLVSKGLAVEDLDFSSASEKDKKAWKEKKKA<br>EATERRALKRQAHEAWKATHVGHLGAGVH<br>WAEDRLADAFDVPHREERARANGLTELDSAE<br>ALAKALGLSVSKLRWFAFHREVDTATHYVSW<br>TIPKRDGSKRTITSPKPELKAAQRWVLSNVVE<br>RLPVHGAAHGFVAGRSILTNALAHQGADVVV<br>KVDLKDFFPSVTWRRVKGLLRKGGLREGTST<br>LLSLLSTEAPREAVQFRGKLLHVAKGPRALPQ<br>GAPTSPGITNALCLKLDKRLSALAKRLGFTYT<br>RYADDLTFSWTKAKQPKPRRTQRPPVAVLLS<br>RVQEVVEAEGFRVHPDKTRVARKGTRQRVTG<br>LVVNAAGKDAPAARVPRDVVRQLRAAIHNR<br>KKGKPGREGESLEQLKGMAAFIHMTDPAKGR<br>AFLAQLTELESTASAAPQAE |
| St85 | ACTCTTTAGCGTTAGGC<br>TTTGATTTATAGCCTTG<br>TCGAGCGTTTCGCCAG<br>ACACTAACTTATTGAG<br>TACTTTTAGGGTTGCGC<br>TAGAAAGTTTTCTACC<br>GATCCTAGAAGTCTCT<br>AGGATCGGTAGAAAAC<br>TTTCTAGCGCCTCCTCT<br>AGTAAAGAGT | MDILQHISDLLLTKKSEIISFSLTAPYRYKIYKI<br>AKRNSDKKRTIAHPSKELKFIQREITEYLTDKL<br>PVHECAFAYKKGSSIKTNAQVHLHTKYLLKM<br>DFENFFPSITPRLFFSKLRLANIDLTADDKVLLE<br>NILFFKSKRNSNLRLSIGAPSSPLISNFVMYFW<br>DIEVQEICSKIGVNYTRYADDLTFSTNNKDVLF<br>DIPDMLENVLPKYSLGRIRINHEKTVFSSKGHN<br>RHVTGITLTNDNKLSIGRERKRKISAMIHHFIN<br>GKLSTDECNKLVGLLAFAKNIEPSFYKSMVIK<br>YGSDNIYKLQKQKDK |

| | | |
|---|---|---|
| Ec67 | CACGCATGTAGGCAGA<br>TTTGTTGGTTGTGAATC<br>GCAACCAGTGGCCTTA<br>ATGGCAGGAGGAATCG<br>CCTCCCTAAAATCCTTG<br>ATTCAGAGCTATACGG<br>CAGGTGTGCTGTGCGA<br>AGGAGTGCCTGCATGC<br>GT | MTKTSKLDALRAATSREDLAKILDIKLVFLTN<br>VLYRIGSDNQYTQFTIPKKGKGVRTISAPTDRL<br>KDIQRRICDLLSDCRDEIFAIRKISNNYSFGFER<br>GKSIILNAYKHRGKQIILNIDLKDFFESFNFGRV<br>RGYFLSNQDFLLNPVVATTLAKAACYNGTLP<br>QGSPCSPIISNLICNIMDMRLAKLAKKYGCTYS<br>RYADDITISTNKNTFPLEMATVQPEGVVLGKV<br>LVKEIENSGFEINDSKTRLTYKTSRQEVTGLTV<br>NRIVNIDRCYYKKTRALAHALYRTGEYKVPD<br>ENGVLVSGGLDKLEGMFGFIDQVDKFNNIKK<br>KLNKQPDRYVLTNATLHGFKLKLNAREKAYS<br>KFIYYKFFHGNTCPTIITEGKTDRIYLKAALHS<br>LETSYPELFREKTDSKKKEINLNIFKSNEKTKY<br>FLDLSGGTADLKKFVERYKNNYASYYGSVPK<br>QPVIMVLDNDTGPSDLLNFLRNKVKSCPDDVT<br>EMRKMKYIHVFYNLYIVLTPLSPSGEQTSMED<br>LFPKDILDIKIDGKKFNKNNDGDSKTEYGKHIF<br>SMRVVRDKKRKIDFKAFCCIFDAIKDIKEHYK<br>LMLNS |
| Sa163 | GTGCGAGCGACCGAGA<br>GAGGTCCCAAGCCATC<br>AGCCTCAGCGCCTCGA<br>GCGCGAGAGCGGCGTT<br>GCGCCGCTCTGGTTGA<br>ATTGCAGGACACTCTC<br>CGCAAGGTAGCCTGTT<br>CTTGGCTCTCTTCCCTC<br>CGGTGAGTACCTCTCC<br>GGCCGGGGAGCTGAAC<br>CAACGACGCAACCGCC<br>GTTTCCCCGGCCGGAG<br>AGGTACTCACCGGAGG<br>GGAGAGCCGGTGAGGC<br>TACCGTGCCCCAGGTG<br>AGAAGGTGGTGCCTTC<br>GGGCCTCCCTCGACCG<br>CTCGCGC | MTAKLESHVPAAPPVSAEAPAPTRPDAAKQE<br>ARRAHHEALRLRWKAIEEAGGTDAWVRQQL<br>VAKGVAAEEVDFESLSDKQKAAWKEKKKAE<br>ATERRAQKRLAWEAWKATHIHHLGVGVHWD<br>EAGGPDKFDVAGREERAKANGLPEGLDSVEA<br>LAKALGISVSRLRWFSFHREVDTGTHYQTWEI<br>PKRDGGKRTLTAPKRELKAVQRWVLANVVE<br>RLPVHGAAHGFVAGRSILTNALAHQGADVVV<br>KVDMKDFFPSVTWPRVKGLLRKGGLPENLAT<br>LLALLSTEAPREVVRFRGETLYVAKGPRALPQ<br>GAPTSPALTNALCLRLDKRLSALSKRLGFTYT<br>RYADDLTFSWRRAKKSRQKELPLADAPVALL<br>LARVKGVLEAEGFTLHPDKTRVQRKGSRQRV<br>TGLVVNEAPEGVPGARVPRDVVRRLRAAIHN<br>REQGKPGPTGETLEQLKGLAAFLHMTDAEKG<br>RAFLRRLEALEKRQTA |

| | | |
|---|---|---|
| Ec73 | AGAGCCAAACCTAGCA<br>TTTATGGGTTAATAGCC<br>CATCGGGCCATGAGTC<br>ATGGTTTCGCCTAGTAT<br>TTTAGCTATGCCCGTCG<br>TTATCGACGTGCTCAA<br>GTAGGTTTGGCTCT | MTKTSKLDALRAATSREDLAKILDIKLVFLTN<br>VLYRIGSDNQYTQFTIPKKGKGVRTISAPTDRL<br>KDIQRRICDLLSDCRDEIFAIRKISNNYSFGFER<br>GKSIILNAYKHRGKQIILNIDLKDFFESFNFGRV<br>RGYFLSNQDFLLNPVVATTLAKAACYNGTLP<br>QGSPCSPIISNLICNIMDMRLAKLAKKYGCTYS<br>RYADDITISTNKNTFPLEMATVQPEGVVLGKV<br>LVKEIENSGFEINDSKTRLTYKTSRQEVTGLTV<br>NRIVNIDRCYYKKTRALAHALYRTGEYKVPD<br>ENGVLVSGGLDKLEGMFGFIDQVDKFNNIKK<br>KLNKQPDRYVLTNATLHGFKLKLNAREKAYS<br>KFIYYKFFHGNTCPTIITEGKTDRIYLKAALHS<br>LETSYPELFREKTDSKKKEINLNIFKSNEKTKY<br>FLDLSGGTADLKKFVERYKNNYASYYGSVPK<br>QPVIMVLDNDTGPSDLLNFLRNKVKSCPDDVT<br>EMRKMKYIHVFYNLYIVLTPLSPSGEQTSMED<br>LFPKDILDIKIDGKKFNKNNDGDSKTEYGKHIF<br>SMRVVRDKKRKIDFKAFCCIFDAIKDIKEHYK<br>LMLNS |

Gene synthesis from Twist:

RT-EcoI/Ec86 in pTwist CMV:

ATGTACCCATACGACGTCCCAGACTACGCTCCTCCCAAGAAGAAAAGGAAAGTT
AAGAGTGCGGAATACCTTAACACATTTAGGCTTAGAAACCTGGGACTCCCCGTTA
TGAATAACCTGCACGATATGTCTAAGGCAACCCGAATCAGTGTAGAGACGCTTA
GACTCTTGATATATACCGCCGACTTTCGATACAGAATATATACCGTCGAGAAGAA
GGGCCCAGAGAAACGAATGCGGACCATATACCAACCTAGTAGGGAGCTGAAAG
CGCTTCAGGGCTGGGTACTTCGAAATATCCTTGACAAACTTTCCAGTTCACCGTT
CAGCATTGGTTTCGAGAAACACCAGAGTATTCTGAATAACGCGACACCTCACAT
AGGAGCCAATTTCATCCTCAATATTGACCTGGAGGATTTCTTCCCTAGCCTTACT
GCCAATAAAGTGTTCGGTGTATTCCACAGCCTCGGCTACAACCGACTGATTTCAT
CTGTACTCACAAAAATTTGTTGCTATAAGAACCTGCTCCCTCAAGGGGCCCCAAG
TAGCCCAAAACTGGCAAACCTCATCTGTTCAAAATTGGACTATCGAATTCAGGGC
TATGCCGGATCCAGGGGCTTGATCTACACGAGATACGCAGACGACCTGACACTTT
CAGCACAATCCATGAAGAAAGTTGTTAAAGCGAGGGATTTTCTTTTTTCCATTAT
TCCGTCTGAAGGATTGGTTATTAATTCTAAAAAAACTTGCATTAGTGGTCCTCGG
TCTCAACGAAAGGTTACAGGCCTGGTAATCTCTCAGGAGAAGGTTGGGATTGGA
AGAGAGAAGTATAAAGAGATTCGCGCCAAAATACATCATATTTTTTGCGGTAAA
TCATCTGAAATCGAGCACGTAAGGGGATGGCTTTCTTTCATTCTTTCTGTAGACTC
CAAATCTCACCGACGACTCATCACATATATAAGTAAACTGGAGAAAAAATATGG
GAAAAATCCGCTTAACAAGGCTAAAACTAAAAGGCCGGCGGCCACGAAAAAGG
CCGGCCAGGCAAAAAAGAAAAAGGACTATAAAGATGACGATGACAAGGACTAC
AAGGATGATGACGATAAAGATTATAAAGACGACGATGATAAGAAAGGTCCGGG
AAGCGGAGCTACTAACTTCAGCCTGCTGAAGCAGGCTGGAGACGTGGAGGAGAA
CCCTGGACCTCCGGTCGCCACCAGCGAGCTGATTAAGGAGAACATGCACATGAA
GCTGTACATGGAGGGCACCGTGGACAACCATCACTTCAAGTGCACATCCGAGGG

CGAAGGCAAGCCCTACGAGGGCACCCAGACCATGAGAATCAAGGTGGTCGAGG
GCGGCCCTCTCCCCTTCGCCTTCGACATCCTGGCTACTAGCTTCCTCTACGGCAGC
AAGACCTTCATCAACCACACCCAGGGCATCCCCGACTTCTTCAAGCAGTCCTTCC
CTGAGGGCTTCACATGGGAGAGAGTCACCACATACGAAGACGGGGGCGTGCTGA
CCGCTACCCAGGACACCAGCCTCCAGGACGGCTGCCTCATCTACAACGTCAAGA
TCAGAGGGGTGAACTTCACATCCAACGGCCCTGTGATGCAGAAGAAAACACTCG
GCTGGGAGGCCTTCACCGAGACGCTGTACCCCGCTGACGGCGGCCTGGAAGGCA
GAAACGACATGGCCCTGAAGCTCGTGGGCGGGAGCCATCTGATCGCAAACATCA
AGACCACATATAGATCCAAGAAACCCGCTAAGAACCTCAAGATGCCTGGCGTCT
ACTATGTGGACTACAGACTGGAAAGAATCAAGGAGGCCAACAACGAGACCTACG
TCGAGCAGCACGAGGTGGCAGTGGCCAGATACTGCGACCTCCCTAGCAAACTGG
GGCACAAGCTTAATTAATTAAGAATTCGTCGAGGGACCTAATAACTTCGTATAGC
ATACATTATACGAAGTTATACATGTTTAAGGGTTCCGGTTCCACTAGGTACAATT
CGATATCAAGCTTATCGATAATCAACCTCTGGATTACAAAATTTGTGAAAGATTG
ACTGGTATTCTTAACTATGTTGCTCCTTTTACGCTATGTGGATACGCTGCTTTAAT
GCCTTTGTATCATGCTATTGCTTCCCGTATGGCTTTCATTTTCTCCTCCTTGTATAA
ATCCTGGTTGCTGTCTCTTTATGAGGAGTTGTGGCCCGTTGTCAGGCAACGTGGC
GTGGTGTGCACTGTGTTTGCTGACGCAACCCCCACTGGTTGGGGCATTGCCACCA
CCTGTCAGCTCCTTTCCGGGACTTTCGCTTTCCCCCTCCCTATTGCCACGGCGGAA
CTCATCGCCGCCTGCCTTGCCCGCTGCTGGACAGGGGCTCGGCTGTTGGGCACTG
ACAATTCCGTGGTGTTGTCGGGGAAATCATCGTCCTTTCCTTGGCTGCTCGCCTGT
GTTGCCACCTGGATTCTGCGCGGGACGTCCTTCTGCTACGTCCCTTCGGCCCTCA
ATCCAGCGGACCTTCCTTCCCGCGGCCTGCTGCCGGCTCTGCGGCCTCTTCCGCG
TCTTCGCCTTCGCCCTCAGACGAGTCGGATCTCCCTTTGGGCCGCCTCCCCGCAG
CGGCCGCGAGGTGCTTGTAGATAACCTCCACGATGGTGCACCTTGGGCAACACA
AAAGTGGCAAATCATCTACAATGCGCACCCTTAGCGAGAGGTTTATCATTAAGGT
CAACCTCTGGATGTTGTTTCGGCATCCTGCATTGAATCTGAGTTACTGTCTGTTTT
CCTGGCCATGCTGGCCAGCGGATAACAAGTTTAAACACGGCCACAGTGGCCAGG
AAACCCGTTTCTTCTGACGTAAGGGTGCGCAGACACTGAGTGAGAAACGTCCCC
GTCGTAGTGTCGGTAATGCGTTGTTTCAACGTAGCCAATTCTCACGGCGCGCC

RT-EcoII/Ec67 in pTwist EF1 Alpha Puro:

ATGTACCCATACGACGTCCCAGACTACGCTCCTCCCAAGAAGAAAAGGAAAGTT
ATGACTAAAACCTCTAAACTCGACGCACTCAGAGCAGCAACATCCCGCGAAGAC
CTCGCCAAAATTTTGGATATTAAACTTGTGTTCCTTACAAATGTCCTGTATCGCAT
AGGGTCAGATAATCAATACACACAGTTTACGATCCCGAAGAAGGGTAAGGGCGT
GCGAACCATATCAGCCCCGACCGACCGGCTGAAGGACATTCAGCGGAGAATCTG
TGATTTGTTGAGCGACTGTCGGGATGAAATCTTTGCGATTCGGAAGATAAGTAAT
AATTATTCTTTCGGATTTGAAAGGGGGAAATCCATAATTCTCAATGCCTATAAAC
ACCGCGGTAAGCAGATTATCCTTAACATTGATCTTAAAGATTTCTTTGAGTCCTTC
AACTTCGGTAGGGTACGAGGTTACTTTCTTTCTAACCAGGATTTTTTGCTCAATCC
TGTTGTGGCGACTACACTGGCAAAAGCGGCATGCTATAATGGACCCTGCCTCA
GGGCAGCCCTTGCAGCCCAATAATATCAAACCTTATCTGCAACATCATGGATATG
CGCTTGGCAAAGCTGGCCAAGAAATACGGATGTACATACAGTCGGTACGCCGAT
GATATCACCATCTCTACAAATAAGAATACTTTTCCTCTCGAAATGGCCACAGTTC

AACCGGAGGGCGTCGTCCTGGGGAAGGTGCTCGTAAAAGAAATCGAAAACTCTG
GTTTCGAGATCAATGATTCTAAGACGAGACTGACTTATAAAACTAGTAGACAGG
AGGTTACGGGGCTGACCGTAAACCGCATAGTAAATATCGATCGATGCTACTATA
AGAAAACAAGGGCGCTTGCACACGCACTTTATCGGACAGGTGAATATAAGGTCC
CTGACGAGAACGGCGTCCTGGTTTCAGGCGGGCTGGATAAGCTCGAAGGAATGT
TTGGCTTTATCGATCAAGTCGACAAATTCAACAATATAAAGAAAAAGCTTAACA
AACAGCCAGATAGGTACGTCTTGACAAACGCCACACTGCATGGGTTCAAGCTGA
AACTCAACGCTCGCGAGAAGGCGTACTCAAAGTTCATTTACTACAAGTTTTTCCA
TGGTAATACGTGTCCTACTATCATAACTGAGGGGAAGACAGACCGCATATATCTG
AAGGCAGCTCTCCACAGTCTCGAGACCTCATATCCCGAACTTTTTCGCGAGAAGA
CGGATTCAAAAAAGAAGGAAATAAATTTGAATATCTTCAAATCAAACGAAAAGA
CAAAATATTTCCTCGACCTCTCTGGCGGGACCGCGGATCTTAAGAAATTCGTAGA
GCGGTACAAGAATAATTACGCGTCCTATTATGGATCCGTTCCCAAGCAGCCAGTC
ATAATGGTTCTCGATAACGACACTGGGCCGAGCGATCTCCTCAATTTTCTGAGAA
ACAAAGTGAAATCCTGTCCTGATGATGTGACTGAAATGCGCAAAATGAAGTATA
TACACGTCTTCTACAATCTCTACATCGTGCTGACGCCATTGAGCCCCTCAGGGGA
GCAAACGTCAATGGAGGACCTCTTCCCCAAGGATATTCTCGACATCAAGATAGA
TGGAAAAAAGTTCAACAAAAACAATGATGGGGATTCAAAGACGGAATATGGAA
AACACATCTTCTCAATGCGAGTCGTACGCGATAAAAAAAGGAAGATCGACTTCA
AAGCTTTTTGTTGCATTTTCGACGCAATCAAGGATATAAAAGAACATTATAAGCT
TATGTTGAACAGCAAAAGGCCGGCGGCCACGAAAAAGGCCGGCCAGGCAAAAA
AGAAAAAGGACTATAAAGATGACGATGACAAGGACTACAAGGATGATGACGAT
AAAGATTATAAAGACGACGATGATAAG

Twist gene fragments:

RT-EcoIII/Ec73

GAAGTGCCATTCCGCCTGACCTATGTACCCATACGACGTCCCAGACTACGCTCCT
CCCAAGAAGAAAAGGAAAGTTATGCGAATATACTCCCTTATAGACAGTCAGACC
CTCATGACCAAGGGGTTTGCCTCAGAGGTCATGCGGTCACCAGAACCCCCGAAG
AAGTGGGACATAGCAAAAAAGAAGGGTGGTATGCGGACAATTTATCACCCATCA
TCTAAAGTTAAACTTATCCAGTACTGGCTCATGAACAACGTATTTAGTAAGCTGC
CAATGCACAACGCGGCTTATGCTTTTGTTAAGAATCGGTCAATCAAGTCTAACGC
GCTTCTCCACGCAGAGTCAAAAAACAAATACTACGTTAAGATCGATCTCAAAGA
CTTCTTTCCCTCAATTAAGTTCACGGATTTTGAGTACGCTTTCACTAGGTATCGCG
ACAGAATCGAATTTACAACCGAGTATGACAAAGAATTGCTGCAGTTGATCAAGA
CTATCTGTTTTATCTCTGACAGCACGCTCCCGATAGGTTTCCCAACCAGCCCACTC
ATTGCGAATTTCGTGGCGAGGGAGCTGGACGAGAAACTTACCCAGAAACTCAAT
GCTATAGATAAACTTAATGCGACATATACCCGCTATGCAGACGATATTATTGTCT
CAACTAATATGAAAGGAGCGAGCAAATTGATCCTTGACTGCTTCAAGCGAACTA
TGAAGGAGATCGGGCCGGACTTCAAGATCAATATCAAAAAGTTCAAAATATGTA
GCGCCAGCGGCGGGTCAATAGTGGTCACGGGGCTGAAAGTGTGCCACGATTTTC
ACATTACGCTTCATCGGAGTATGAAAGATAAAATAAGGTTGCACTTGAGCTTGCT
CTCTAAGGGAATTTTGAAAGACGAAGACCATAACAAACTTTCAGGTTATATTGCA
TACGCTAAAGACATAGACCCCCACTTTTATACCAAACTTAACCGGAAATACTTCC
AAGAAATTAAGTGGATCCAAAACCTGCATAATAAAGTTGAAAAAAGGCCGGCGG

CCACGAAAAAGGCCGGCCAGGCAAAAAAGAAAAAGGACTATAAAGATGACGAT
GACAAGGACTACAAGGATGATGACGATAAAGATTATAAAGACGACGATGATAA
GAGGCTAGGTGGAGGCTCAGTG

RT-EcoV/Ec107

GAAGTGCCATTCCGCCTGACCTATGTACCCATACGACGTCCCAGACTACGCTCCT
CCCAAGAAGAAAAGGAAAGTTATGGACGCCACCCGAACCACTCTTCTTGCGTTG
GATCTCTTCGGGTCACCAGGGTGGTCCGCCGACAAGGAGATACAGCGCCTTCAC
GCTCTGAGCAATCATGCTGGCCGCCATTATAGGAGGATCATTCTTTCTAAACGGC
ACGGAGGCCAGAGACTCGTCTTGGCGCCGGATTACTTGTTGAAAACTGTCCAAC
GGAATATACTGAAAAACGTATTGTCCCAATTTCCGTTGTCACCCTTTGCCACTGC
CTATCGACCAGGATGCCCTATCGTATCTAACGCGCAACCACACTGCCAGCAACCA
CAGATACTGAAATTGGACATTGAAAATTTCTTTGACTCTATCAGTTGGCTCCAAG
TATGGCGCGTTTTCAGACAGGCTCAGCTCCCTCGAAATGTTGTGACAATGCTTAC
CTGGATTTGTTGTTACAACGATGCCCTCCCGCAAGGTGCGCCAACATCCCCTGCG
ATATCTAATCTCGTCATGAGGCGGTTCGATGAGAGAATCGGGGAATGGTGCCAG
GCTCGCGGAATCACATACACGCGGTATTGTGACGACATGACATTTTCCGGCCATT
TCAACGCCCGACAAGTAAAGAACAAAGTGTGTGGCCTTCTCGCAGAGCTTGGGT
TGTCTCTGAATAAGCGAAAAGGGTGCCTTATTGCGGCTTGTAAGCGGCAGCAAG
TGACAGGTATTGTGGTGAACCATAAACCGCAACTTGCAAGAGAGGCCCGCCGGG
CTCTCCGACAGGAGGTGCATCTCTGTCAGAAATATGGGGTCATCTCACACTTGAG
TCACAGGGGTGAACTCGACCCGAGTGGAGACCTGCACGCCCAGGCGACGGCCTA
CTTGTACGCGCTTCAAGGAAGGATAAACTGGCTCCTTCAAATCAATCCTGAAGAC
GAAGCGTTCCAGCAGGCCAGGGAGTCTGTTAAACGGATGCTCGTAGCATGGAAA
AGGCCGGCGGCCACGAAAAAGGCCGGCCAGGCAAAAAAGAAAAAGGACTATAA
AGATGACGATGACAAGGACTACAAGGATGATGACGATAAAGATTATAAAGACG
ACGATGATAAGAGGCTAGGTGGAGGCTCAGTG

RT-MaxII/Mx65

GAAGTGCCATTCCGCCTGACCTATGTACCCATACGACGTCCCAGACTACGCTCCT
CCCAAGAAGAAAAGGAAAGTTATGAGTTGGTTCGATACAACTCTGTCTCGGTTG
AAAGGGCTTTTTTCAAGGCCGGTTACTCGGAGCACAACAGGACTGGATGTCCCAT
TGGACGCGCACGGGAGGCCGCAGGACGTTGTTACAGAAACTGTCAGTACCAGTG
GTCCTCTCAAGCCAGGTCATCTTCGCCAAGTGAGACGAGATGCTCGGCTTCTCCC
GAAAGGTGTTCGACGCTACACCCCCGGCCGAAAGAAATGGATGGAGGCCGCGGA
GGCCAGGCGACTCTTTTCTGCCACGCTGCGAACGAGAAACCGAAATCTGAGAGA
TCTGTTGCCAGACGAAGCACAACTTGCTCGCTATGGTTTGCCAGTCTGGCGCACC
GAAGAAGACGTGGCAGCGGCCCTCGGAGTATCAGTTGGCGTCCTTAGGCATTAC
TCTATACATAGGCCGCGCGAGCGGGTGAGGCACTACGTAACGTTCGCGGTCCCA
AAACGAAGCGGTGGCGTCAGGTTGCTCCATGCACCTAAACGGCGGCTTAAAGCA
CTCCAGCGCCGGATGCTGGCACTTCTGGTAAGCAAACTCCCTGTGTCCCCACAGG
CTCATGGTTTCGTGCCCGGAAGAAGTATAAAGACTGGGGCGGCCCCTCATGTAG

GTAGGCGCGTAGTCCTCAAGCTCGATTTGAAGGACTTCTTTCCGTCTGTTACGTTT
GCGCGGGTTAGAGGCCTGCTTATTGCACTCGGCTATGGTTATCCTGTTGCTGCTA
CACTCGCAGTCCTTATGACGGAATCAGAGCGCCAACCTGTCGAGTTGGAAGGAA
TCCTGTTTCATGTCCCAGTTGGCCCAAGAGTTTGCGTACAGGGGGCGCCTACTTC
CCCAGCACTTTGCAACGCTGTTCTTCTTCGATTGGACAGGAGACTTGCCGGCCTT
GCAAGACGATATGGTTACACGTACACACGGTATGCGGACGACTTGACATTCTCA
GGAGACGACGTCACGGCGTTGGAGCGAGTTAGAGCTCTTGCGGCAAGATATGTC
CAAGAAGAAGGTTTCGAGGTGAACAGAGAGAAAACTCGCGTGCAGAGGCGAGG
CGGAGCCCAGCGAGTGACGGGCGTGACCGTGAACACGACACTGGGACTGTCAAG
GGAAGAACGCCCTAGGTTGCGAGCAATGCTCCACCAGGAGGCACGATCTGAAGA
TGTGGAAGCACACCGAGCCCATCTGGATGGCTTGTTGGCCTACGTCAAAATGCTG
AACCCCGAACAAGCCGAACGGTTGGCCCGACGCCGCAAGCCGCGGGGGACCAA
AAGGCCGGCGGCCACGAAAAAGGCCGGCCAGGCAAAAAAGAAAAAGGACTATA
AAGATGACGATGACAAGGACTACAAGGATGATGACGATAAAGATTATAAAGAC
GACGATGATAAGAGGCTAGGTGGAGGCTCAGTG

RT-MxaI/Mx162

GAAGTGCCATTCCGCCTGACCTATGTACCCATACGACGTCCCAGACTACGCTCCT
CCCAAGAAGAAAAGGAAAGTTATGACAGCAAGGTTGGACCCCTTTGTCCCAGCA
GCTTCACCCCAGGCAGTGCCAACCCCAGAACTGACGGCGCCCAGCTCCGATGCG
GCTGCAAAACGAGAGGCACGGCGCCTGGCGCATGAAGCGTTGTTGGTAAGGGCG
AAAGCTATCGATGAAGCTGGGGGTGCCGACGATTGGGTTCAGGCCCAACTGGTG
TCAAAAGGATTGGCGGTGGAAGATCTCGACTTCTCTTCTGCCTCAGAGAAAGATA
AAAAGGCATGGAAGGAAAAGAAGAAAGCGGAGGCAACAGAAAGGCGGGCCCTT
AAGAGGCAAGCCCATGAGGCGTGGAAAGCAACCCATGTTGGACACTTGGGCGCG
GGAGTCCACTGGGCCGAAGACAGACTCGCGGATGCCTTCGACGTACCGCATAGG
GAAGAACGGGCACGGGCGAATGGGCTGACCGAGCTGGACAGTGCGGAAGCGTT
GGCAAAAGCCCTTGGCTTGTCAGTGTCAAAACTCCGATGGTTTGCATTCCATAGA
GAGGTAGATACAGCCACTCATTATGTATCTTGGACGATCCCAAAGAGGGACGGC
TCCAAGCGAACGATAACATCCCCCAAGCCCGAGTTGAAAGCCGCTCAAAGGTGG
GTTCTCTCAAATGTGGTGGAAAGACTCCCGGTCCATGGTGCTGCTCACGGGTTTG
TCGCTGGTCGCAGCATACTCACTAACGCTCTTGCTCATCAGGGAGCGGATGTAGT
CGTAAAAGTTGACCTTAAAGACTTTTTTCCTAGTGTGACCTGGCGCAGGGTAAAG
GGCCTCTTGAGGAAAGGTGGACTCAGAGAGGGCACAAGTACGTTGCTTAGTCTT
CTTTCTACCGAAGCACCTCGCGAGGCTGTCCAATTTCGAGGCAAATTGCTCCATG
TTGCAAAGGGTCCACGGGCGCTTCCACAGGGAGCCCCTACCTCACCGGGGATCA
CCAACGCACTCTGTCTCAAGCTTGATAAGCGCCTTTCAGCCCTCGCCAAGAGGTT
GGGTTTTACTTACACGCGCTACGCGGACGATTTGACCTTTTCTTGGACTAAGGCG
AAGCAGCCAAAGCCACGACGGACACAACGGCCTCCAGTCGCGGTTCTCTTGTCT
CGCGTCCAAGAAGTGGTTGAGGCCGAAGGCTTTAGGGTCCATCCTGACAAGACC
CGAGTTGCCCGGAAGGGAACGCGACAACGCGTTACGGGGTTGGTGGTAAATGCG
GCGGGGAAAGACGCCCCAGCCGCTCGGGTCCCCCGCGACGTGGTAAGGCAGCTT
AGGGCCGCGATTCATAACCGAAAGAAGGGCAAGCCCGGACGGGAAGGGGAGTC
TCTGGAACAATTGAAGGGCATGGCTGCATTTATTCACATGACAGACCCCGCAAA

AGGTCGGGCATTTCTGGCACAGCTTACGGAGCTGGAAAGTACTGCGTCCGCCGC
ACCTCAAGCAGAAAAAAGGCCGGCGGCCACGAAAAAGGCCGGCCAGGCAAAAA
AGAAAAAGGACTATAAAGATGACGATGACAAGGACTACAAGGATGATGACGAT
AAAGATTATAAAGACGACGATGATAAGAGGCTAGGTGGAGGCTCAGTG

RT-SenII/St85

GAAGTGCCATTCCGCCTGACCTATGTACCCATACGACGTCCCAGACTACGCTCCT
CCCAAGAAGAAAAGGAAAGTTATGGATATTCTGCAACACATATCCGACCTGCTG
CTGACAAAGAAGTCAGAAATTATTTCTTTTAGTCTCACCGCACCTTATCGGTACA
AAATTTATAAGATCGCTAAAAGAAACTCCGACAAGAAAAGAACCATCGCCCACC
CGAGTAAAGAACTTAAATTCATACAGAGAGAGATTACTGAGTACCTGACAGATA
AGCTGCCCGTGCACGAATGCGCCTTTGCTTACAAAAAGGGCTCTAGTATAAAAA
CAAACGCACAAGTTCATCTCCATACGAAATATCTGTTGAAAATGGATTTTGAAAA
CTTTTTTCCCTCCATAACGCCGCGGCTCTTCTTCTCAAAACTGCGCTTGGCTAACA
TCGACTTGACTGCTGATGATAAGGTTTTGTTGGAGAATATCTTGTTTTTTAAATCC
AAACGCAACAGTAATCTGCGCTTGAGTATCGGTGCGCCGTCTTCTCCATTGATTT
CCAACTTCGTTATGTACTTCTGGGACATTGAGGTGCAAGAGATATGCTCCAAGAT
CGGGGTTAATTATACCCGCTACGCAGATGATTTGACCTTTAGCACAAATAATAAA
GACGTGCTGTTTGATATACCTGATATGTTGGAAAACGTTCTGCCGAAGTATAGTT
TGGGGCGGATACGGATTAACCACGAAAAGACTGTGTTCTCTAGCAAGGGCCACA
ACCGACATGTAACCGGGATAACTCTTACTAATGATAACAAGCTGTCTATCGGTCG
GGAGCGGAAAAGGAAGATCAGCGCGATGATCCACCACTTCATCAATGGAAAGCT
GTCAACAGATGAGTGCAACAAATTGGTGGGCCTCCTGGCCTTCGCCAAAAATAT
AGAGCCATCCTTTTACAAAAGCATGGTTATTAAGTATGGGAGTGATAACATTTAT
AAACTCCAGAAGCAGAAAGACAAGAAAAGGCCGGCGGCCACGAAAAAGGCCGG
CCAGGCAAAAAAGAAAAAGGACTATAAAGATGACGATGACAAGGACTACAAGG
ATGATGACGATAAAGATTATAAAGACGACGATGATAAGAGGCTAGGTGGAGGCT
CAGTG

gBlock Gene Fragments ordered from IDT:

RT-SauI_Sa163

ATGTACCCATACGACGTCCCAGACTACGCTCCTCCCAAGAAGAAAAGGAAAGTT
ATGACGGCAAAACTGGAAAGCCACGTGCCTGCTGCACCACCAGTCTCAGCGGAA
GCGCCGGCACCGACGCGACCTGACGCTGCTAAGCAGGAGGCTCGACGAGCCCAC
CATGAAGCCCTGAGACTGCGGTGGAAAGCAATCGAGGAAGCTGGGGGCACTGAC
GCCTGGGTACGACAGCAGTTGGTTGCGAAAGGCGTAGCGGCGGAAGAGGTGGAC
TTCGAAAGTCTCTCCGACAAGCAAAAAGCGGCTTGGAAAGAGAAAAAAAAAGCT
GAGGCAACCGAGCGGAGGGCTCAAAAACGACTTGCCTGGGAAGCCTGGAAAGC
GACCCATATTCATCATTTGGGCGTTGGAGTTCACTGGGACGAAGCCGGCGGGCCT
GACAAGTTCGATGTGGCAGGGAGGGAAGAAAGGGCAAAGGCAAACGGGTTGCC
CGAAGGTCTTGACAGCGTAGAAGCATTGGCGAAAGCGCTGGGAATCTCAGTATC
CCGATTGCGCTGGTTCTCTTTTCACAGGGAAGTTGATACAGGGACACACTACCAA
ACTTGGGAGATCCCAAAGAGGGATGGGGGCAAACGCACCCTCACGGCCCCGAAA

AGGGAGCTTAAGGCAGTACAAAGGTGGGTACTCGCCAATGTGGTGGAAAGACTT
CCCGTACACGGAGCTGCTCATGGGTTTGTTGCAGGACGCTCCATACTCACTAATG
CGCTCGCTCACCAAGGAGCTGACGTGGTTGTTAAAGTTGATATGAAAGATTTTTT
TCCCAGTGTCACATGGCCTCGAGTTAAAGGCCTGCTCAGAAAGGGAGGTTTGCCT
GAAAACTTGGCGACCCTGCTCGCCCTTCTGAGTACCGAAGCTCCACGGGAAGTA
GTCCGATTCAGAGGGGAAACGCTGTACGTCGCTAAGGGCCCACGCGCTCTGCCA
CAGGGCGCGCCTACCAGTCCTGCGCTTACTAACGCATTGTGTCTTAGGTTGGATA
AGCGGTTGAGCGCGCTGAGTAAGCGGTTGGGTTTCACATACACCAGATACGCTG
ACGATCTTACATTTAGTTGGAGGCGCGCTAAAAAATCTCGCCAGAAAGAGTTGC
CGCTGGCTGATGCTCCCGTCGCGCTTCTCCTGGCCCGGGTTAAGGGAGTGTTGGA
AGCAGAAGGTTTCACTTTGCATCCTGATAAGACCAGGGTTCAGCGGAAGGGAAG
CCGGCAACGAGTGACCGGCCTCGTGGTTAATGAAGCGCCCGAAGGAGTGCCAGG
TGCCCGAGTTCCCCGAGATGTTGTTCGGAGATTGCGAGCGGCAATACACAATAG
AGAGCAAGGTAAACCAGGGCCGACTGGAGAAACGTTGGAACAACTCAAAGGCT
TGGCAGCTTTTCTGCACATGACAGACGCTGAAAAAGGAAGAGCGTTTCTGAGAA
GACTTGAGGCCCTGGAGAAGAGGCAAACTGCGAAAAGGCCGGCGGCCACGAAA
AAGGCCGGCCAGGCAAAAAAGAAAAAGGACTATAAAGATGACGATGACAAGGA
CTACAAGGATGATGACGATAAAGATTATAAAGACGACGATGATAAG

**Supplemental Note 2**

**Plasmids used in FACS plots**

**pBZ186-pU6-(BbsI)_CBh-sv40NLS-Cas9-NLS T2A-mCherry**

gagggcctatttcccatgattccttcatatttgcatatacgatacaaggctgttagagagataattggaattaatttgactgtaaacacaaag
atattagtacaaaatacgtgacgtagaaagtaataatttcttgggtagtttgcagttttaaaattatgttttaaaatggactatcatatgcttacc
gtaacttgaaagtatttcgatttcttggctttatatatcttgtggaaaggacgaaacaccgggtcttcgagaagacctgttttagagctagaa
atagcaagttaaaataaggctagtccgttatcaacttgaaaaagtggcaccgagtcggtgctttttttgttttagagctagaaatagcaagtt
aaaataaggctagtccgtttttagcgcgtgcgccaattctgcagacaaatggctctagaggtacccgttacataacttacggtaaatggc
ccgcctggctgaccgcccaacgacccccgcccattgacgtcaatagtaacgccaatagggactttccattgacgtcaatgggtggagt
atttacggtaaactgcccacttggcagtacatcaagtgtatcatatgccaagtacgccccctattgacgtcaatgacggtaaatggcccg
cctggcattgtgcccagtacatgaccttatgggactttcctacttggcagtacatctacgtattagtcatcgctattaccatggtcgaggtg
agccccacgttctgcttcactctccccatctcccccccctccccacccccaattttgtatttatttattttttaattattttgtgcagcgatggggg
gcgggggggggggggggcgcgcgccaggcggggcggggcggggcgaggggcggggcggggcgaggcggagaggtgc
ggcggcagccaatcagagcggcgcgctccgaaagtttccttttatggcgaggcggcggcggcggcggccctataaaaagcgaagc
gcgcggcgggcgggagtcgctgcgacgctgccttcgccccgtgccccgctccgccgccgcctcgcgccgcccgccccggctctg
actgaccgcgttactcccacaggtgagcgggcgggacggcccttctcctccgggctgtaattagctgagcaagaggtaagggtttaa
gggatggttggttggtggggtattaatgtttaattacctggagcacctgcctgaaatcactttttttcaggttggaccggtgccaccatgga
ctataaggaccacgacggagactacaaggatcatgatattgattacaaagacgatgacgataagatggccccaaagaagaagcgga
aggtcggtatccacggagtcccagcagccgacaagaagtacagcatcggcctggacatcggcaccaactctgtgggctgggccgtg
atcaccgacgagtacaaggtgcccagcaagaaattcaaggtgctgggcaacaccgaccggcacagcatcaagaagaacctgatcg
gagccctgctgttcgacagcggcgaaacagccgaggccacccggctgaagagaaccgccagaagaagatacaccagacggaag
aaccggatctgctatctgcaagagatcttcagcaacgagatggccaaggtggacgacagcttcttccacagactggaagagtccttcct
ggtggaagaggataagaagcacgagcggcaccccatcttcggcaacatcgtggacgaggtggcctaccacgagaagtaccccacc
atctaccacctgagaaagaaactggtggacagcaccgacaaggccgacctgcggctgatctatctggccctggcccacatgatcaag
ttccggggccacttcctgatcgagggcgacctgaaccccgacaacagcgacgtggacaagctgttcatccagctggtgcagacctac
aaccagctgttcgaggaaaaccccatcaacgccagcggcgtggacgccaaggccatcctgtctgccagactgagcaagagcagac
ggctggaaaatctgatcgcccagctgcccggcgagaagaagaatggcctgttcggaaacctgattgccctgagcctgggcctgacc

cccaacttcaagagcaacttcgacctggccgaggatgccaaactgcagctgagcaaggacacctacgacgacgacctggacaacct
gctggcccagatcggcgaccagtacgccgacctgtttctggccgccaagaacctgtccgacgccatcctgctgagcgacatcctgag
agtgaacaccgagatcaccaaggccccctgagcgcctctatgatcaagagatacgacgagcaccaccaggacctgaccctgctga
aagctctcgtgcggcagcagctgcctgagaagtacaaagagattttcttcgaccagagcaagaacggctacgccggctacattgacg
gcggagccagccaggaagagttctacaagttcatcaagcccatcctggaaaagatggacggcaccgaggaactgctcgtgaagctg
aacagagaggacctgctgcggaagcagcggaccttcgacaacggcagcatcccccaccagatccacctgggagagctgcacgcc
attctgcggcggcaggaagatttttacccattcctgaaggacaaccgggaaaagatcgagaagatcctgaccttccgcatcccctacta
cgtgggccctctggccaggggaaacagcagattcgcctggatgaccagaaagagcgaggaaaccatcacccccctggaacttcgag
gaagtggtggacaagggcgcttccgcccagagcttcatcgagcggatgaccaacttcgataagaacctgcccaacgagaaggtgct
gcccaagcacagcctgctgtacgagtacttcaccgtgtataacgagctgaccaaagtgaaatacgtgaccgagggaatgagaaagcc
cgccttcctgagcggcgagcagaaaaaggccatcgtggacctgctgttcaagaccaaccggaaagtgaccgtgaagcagctgaaa
gaggactacttcaagaaaatcgagtgcttcgactccgtggaaatctccggcgtggaagatcggttcaacgcctccctgggcacatacc
acgatctgctgaaaattatcaaggacaaggacttcctggacaatgaggaaaacgaggacattctggaagatatcgtgctgaccctgac
actgtttgaggacagagagatgatcgaggaacggctgaaaacctatgcccacctgttcgacgacaaagtgatgaagcagctgaagcg
gcggagatacaccggctggggcaggctgagccggaagctgatcaacggcatccgggacaagcagtccggcaagacaatcctgga
tttcctgaagtccgacggcttcgccaacagaaacttcatgcagctgatccacgacgacagcctgacctttaaagaggacatccagaaa
gcccaggtgtccggccagggcgatagcctgcacgagcacattgccaatctggccggcagccccgccattaagaagggcatcctgc
agacagtgaaggtggtggacgagctcgtgaaagtgatgggccggcacaagcccgagaacatcgtgatcgaaatggccagagaga
accagaccacccagaagggacagaagaacagccgcgagagaatgaagcggatcgaagagggcatcaaagagctgggcagcca
gatcctgaaagaacaccccgtggaaaacacccagctgcagaacgagaagctgtacctgtactacctgcagaatgggcgggatatgta
cgtggaccaggaactggacatcaaccggctgtccgactacgatgtggaccatatcgtgcctcagagctttctgaaggacgactccatc
gacaacaaggtgctgaccagaagcgacaagaaccggggcaagagcgacaacgtgccctccgaagaggtcgtgaagaagatgaa
gaactactggcggcagctgctgaacgccaagctgattacccagagaaagttcgacaatctgaccaaggccgagagaggcggcctg
agcgaactggataaggccggcttcatcaagagacagctggtggaaacccggcagatcacaaagcacgtggcacagatcctggactc
ccggatgaacactaagtacgacgagaatgacaagctgatccgggaagtgaaagtgatcaccctgaagtccaagctggtgtccgatttc
cggaaggatttccagttttacaaagtgcgcgagatcaacaactaccaccacgcccacgacgcctacctgaacgccgtcgtgggaacc
gccctgatcaaaaagtaccctaagctggaaagcgagttcgtgtacggcgactacaaggtgtacgacgtgcggaagatgatcgccaag
agcgagcaggaaatcggcaaggctaccgccaagtacttcttctacagcaacatcatgaactttttcaagaccgagattaccctggccaa
cggcgagatccggaagcggcctctgatcgagacaaacggcgaaaccggggagatcgtgtgggataagggccgggattttgccacc
gtgcggaaagtgctgagcatgccccaagtgaatatcgtgaaaaagaccgaggtgcagacaggcggcttcagcaaagagtctatcct
gcccaagaggaacagcgataagctgatcgccagaaagaaggactgggaccctaagaagtacggcggcttcgacagccccaccgt
ggcctattctgtgctggtggtggccaaagtggaaaagggcaagtccaagaaactgaagagtgtgaaagagctgctggggatcaccat
catggaaagaagcagcttcgagaagaatcccatcgactttctggaagccaagggctacaaagaagtgaaaaaggacctgatcatcaa
gctgcctaagtactccctgttcgagctggaaaacggccggaagagaatgctggcctctgccggcgaactgcagaagggaaacgaac
tggccctgccctccaaatatgtgaacttcctgtacctggccagccactatgagaagctgaagggctcccccgaggataatgagcagaa
acagctgtttgtggaacagcacaagcactacctggacgagatcatcgagcagatcagcgagttctccaagagagtgatcctggccga
cgctaatctggacaaagtgctgtccgcctacaacaagcaccgggataagcccatcagagagcaggccgagaatatcatccacctgttt
accctgaccaatctgggagcccctgccgccttcaagtactttgacaccaccatcgaccggaagaggtacaccagcaccaaagaggtg
ctggacgccaccctgatccaccagagcatcaccggcctgtacgagacacggatcgacctgtctcagctgggaggcgacaaaaggc
cggcggccacgaaaaaggccggccaggcaaaaaagaaaaagcttgagggcagaggaagtctgctaacatgcggtgacgtggag
gagaatcccggccctgctagcatggtgagcaagggcgaggaggataacatggccatcatcaaggagttcatgcgcttcaaggtgca
catggagggctccgtgaacggccacgagttcgagatcgagggcgagggcgagggccgcccctacgagggcacccagaccgcca
agctgaaggtgaccaagggtggccccctgcccttcgcctgggacatcctgtcccctcagttcatgtacggctccaaggcctacgtgaa
gcaccccgccgacatccccgactacttgaagctgtccttccccgagggcttcaagtgggagcgcgtgatgaacttcgaggacggcg
gcgtggtgaccgtgacccaggactcctccctgcaggacggcgagttcatctacaaggtgaagctgcgcggcaccaacttcccctccg
acggccccgtaatgcagaagaagaccatgggctgggaggcctcctccgagcggatgtaccccgaggacggcgccctgaagggcg
agatcaagcagaggctgaagctgaaggacggcggccactacgacgctgaggtcaagaccacctacaaggccaagaagcccgtgc
agctgcccggcgcctacaacgtcaacatcaagttggacatcacctcccacaacgaggactacaccatcgtggaacagtacgaacgc
gccgagggccgccactccaccggcggcatggacgagctgtacaagtgagaattcctagagctcgctgatcagcctcgactgtgcctt
ctagttgccagccatctgttgtttgcccctcccccgtgccttccttgaccctggaaggtgccactcccactgtcctttcctaataaaatgag

gaaattgcatcgcattgtctgagtaggtgtcattctattctggggggtggggtggggcaggacagcaagggggaggattgggaagag
aatagcaggcatgctggggagcggccgcaggaacccctagtgatggagttggccactccctctctgcgcgctcgctcgctcactgag
gccgggcgaccaaaggtcgcccgacgcccgggctttgcccgggcggcctcagtgagcgagcgagcgcgcagctgcctgcaggg
gcgcctgatgcggtattttctccttacgcatctgtgcggtatttcacaccgcatacgtcaaagcaaccatagtacgcgccctgtagcggc
gcattaagcgcggcgggtgtggtggttacgcgcagcgtgaccgctacacttgccagcgccctagcgcccgctcctttcgctttcttccc
ttcctttctcgccacgttcgccggctttccccgtcaagctctaaatcgggggctccctttagggttccgatttagtgctttacggcacctcg
accccaaaaaacttgatttgggtgatggttcacgtagtgggccatcgccctgatagacggtttttcgccctttgacgttggagtccacgtt
ctttaatagtggactcttgttccaaactggaacaacactcaaccctatctcgggctattcttttgatttataagggattttgccgatttcggCc
tattggttaaaaaatgagctgatttaacaaaaatttaacgcgaattttaacaaaatattaacgtttacaattttatggtgcactctcagtacaat
ctgctctgatgccgcatagttaagccagccccgacacccgccaacacccgctgacgcgccctgacgggcttgtctgctcccggcatc
cgcttacagacaagctgtgaccgtctccgggagctgcatgtgtcagaggttttcaccgtcatcaccgaaacgcgcgagacgaaaggg
cctcgtgatacgcctatttttataggttaatgtcatgataataatggtttcttagacgtcaggtggcacttttcggggaaatgtgcgcggaac
ccctatttgtttatttttctaaatacattcaaatatgtatccgctcatgagacaataaccctgataaatgcttcaataatattgaaaaaggaaga
gtatgagtattcaacatttccgtgtcgcccttattcccttttttgcggcattttgccttcctgtttttgctcacccagaaacgctggtgaaagta
aaagatgctgaagatcagttgggtgcacgagtgggttacatcgaactggatctcaacagcggtaagatccttgagagttttcgccccga
agaacgttttccaatgatgagcacttttaaagttctgctatgtggcgcggtattatcccgtattgacgccgggcaagagcaactcggtcgc
cgcatacactattctcagaatgacttggttgagtactcaccagtcacagaaaagcatcttacggatggcatgacagtaagagaattatgc
agtgctgccataaccatgagtgataacactgcggccaacttacttctgacaacgatcggaggaccgaaggagctaaccgcttttttgca
caacatgggggatcatgtaactcgccttgatcgttgggaaccggagctgaatgaagccataccaaacgacgagcgtgacaccacgat
gcctgtagcaatggcaacaacgttgcgcaaactattaactggcgaactacttactctagcttcccggcaacaattaatagactggatgga
ggcggataaagttgcaggaccacttctgcgctcggcccttccggctggctggtttattgctgataaatctggagccggtgagcgtggaa
gccgcggtatcattgcagcactggggccagatggtaagccctcccgtatcgtagttatctacacgacggggagtcaggcaactatgga
tgaacgaaatagacagatcgctgagataggtgcctcactgattaagcattggtaactgtcagaccaagtttactcatatatactttagattg
atttaaaacttcatttttaatttaaaaggatctaggtgaagatcctttttgataatctcatgaccaaaatcccttaacgtgagttttcgttccact
gagcgtcagaccccgtagaaaagatcaaaggatcttcttgagatcctttttttctgcgcgtaatctgctgcttgcaaacaaaaaaaccacc
gctaccagcggtggtttgtttgccggatcaagagctaccaactctttttccgaaggtaactggcttcagcagagcgcagataccaaatac
tgtccttctagtgtagccgtagttaggccaccacttcaagaactctgtagcaccgcctacatacctcgctctgctaatcctgttaccagtgg
ctgctgccagtggcgataagtcgtgtcttaccgggttggactcaagacgatagttaccggataaggcgcagcggtcgggctgaacgg
ggggttcgtgcacacagcccagcttggagcgaacgacctacaccgaactgagatacctacagcgtgagctatgagaaagcgccacg
cttcccgaagggagaaaggcggacaggtatccggtaagcggcagggtcggaacaggagagcgcacgagggagcttccaggggg
aaacgcctggtatctttatagtcctgtcgggtttcgccacctctgacttgagcgtcgatttttgtgatgctcgtcaggggggcggagcctat
ggaaaaacgccagcaacgcggcctttttacggttcctggccttttgctggccttttgctcacatgt

**pBZ191-pU6-sgBFP-CBh-sv40NLS-Cas9-NLS-T2A-mCherry**

gagggcctatttcccatgattccttcatatttgcatatacgatacaaggctgttagagagataattggaattaatttgactgtaaacacaaag
atattagtacaaaatacgtgacgtagaaagtaataatttcttgggtagtttgcagttttaaaattatgttttaaaatggactatcatatgcttacc
gtaacttgaaagtatttcgatttcttggctttatatatcttgtggaaaggacgaaacaccGCTGAAGCACTGCACGCCAT
gttttagagctagaaatagcaagttaaaataaggctagtccgttatcaacttgaaaaagtggcaccgagtcggtgcttttttgttttagagct
agaaatagcaagttaaaataaggctagtccgtttttagcgcgtgcgccaattctgcagacaaatggctctagaggtacccgttacataac
ttacggtaaatggcccgcctggctgaccgcccaacgacccccgcccattgacgtcaatagtaacgccaatagggactttccattgacg
tcaatgggtggagtatttacggtaaactgcccacttggcagtacatcaagtgtatcatatgccaagtacgccccctattgacgtcaatgac
ggtaaatggcccgcctggcattgtgcccagtacatgaccttatgggactttcctacttggcagtacatctacgtattagtcatcgctattac
catggtcgaggtgagccccacgttctgcttcactctccccatctcccccccctccccacccccaattttgtatttatttattttttaattattttgt
gcagcgatgggggcgggggggggggggggcgcgcgccaggcggggcggggcggggcgaggggcggggcggggcgag
gcggagaggtgcggcggcagccaatcagagcggcgcgctccgaaagtttccttttatggcgaggcggcggcggcggcggccctat
aaaaagcgaagcgcgcggcgggcgggagtcgctgcgacgctgccttcgccccgtgccccgctccgccgccgcctcgcgccgccc
gccccggctctgactgaccgcgttactcccacaggtgagcgggcgggacggcccttctcctccgggctgtaattagctgagcaagag
gtaagggtttaagggatggttggttggtggggtattaatgtttaattacctggagcacctgcctgaaatcactttttttcaggttggaccggt

gccaccatggactataaggaccacgacggagactacaaggatcatgatattgattacaaagacgatgacgataagatggccccaaag
aagaagcggaaggtcggtatccacggagtcccagcagccgacaagaagtacagcatcggcctggacatcggcaccaactctgtgg
gctgggccgtgatcaccgacgagtacaaggtgcccagcaagaaattcaaggtgctgggcaacaccgaccggcacagcatcaagaa
gaacctgatcggagccctgctgttcgacagcggcgaaacagccgaggccacccggctgaagagaaccgccagaagaagatacac
cagacggaagaaccggatctgctatctgcaagagatcttcagcaacgagatggccaaggtggacgacagcttcttccacagactgga
agagtccttcctggtggaagaggataagaagcacgagcggcaccccatcttcggcaacatcgtggacgaggtggcctaccacgaga
agtaccccaccatctaccacctgagaaagaaactggtggacagcaccgacaaggccgacctgcggctgatctatctggccctggcc
cacatgatcaagttccggggccacttcctgatcgagggcgacctgaaccccgacaacagcgacgtggacaagctgttcatccagctg
gtgcagacctacaaccagctgttcgaggaaaaccccatcaacgccagcggcgtggacgccaaggccatcctgtctgccagactgag
caagagcagacggctggaaaatctgatcgcccagctgcccggcgagaagaagaatggcctgttcggaaacctgattgccctgagcc
tgggcctgacccccaacttcaagagcaacttcgacctggccgaggatgccaaactgcagctgagcaaggacacctacgacgacgac
ctggacaacctgctggcccagatcggcgaccagtacgccgacctgtttctggccgccaagaacctgtccgacgccatcctgctgagc
gacatcctgagagtgaacaccgagatcaccaaggcccccctgagcgcctctatgatcaagagatacgacgagcaccaccaggacct
gaccctgctgaaagctctcgtgcggcagcagctgcctgagaagtacaaagagattttcttcgaccagagcaagaacggctacgccgg
ctacattgacggcggagccagccaggaagagttctacaagttcatcaagcccatcctggaaaagatggacggcaccgaggaactgc
tcgtgaagctgaacagagaggacctgctgcggaagcagcggaccttcgacaacggcagcatcccccaccagatccacctgggaga
gctgcacgccattctgcggcggcaggaagatttttacccattcctgaaggacaaccgggaaaagatcgagaagatcctgaccttccgc
atcccctactacgtgggccctctggccaggggaaacagcagattcgcctggatgaccagaaagagcgaggaaaccatcaccccctg
gaacttcgaggaagtggtggacaagggcgcttccgcccagagcttcatcgagcggatgaccaacttcgataagaacctgcccaacg
agaaggtgctgcccaagcacagcctgctgtacgagtacttcaccgtgtataacgagctgaccaaagtgaaatacgtgaccgagggaa
tgagaaagcccgccttcctgagcggcgagcagaaaaaggccatcgtggacctgctgttcaagaccaaccggaaagtgaccgtgaa
gcagctgaaagaggactacttcaagaaaatcgagtgcttcgactccgtggaaatctccggcgtggaagatcggttcaacgcctccctg
ggcacataccacgatctgctgaaaattatcaaggacaaggacttcctggacaatgaggaaaacgaggacattctggaagatatcgtgc
tgaccctgacactgtttgaggacagagagatgatcgaggaacggctgaaaacctatgcccacctgttcgacgacaaagtgatgaagc
agctgaagcggcggagatacaccggctggggcaggctgagccggaagctgatcaacggcatccgggacaagcagtccggcaag
acaatcctggatttcctgaagtccgacggcttcgccaacagaaacttcatgcagctgatccacgacgacagcctgacctttaaagagga
catccagaaagcccaggtgtccggccagggcgatagcctgcacgagcacattgccaatctggccggcagccccgccattaagaag
ggcatcctgcagacagtgaaggtggtggacgagctcgtgaaagtgatgggccggcacaagcccgagaacatcgtgatcgaaatgg
ccagagagaaccagaccacccagaagggacagaagaacagccgcgagagaatgaagcggatcgaagagggcatcaaagagct
gggcagccagatcctgaaagaacaccccgtggaaaacacccagctgcagaacgagaagctgtacctgtactacctgcagaatggg
cgggatatgtacgtggaccaggaactggacatcaaccggctgtccgactacgatgtggaccatatcgtgcctcagagctttctgaagg
acgactccatcgacaacaaggtgctgaccagaagcgacaagaaccggggcaagagcgacaacgtgccctccgaagaggtcgtga
agaagatgaagaactactggcggcagctgctgaacgccaagctgattacccagagaaagttcgacaatctgaccaaggccgagaga
ggcggcctgagcgaactggataaggccggcttcatcaagagacagctggtggaaacccggcagatcacaaagcacgtggcacag
atcctggactcccggatgaacactaagtacgacgagaatgacaagctgatccgggaagtgaaagtgatcaccctgaagtccaagctg
gtgtccgatttccggaaggatttccagttttacaaagtgcgcgagatcaacaactaccaccacgcccacgacgcctacctgaacgccgt
cgtgggaaccgccctgatcaaaaagtaccctaagctggaaagcgagttcgtgtacggcgactacaaggtgtacgacgtgcggaaga
tgatcgccaagagcgagcaggaaatcggcaaggctaccgccaagtacttcttctacagcaacatcatgaactttttcaagaccgagatt
accctggccaacggcgagatccggaagcggcctctgatcgagacaaacggcgaaaccggggagatcgtgtgggataagggccgg
gattttgccaccgtgcggaaagtgctgagcatgccccaagtgaatatcgtgaaaaagaccgaggtgcagacaggcggcttcagcaaa
gagtctatcctgcccaagaggaacagcgataagctgatcgccagaaagaaggactgggaccctaagaagtacggcggcttcgaca
gccccaccgtggcctattctgtgctggtggtggccaaagtggaaaagggcaagtccaagaaactgaagagtgtgaaagagctgctg
gggatcaccatcatggaaagaagcagcttcgagaagaatcccatcgactttctggaagccaagggctacaaagaagtgaaaaagga
cctgatcatcaagctgcctaagtactccctgttcgagctggaaaacggccggaagagaatgctggcctctgccggcgaactgcagaa
gggaaacgaactggccctgccctccaaatatgtgaacttcctgtacctggccagccactatgagaagctgaagggctcccccgagga
taatgagcagaaacagctgtttgtggaacagcacaagcactacctggacgagatcatcgagcagatcagcgagttctccaagagagt
gatcctggccgacgctaatctggacaaagtgctgtccgcctacaacaagcaccgggataagcccatcagagagcaggccgagaata
tcatccacctgtttaccctgaccaatctgggagcccctgccgccttcaagtactttgacaccaccatcgaccggaagaggtacaccagc
accaaagaggtgctggacgccaccctgatccaccagagcatcaccggcctgtacgagacacggatcgacctgtctcagctgggagg
cgacaaaaggccggcggccacgaaaaaggccggccaggcaaaaaagaaaaagcttgagggcagaggaagtctgctaacatgcg

gtgacgtggaggagaatcccggccctgctagcatggtgagcaagggcgaggaggataacatggccatcatcaaggagttcatgcgc
ttcaaggtgcacatggagggctccgtgaacggccacgagttcgagatcgagggcgagggcgagggccgcccctacgagggcacc
cagaccgccaagctgaaggtgaccaagggtggcccccctgcccttcgcctgggacatcctgtcccctcagttcatgtacggctccaag
gcctacgtgaagcaccccgccgacatccccgactacttgaagctgtccttccccgagggcttcaagtgggagcgcgtgatgaacttcg
aggacggcggcgtggtgaccgtgacccaggactcctccctgcaggacggcgagttcatctacaaggtgaagctgcgcggcaccaa
cttcccctccgacggccccgtaatgcagaagaagaccatgggctgggaggcctcctccgagcggatgtaccccgaggacggcgcc
ctgaagggcgagatcaagcagaggctgaagctgaaggacggcggccactacgacgctgaggtcaagaccacctacaaggccaag
aagcccgtgcagctgcccggcgcctacaacgtcaacatcaagttggacatcacctcccacaacgaggactacaccatcgtggaaca
gtacgaacgcgccgagggccgccactccaccggcggcatggacgagctgtacaagtgagaattcctagagctcgctgatcagcctc
gactgtgccttctagttgccagccatctgttgtttgcccctcccccgtgccttccttgaccctggaaggtgccactcccactgtcctttccta
ataaaatgaggaaattgcatcgcattgtctgagtaggtgtcattctattctggggggtggggtggggcaggacagcaagggggaggat
tgggaagagaatagcaggcatgctggggagcggccgcaggaacccctagtgatggagttggccactccctctctgcgcgctcgctc
gctcactgaggccgggcgaccaaaggtcgcccgacgcccgggctttgcccgggcggcctcagtgagcgagcgagcgcgcagct
gcctgcaggggcgcctgatgcggtattttctccttacgcatctgtgcggtatttcacaccgcatacgtcaaagcaaccatagtacgcgcc
ctgtagcggcgcattaagcgcggcgggtgtggtggttacgcgcagcgtgaccgctacacttgccagcgccctagcgcccgctccttt
cgctttcttcccttcctttctcgccacgttcgccggctttccccgtcaagctctaaatcgggggctccctttagggttccgatttagtgcttta
cggcacctcgaccccaaaaaacttgatttgggtgatggttcacgtagtgggccatcgccctgatagacggtttttcgccctttgacgttg
gagtccacgttctttaatagtggactcttgttccaaactggaacaacactcaaccctatctcgggctattcttttgatttataagggattttgc
cgatttcggCctattggttaaaaaatgagctgatttaacaaaaatttaacgcgaattttaacaaaatattaacgtttacaattttatggtgcac
tctcagtacaatctgctctgatgccgcatagttaagccagccccgacacccgccaacacccgctgacgcgccctgacgggcttgtctg
ctcccggcatccgcttacagacaagctgtgaccgtctccgggagctgcatgtgtcagaggttttcaccgtcatcaccgaaacgcgcga
gacgaaagggcctcgtgatacgcctatttttataggttaatgtcatgataataatggtttcttagacgtcaggtggcacttttcggggaaat
gtgcgcggaacccctatttgtttatttttctaaatacattcaaatatgtatccgctcatgagacaataaccctgataaatgcttcaataatattg
aaaaaggaagagtatgagtattcaacatttccgtgtcgcccttattcccttttttgcggcattttgccttcctgtttttgctcacccagaaacg
ctggtgaaagtaaaagatgctgaagatcagttgggtgcacgagtgggttacatcgaactggatctcaacagcggtaagatccttgaga
gttttcgccccgaagaacgttttccaatgatgagcacttttaaagttctgctatgtggcgcggtattatcccgtattgacgccgggcaaga
gcaactcggtcgccgcatacactattctcagaatgacttggttgagtactcaccagtcacagaaaagcatcttacggatggcatgacagt
aagagaattatgcagtgctgccataaccatgagtgataacactgcggccaacttacttctgacaacgatcggaggaccgaaggagcta
accgcttttttgcacaacatgggggatcatgtaactcgccttgatcgttgggaaccggagctgaatgaagccataccaaacgacgagc
gtgacaccacgatgcctgtagcaatggcaacaacgttgcgcaaactattaactggcgaactacttactctagcttcccggcaacaattaa
tagactggatggaggcggataaagttgcaggaccacttctgcgctcggcccttccggctggctggtttattgctgataaatctggagcc
ggtgagcgtggaagccgcggtatcattgcagcactggggccagatggtaagccctcccgtatcgtagttatctacacgacggggagt
caggcaactatggatgaacgaaatagacagatcgctgagataggtgcctcactgattaagcattggtaactgtcagaccaagtttactc
atatatactttagattgatttaaaacttcatttttaatttaaaaggatctaggtgaagatcctttttgataatctcatgaccaaaatcccttaacgt
gagttttcgttccactgagcgtcagaccccgtagaaaagatcaaaggatcttcttgagatcctttttttctgcgcgtaatctgctgcttgcaa
acaaaaaaaccaccgctaccagcggtggtttgtttgccggatcaagagctaccaactctttttccgaaggtaactggcttcagcagagc
gcagataccaaatactgtccttctagtgtagccgtagttaggccaccacttcaagaactctgtagcaccgcctacatacctcgctctgcta
atcctgttaccagtggctgctgccagtggcgataagtcgtgtcttaccgggttggactcaagacgatagttaccggataaggcgcagcg
gtcgggctgaacggggggttcgtgcacacagcccagcttggagcgaacgacctacaccgaactgagatacctacagcgtgagctat
gagaaagcgccacgcttcccgaagggagaaaggcggacaggtatccggtaagcggcagggtcggaacaggagagcgcacgag
ggagcttccagggggaaacgcctggtatctttatagtcctgtcgggtttcgccacctctgacttgagcgtcgatttttgtgatgctcgtcag
gggggcggagcctatggaaaaacgccagcaacgcggcctttttacggttcctggccttttgctggccttttgct

**pBZ193-pU6-sgBFP-Ht_CBh-sv40NLS-Cas9-NLS-T2A-mCherry-P2A-Sa163RT**

gagggcctatttcccatgattccttcatatttgcatatacgatacaaggctgttagagagataattggaattaatttgactgtaaacacaaag
atattagtacaaaatacgtgacgtagaaagtaataatttcttgggtagtttgcagttttaaaattatgttttaaaatggactatcatatgcttacc
gtaacttgaaagtatttcgatttcttggctttatatatcttgtggaaaggacgaaaCACCGCTGAAGCACTGCACGCC
ATgttttagagctagaaatagcaagttaaaataaggctagtccgttatcaacttgaaaaagtggcaccgagtcggtgcCGTACG
GTGCGAGCGACCGAGAGAGGTCCCAAGCCATCAGCCTCAGCGCCTCGAGCGCGA
GAGCGGCGTTGCGCCGCTCTGGTTGAATTGCAGGACACTCTCCGCAAGGTAGCCT

GTTCTTGGCTCTCTTCCCTCCGGTGAGTACCTCTCCGGCCGGGGAGCTGAACCAA
CGAACTAGTGCCACCTACGGCAAGCTGACCCTGAAGTTCATCTGCACCACCGGC
AAGCTGCCCGTGCCCTGGCCCACCCTCGTGACCACCCTGACGTACGGCGTGCAGT
GCTTCAGCCGCTACCCCGACCACATGACCTAGGCGCAACCGCCGTTTCCCCGGCC
GGAGAGGTACTCACCGGAGGGGAGAGCCGGTGAGGCTACCGTGCCCCAGGTGA
GAAGGTGGTGCCTTCGGGCCTCCCTCGACCGCTCGCGCtttttttctagaggtacccgttacataactt
acggtaaatggcccgcctggctgaccgcccaacgacccccgcccattgacgtcaatagtaacgccaatagggactttccattgacgtc
aatgggtggagtatttacggtaaactgcccacttggcagtacatcaagtgtatcatatgccaagtacgccccctattgacgtcaatgacg
gtaaatggcccgcctggcattgtgcccagtacatgaccttatgggactttcctacttggcagtacatctacgtattagtcatcgctattacc
atggtcgaggtgagccccacgttctgcttcactctccccatctcccccccctccccacccccaattttgtatttatttattttttaattattttgtg
cagcgatgggggcgggggggggggggggcgcgcgccaggcggggcggggcggggcgaggggcggggcggggcgaggc
ggagaggtgcggcggcagccaatcagagcggcgcgctccgaaagtttccttttatggcgaggcggcggcggcggcggccctataa
aaagcgaagcgcgcggcgggcgggagtcgctgcgacgctgccttcgccccgtgccccgctccgccgccgcctcgcgccgcccg
ccccggctctgactgaccgcgttactcccacaggtgagcgggcgggacggcccttctcctccgggctgtaattagctgagcaagagg
taagggtttaagggatggttggttggtggggtattaatgtttaattacctggagcacctgcctgaaatcactttttttcaggttggaccggtg
ccaccatggactataaggaccacgacggagactacaaggatcatgatattgattacaaagacgatgacgataagatggccccaaaga
agaagcggaaggtcggtatccacggagtcccagcagccgacaagaagtacagcatcggcctggacatcggcaccaactctgtggg
ctgggccgtgatcaccgacgagtacaaggtgcccagcaagaaattcaaggtgctgggcaacaccgaccggcacagcatcaagaag
aacctgatcggagccctgctgttcgacagcggcgaaacagccgaggccacccggctgaagagaaccgccagaagaagatacacc
agacggaagaaccggatctgctatctgcaagagatcttcagcaacgagatggccaaggtggacgacagcttcttccacagactggaa
gagtccttcctggtggaagaggataagaagcacgagcggcaccccatcttcggcaacatcgtggacgaggtggcctaccacgagaa
gtaccccaccatctaccacctgagaaagaaactggtggacagcaccgacaaggccgacctgcggctgatctatctggccctggccc
acatgatcaagttccggggccacttcctgatcgagggcgacctgaaccccgacaacagcgacgtggacaagctgttcatccagctgg
tgcagacctacaaccagctgttcgaggaaaaccccatcaacgccagcggcgtggacgccaaggccatcctgtctgccagactgagc
aagagcagacggctggaaaatctgatcgcccagctgcccggcgagaagaagaatggcctgttcggaaacctgattgccctgagcct
gggcctgacccccaacttcaagagcaacttcgacctggccgaggatgccaaactgcagctgagcaaggacacctacgacgacgac
ctggacaacctgctggcccagatcggcgaccagtacgccgacctgtttctggccgccaagaacctgtccgacgccatcctgctgagc
gacatcctgagagtgaacaccgagatcaccaaggcccccctgagcgcctctatgatcaagagatacgacgagcaccaccaggacct
gaccctgctgaaagctctcgtgcggcagcagctgcctgagaagtacaaagagattttcttcgaccagagcaagaacggctacgccgg
ctacattgacggcggagccagccaggaagagttctacaagttcatcaagcccatcctggaaaagatggacggcaccgaggaactgc
tcgtgaagctgaacagagaggacctgctgcggaagcagcggaccttcgacaacggcagcatcccccaccagatccacctgggaga
gctgcacgccattctgcggcggcaggaagatttttacccattcctgaaggacaaccgggaaaagatcgagaagatcctgaccttccgc
atcccctactacgtgggccctctggccaggggaaacagcagattcgcctggatgaccagaaagagcgaggaaaccatcaccccctg
gaacttcgaggaagtggtggacaagggcgcttccgcccagagcttcatcgagcggatgaccaacttcgataagaacctgcccaacg
agaaggtgctgcccaagcacagcctgctgtacgagtacttcaccgtgtataacgagctgaccaaagtgaaatacgtgaccgagggaa
tgagaaagcccgccttcctgagcggcgagcagaaaaaggccatcgtggacctgctgttcaagaccaaccggaaagtgaccgtgaa
gcagctgaaagaggactacttcaagaaaatcgagtgcttcgactccgtggaaatctccggcgtggaagatcggttcaacgcctccctg
ggcacataccacgatctgctgaaaattatcaaggacaaggacttcctggacaatgaggaaaacgaggacattctggaagatatcgtgc
tgaccctgacactgtttgaggacagagagatgatcgaggaacggctgaaaacctatgcccacctgttcgacgacaaagtgatgaagc
agctgaagcggcggagatacaccggctggggcaggctgagccggaagctgatcaacggcatccgggacaagcagtccggcaag
acaatcctggatttcctgaagtccgacggcttcgccaacagaaacttcatgcagctgatccacgacgacagcctgacctttaaagagga
catccagaaagcccaggtgtccggccagggcgatagcctgcacgagcacattgccaatctggccggcagccccgccattaagaag
ggcatcctgcagacagtgaaggtggtggacgagctcgtgaaagtgatgggccggcacaagcccgagaacatcgtgatcgaaatgg
ccagagagaaccagaccacccagaagggacagaagaacagccgcgagagaatgaagcggatcgaagagggcatcaaagagct
gggcagccagatcctgaaagaacaccccgtggaaaacacccagctgcagaacgagaagctgtacctgtactacctgcagaatggg
cgggatatgtacgtggaccaggaactggacatcaaccggctgtccgactacgatgtggaccatatcgtgcctcagagctttctgaagg
acgactccatcgacaacaaggtgctgaccagaagcgacaagaaccggggcaagagcgacaacgtgccctccgaagaggtcgtga
agaagatgaagaactactggcggcagctgctgaacgccaagctgattacccagagaaagttcgacaatctgaccaaggccgagaga
ggcggcctgagcgaactggataaggccggcttcatcaagagacagctggtggaaacccggcagatcacaaagcacgtggcacag
atcctggactcccggatgaacactaagtacgacgagaatgacaagctgatccgggaagtgaaagtgatcaccctgaagtccaagctg

gtgtccgatttccggaaggatttccagttttacaaagtgcgcgagatcaacaactaccaccacgcccacgacgcctacctgaacgccgt
cgtgggaaccgccctgatcaaaaagtaccctaagctggaaagcgagttcgtgtacggcgactacaaggtgtacgacgtgcggaaga
tgatcgccaagagcgagcaggaaatcggcaaggctaccgccaagtacttcttctacagcaacatcatgaactttttcaagaccgagatt
accctggccaacggcgagatccggaagcggcctctgatcgagacaaacggcgaaaccggggagatcgtgtgggataagggccgg
gattttgccaccgtgcggaaagtgctgagcatgccccaagtgaatatcgtgaaaaagaccgaggtgcagacaggcggcttcagcaaa
gagtctatcctgcccaagaggaacagcgataagctgatcgccagaaagaaggactgggaccctaagaagtacggcggcttcgaca
gccccaccgtggcctattctgtgctggtggtggccaaagtggaaaagggcaagtccaagaaactgaagagtgtgaaagagctgctg
gggatcaccatcatggaaagaagcagcttcgagaagaatcccatcgactttctggaagccaagggctacaaagaagtgaaaaagga
cctgatcatcaagctgcctaagtactccctgttcgagctggaaaacggccggaagagaatgctggcctctgccggcgaactgcagaa
gggaaacgaactggccctgccctccaaatatgtgaacttcctgtacctggccagccactatgagaagctgaagggctcccccgagga
taatgagcagaaacagctgtttgtggaacagcacaagcactacctggacgagatcatcgagcagatcagcgagttctccaagagagt
gatcctggccgacgctaatctggacaaagtgctgtccgcctacaacaagcaccgggataagcccatcagagagcaggccgagaata
tcatccacctgtttaccctgaccaatctgggagcccctgccgccttcaagtactttgacaccaccatcgaccggaagaggtacaccagc
accaaagaggtgctggacgccaccctgatccaccagagcatcaccggcctgtacgagacacggatcgacctgtctcagctgggagg
cgacaaaaggccggcggccacgaaaaaggccggccaggcaaaaaagaaaaagcttgagggcagaggaagtctgctaacatgcg
gtgacgtggaggagaatcccggccctgctagcatggtgagcaagggcgaggaggataacatggccatcatcaaggagttcatgcgc
ttcaaggtgcacatggagggctccgtgaacggccacgagttcgagatcgagggcgagggcgagggccgcccctacgagggcacc
cagaccgccaagctgaaggtgaccaagggtggccccctgcccttcgcctgggacatcctgtcccctcagttcatgtacggctccaag
gcctacgtgaagcaccccgccgacatccccgactacttgaagctgtccttccccgagggcttcaagtgggagcgcgtgatgaacttcg
aggacggcggcgtggtgaccgtgacccaggactcctccctgcaggacggcgagttcatctacaaggtgaagctgcgcggcaccaa
cttcccctccgacggccccgtaatgcagaagaagaccatgggctgggaggcctcctccgagcggatgtaccccgaggacggcgcc
ctgaagggcgagatcaagcagaggctgaagctgaaggacggcggccactacgacgctgaggtcaagaccacctacaaggccaag
aagcccgtgcagctgcccggcgcctacaacgtcaacatcaagttggacatcacctcccacaacgaggactacaccatcgtggaaca
gtacgaacgcgccgagggccgccactccaccggcggcatggacgagctgtacaagctcgaggcaacaaacttctcactactcaaac
aagcaggtgacgtggaggagaatcccgggcctcttaagATGACGGCAAAACTGGAAAGCCACGTGCCT
GCTGCACCACCAGTCTCAGCGGAAGCGCCGGCACCGACGCGACCTGACGCTGCT
AAGCAGGAGGCTCGACGAGCCCACCATGAAGCCCTGAGACTGCGGTGGAAAGC
AATCGAGGAAGCTGGGGGCACTGACGCCTGGGTACGACAGCAGTTGGTTGCGAA
AGGCGTAGCGGCGGAAGAGGTGGACTTCGAAAGTCTCTCCGACAAGCAAAAAGC
GGCTTGGAAAGAGAAAAAAAAAGCTGAGGCAACCGAGCGGAGGGCTCAAAAAC
GACTTGCCTGGGAAGCCTGGAAAGCGACCCATATTCATCATTTGGGCGTTGGAGT
TCACTGGGACGAAGCCGGCGGGCCTGACAAGTTCGATGTGGCAGGGAGGGAAG
AAAGGGCAAAGGCAAACGGGTTGCCCGAAGGTCTTGACAGCGTAGAAGCATTGG
CGAAAGCGCTGGGAATCTCAGTATCCCGATTGCGCTGGTTCTCTTTTCACAGGGA
AGTTGATACAGGGACACACTACCAAACTTGGGAGATCCCAAAGAGGGATGGGGG
CAAACGCACCCTCACGGCCCCGAAAAGGGAGCTTAAGGCAGTACAAAGGTGGGT
ACTCGCCAATGTGGTGGAAAGACTTCCCGTACACGGAGCTGCTCATGGGTTTGTT
GCAGGACGCTCCATACTCACTAATGCGCTCGCTCACCAAGGAGCTGACGTGGTTG
TTAAAGTTGATATGAAAGATTTTTTTCCCAGTGTCACATGGCCTCGAGTTAAAGG
CCTGCTCAGAAAGGGAGGTTTGCCTGAAAACTTGGCGACCCTGCTCGCCCTTCTG
AGTACCGAAGCTCCACGGGAAGTAGTCCGATTCAGAGGGGAAACGCTGTACGTC
GCTAAGGGCCCACGCGCTCTGCCACAGGGCGCGCCTACCAGTCCTGCGCTTACTA
ACGCATTGTGTCTTAGGTTGGATAAGCGGTTGAGCGCGCTGAGTAAGCGGTTGGG
TTTCACATACACCAGATACGCTGACGATCTTACATTTAGTTGGAGGCGCGCTAAA
AAATCTCGCCAGAAAGAGTTGCCGCTGGCTGATGCTCCCGTCGCGCTTCTCCTGG
CCCGGGTTAAGGGAGTGTTGGAAGCAGAAGGTTTCACTTTGCATCCTGATAAGA
CCAGGGTTCAGCGGAAGGGAAGCCGGCAACGAGTGACCGGCCTCGTGGTTAATG
AAGCGCCCGAAGGAGTGCCAGGTGCCCGAGTTCCCCGAGATGTTGTTCGGAGAT
TGCGAGCGGCAATACACAATAGAGAGCAAGGTAAACCAGGGCCGACTGGAGAA
ACGTTGGAACAACTCAAAGGCTTGGCAGCTTTTCTGCACATGACAGACGCTGAA

AAAGGAAGAGCGTTTCTGAGAAGACTTGAGGCCCTGGAGAAGAGGCAAACTGC
Gtaggaattcctagagctcgctgatcagcctcgactgtgccttctagttgccagccatctgttgtttgcccctcccccgtgccttccttgac
cctggaaggtgccactcccactgtcctttcctaataaaatgaggaaattgcatcgcattgtctgagtaggtgtcattctattctggggggtg
gggtggggcaggacagcaagggggaggattgggaagagaatagcaggcatgctggggagcggccgcaggaacccctagtgatg
gagttggccactccctctctgcgcgctcgctcgctcactgaggccgggcgaccaaaggtcgcccgacgcccgggctttgcccgggc
ggcctcagtgagcgagcgagcgcgcagctgcctgcaggggcgcctgatgcggtattttctccttacgcatctgtgcggtatttcacacc
gcatacgtcaaagcaaccatagtacgcgccctgtagcggcgcattaagcgcggcgggtgtggtggttacgcgcagcgtgaccgcta
cacttgccagcgccctagcgcccgctcctttcgctttcttcccttcctttctcgccacgttcgccggctttccccgtcaagctctaaatcgg
gggctccctttagggttccgatttagtgctttacggcacctcgaccccaaaaaacttgatttgggtgatggttcacgtagtgggccatcgc
cctgatagacggtttttcgccctttgacgttggagtccacgttctttaatagtggactcttgttccaaactggaacaacactcaaccctatct
cgggctattcttttgatttataagggattttgccgatttcggcctattggttaaaaaatgagctgatttaacaaaaatttaacgcgaattttaac
aaaatattaacgtttacaattttatggtgcactctcagtacaatctgctctgatgccgcatagttaagccagccccgacacccgccaacac
ccgctgacgcgccctgacgggcttgtctgctcccggcatccgcttacagacaagctgtgaccgtctccgggagctgcatgtgtcagag
gttttcaccgtcatcaccgaaacgcgcgagacgaaagggcctcgtgatacgcctatttttataggttaatgtcatgataataatggtttctta
gacgtcaggtggcacttttcggggaaatgtgcgcggaacccctatttgtttatttttctaaatacattcaaatatgtatccgctcatgagaca
ataaccctgataaatgcttcaataatattgaaaaaggaagagtatgagtattcaacatttccgtgtcgcccttattcccttttttgcggcatttt
gccttcctgtttttgctcacccagaaacgctggtgaaagtaaaagatgctgaagatcagttgggtgcacgagtgggttacatcgaactgg
atctcaacagcggtaagatccttgagagttttcgccccgaagaacgttttccaatgatgagcacttttaaagttctgctatgtggcgcggta
ttatcccgtattgacgccgggcaagagcaactcggtcgccgcatacactattctcagaatgacttggttgagtactcaccagtcacagaa
aagcatcttacggatggcatgacagtaagagaattatgcagtgctgccataaccatgagtgataacactgcggccaacttacttctgaca
acgatcggaggaccgaaggagctaaccgcttttttgcacaacatgggggatcatgtaactcgccttgatcgttgggaaccggagctga
atgaagccataccaaacgacgagcgtgacaccacgatgcctgtagcaatggcaacaacgttgcgcaaactattaactggcgaactac
ttactctagcttcccggcaacaattaatagactggatggaggcggataaagttgcaggaccacttctgcgctcggcccttccggctggc
tggtttattgctgataaatctggagccggtgagcgtggaagccgcggtatcattgcagcactggggccagatggtaagccctcccgtat
cgtagttatctacacgacggggagtcaggcaactatggatgaacgaaatagacagatcgctgagataggtgcctcactgattaagcatt
ggtaactgtcagaccaagtttactcatatatactttagattgatttaaaacttcatttttaatttaaaaggatctaggtgaagatcctttttgataa
tctcatgaccaaaatcccttaacgtgagttttcgttccactgagcgtcagaccccgtagaaaagatcaaaggatcttcttgagatcctttttt
tctgcgcgtaatctgctgcttgcaaacaaaaaaaccaccgctaccagcggtggtttgtttgccggatcaagagctaccaactctttttccg
aaggtaactggcttcagcagagcgcagataccaaatactgtccttctagtgtagccgtagttaggccaccacttcaagaactctgtagca
ccgcctacatacctcgctctgctaatcctgttaccagtggctgctgccagtggcgataagtcgtgtcttaccgggttggactcaagacga
tagttaccggataaggcgcagcggtcgggctgaacggggggttcgtgcacacagcccagcttggagcgaacgacctacaccgaac
tgagatacctacagcgtgagctatgagaaagcgccacgcttcccgaagggagaaaggcggacaggtatccggtaagcggcagggt
cggaacaggagagcgcacgagggagcttccagggggaaacgcctggtatctttatagtcctgtcgggtttcgccacctctgacttgag
cgtcgatttttgtgatgctcgtcaggggggcggagcctatggaaaaacgccagcaacgcggcctttttacggttcctggccttttgctgg
ccttttgctcacatgt

**pBZ194-pU6-sgBFP-Hn_CBh-sv40NLS-Cas9-NLS-T2A-mCherry-P2A-Sa163RT**

gagggcctatttcccatgattccttcatatttgcatatacgatacaaggctgttagagagataattggaattaatttgactgtaaacacaaag
atattagtacaaaatacgtgacgtagaaagtaataatttcttgggtagtttgcagttttaaaattatgttttaaaatggactatcatatgcttacc
gtaacttgaaagtatttcgatttcttggctttatatatcttgtggaaaggacgaaaCACCGCTGAAGCACTGCACGCC
ATgttttagagctagaaatagcaagttaaaataaggctagtccgttatcaacttgaaaaagtggcaccgagtcggtgcCGTACG
GTGCGAGCGACCGAGAGAGGTCCCAAGCCATCAGCCTCAGCGCCTCGAGCGCGA
GAGCGGCGTTGCGCCGCTCTGGTTGAATTGCAGGACACTCTCCGCAAGGTAGCCT
GTTCTTGGCTCTCTTCCCTCCGGTGAGTACCTCTCCGGCCGGGGAGCTGAACCAA
CGAACTAGTTCATGTGGTCGGGGTAGCGGCTGAAGCACTGCACGCCGTACGTCA
GGGTGGTCACGAGGGTGGGCCAGGGCACGGGCAGCTTGCCGGTGGTGCAGATGA
ACTTCAGGGTCAGCTTGCCGTAGGTGGCCCTAGGCGCAACCGCCGTTTCCCCGGC
CGGAGAGGTACTCACCGGAGGGGAGAGCCGGTGAGGCTACCGTGCCCCAGGTGA
GAAGGTGGTGCCTTCGGGCCTCCCTCGACCGCTCGCGCtttttttctagaggtacccgttacataactt
acggtaaatggcccgcctggctgaccgcccaacgacccccgcccattgacgtcaataatgacgtatgttcccatagtaacgccaatagggactttccattgacgtc

aatgggtggagtatttacggtaaactgcccacttggcagtacatcaagtgtatcatatgccaagtacgccccctattgacgtcaatgacg
gtaaatggcccgcctggcattgtgcccagtacatgaccttatgggactttcctacttggcagtacatctacgtattagtcatcgctattacc
atggtcgaggtgagccccacgttctgcttcactctccccatctcccccccctccccacccccaattttgtatttatttattttttaattattttgtg
cagcgatgggggcgggggggggggggggggcgcgcgccaggcggggcggggcggggcgaggggcggggcggggcgaggc
ggagaggtgcggcggcagccaatcagagcggcgcgctccgaaagtttccttttatggcgaggcggcggcggcggcggccctataa
aaagcgaagcgcgcggcgggcgggagtcgctgcgacgctgccttcgccccgtgccccgctccgccgccgcctcgcgccgcccg
ccccggctctgactgaccgcgttactcccacaggtgagcgggcgggacggcccttctcctccgggctgtaattagctgagcaagagg
taagggtttaagggatggttggttggtggggtattaatgtttaattacctggagcacctgcctgaaatcactttttttcaggttggaccggtg
ccaccatggactataaggaccacgacggagactacaaggatcatgatattgattacaaagacgatgacgataagatggccccaaaga
agaagcggaaggtcggtatccacggagtcccagcagccgacaagaagtacagcatcggcctggacatcggcaccaactctgtggg
ctgggccgtgatcaccgacgagtacaaggtgcccagcaagaaattcaaggtgctgggcaacaccgaccggcacagcatcaagaag
aacctgatcggagccctgctgttcgacagcggcgaaacagccgaggccacccggctgaagagaaccgccagaagaagatacacc
agacggaagaaccggatctgctatctgcaagagatcttcagcaacgagatggccaaggtggacgacagcttcttccacagactggaa
gagtccttcctggtggaagaggataagaagcacgagcggcaccccatcttcggcaacatcgtggacgaggtggcctaccacgagaa
gtaccccaccatctaccacctgagaaagaaactggtggacagcaccgacaaggccgacctgcggctgatctatctggccctggccc
acatgatcaagttccggggccacttcctgatcgagggcgacctgaaccccgacaacagcgacgtggacaagctgttcatccagctgg
tgcagacctacaaccagctgttcgaggaaaaccccatcaacgccagcggcgtggacgccaaggccatcctgtctgccagactgagc
aagagcagacggctggaaaatctgatcgcccagctgcccggcgagaagaagaatggcctgttcggaaacctgattgccctgagcct
gggcctgacccccaacttcaagagcaacttcgacctggccgaggatgccaaactgcagctgagcaaggacacctacgacgacgac
ctggacaacctgctggcccagatcggcgaccagtacgccgacctgtttctggccgccaagaacctgtccgacgccatcctgctgagc
gacatcctgagagtgaacaccgagatcaccaaggcccccctgagcgcctctatgatcaagagatacgacgagcaccaccaggacct
gaccctgctgaaagctctcgtgcggcagcagctgcctgagaagtacaaagagattttcttcgaccagagcaagaacggctacgccgg
ctacattgacggcggagccagccaggaagagttctacaagttcatcaagcccatcctggaaaagatggacggcaccgaggaactgc
tcgtgaagctgaacagagaggacctgctgcggaagcagcggaccttcgacaacggcagcatcccccaccagatccacctgggaga
gctgcacgccattctgcggcggcaggaagatttttacccattcctgaaggacaaccgggaaaagatcgagaagatcctgaccttccgc
atcccctactacgtgggccctctggccaggggaaacagcagattcgcctggatgaccagaaagagcgaggaaaccatcaccccctg
gaacttcgaggaagtggtggacaagggcgcttccgcccagagcttcatcgagcggatgaccaacttcgataagaacctgcccaacg
agaaggtgctgcccaagcacagcctgctgtacgagtacttcaccgtgtataacgagctgaccaaagtgaaatacgtgaccgagggaa
tgagaaagcccgccttcctgagcggcgagcagaaaaaggccatcgtggacctgctgttcaagaccaaccggaaagtgaccgtgaa
gcagctgaaagaggactacttcaagaaaatcgagtgcttcgactccgtggaaatctccggcgtggaagatcggttcaacgcctccctg
ggcacataccacgatctgctgaaaattatcaaggacaaggacttcctggacaatgaggaaaacgaggacattctggaagatatcgtgc
tgaccctgacactgtttgaggacagagagatgatcgaggaacggctgaaaacctatgcccacctgttcgacgacaaagtgatgaagc
agctgaagcggcggagatacaccggctggggcaggctgagccggaagctgatcaacggcatccgggacaagcagtccggcaag
acaatcctggatttcctgaagtccgacggcttcgccaacagaaacttcatgcagctgatccacgacgacagcctgacctttaaagagga
catccagaaagcccaggtgtccggccagggcgatagcctgcacgagcacattgccaatctggccggcagccccgccattaagaag
ggcatcctgcagacagtgaaggtggtggacgagctcgtgaaagtgatgggccggcacaagcccgagaacatcgtgatcgaaatgg
ccagagagaaccagaccacccagaagggacagaagaacagccgcgagagaatgaagcggatcgaagagggcatcaaagagct
gggcagccagatcctgaaagaacaccccgtggaaaacacccagctgcagaacgagaagctgtacctgtactacctgcagaatggg
cgggatatgtacgtggaccaggaactggacatcaaccggctgtccgactacgatgtggaccatatcgtgcctcagagctttctgaagg
acgactccatcgacaacaaggtgctgaccagaagcgacaagaaccggggcaagagcgacaacgtgccctccgaagaggtcgtga
agaagatgaagaactactggcggcagctgctgaacgccaagctgattacccagagaaagttcgacaatctgaccaaggccgagaga
ggcggcctgagcgaactggataaggccggcttcatcaagagacagctggtggaaacccggcagatcacaaagcacgtggcacag
atcctggactcccggatgaacactaagtacgacgagaatgacaagctgatccgggaagtgaaagtgatcaccctgaagtccaagctg
gtgtccgatttccggaaggatttccagttttacaaagtgcgcgagatcaacaactaccaccacgcccacgacgcctacctgaacgccgt
cgtgggaaccgccctgatcaaaaagtaccctaagctggaaagcgagttcgtgtacggcgactacaaggtgtacgacgtgcggaaga
tgatcgccaagagcgagcaggaaatcggcaaggctaccgccaagtacttcttctacagcaacatcatgaactttttcaagaccgagatt
accctggccaacggcgagatccggaagcggcctctgatcgagacaaacggcgaaaccggggagatcgtgtgggataagggccgg
gattttgccaccgtgcggaaagtgctgagcatgccccaagtgaatatcgtgaaaaagaccgaggtgcagacaggcggcttcagcaaa
gagtctatcctgcccaagaggaacagcgataagctgatcgccagaaagaaggactgggaccctaagaagtacggcggcttcgaca
gccccaccgtggcctattctgtgctggtggtggccaaagtggaaaagggcaagtccaagaaactgaagagtgtgaaagagctgctg

gggatcaccatcatggaaagaagcagcttcgagaagaatcccatcgactttctggaagccaagggctacaaagaagtgaaaaagga
cctgatcatcaagctgcctaagtactccctgttcgagctggaaaacggccggaagagaatgctggcctctgccggcgaactgcagaa
gggaaacgaactggccctgccctccaaatatgtgaacttcctgtacctggccagccactatgagaagctgaagggctcccccgagga
taatgagcagaaacagctgtttgtggaacagcacaagcactacctggacgagatcatcgagcagatcagcgagttctccaagagagt
gatcctggccgacgctaatctggacaaagtgctgtccgcctacaacaagcaccgggataagcccatcagagagcaggccgagaata
tcatccacctgtttaccctgaccaatctgggagcccctgccgccttcaagtactttgacaccaccatcgaccggaagaggtacaccagc
accaaagaggtgctggacgccaccctgatccaccagagcatcaccggcctgtacgagacacggatcgacctgtctcagctgggagg
cgacaaaaggccggcggccacgaaaaaggccggccaggcaaaaaagaaaaagcttgagggcagaggaagtctgctaacatgcg
gtgacgtggaggagaatcccggccctgctagcatggtgagcaagggcgaggaggataacatggccatcatcaaggagttcatgcgc
ttcaaggtgcacatggagggctccgtgaacggccacgagttcgagatcgagggcgagggcgagggccgcccctacgagggcacc
cagaccgccaagctgaaggtgaccaagggtggcccccctgcccttcgcctgggacatcctgtcccctcagttcatgtacggctccaag
gcctacgtgaagcaccccgccgacatccccgactacttgaagctgtccttccccgagggcttcaagtgggagcgcgtgatgaacttcg
aggacggcggcgtggtgaccgtgacccaggactcctccctgcaggacggcgagttcatctacaaggtgaagctgcgcggcaccaa
cttcccctccgacggccccgtaatgcagaagaagaccatgggctgggaggcctcctccgagcggatgtaccccgaggacggcgcc
ctgaagggcgagatcaagcagaggctgaagctgaaggacggcggccactacgacgctgaggtcaagaccacctacaaggccaag
aagcccgtgcagctgcccggcgcctacaacgtcaacatcaagttggacatcacctcccacaacgaggactacaccatcgtggaaca
gtacgaacgcgccgagggccgccactccaccggcggcatggacgagctgtacaagctcgaggcaacaaacttctcactactcaaac
aagcaggtgacgtggaggagaatcccgggcctcttaagATGACGGCAAAACTGGAAAGCCACGTGCCT
GCTGCACCACCAGTCTCAGCGGAAGCGCCGGCACCGACGCGACCTGACGCTGCT
AAGCAGGAGGCTCGACGAGCCCACCATGAAGCCCTGAGACTGCGGTGGAAAGC
AATCGAGGAAGCTGGGGGCACTGACGCCTGGGTACGACAGCAGTTGGTTGCGAA
AGGCGTAGCGGCGGAAGAGGTGGACTTCGAAAGTCTCTCCGACAAGCAAAAAGC
GGCTTGGAAAGAGAAAAAAAAAGCTGAGGCAACCGAGCGGAGGGCTCAAAAAC
GACTTGCCTGGGAAGCCTGGAAAGCGACCCATATTCATCATTTGGGCGTTGGAGT
TCACTGGGACGAAGCCGGCGGGCCTGACAAGTTCGATGTGGCAGGGAGGGAAG
AAAGGGCAAAGGCAAACGGGTTGCCCGAAGGTCTTGACAGCGTAGAAGCATTGG
CGAAAGCGCTGGGAATCTCAGTATCCCGATTGCGCTGGTTCTCTTTTCACAGGGA
AGTTGATACAGGGACACACTACCAAACTTGGGAGATCCCAAAGAGGGATGGGGG
CAAACGCACCCTCACGGCCCCGAAAAGGGAGCTTAAGGCAGTACAAAGGTGGGT
ACTCGCCAATGTGGTGGAAAGACTTCCCGTACACGGAGCTGCTCATGGGTTTGTT
GCAGGACGCTCCATACTCACTAATGCGCTCGCTCACCAAGGAGCTGACGTGGTTG
TTAAAGTTGATATGAAAGATTTTTTTCCCAGTGTCACATGGCCTCGAGTTAAAGG
CCTGCTCAGAAAGGGAGGTTTGCCTGAAAACTTGGCGACCCTGCTCGCCCTTCTG
AGTACCGAAGCTCCACGGGAAGTAGTCCGATTCAGAGGGGAAACGCTGTACGTC
GCTAAGGGCCCACGCGCTCTGCCACAGGGCGCGCCTACCAGTCCTGCGCTTACTA
ACGCATTGTGTCTTAGGTTGGATAAGCGGTTGAGCGCGCTGAGTAAGCGGTTGGG
TTTCACATACACCAGATACGCTGACGATCTTACATTTAGTTGGAGGCGCGCTAAA
AAATCTCGCCAGAAAGAGTTGCCGCTGGCTGATGCTCCCGTCGCGCTTCTCCTGG
CCCGGGTTAAGGGAGTGTTGGAAGCAGAAGGTTTCACTTTGCATCCTGATAAGA
CCAGGGTTCAGCGGAAGGGAAGCCGGCAACGAGTGACCGGCCTCGTGGTTAATG
AAGCGCCCGAAGGAGTGCCAGGTGCCCGAGTTCCCCGAGATGTTGTTCGGAGAT
TGCGAGCGGCAATACACAATAGAGAGCAAGGTAAACCAGGGCCGACTGGAGAA
ACGTTGGAACAACTCAAAGGCTTGGCAGCTTTTCTGCACATGACAGACGCTGAA
AAAGGAAGAGCGTTTCTGAGAAGACTTGAGGCCCTGGAGAAGAGGCAAACTGC
Gtaggaattcctagagctcgctgatcagcctcgactgtgccttctagttgccagccatctgttgtttgcccctcccccgtgccttccttgac
cctggaaggtgccactcccactgtcctttcctaataaaatgaggaaattgcatcgcattgtctgagtaggtgtcattctattctggggggtg
gggtggggcaggacagcaagggggaggattgggaagagaatagcaggcatgctggggagcggccgcaggaacccctagtgatg
gagttggccactccctctctgcgcgctcgctcgctcactgaggccgggcgaccaaaggtcgcccgacgcccgggctttgcccgggc
ggcctcagtgagcgagcgagcgcgcagctgcctgcaggggcgcctgatgcggtattttctccttacgcatctgtgcggtatttcacacc
gcatacgtcaaagcaaccatagtacgcgccctgtagcggcgcattaagcgcggcgggtgtggtggttacgcgcagcgtgaccgcta

cacttgccagcgccctagcgcccgctcctttcgctttcttcccttcctttctcgccacgttcgccggctttccccgtcaagctctaaatcgg
gggctccctttagggttccgatttagtgctttacggcacctcgaccccaaaaaacttgatttgggtgatggttcacgtagtgggccatcgc
cctgatagacggtttttcgccctttgacgttggagtccacgttctttaatagtggactcttgttccaaactggaacaacactcaaccctatct
cgggctattcttttgatttataagggattttgccgatttcggcctattggttaaaaaatgagctgatttaacaaaaatttaacgcgaattttaac
aaaatattaacgtttacaattttatggtgcactctcagtacaatctgctctgatgccgcatagttaagccagccccgacacccgccaacac
ccgctgacgcgccctgacgggcttgtctgctcccggcatccgcttacagacaagctgtgaccgtctccgggagctgcatgtgtcagag
gttttcaccgtcatcaccgaaacgcgcgagacgaaagggcctcgtgatacgcctatttttataggttaatgtcatgataataatggtttctta
gacgtcaggtggcacttttcggggaaatgtgcgcggaacccctatttgtttatttttctaaatacattcaaatatgtatccgctcatgagaca
ataaccctgataaatgcttcaataatattgaaaaaggaagagtatgagtattcaacatttccgtgtcgcccttattcccttttttgcggcatttt
gccttcctgtttttgctcacccagaaacgctggtgaaagtaaaagatgctgaagatcagttgggtgcacgagtgggttacatcgaactgg
atctcaacagcggtaagatccttgagagttttcgccccgaagaacgttttccaatgatgagcacttttaaagttctgctatgtggcgcggta
ttatcccgtattgacgccgggcaagagcaactcggtcgccgcatacactattctcagaatgacttggttgagtactcaccagtcacagaa
aagcatcttacggatggcatgacagtaagagaattatgcagtgctgccataaccatgagtgataacactgcggccaacttacttctgaca
acgatcggaggaccgaaggagctaaccgcttttttgcacaacatgggggatcatgtaactcgccttgatcgttgggaaccggagctga
atgaagccataccaaacgacgagcgtgacaccacgatgcctgtagcaatggcaacaacgttgcgcaaactattaactggcgaactac
ttactctagcttcccggcaacaattaatagactggatggaggcggataaagttgcaggaccacttctgcgctcggcccttccggctggc
tggtttattgctgataaatctggagccggtgagcgtggaagccgcggtatcattgcagcactggggccagatggtaagccctcccgtat
cgtagttatctacacgacggggagtcaggcaactatggatgaacgaaatagacagatcgctgagataggtgcctcactgattaagcatt
ggtaactgtcagaccaagtttactcatatatactttagattgatttaaaacttcatttttaatttaaaaggatctaggtgaagatcctttttgataa
tctcatgaccaaaatcccttaacgtgagttttcgttccactgagcgtcagaccccgtagaaaagatcaaaggatcttcttgagatccttttt
tctgcgcgtaatctgctgcttgcaaacaaaaaaaccaccgctaccagcggtggtttgtttgccggatcaagagctaccaactctttttccg
aaggtaactggcttcagcagagcgcagataccaaatactgtccttctagtgtagccgtagttaggccaccacttcaagaactctgtagca
ccgcctacatacctcgctctgctaatcctgttaccagtggctgctgccagtggcgataagtcgtgtcttaccgggttggactcaagacga
tagttaccggataaggcgcagcggtcgggctgaacggggggttcgtgcacacagcccagcttggagcgaacgacctacaccgaac
tgagatacctacagcgtgagctatgagaaagcgccacgcttcccgaagggagaaaggcggacaggtatccggtaagcggcagggt
cggaacaggagagcgcacgagggagcttccagggggaaacgcctggtatctttatagtcctgtcgggtttcgccacctctgacttgag
cgtcgatttttgtgatgctcgtcaggggggcggagcctatggaaaaacgccagcaacgcggcctttttacggttcctggccttttgctgg
ccttttgctcacatgt

**pBZ195-pU6-sgBFP-At_CBh-sv40NLS-Cas9-NLS-T2A-mCherry-P2A-Sa163RT**

gagggcctatttcccatgattccttcatatttgcatatacgatacaaggctgttagagagataattggaattaatttgactgtaaacacaaag
atattagtacaaaatacgtgacgtagaaagtaataatttcttgggtagtttgcagttttaaaattatgttttaaaatggactatcatatgcttacc
gtaacttgaaagtatttcgatttcttggctttatatatcttgtggaaaggacgaaaCACCGCTGAAGCACTGCACGCC
ATgttttagagctagaaatagcaagttaaaataaggctagtccgttatcaacttgaaaaagtggcaccgagtcggtgcCGTACG
GTGCGAGCGACCGAGAGAGGTCCCAAGCCATCAGCCTCAGCGCCTCGAGCGCGA
GAGCGGCGTTGCGCCGCTCTGGTTGAATTGCAGGACACTCTCCGCAAGGTAGCCT
GTTCTTGGCTCTCTTCCCTCCGGTGAGTACCTCTCCGGCCGGGGAGCTGAACCAA
CGAACTAGTCCTGAAGTTCATCTGCACCACCGGCAAGCTGCCCGTGCCCTGGCCC
ACCCTCGTGACCACCCTGACGTACGGCGTGCAGTGCTTCAGCCGCTACCCCGACC
ACATGAAGCAGCACGACTTCCTAGGCGCAACCGCCGTTTCCCCGGCCGGAGAGG
TACTCACCGGAGGGGAGAGCCGGTGAGGCTACCGTGCCCCAGGTGAGAAGGTGG
TGCCTTCGGGCCTCCCTCGACCGCTCGCGCtttttttctagaggtacccgttacataacttacggtaaatggcc
cgcctggctgaccgcccaacgacccccgcccattgacgtcaatagtaacgccaatagggactttccattgacgtcaatgggtggagta
tttacggtaaactgcccacttggcagtacatcaagtgtatcatatgccaagtacgccccctattgacgtcaatgacggtaaatggcccgc
ctggcattgtgcccagtacatgaccttatgggactttcctacttggcagtacatctacgtattagtcatcgctattaccatggtcgaggtga
gccccacgttctgcttcactctccccatctcccccccctccccacccccaattttgtatttatttattttttaattattttgtgcagcgatgggg
cgggggggggggggggcgcgcgccaggcggggcggggcggggcgaggggcggggcggggcgaggcggagaggtgcg
gcggcagccaatcagagcggcgcgctccgaaagtttccttttatggcgaggcggcggcggcggcggccctataaaaagcgaagcg
cgcggcgggcgggagtcgctgcgacgctgccttcgccccgtgccccgctccgccgccgcctcgcgccgcccgccccggctctga
ctgaccgcgttactcccacaggtgagcgggcgggacggcccttctcctccgggctgtaattagctgagcaagaggtaagggtttaag

ggatggttggttggtggggtattaatgtttaattacctggagcacctgcctgaaatcactttttttcaggttggaccggtgccaccatggac
tataaggaccacgacggagactacaaggatcatgatattgattacaaagacgatgacgataagatggccccaaagaagaagcggaa
ggtcggtatccacggagtcccagcagccgacaagaagtacagcatcggcctggacatcggcaccaactctgtgggctgggccgtga
tcaccgacgagtacaaggtgcccagcaagaaattcaaggtgctgggcaacaccgaccggcacagcatcaagaagaacctgatcgg
agccctgctgttcgacagcggcgaaacagccgaggccacccggctgaagagaaccgccagaagaagatacaccagacggaaga
accggatctgctatctgcaagagatcttcagcaacgagatggccaaggtggacgacagcttcttccacagactggaagagtccttcct
ggtggaagaggataagaagcacgagcggcaccccatcttcggcaacatcgtggacgaggtggcctaccacgagaagtaccccacc
atctaccacctgagaaagaaactggtggacagcaccgacaaggccgacctgcggctgatctatctggccctggcccacatgatcaag
ttccggggccacttcctgatcgagggcgacctgaaccccgacaacagcgacgtggacaagctgttcatccagctggtgcagacctac
aaccagctgttcgaggaaaaccccatcaacgccagcggcgtggacgccaaggccatcctgtctgccagactgagcaagagcagac
ggctggaaaatctgatcgcccagctgcccggcgagaagaagaatggcctgttcggaaacctgattgccctgagcctgggcctgacc
cccaacttcaagagcaacttcgacctggccgaggatgccaaactgcagctgagcaaggacacctacgacgacgacctggacaacct
gctggcccagatcggcgaccagtacgccgacctgtttctggccgccaagaacctgtccgacgccatcctgctgagcgacatcctgag
agtgaacaccgagatcaccaaggcccccctgagcgcctctatgatcaagagatacgacgagcaccaccaggacctgaccctgctga
aagctctcgtgcggcagcagctgcctgagaagtacaaagagattttcttcgaccagagcaagaacggctacgccggctacattgacg
gcggagccagccaggaagagttctacaagttcatcaagcccatcctggaaaagatggacggcaccgaggaactgctcgtgaagctg
aacagagaggacctgctgcggaagcagcggaccttcgacaacggcagcatcccccaccagatccacctgggagagctgcacgcc
attctgcggcggcaggaagatttttacccattcctgaaggacaaccgggaaaagatcgagaagatcctgaccttccgcatcccctacta
cgtgggccctctggccaggggaaacagcagattcgcctggatgaccagaaagagcgaggaaaccatcaccccctggaacttcgag
gaagtggtggacaagggcgcttccgcccagagcttcatcgagcggatgaccaacttcgataagaacctgcccaacgagaaggtgct
gcccaagcacagcctgctgtacgagtacttcaccgtgtataacgagctgaccaaagtgaaatacgtgaccgagggaatgagaaagcc
cgccttcctgagcggcgagcagaaaaaggccatcgtggacctgctgttcaagaccaaccggaaagtgaccgtgaagcagctgaaa
gaggactacttcaagaaaatcgagtgcttcgactccgtggaaatctccggcgtggaagatcggttcaacgcctccctgggcacatacc
acgatctgctgaaaattatcaaggacaaggacttcctggacaatgaggaaaacgaggacattctggaagatatcgtgctgaccctgac
actgtttgaggacagagagatgatcgaggaacggctgaaaacctatgcccacctgttcgacgacaaagtgatgaagcagctgaagcg
gcggagatacaccggctggggcaggctgagccggaagctgatcaacggcatccgggacaagcagtccggcaagacaatcctgga
tttcctgaagtccgacggcttcgccaacagaaacttcatgcagctgatccacgacgacagcctgacctttaaagaggacatccagaaa
gcccaggtgtccggccagggcgatagcctgcacgagcacattgccaatctggccggcagccccgccattaagaagggcatcctgc
agacagtgaaggtggtggacgagctcgtgaaagtgatgggccggcacaagcccgagaacatcgtgatcgaaatggccagagaga
accagaccacccagaagggacagaagaacagccgcgagagaatgaagcggatcgaagagggcatcaaagagctgggcagcca
gatcctgaaagaacaccccgtggaaaacacccagctgcagaacgagaagctgtacctgtactacctgcagaatgggcgggatatgta
cgtggaccaggaactggacatcaaccggctgtccgactacgatgtggaccatatcgtgcctcagagctttctgaaggacgactccatc
gacaacaaggtgctgaccagaagcgacaagaaccggggcaagagcgacaacgtgccctccgaagaggtcgtgaagaagatgaa
gaactactggcggcagctgctgaacgccaagctgattacccagagaaagttcgacaatctgaccaaggccgagagaggcggcctg
agcgaactggataaggccggcttcatcaagagacagctggtggaaacccggcagatcacaaagcacgtggcacagatcctggactc
ccggatgaacactaagtacgacgagaatgacaagctgatccgggaagtgaaagtgatcaccctgaagtccaagctggtgtccgatttc
cggaaggatttccagttttacaaagtgcgcgagatcaacaactaccaccacgcccacgacgcctacctgaacgccgtcgtgggaacc
gccctgatcaaaaagtaccctaagctggaaagcgagttcgtgtacggcgactacaaggtgtacgacgtgcggaagatgatcgccaag
agcgagcaggaaatcggcaaggctaccgccaagtacttcttctacagcaacatcatgaactttttcaagaccgagattaccctggccaa
cggcgagatccggaagcggcctctgatcgagacaaacggcgaaaccggggagatcgtgtgggataagggccgggattttgccacc
gtgcggaaagtgctgagcatgccccaagtgaatatcgtgaaaaagaccgaggtgcagacaggcggcttcagcaaagagtctatcct
gcccaagaggaacagcgataagctgatcgccagaaagaaggactgggaccctaagaagtacggcggcttcgacagccccaccgt
ggcctattctgtgctggtggtggccaaagtggaaaagggcaagtccaagaaactgaagagtgtgaaagagctgctggggatcaccat
catggaaagaagcagcttcgagaagaatcccatcgactttctggaagccaagggctacaaagaagtgaaaaaggacctgatcatcaa
gctgcctaagtactccctgttcgagctggaaaacggccggaagagaatgctggcctctgccggcgaactgcagaagggaaacgaac
tggccctgccctccaaatatgtgaacttcctgtacctggccagccactatgagaagctgaagggctcccccgaggataatgagcagaa
acagctgtttgtggaacagcacaagcactacctggacgagatcatcgagcagatcagcgagttctccaagagagtgatcctggccga
cgctaatctggacaaagtgctgtccgcctacaacaagcaccgggataagcccatcagagagcaggccgagaatatcatccacctgttt
accctgaccaatctgggagcccctgccgccttcaagtactttgacaccaccatcgaccggaagaggtacaccagcaccaaagaggtg
ctggacgccaccctgatccaccagagcatcaccggcctgtacgagacacggatcgacctgtctcagctgggaggcgacaaaaggc

cggcggccacgaaaaaggccggccaggcaaaaaagaaaaagcttgagggcagaggaagtctgctaacatgcggtgacgtggag
gagaatcccggccctgctagcatggtgagcaagggcgaggaggataacatggccatcatcaaggagttcatgcgcttcaaggtgca
catggagggctccgtgaacggccacgagttcgagatcgagggcgagggcgagggccgcccctacgagggcacccagaccgcca
agctgaaggtgaccaagggtggcccctgcccttcgcctgggacatcctgtcccctcagttcatgtacggctccaaggcctacgtgaa
gcaccccgccgacatccccgactacttgaagctgtccttccccgagggcttcaagtgggagcgcgtgatgaacttcgaggacggcg
gcgtggtgaccgtgacccaggactcctccctgcaggacggcgagttcatctacaaggtgaagctgcgcggcaccaacttcccctccg
acggccccgtaatgcagaagaagaccatgggctgggaggcctcctccgagcggatgtaccccgaggacggcgccctgaagggcg
agatcaagcagaggctgaagctgaaggacggcggccactacgacgctgaggtcaagaccacctacaaggccaagaagcccgtgc
agctgcccggcgcctacaacgtcaacatcaagttggacatcacctcccacaacgaggactacaccatcgtggaacagtacgaacgc
gccgagggccgccactccaccggcggcatggacgagctgtacaagctcgaggcaacaaacttctcactactcaaacaagcaggtg
acgtggaggagaatcccgggcctcttaagATGACGGCAAAACTGGAAAGCCACGTGCCTGCTGCA
CCACCAGTCTCAGCGGAAGCGCCGGCACCGACGCGACCTGACGCTGCTAAGCAG
GAGGCTCGACGAGCCCACCATGAAGCCCTGAGACTGCGGTGGAAAGCAATCGAG
GAAGCTGGGGGCACTGACGCCTGGGTACGACAGCAGTTGGTTGCGAAAGGCGTA
GCGGCGGAAGAGGTGGACTTCGAAAGTCTCTCCGACAAGCAAAAAGCGGCTTGG
AAAGAGAAAAAAAAAGCTGAGGCAACCGAGCGGAGGGCTCAAAAACGACTTGC
CTGGGAAGCCTGGAAAGCGACCCATATTCATCATTTGGGCGTTGGAGTTCACTGG
GACGAAGCCGGCGGGCCTGACAAGTTCGATGTGGCAGGGAGGGAAGAAAGGGC
AAAGGCAAACGGGTTGCCCGAAGGTCTTGACAGCGTAGAAGCATTGGCGAAAGC
GCTGGGAATCTCAGTATCCCGATTGCGCTGGTTCTCTTTTCACAGGGAAGTTGAT
ACAGGGACACACTACCAAACTTGGGAGATCCCAAAGAGGGATGGGGGCAAACG
CACCCTCACGGCCCCGAAAAGGGAGCTTAAGGCAGTACAAAGGTGGGTACTCGC
CAATGTGGTGGAAAGACTTCCCGTACACGGAGCTGCTCATGGGTTTGTTGCAGGA
CGCTCCATACTCACTAATGCGCTCGCTCACCAAGGAGCTGACGTGGTTGTTAAAG
TTGATATGAAAGATTTTTTTCCCAGTGTCACATGGCCTCGAGTTAAAGGCCTGCT
CAGAAAGGGAGGTTTGCCTGAAAACTTGGCGACCCTGCTCGCCCTTCTGAGTACC
GAAGCTCCACGGGAAGTAGTCCGATTCAGAGGGGAAACGCTGTACGTCGCTAAG
GGCCCACGCGCTCTGCCACAGGGCGCGCCTACCAGTCCTGCGCTTACTAACGCAT
TGTGTCTTAGGTTGGATAAGCGGTTGAGCGCGCTGAGTAAGCGGTTGGGTTTCAC
ATACACCAGATACGCTGACGATCTTACATTTAGTTGGAGGCGCGCTAAAAAATCT
CGCCAGAAAGAGTTGCCGCTGGCTGATGCTCCCGTCGCGCTTCTCCTGGCCCGGG
TTAAGGGAGTGTTGGAAGCAGAAGGTTTCACTTTGCATCCTGATAAGACCAGGGT
TCAGCGGAAGGGAAGCCGGCAACGAGTGACCGGCCTCGTGGTTAATGAAGCGCC
CGAAGGAGTGCCAGGTGCCCGAGTTCCCCGAGATGTTGTTCGGAGATTGCGAGC
GGCAATACACAATAGAGAGCAAGGTAAACCAGGGCCGACTGGAGAAACGTTGG
AACAACTCAAAGGCTTGGCAGCTTTTCTGCACATGACAGACGCTGAAAAAGGAA
GAGCGTTTCTGAGAAGACTTGAGGCCCTGGAGAAGAGGCAAACTGCGtaggaattccta
gagctcgctgatcagcctcgactgtgccttctagttgccagccatctgttgtttgcccctcccccgtgccttccttgaccctggaaggtgc
cactcccactgtcctttcctaataaaatgaggaaattgcatcgcattgtctgagtaggtgtcattctattctggggggtggggtggggcag
gacagcaagggggaggattgggaagagaatagcaggcatgctggggagcggccgcaggaacccctagtgatggagttggccact
ccctctctgcgcgctcgctcgctcactgaggccgggcgaccaaaggtcgcccgacgcccgggctttgcccgggcggcctcagtga
gcgagcgagcgcgcagctgcctgcaggggcgcctgatgcggtattttctccttacgcatctgtgcggtatttcacaccgcatacgtcaa
agcaaccatagtacgcgccctgtagcggcgcattaagcgcggcgggtgtggtggttacgcgcagcgtgaccgctacacttgccagc
gccctagcgcccgctcctttcgctttcttcccttcctttctcgccacgttcgccggctttccccgtcaagctctaaatcgggggctcccttta
gggttccgatttagtgctttacggcacctcgaccccaaaaaacttgatttgggtgatggttcacgtagtgggccatcgccctgatagacg
gtttttcgccctttgacgttggagtccacgttctttaatagtggactcttgttccaaactggaacaacactcaaccctatctcgggctattcttt
tgatttataagggattttgccgatttcggcctattggttaaaaaatgagctgatttaacaaaaatttaacgcgaattttaacaaaatattaacgt
ttacaattttatggtgcactctcagtacaatctgctctgatgccgcatagttaagccagccccgacacccgccaacacccgctgacgcgc
cctgacgggcttgtctgctcccggcatccgcttacagacaagctgtgaccgtctccgggagctgcatgtgtcagaggttttcaccgtcat
caccgaaacgcgcgagacgaaagggcctcgtgatacgcctatttttataggttaatgtcatgataataatggtttcttagacgtcaggtgg

cacttttcggggaaatgtgcgcggaacccctatttgtttatttttctaaatacattcaaatatgtatccgctcatgagacaataaccctgataa
atgcttcaataatattgaaaaaggaagagtatgagtattcaacatttccgtgtcgcccttattcccttttttgcggcattttgccttcctgtttttg
ctcacccagaaacgctggtgaaagtaaaagatgctgaagatcagttgggtgcacgagtgggttacatcgaactggatctcaacagcg
gtaagatccttgagagttttcgccccgaagaacgttttccaatgatgagcacttttaaagttctgctatgtggcgcggtattatcccgtattg
acgccgggcaagagcaactcggtcgccgcatacactattctcagaatgacttggttgagtactcaccagtcacagaaaagcatcttac
ggatggcatgacagtaagagaattatgcagtgctgccataaccatgagtgataacactgcggccaacttacttctgacaacgatcgga
ggaccgaaggagctaaccgcttttttgcacaacatgggggatcatgtaactcgccttgatcgttgggaaccggagctgaatgaagcca
taccaaacgacgagcgtgacaccacgatgcctgtagcaatggcaacaacgttgcgcaaactattaactggcgaactacttactctagct
tcccggcaacaattaatagactggatggaggcggataaagttgcaggaccacttctgcgctcggcccttccggctggctggtttattgct
gataaatctggagccggtgagcgtggaagccgcggtatcattgcagcactggggccagatggtaagccctcccgtatcgtagttatct
acacgacggggagtcaggcaactatggatgaacgaaatagacagatcgctgagataggtgcctcactgattaagcattggtaactgtc
agaccaagtttactcatatatactttagattgatttaaaacttcatttttaatttaaaaggatctaggtgaagatcctttttgataatctcatgacc
aaaatcccttaacgtgagttttcgttccactgagcgtcagaccccgtagaaaagatcaaaggatcttcttgagatcctttttttctgcgcgta
atctgctgcttgcaaacaaaaaaaccaccgctaccagcggtggtttgtttgccggatcaagagctaccaactctttttccgaaggtaact
ggcttcagcagagcgcagataccaaatactgtccttctagtgtagccgtagttaggccaccacttcaagaactctgtagcaccgcctac
atacctcgctctgctaatcctgttaccagtggctgctgccagtggcgataagtcgtgtcttaccgggttggactcaagacgatagttaccg
gataaggcgcagcggtcgggctgaacggggggttcgtgcacacagcccagcttggagcgaacgacctacaccgaactgagatacc
tacagcgtgagctatgagaaagcgccacgcttcccgaagggagaaaggcggacaggtatccggtaagcggcagggtcggaacag
gagagcgcacgagggagcttccagggggaaacgcctggtatctttatagtcctgtcgggtttcgccacctctgacttgagcgtcgatttt
tgtgatgctcgtcaggggggcggagcctatggaaaaacgccagcaacgcggcctttttacggttcctggccttttgctggccttttgctc
acatgt

**pBZ196-pU6-sgBFP-An_CBh-sv40NLS-Cas9-NLS-T2A-mCherry-P2A-Sa163RT**

gagggcctatttcccatgattccttcatatttgcatatacgatacaaggctgttagagagataattggaattaatttgactgtaaacacaaag
atattagtacaaaatacgtgacgtagaaagtaataatttcttgggtagtttgcagttttaaaattatgttttaaaatggactatcatatgcttacc
gtaacttgaaagtatttcgatttcttggctttatatatcttgtggaaaggacgaaaCACCGCTGAAGCACTGCACGCC
ATgttttagagctagaaatagcaagttaaaataaggctagtccgttatcaacttgaaaaagtggcaccgagtcggtgcCGTACG
GTGCGAGCGACCGAGAGAGGTCCCAAGCCATCAGCCTCAGCGCCTCGAGCGCGA
GAGCGGCGTTGCGCCGCTCTGGTTGAATTGCAGGACACTCTCCGCAAGGTAGCCT
GTTCTTGGCTCTCTTCCCTCCGGTGAGTACCTCTCCGGCCGGGGAGCTGAACCAA
CGAACTAGTAAGTCGTGCTGCTTCATGTGGTCGGGGTAGCGGCTGAAGCACTGC
ACGCCGTACGTCAGGGTGGTCACGAGGGTGGGCCAGGGCACGGGCAGCTTGCCG
GTGGTGCAGATGAACTTCAGGCCTAGGCGCAACCGCCGTTTCCCCGGCCGGAGA
GGTACTCACCGGAGGGGAGAGCCGGTGAGGCTACCGTGCCCCAGGTGAGAAGGT
GGTGCCTTCGGGCCTCCCTCGACCGCTCGCGCtttttttctagaggtacccgttacataacttacggtaaatg
gcccgcctggctgaccgcccaacgacccccgcccattgacgtcaatagtaacgccaatagggactttccattgacgtcaatgggtgg
agtatttacggtaaactgcccacttggcagtacatcaagtgtatcatatgccaagtacgccccctattgacgtcaatgacggtaaatggc
ccgcctggcattgtgcccagtacatgaccttatgggactttcctacttggcagtacatctacgtattagtcatcgctattaccatggtcgag
gtgagccccacgttctgcttcactctccccatctcccccccctccccacccccaattttgtatttatttattttttaattattttgtgcagcgatg
ggggcggggggggggggggggcgcgcgccaggcggggcggggcggggcgaggggcggggcggggcgaggcggagagg
tgcggcggcagccaatcagagcggcgcgctccgaaagtttccttttatggcgaggcggcggcggcggcggccctataaaaagcga
agcgcgcggcgggcgggagtcgctgcgacgctgccttcgccccgtgccccgctccgccgccgcctcgcgccgcccgccccggct
ctgactgaccgcgttactcccacaggtgagcgggcgggacggcccttctcctccgggctgtaattagctgagcaagaggtaagggttt
aagggatggttggttggtggggtattaatgtttaattacctggagcacctgcctgaaatcactttttttcaggttggaccggtgccaccatg
gactataaggaccacgacggagactacaaggatcatgatattgattacaaagacgatgacgataagatggccccaaagaagaagcg
gaaggtcggtatccacggagtcccagcagccgacaagaagtacagcatcggcctggacatcggcaccaactctgtgggctgggcc
gtgatcaccgacgagtacaaggtgcccagcaagaaattcaaggtgctgggcaacaccgaccggcacagcatcaagaagaacctga
tcggagccctgctgttcgacagcggcgaaacagccgaggccacccggctgaagagaaccgccagaagaagatacaccagacgga
agaaccggatctgctatctgcaagagatcttcagcaacgagatggccaaggtggacgacagcttcttccacagactggaagagtcctt
cctggtggaagaggataagaagcacgagcggcaccccatcttcggcaacatcgtggacgaggtggcctaccacgagaagtacccc

accatctaccacctgagaaagaaactggtggacagcaccgacaaggccgacctgcggctgatctatctggccctggcccacatgatc
aagttccggggccacttcctgatcgagggcgacctgaaccccgacaacagcgacgtggacaagctgttcatccagctggtgcagac
ctacaaccagctgttcgaggaaaaccccatcaacgccagcggcgtggacgccaaggccatcctgtctgccagactgagcaagagca
gacggctggaaaatctgatcgcccagctgcccggcgagaagaagaatggcctgttcggaaacctgattgccctgagcctgggcctg
acccccaacttcaagagcaacttcgacctggccgaggatgccaaactgcagctgagcaaggacacctacgacgacgacctggaca
acctgctggcccagatcggcgaccagtacgccgacctgtttctggccgccaagaacctgtccgacgccatcctgctgagcgacatcc
tgagagtgaacaccgagatcaccaaggcccccctgagcgcctctatgatcaagagatacgacgagcaccaccaggacctgaccctg
ctgaaagctctcgtgcggcagcagctgcctgagaagtacaaagagattttcttcgaccagagcaagaacggctacgccggctacattg
acggcggagccagccaggaagagttctacaagttcatcaagcccatcctggaaaagatggacggcaccgaggaactgctcgtgaa
gctgaacagagaggacctgctgcggaagcagcggaccttcgacaacggcagcatcccccaccagatccacctgggagagctgca
cgccattctgcggcggcaggaagatttttacccattcctgaaggacaaccgggaaaagatcgagaagatcctgaccttccgcatcccc
tactacgtgggccctctggccaggggaaacagcagattcgcctggatgaccagaaagagcgaggaaaccatcaccccctggaactt
cgaggaagtggtggacaagggcgcttccgcccagagcttcatcgagcggatgaccaacttcgataagaacctgcccaacgagaag
gtgctgcccaagcacagcctgctgtacgagtacttcaccgtgtataacgagctgaccaaagtgaaatacgtgaccgagggaatgaga
aagcccgccttcctgagcggcgagcagaaaaaggccatcgtggacctgctgttcaagaccaaccggaaagtgaccgtgaagcagct
gaaagaggactacttcaagaaaatcgagtgcttcgactccgtggaaatctccggcgtggaagatcggttcaacgcctccctgggcaca
taccacgatctgctgaaaattatcaaggacaaggacttcctggacaatgaggaaaacgaggacattctggaagatatcgtgctgaccct
gacactgtttgaggacagagagatgatcgaggaacggctgaaaacctatgcccacctgttcgacgacaaagtgatgaagcagctgaa
gcggcggagatacaccggctggggcaggctgagccggaagctgatcaacggcatccgggacaagcagtccggcaagacaatcct
ggatttcctgaagtccgacggcttcgccaacagaaacttcatgcagctgatccacgacgacagcctgacctttaaagaggacatccag
aaagcccaggtgtccggccagggcgatagcctgcacgagcacattgccaatctggccggcagccccgccattaagaagggcatcct
gcagacagtgaaggtggtggacgagctcgtgaaagtgatgggccggcacaagcccgagaacatcgtgatcgaaatggccagaga
gaaccagaccacccagaagggacagaagaacagccgcgagagaatgaagcggatcgaagagggcatcaaagagctgggcagc
cagatcctgaaagaacaccccgtggaaaacacccagctgcagaacgagaagctgtacctgtactacctgcagaatgggcgggatat
gtacgtggaccaggaactggacatcaaccggctgtccgactacgatgtggaccatatcgtgcctcagagctttctgaaggacgactcc
atcgacaacaaggtgctgaccagaagcgacaagaaccggggcaagagcgacaacgtgccctccgaagaggtcgtgaagaagatg
aagaactactggcggcagctgctgaacgccaagctgattacccagagaaagttcgacaatctgaccaaggccgagagaggcggcct
gagcgaactggataaggccggcttcatcaagagacagctggtggaaacccggcagatcacaaagcacgtggcacagatcctggac
tcccggatgaacactaagtacgacgagaatgacaagctgatccgggaagtgaaagtgatcaccctgaagtccaagctggtgtccgatt
tccggaaggatttccagttttacaaagtgcgcgagatcaacaactaccaccacgcccacgacgcctacctgaacgccgtcgtgggaa
ccgccctgatcaaaaagtaccctaagctggaaagcgagttcgtgtacggcgactacaaggtgtacgacgtgcggaagatgatcgcca
agagcgagcaggaaatcggcaaggctaccgccaagtacttcttctacagcaacatcatgaactttttcaagaccgagattaccctggcc
aacggcgagatccggaagcggcctctgatcgagacaaacggcgaaaccggggagatcgtgtgggataagggccgggattttgcca
ccgtgcggaaagtgctgagcatgccccaagtgaatatcgtgaaaaagaccgaggtgcagacaggcggcttcagcaaagagtctatc
ctgcccaagaggaacagcgataagctgatcgccagaaagaaggactgggaccctaagaagtacggcggcttcgacagccccacc
gtggcctattctgtgctggtggtggccaaagtggaaaagggcaagtccaagaaactgaagagtgtgaaagagctgctggggatcacc
atcatggaaagaagcagcttcgagaagaatcccatcgactttctggaagccaagggctacaaagaagtgaaaaaggacctgatcatc
aagctgcctaagtactccctgttcgagctggaaaacggccggaagagaatgctggcctctgccggcgaactgcagaagggaaacga
actggccctgccctccaaatatgtgaacttcctgtacctggccagccactatgagaagctgaagggctcccccgaggataatgagcag
aaacagctgtttgtggaacagcacaagcactacctggacgagatcatcgagcagatcagcgagttctccaagagagtgatcctggcc
gacgctaatctggacaaagtgctgtccgcctacaacaagcaccgggataagcccatcagagagcaggccgagaatatcatccacct
gtttaccctgaccaatctgggagcccctgccgccttcaagtactttgacaccaccatcgaccggaagaggtacaccagcaccaaaga
ggtgctggacgccaccctgatccaccagagcatcaccggcctgtacgagacacggatcgacctgtctcagctgggaggcgacaaaa
ggccggcggccacgaaaaaggccggccaggcaaaaaagaaaaagcttgagggcagaggaagtctgctaacatgcggtgacgtg
gaggagaatcccggccctgctagcatggtgagcaagggcgaggaggataacatggccatcatcaaggagttcatgcgcttcaaggt
gcacatggagggctccgtgaacggccacgagttcgagatcgagggcgagggcgagggccgcccctacgagggcacccagaccg
ccaagctgaaggtgaccaagggtggccccctgcccttcgcctgggacatcctgtcccctcagttcatgtacggctccaaggcctacgt
gaagcaccccgccgacatccccgactacttgaagctgtccttccccgagggcttcaagtgggagcgcgtgatgaacttcgaggacgg
cggcgtggtgaccgtgacccaggactcctccctgcaggacggcgagttcatctacaaggtgaagctgcgcggcaccaacttcccctc
cgacggccccgtaatgcagaagaagaccatgggctgggaggcctcctccgagcggatgtaccccgaggacggcgccctgaaggg

cgagatcaagcagaggctgaagctgaaggacggcggccactacgacgctgaggtcaagaccacctacaaggccaagaagcccgt
gcagctgcccggcgcctacaacgtcaacatcaagttggacatcacctcccacaacgaggactacaccatcgtggaacagtacgaac
gcgccgagggccgccactccaccggcggcatggacgagctgtacaagctcgaggcaacaaacttctcactactcaaacaagcagg
tgacgtggaggagaatcccgggcctcttaagATGACGGCAAAACTGGAAAGCCACGTGCCTGCTGC
ACCACCAGTCTCAGCGGAAGCGCCGGCACCGACGCGACCTGACGCTGCTAAGCA
GGAGGCTCGACGAGCCCACCATGAAGCCCTGAGACTGCGGTGGAAAGCAATCGA
GGAAGCTGGGGGCACTGACGCCTGGGTACGACAGCAGTTGGTTGCGAAAGGCGT
AGCGGCGGAAGAGGTGGACTTCGAAAGTCTCTCCGACAAGCAAAAAGCGGCTTG
GAAAGAGAAAAAAAAAGCTGAGGCAACCGAGCGGAGGGCTCAAAAACGACTTG
CCTGGGAAGCCTGGAAAGCGACCCATATTCATCATTTGGGCGTTGGAGTTCACTG
GGACGAAGCCGGCGGGCCTGACAAGTTCGATGTGGCAGGGAGGGAAGAAAGGG
CAAAGGCAAACGGGTTGCCCGAAGGTCTTGACAGCGTAGAAGCATTGGCGAAAG
CGCTGGGAATCTCAGTATCCCGATTGCGCTGGTTCTCTTTTCACAGGGAAGTTGA
TACAGGGACACACTACCAAACTTGGGAGATCCCAAAGAGGGATGGGGGCAAAC
GCACCCTCACGGCCCCGAAAAGGGAGCTTAAGGCAGTACAAAGGTGGGTACTCG
CCAATGTGGTGGAAAGACTTCCCGTACACGGAGCTGCTCATGGGTTTGTTGCAGG
ACGCTCCATACTCACTAATGCGCTCGCTCACCAAGGAGCTGACGTGGTTGTTAAA
GTTGATATGAAAGATTTTTTTCCCAGTGTCACATGGCCTCGAGTTAAAGGCCTGC
TCAGAAAGGGAGGTTTGCCTGAAAACTTGGCGACCCTGCTCGCCCTTCTGAGTAC
CGAAGCTCCACGGGAAGTAGTCCGATTCAGAGGGGAAACGCTGTACGTCGCTAA
GGGCCCACGCGCTCTGCCACAGGGCGCGCCTACCAGTCCTGCGCTTACTAACGCA
TTGTGTCTTAGGTTGGATAAGCGGTTGAGCGCGCTGAGTAAGCGGTTGGGTTTCA
CATACACCAGATACGCTGACGATCTTACATTTAGTTGGAGGCGCGCTAAAAAATC
TCGCCAGAAAGAGTTGCCGCTGGCTGATGCTCCCGTCGCGCTTCTCCTGGCCCGG
GTTAAGGGAGTGTTGGAAGCAGAAGGTTTCACTTTGCATCCTGATAAGACCAGG
GTTCAGCGGAAGGGAAGCCGGCAACGAGTGACCGGCCTCGTGGTTAATGAAGCG
CCCGAAGGAGTGCCAGGTGCCCGAGTTCCCCGAGATGTTGTTCGGAGATTGCGA
GCGGCAATACACAATAGAGAGCAAGGTAAACCAGGGCCGACTGGAGAAACGTT
GGAACAACTCAAAGGCTTGGCAGCTTTTCTGCACATGACAGACGCTGAAAAAGG
AAGAGCGTTTCTGAGAAGACTTGAGGCCCTGGAGAAGAGGCAAACTGCGtaggaattc
ctagagctcgctgatcagcctcgactgtgccttctagttgccagccatctgttgtttgcccctcccccgtgccttccttgaccctggaaggt
gccactcccactgtcctttcctaataaaatgaggaaattgcatcgcattgtctgagtaggtgtcattctattctggggggtggggtggggc
aggacagcaagggggaggattgggaagagaatagcaggcatgctggggagcggccgcaggaacccctagtgatggagttggcc
actccctctctgcgcgctcgctcgctcactgaggccgggcgaccaaaggtcgcccgacgcccgggctttgcccgggcggcctcagt
gagcgagcgagcgcgcagctgcctgcaggggcgcctgatgcggtattttctccttacgcatctgtgcggtatttcacaccgcatacgtc
aaagcaaccatagtacgcgccctgtagcggcgcattaagcgcggcgggtgtggtggttacgcgcagcgtgaccgctacacttgcca
gcgccctagcgcccgctcctttcgctttcttcccttcctttctcgccacgttcgccggctttccccgtcaagctctaaatcgggggctccct
ttagggttccgatttagtgctttacggcacctcgaccccaaaaaacttgatttgggtgatggttcacgtagtgggccatcgccctgataga
cggtttttcgccctttgacgttggagtccacgttctttaatagtggactcttgttccaaactggaacaacactcaaccctatctcgggctattc
ttttgatttataagggattttgccgatttcggcctattggttaaaaaatgagctgatttaacaaaaatttaacgcgaattttaacaaaatattaa
cgtttacaattttatggtgcactctcagtacaatctgctctgatgccgcatagttaagccagccccgacacccgccaacacccgctgacg
cgccctgacgggcttgtctgctcccggcatccgcttacagacaagctgtgaccgtctccgggagctgcatgtgtcagaggttttcaccg
tcatcaccgaaacgcgcgagacgaaagggcctcgtgatacgcctatttttataggttaatgtcatgataataatggtttcttagacgtcag
gtggcacttttcggggaaatgtgcgcggaacccctatttgtttatttttctaaatacattcaaatatgtatccgctcatgagacaataaccctg
ataaatgcttcaataatattgaaaaaggaagagtatgagtattcaacatttccgtgtcgcccttattcccttttttgcggcattttgccttcctgt
ttttgctcacccagaaacgctggtgaaagtaaaagatgctgaagatcagttgggtgcacgagtgggttacatcgaactggatctcaaca
gcggtaagatccttgagagttttcgccccgaagaacgttttccaatgatgagcacttttaaagttctgctatgtggcgcggtattatcccgt
attgacgccgggcaagagcaactcggtcgccgcatacactattctcagaatgacttggttgagtactcaccagtcacagaaaagcatct
tacggatggcatgacagtaagagaattatgcagtgctgccataaccatgagtgataacactgcggccaacttacttctgacaacgatcg
gaggaccgaaggagctaaccgcttttttgcacaacatgggggatcatgtaactcgccttgatcgttgggaaccggagctgaatgaagc

cataccaaacgacgagcgtgacaccacgatgcctgtagcaatggcaacaacgttgcgcaaactattaactggcgaactacttactcta
gcttcccggcaacaattaatagactggatggaggcggataaagttgcaggaccacttctgcgctcggcccttccggctggctggtttatt
gctgataaatctggagccggtgagcgtggaagccgcggtatcattgcagcactggggccagatggtaagccctcccgtatcgtagtta
tctacacgacggggagtcaggcaactatggatgaacgaaatagacagatcgctgagataggtgcctcactgattaagcattggtaact
gtcagaccaagtttactcatatatactttagattgatttaaaacttcatttttaatttaaaaggatctaggtgaagatcctttttgataatctcatg
accaaaatcccttaacgtgagttttcgttccactgagcgtcagaccccgtagaaaagatcaaaggatcttcttgagatcctttttttctgcgc
gtaatctgctgcttgcaaacaaaaaaaccaccgctaccagcggtggtttgtttgccggatcaagagctaccaactctttttccgaaggta
actggcttcagcagagcgcagataccaaatactgtccttctagtgtagccgtagttaggccaccacttcaagaactctgtagcaccgcct
acatacctcgctctgctaatcctgttaccagtggctgctgccagtggcgataagtcgtgtcttaccgggttggactcaagacgatagttac
cggataaggcgcagcggtcgggctgaacggggggttcgtgcacacagcccagcttggagcgaacgacctacaccgaactgagata
cctacagcgtgagctatgagaaagcgccacgcttcccgaagggagaaaggcggacaggtatccggtaagcggcagggtcggaac
aggagagcgcacgagggagcttccagggggaaacgcctggtatctttatagtcctgtcgggtttcgccacctctgacttgagcgtcga
tttttgtgatgctcgtcaggggggcggagcctatggaaaaacgccagcaacgcggcctttttacggttcctggccttttgctggccttttg
ctcacatgt

**pBZ197-pU6-sgBFP-Dt_CBh-sv40NLS-Cas9-NLS-T2A-mCherry-P2A-Sa163RT**

gagggcctatttcccatgattccttcatatttgcatatacgatacaaggctgttagagagataattggaattaatttgactgtaaacacaaag
atattagtacaaaatacgtgacgtagaaagtaataatttcttgggtagtttgcagttttaaaattatgttttaaaatggactatcatatgcttacc
gtaacttgaaagtatttcgatttcttggctttatatatcttgtggaaaggacgaaaCACCGCTGAAGCACTGCACGCC
ATgttttagagctagaaatagcaagttaaaataaggctagtccgttatcaacttgaaaaagtggcaccgagtcggtgcCGTACG
GTGCGAGCGACCGAGAGAGGTCCCAAGCCATCAGCCTCAGCGCCTCGAGCGCGA
GAGCGGCGTTGCGCCGCTCTGGTTGAATTGCAGGACACTCTCCGCAAGGTAGCCT
GTTCTTGGCTCTCTTCCCTCCGGTGAGTACCTCTCCGGCCGGGGAGCTGAACCAA
CGAACTAGTCCCTCGTGACCACCCTGACGTACGGCGTGCAGTGCTTCAGCCGCTA
CCCCGACCACATGAAGCAGCACGACTTCTTCAAGTCCGCCATGCCCGAAGGCTA
CGTCCAGGAGCGCCCTAGGCGCAACCGCCGTTTCCCCGGCCGGAGAGGTACTCA
CCGGAGGGGAGAGCCGGTGAGGCTACCGTGCCCCAGGTGAGAAGGTGGTGCCTT
CGGGCCTCCCTCGACCGCTCGCGCtttttttctagaggtacccgttacataacttacggtaaatggcccgcctggct
gaccgcccaacgacccccgcccattgacgtcaatagtaacgccaatagggactttccattgacgtcaatgggtggagtatttacggtaa
actgcccacttggcagtacatcaagtgtatcatatgccaagtacgccccctattgacgtcaatgacggtaaatggcccgcctggcattgt
gcccagtacatgaccttatgggactttcctacttggcagtacatctacgtattagtcatcgctattaccatggtcgaggtgagccccacgtt
ctgcttcactctccccatctcccccccctccccacccccaattttgtatttatttattttttaattattttgtgcagcgatgggggcgggggggg
ggggggggcgcgcgccaggcggggcggggcggggcgaggggcggggcggggcgaggcggagaggtgcggcggcagcc
aatcagagcggcgcgctccgaaagtttccttttatggcgaggcggcggcggcggcggccctataaaaagcgaagcgcgcggcggg
cgggagtcgctgcgacgctgccttcgccccgtgccccgctccgccgccgcctcgcgccgcccgccccggctctgactgaccgcgtt
actcccacaggtgagcgggcgggacggcccttctcctccgggctgtaattagctgagcaagaggtaagggtttaagggatggttggtt
ggtggggtattaatgtttaattacctggagcacctgcctgaaatcactttttttcaggttggaccggtgccaccatggactataaggacca
cgacggagactacaaggatcatgatattgattacaaagacgatgacgataagatggccccaaagaagaagcggaaggtcggtatcc
acggagtcccagcagccgacaagaagtacagcatcggcctggacatcggcaccaactctgtgggctgggccgtgatcaccgacga
gtacaaggtgcccagcaagaaattcaaggtgctgggcaacaccgaccggcacagcatcaagaagaacctgatcggagccctgctgt
tcgacagcggcgaaacagccgaggccacccggctgaagagaaccgccagaagaagatacaccagacggaagaaccggatctgc
tatctgcaagagatcttcagcaacgagatggccaaggtggacgacagcttcttccacagactggaagagtccttcctggtggaagagg
ataagaagcacgagcggcaccccatcttcggcaacatcgtggacgaggtggcctaccacgagaagtaccccaccatctaccacctg
agaaagaaactggtggacagcaccgacaaggccgacctgcggctgatctatctggccctggcccacatgatcaagttccggggcca
cttcctgatcgagggcgacctgaaccccgacaacagcgacgtggacaagctgttcatccagctggtgcagacctacaaccagctgtt
cgaggaaaaccccatcaacgccagcggcgtggacgccaaggccatcctgtctgccagactgagcaagagcagacggctggaaaa
tctgatcgcccagctgcccggcgagaagaagaatggcctgttcggaaacctgattgccctgagcctgggcctgacccccaacttcaa
gagcaacttcgacctggccgaggatgccaaactgcagctgagcaaggacacctacgacgacgacctggacaacctgctggcccag
atcggcgaccagtacgccgacctgtttctggccgccaagaacctgtccgacgccatcctgctgagcgacatcctgagagtgaacacc
gagatcaccaaggcccccctgagcgcctctatgatcaagagatacgacgagcaccaccaggacctgaccctgctgaaagctctcgt

gcggcagcagctgcctgagaagtacaaagagattttcttcgaccagagcaagaacggctacgccggctacattgacggcggagcca
gccaggaagagttctacaagttcatcaagcccatcctggaaaagatggacggcaccgaggaactgctcgtgaagctgaacagagag
gacctgctgcggaagcagcggaccttcgacaacggcagcatccccccaccagatccacctgggagagctgcacgccattctgcggc
ggcaggaagatttttacccattcctgaaggacaaccgggaaaagatcgagaagatcctgaccttccgcatcccctactacgtgggccc
tctggccaggggaaacagcagattcgcctggatgaccagaaagagcgaggaaaccatcacccccctggaacttcgaggaagtggtg
gacaagggcgcttccgcccagagcttcatcgagcggatgaccaacttcgataagaacctgcccaacgagaaggtgctgcccaagca
cagcctgctgtacgagtacttcaccgtgtataacgagctgaccaaagtgaaatacgtgaccgagggaatgagaaagcccgccttcctg
agcggcgagcagaaaaaggccatcgtggacctgctgttcaagaccaaccggaaagtgaccgtgaagcagctgaaagaggactact
tcaagaaaatcgagtgcttcgactccgtggaaatctccggcgtggaagatcggttcaacgcctccctgggcacataccacgatctgct
gaaaattatcaaggacaaggacttcctggacaatgaggaaaacgaggacattctggaagatatcgtgctgaccctgacactgtttgag
gacagagagatgatcgaggaacggctgaaaacctatgcccacctgttcgacgacaaagtgatgaagcagctgaagcggcggagat
acaccggctggggcaggctgagccggaagctgatcaacggcatccgggacaagcagtccggcaagacaatcctggatttcctgaa
gtccgacggcttcgccaacagaaacttcatgcagctgatccacgacgacagcctgacctttaaagaggacatccagaaagcccaggt
gtccggccagggcgatagcctgcacgagcacattgccaatctggccggcagccccgccattaagaagggcatcctgcagacagtg
aaggtggtggacgagctcgtgaaagtgatgggccggcacaagcccgagaacatcgtgatcgaaatggccagagagaaccagacc
acccagaagggacagaagaacagccgcgagagaatgaagcggatcgaagagggcatcaaagagctgggcagccagatcctgaa
agaacaccccgtggaaaacacccagctgcagaacgagaagctgtacctgtactacctgcagaatgggcgggatatgtacgtggacc
aggaactggacatcaaccggctgtccgactacgatgtggaccatatcgtgcctcagagctttctgaaggacgactccatcgacaacaa
ggtgctgaccagaagcgacaagaaccggggcaagagcgacaacgtgccctccgaagaggtcgtgaagaagatgaagaactactg
gcggcagctgctgaacgccaagctgattacccagagaaagttcgacaatctgaccaaggccgagagaggcggcctgagcgaactg
gataaggccggcttcatcaagagacagctggtggaaacccggcagatcacaaagcacgtggcacagatcctggactcccggatgaa
cactaagtacgacgagaatgacaagctgatccgggaagtgaaagtgatcaccctgaagtccaagctggtgtccgatttccggaagga
tttccagttttacaaagtgcgcgagatcaacaactaccaccacgcccacgacgcctacctgaacgccgtcgtgggaaccgccctgatc
aaaaagtaccctaagctggaaagcgagttcgtgtacggcgactacaaggtgtacgacgtgcggaagatgatcgccaagagcgagca
ggaaatcggcaaggctaccgccaagtacttcttctacagcaacatcatgaactttttcaagaccgagattaccctggccaacggcgaga
tccggaagcggcctctgatcgagacaaacggcgaaaccggggagatcgtgtgggataagggccgggattttgccaccgtgcggaa
agtgctgagcatgccccaagtgaatatcgtgaaaaagaccgaggtgcagacaggcggcttcagcaaagagtctatcctgcccaaga
ggaacagcgataagctgatcgccagaaagaaggactgggaccctaagaagtacggcggcttcgacagccccaccgtggcctattct
gtgctggtggtggccaaagtggaaaagggcaagtccaagaaactgaagagtgtgaaagagctgctggggatcaccatcatggaaag
aagcagcttcgagaagaatcccatcgactttctggaagccaagggctacaaagaagtgaaaaaggacctgatcatcaagctgcctaa
gtactccctgttcgagctggaaaacggccggaagagaatgctggcctctgccggcgaactgcagaagggaaacgaactggccctgc
cctccaaatatgtgaacttcctgtacctggccagccactatgagaagctgaagggctcccccgaggataatgagcagaaacagctgttt
gtggaacagcacaagcactacctggacgagatcatcgagcagatcagcgagttctccaagagagtgatcctggccgacgctaatctg
gacaaagtgctgtccgcctacaacaagcaccgggataagcccatcagagagcaggccgagaatatcatccacctgtttaccctgacc
aatctgggagcccctgccgccttcaagtactttgacaccaccatcgaccggaagaggtacaccagcaccaaagaggtgctggacgc
caccctgatccaccagagcatcaccggcctgtacgagacacggatcgacctgtctcagctgggaggcgacaaaaggccggcggcc
acgaaaaaggccggccaggcaaaaaagaaaaagcttgagggcagaggaagtctgctaacatgcggtgacgtggaggagaatccc
ggccctgctagcatggtgagcaagggcgaggaggataacatggccatcatcaaggagttcatgcgcttcaaggtgcacatggaggg
ctccgtgaacggccacgagttcgagatcgagggcgagggcgagggccgcccctacgagggcacccagaccgccaagctgaagg
tgaccaagggtggcccccctgcccttcgcctgggacatcctgtcccctcagttcatgtacggctccaaggcctacgtgaagcaccccgc
cgacatccccgactacttgaagctgtccttccccgagggcttcaagtgggagcgcgtgatgaacttcgaggacggcggcgtggtgac
cgtgacccaggactcctccctgcaggacggcgagttcatctacaaggtgaagctgcgcggcaccaacttcccctccgacggccccgt
aatgcagaagaagaccatgggctgggaggcctcctccgagcggatgtaccccgaggacggcgccctgaagggcgagatcaagca
gaggctgaagctgaaggacggcggccactacgacgctgaggtcaagaccacctacaaggccaagaagcccgtgcagctgcccgg
cgcctacaacgtcaacatcaagttggacatcacctcccacaacgaggactacaccatcgtggaacagtacgaacgcgccgagggcc
gccactccaccggcggcatggacgagctgtacaagctcgaggcaacaaacttctcactactcaaacaagcaggtgacgtggaggag
aatcccgggcctcttaagATGACGGCAAAACTGGAAAGCCACGTGCCTGCTGCACCACCAG
TCTCAGCGGAAGCGCCGGCACCGACGCGACCTGACGCTGCTAAGCAGGAGGCTC
GACGAGCCCACCATGAAGCCCTGAGACTGCGGTGGAAAGCAATCGAGGAAGCTG
GGGGCACTGACGCCTGGGTACGACAGCAGTTGGTTGCGAAAGGCGTAGCGGCGG

AAGAGGTGGACTTCGAAAGTCTCTCCGACAAGCAAAAAGCGGCTTGGAAAGAGA
AAAAAAAAGCTGAGGCAACCGAGCGGAGGGCTCAAAAACGACTTGCCTGGGAA
GCCTGGAAAGCGACCCATATTCATCATTTGGGCGTTGGAGTTCACTGGGACGAA
GCCGGCGGGCCTGACAAGTTCGATGTGGCAGGGAGGGAAGAAAGGGCAAAGGC
AAACGGGTTGCCCGAAGGTCTTGACAGCGTAGAAGCATTGGCGAAAGCGCTGGG
AATCTCAGTATCCCGATTGCGCTGGTTCTCTTTTCACAGGGAAGTTGATACAGGG
ACACACTACCAAACTTGGGAGATCCCAAAGAGGGATGGGGGCAAACGCACCCTC
ACGGCCCCGAAAAGGGAGCTTAAGGCAGTACAAAGGTGGGTACTCGCCAATGTG
GTGGAAAGACTTCCCGTACACGGAGCTGCTCATGGGTTTGTTGCAGGACGCTCCA
TACTCACTAATGCGCTCGCTCACCAAGGAGCTGACGTGGTTGTTAAAGTTGATAT
GAAAGATTTTTTTCCCAGTGTCACATGGCCTCGAGTTAAAGGCCTGCTCAGAAAG
GGAGGTTTGCCTGAAAACTTGGCGACCCTGCTCGCCCTTCTGAGTACCGAAGCTC
CACGGGAAGTAGTCCGATTCAGAGGGGAAACGCTGTACGTCGCTAAGGGCCCAC
GCGCTCTGCCACAGGGCGCGCCTACCAGTCCTGCGCTTACTAACGCATTGTGTCT
TAGGTTGGATAAGCGGTTGAGCGCGCTGAGTAAGCGGTTGGGTTTCACATACAC
CAGATACGCTGACGATCTTACATTTAGTTGGAGGCGCGCTAAAAAATCTCGCCAG
AAAGAGTTGCCGCTGGCTGATGCTCCCGTCGCGCTTCTCCTGGCCCGGGTTAAGG
GAGTGTTGGAAGCAGAAGGTTTCACTTTGCATCCTGATAAGACCAGGGTTCAGC
GGAAGGGAAGCCGGCAACGAGTGACCGGCCTCGTGGTTAATGAAGCGCCCGAA
GGAGTGCCAGGTGCCCGAGTTCCCCGAGATGTTGTTCGGAGATTGCGAGCGGCA
ATACACAATAGAGAGCAAGGTAAACCAGGGCCGACTGGAGAAACGTTGGAACA
ACTCAAAGGCTTGGCAGCTTTTCTGCACATGACAGACGCTGAAAAAGGAAGAGC
GTTTCTGAGAAGACTTGAGGCCCTGGAGAAGAGGCAAACTGCGtaggaattcctagagctcg
ctgatcagcctcgactgtgccttctagttgccagccatctgttgtttgcccctcccccgtgccttccttgaccctggaaggtgccactccca
ctgtcctttcctaataaaatgaggaaattgcatcgcattgtctgagtaggtgtcattctattctggggggtggggtggggcaggacagca
agggggaggattgggaagagaatagcaggcatgctggggagcggccgcaggaacccctagtgatggagttggccactccctctct
gcgcgctcgctcgctcactgaggccgggcgaccaaaggtcgcccgacgcccgggctttgcccgggcggcctcagtgagcgagcg
agcgcgcagctgcctgcaggggcgcctgatgcggtattttctccttacgcatctgtgcggtatttcacaccgcatacgtcaaagcaacc
atagtacgcgccctgtagcggcgcattaagcgcggcgggtgtggtggttacgcgcagcgtgaccgctacacttgccagcgccctag
cgcccgctcctttcgctttcttcccttcctttctcgccacgttcgccggctttccccgtcaagctctaaatcgggggctccctttagggttcc
gatttagtgctttacggcacctcgaccccaaaaaacttgatttgggtgatggttcacgtagtgggccatcgccctgatagacggtttttcg
ccctttgacgttggagtccacgttctttaatagtggactcttgttccaaactggaacaacactcaaccctatctcgggctattcttttgatttat
aagggattttgccgatttcggcctattggttaaaaaatgagctgatttaacaaaaatttaacgcgaattttaacaaaatattaacgtttacaat
tttatggtgcactctcagtacaatctgctctgatgccgcatagttaagccagccccgacacccgccaacacccgctgacgcgccctgac
gggcttgtctgctcccggcatccgcttacagacaagctgtgaccgtctccgggagctgcatgtgtcagaggttttcaccgtcatcaccg
aaacgcgcgagacgaaagggcctcgtgatacgcctatttttataggttaatgtcatgataataatggtttcttagacgtcaggtggcactt
ttcggggaaatgtgcgcggaacccctatttgtttatttttctaaatacattcaaatatgtatccgctcatgagacaataaccctgataaatgctt
caataatattgaaaaaggaagagtatgagtattcaacatttccgtgtcgcccttattcccttttttgcggcattttgccttcctgtttttgctcac
ccagaaacgctggtgaaagtaaaagatgctgaagatcagttgggtgcacgagtgggttacatcgaactggatctcaacagcggtaag
atccttgagagttttcgccccgaagaacgttttccaatgatgagcacttttaaagttctgctatgtggcgcggtattatcccgtattgacgcc
gggcaagagcaactcggtcgccgcatacactattctcagaatgacttggttgagtactcaccagtcacagaaaagcatcttacggatgg
catgacagtaagagaattatgcagtgctgccataaccatgagtgataacactgcggccaacttacttctgacaacgatcggaggaccg
aaggagctaaccgcttttttgcacaacatgggggatcatgtaactcgccttgatcgttgggaaccggagctgaatgaagccataccaaa
cgacgagcgtgacaccacgatgcctgtagcaatggcaacaacgttgcgcaaactattaactggcgaactacttactctagcttcccgg
caacaattaatagactggatggaggcggataaagttgcaggaccacttctgcgctcggcccttccggctggctggtttattgctgataaa
tctggagccggtgagcgtggaagccgcggtatcattgcagcactggggccagatggtaagccctcccgtatcgtagttatctacacga
cggggagtcaggcaactatggatgaacgaaatagacagatcgctgagataggtgcctcactgattaagcattggtaactgtcagacca
agtttactcatatatactttagattgatttaaaacttcatttttaatttaaaaggatctaggtgaagatcctttttgataatctcatgaccaaaatc
ccttaacgtgagttttcgttccactgagcgtcagaccccgtagaaaagatcaaaggatcttcttgagatcctttttttctgcgcgtaatctgc
tgcttgcaaacaaaaaaaccaccgctaccagcggtggtttgtttgccggatcaagagctaccaactctttttccgaaggtaactggcttc

agcagagcgcagataccaaatactgtccttctagtgtagccgtagttaggccaccacttcaagaactctgtagcaccgcctacatacctc
gctctgctaatcctgttaccagtggctgctgccagtggcgataagtcgtgtcttaccgggttggactcaagacgatagttaccggataag
gcgcagcggtcgggctgaacggggggttcgtgcacacagcccagcttggagcgaacgacctacaccgaactgagatacctacagc
gtgagctatgagaaagcgccacgcttcccgaagggagaaaggcggacaggtatccggtaagcggcagggtcggaacaggagag
cgcacgagggagcttccagggggaaacgcctggtatctttatagtcctgtcgggtttcgccacctctgacttgagcgtcgatttttgtgat
gctcgtcaggggggcggagcctatggaaaaacgccagcaacgcggcctttttacggttcctggccttttgctggccttttgctcacatgt

**pBZ198-pU6-sgBFP-Dn_CBh-sv40NLS-Cas9-NLS-T2A-mCherry-P2A-Sa163RT**

gagggcctatttcccatgattccttcatatttgcatatacgatacaaggctgttagagagataattggaattaatttgactgtaaacacaaag
atattagtacaaaatacgtgacgtagaaagtaataatttcttgggtagtttgcagttttaaaattatgttttaaaatggactatcatatgcttacc
gtaacttgaaagtatttcgatttcttggctttatatatcttgtggaaaggacgaaaCACCGCTGAAGCACTGCACGCC
ATgttttagagctagaaatagcaagttaaaataaggctagtccgttatcaacttgaaaaagtggcaccgagtcggtgcCGTACG
GTGCGAGCGACCGAGAGAGGTCCCAAGCCATCAGCCTCAGCGCCTCGAGCGCGA
GAGCGGCGTTGCGCCGCTCTGGTTGAATTGCAGGACACTCTCCGCAAGGTAGCCT
GTTCTTGGCTCTCTTCCCTCCGGTGAGTACCTCTCCGGCCGGGGAGCTGAACCAA
CGAACTAGTGCGCTCCTGGACGTAGCCTTCGGGCATGGCGGACTTGAAGAAGTC
GTGCTGCTTCATGTGGTCGGGGTAGCGGCTGAAGCACTGCACGCCGTACGTCAG
GGTGGTCACGAGGGCCTAGGCGCAACCGCCGTTTCCCCGGCCGGAGAGGTACTC
ACCGGAGGGGAGAGCCGGTGAGGCTACCGTGCCCCAGGTGAGAAGGTGGTGCCT
TCGGGCCTCCCTCGACCGCTCGCGCtttttttctagaggtacccgttacataacttacggtaaatggcccgcctgg
ctgaccgcccaacgacccccgcccattgacgtcaatagtaacgccaatagggactttccattgacgtcaatgggtggagtatttacggt
aaactgcccacttggcagtacatcaagtgtatcatatgccaagtacgccccctattgacgtcaatgacggtaaatggcccgcctggcatt
gtgcccagtacatgaccttatgggactttcctacttggcagtacatctacgtattagtcatcgctattaccatggtcgaggtgagccccac
gttctgcttcactctccccatctcccccccctccccacccccaattttgtatttatttattttttaattattttgtgcagcgatgggggcggggg
gggggggggggcgcgcgccaggcggggcggggcggggcgaggggcggggcggggcgaggcggagaggtgcggcggcag
ccaatcagagcggcgcgctccgaaagtttccttttatggcgaggcggcggcggcggcggccctataaaaagcgaagcgcgcggcg
ggcgggagtcgctgcgacgctgccttcgccccgtgccccgctccgccgccgcctcgcgccgcccgccccggctctgactgaccgc
gttactcccacaggtgagcgggcgggacggcccttctcctccgggctgtaattagctgagcaagaggtaagggtttaagggatggttg
gttggtggggtattaatgtttaattacctggagcacctgcctgaaatcactttttttcaggttggaccggtgccaccatggactataaggac
cacgacggagactacaaggatcatgatattgattacaaagacgatgacgataagatggccccaaagaagaagcggaaggtcggtat
ccacggagtcccagcagccgacaagaagtacagcatcggcctggacatcggcaccaactctgtgggctgggccgtgatcaccgac
gagtacaaggtgcccagcaagaaattcaaggtgctgggcaacaccgaccggcacagcatcaagaagaacctgatcggagccctgc
tgttcgacagcggcgaaacagccgaggccacccggctgaagagaaccgccagaagaagatacaccagacggaagaaccggatct
gctatctgcaagagatcttcagcaacgagatggccaaggtggacgacagcttcttccacagactggaagagtccttcctggtggaaga
ggataagaagcacgagcggcaccccatcttcggcaacatcgtggacgaggtggcctaccacgagaagtaccccaccatctaccacc
tgagaaagaaactggtggacagcaccgacaaggccgacctgcggctgatctatctggccctggcccacatgatcaagttccggggc
cacttcctgatcgagggcgacctgaaccccgacaacagcgacgtggacaagctgttcatccagctggtgcagacctacaaccagctg
ttcgaggaaaaccccatcaacgccagcggcgtggacgccaaggccatcctgtctgccagactgagcaagagcagacggctggaaa
atctgatcgcccagctgcccggcgagaagaagaatggcctgttcggaaacctgattgccctgagcctgggcctgacccccaacttca
agagcaacttcgacctggccgaggatgccaaactgcagctgagcaaggacacctacgacgacgacctggacaacctgctggccca
gatcggcgaccagtacgccgacctgtttctggccgccaagaacctgtccgacgccatcctgctgagcgacatcctgagagtgaacac
cgagatcaccaaggcccccctgagcgcctctatgatcaagagatacgacgagcaccaccaggacctgaccctgctgaaagctctcgt
gcggcagcagctgcctgagaagtacaaagagattttcttcgaccagagcaagaacggctacgccggctacattgacggcggagcca
gccaggaagagttctacaagttcatcaagcccatcctggaaaagatggacggcaccgaggaactgctcgtgaagctgaacagagag
gacctgctgcggaagcagcggaccttcgacaacggcagcatcccccaccagatccacctgggagagctgcacgccattctgcggc
ggcaggaagatttttacccattcctgaaggacaaccgggaaaagatcgagaagatcctgaccttccgcatcccctactacgtgggccc
tctggccaggggaaacagcagattcgcctggatgaccagaaagagcgaggaaaccatcaccccctggaacttcgaggaagtggtg
gacaagggcgcttccgcccagagcttcatcgagcggatgaccaacttcgataagaacctgcccaacgagaaggtgctgcccaagca
cagcctgctgtacgagtacttcaccgtgtataacgagctgaccaaagtgaaatacgtgaccgagggaatgagaaagcccgccttcctg
agcggcgagcagaaaaaggccatcgtggacctgctgttcaagaccaaccggaaagtgaccgtgaagcagctgaaagaggactact

tcaagaaaatcgagtgcttcgactccgtggaaatctccggcgtggaagatcggttcaacgcctccctgggcacataccacgatctgct
gaaaattatcaaggacaaggacttcctggacaatgaggaaaacgaggacattctggaagatatcgtgctgaccctgacactgtttgag
gacagagagatgatcgaggaacggctgaaaacctatgcccacctgttcgacgacaaagtgatgaagcagctgaagcggcggagat
acaccggctggggcaggctgagccggaagctgatcaacggcatccgggacaagcagtccggcaagacaatcctggatttcctgaa
gtccgacggcttcgccaacagaaacttcatgcagctgatccacgacgacagcctgacctttaaagaggacatccagaaagcccaggt
gtccggccagggcgatagcctgcacgagcacattgccaatctggccggcagccccgccattaagaagggcatcctgcagacagtg
aaggtggtggacgagctcgtgaaagtgatgggccggcacaagcccgagaacatcgtgatcgaaatggccagagagaaccagacc
acccagaagggacagaagaacagccgcgagagaatgaagcggatcgaagagggcatcaaagagctgggcagccagatcctgaa
agaacaccccgtggaaaacacccagctgcagaacgagaagctgtacctgtactacctgcagaatgggcgggatatgtacgtggacc
aggaactggacatcaaccggctgtccgactacgatgtggaccatatcgtgcctcagagctttctgaaggacgactccatcgacaacaa
ggtgctgaccagaagcgacaagaaccggggcaagagcgacaacgtgccctccgaagaggtcgtgaagaagatgaagaactactg
gcggcagctgctgaacgccaagctgattacccagagaaagttcgacaatctgaccaaggccgagagaggcggcctgagcgaactg
gataaggccggcttcatcaagagacagctggtggaaacccggcagatcacaaagcacgtggcacagatcctggactcccggatgaa
cactaagtacgacgagaatgacaagctgatccgggaagtgaaagtgatcaccctgaagtccaagctggtgtccgatttccggaagga
tttccagttttacaaagtgcgcgagatcaacaactaccaccacgcccacgacgcctacctgaacgccgtcgtgggaaccgccctgatc
aaaaagtaccctaagctggaaagcgagttcgtgtacggcgactacaaggtgtacgacgtgcggaagatgatcgccaagagcgagca
ggaaatcggcaaggctaccgccaagtacttcttctacagcaacatcatgaactttttcaagaccgagattaccctggccaacggcgaga
tccggaagcggcctctgatcgagacaaacggcgaaaccggggagatcgtgtgggataagggccgggattttgccaccgtgcggaa
agtgctgagcatgccccaagtgaatatcgtgaaaaagaccgaggtgcagacaggcggcttcagcaaagagtctatcctgcccaaga
ggaacagcgataagctgatcgccagaaagaaggactgggaccctaagaagtacggcggcttcgacagccccaccgtggcctattct
gtgctggtggtggccaaagtggaaaagggcaagtccaagaaactgaagagtgtgaaagagctgctggggatcaccatcatggaaag
aagcagcttcgagaagaatcccatcgactttctggaagccaagggctacaaagaagtgaaaaaggacctgatcatcaagctgcctaa
gtactccctgttcgagctggaaaacggccggaagagaatgctggcctctgccggcgaactgcagaagggaaacgaactggccctgc
cctccaaatatgtgaacttcctgtacctggccagccactatgagaagctgaagggctcccccgaggataatgagcagaaacagctgttt
gtggaacagcacaagcactacctggacgagatcatcgagcagatcagcgagttctccaagagagtgatcctggccgacgctaatctg
gacaaagtgctgtccgcctacaacaagcaccgggataagcccatcagagagcaggccgagaatatcatccacctgtttaccctgacc
aatctgggagcccctgccgccttcaagtactttgacaccaccatcgaccggaagaggtacaccagcaccaaagaggtgctggacgc
caccctgatccaccagagcatcaccggcctgtacgagacacggatcgacctgtctcagctgggaggcgacaaaaggccggcggcc
acgaaaaaggccggccaggcaaaaaagaaaaagcttgagggcagaggaagtctgctaacatgcggtgacgtggaggagaatccc
ggccctgctagcatggtgagcaagggcgaggaggataacatggccatcatcaaggagttcatgcgcttcaaggtgcacatggaggg
ctccgtgaacggccacgagttcgagatcgagggcgagggcgagggccgcccctacgagggcacccagaccgccaagctgaagg
tgaccaagggtggcccctgcccttcgcctgggacatcctgtcccctcagttcatgtacggctccaaggcctacgtgaagcaccccgc
cgacatccccgactacttgaagctgtccttccccgagggcttcaagtgggagcgcgtgatgaacttcgaggacggcggcgtggtgac
cgtgacccaggactcctccctgcaggacggcgagttcatctacaaggtgaagctgcgcggcaccaacttcccctccgacggccccgt
aatgcagaagaagaccatgggctgggaggcctcctccgagcggatgtaccccgaggacggcgccctgaagggcgagatcaagca
gaggctgaagctgaaggacggcggccactacgacgctgaggtcaagaccacctacaaggccaagaagcccgtgcagctgcccgg
cgcctacaacgtcaacatcaagttggacatcacctcccacaacgaggactacaccatcgtggaacagtacgaacgcgccgagggcc
gccactccaccggcggcatggacgagctgtacaagctcgaggcaacaaacttctcactactcaaacaagcaggtgacgtggaggag
aatcccgggcctcttaagATGACGGCAAAACTGGAAAGCCACGTGCCTGCTGCACCACCAG
TCTCAGCGGAAGCGCCGGCACCGACGCGACCTGACGCTGCTAAGCAGGAGGCTC
GACGAGCCCACCATGAAGCCCTGAGACTGCGGTGGAAAGCAATCGAGGAAGCTG
GGGGCACTGACGCCTGGGTACGACAGCAGTTGGTTGCGAAAGGCGTAGCGGCGG
AAGAGGTGGACTTCGAAAGTCTCTCCGACAAGCAAAAAGCGGCTTGGAAAGAGA
AAAAAAAAGCTGAGGCAACCGAGCGGAGGGCTCAAAAACGACTTGCCTGGGAA
GCCTGGAAAGCGACCCATATTCATCATTTGGGCGTTGGAGTTCACTGGGACGAA
GCCGGCGGGCCTGACAAGTTCGATGTGGCAGGGAGGGAAGAAAGGGCAAAGGC
AAACGGGTTGCCCGAAGGTCTTGACAGCGTAGAAGCATTGGCGAAAGCGCTGGG
AATCTCAGTATCCCGATTGCGCTGGTTCTCTTTTCACAGGGAAGTTGATACAGGG
ACACACTACCAAACTTGGGAGATCCCAAAGAGGGATGGGGGCAAACGCACCCTC
ACGGCCCCGAAAAGGGAGCTTAAGGCAGTACAAAGGTGGGTACTCGCCAATGTG

GTGGAAAGACTTCCCGTACACGGAGCTGCTCATGGGTTTGTTGCAGGACGCTCCA
TACTCACTAATGCGCTCGCTCACCAAGGAGCTGACGTGGTTGTTAAAGTTGATAT
GAAAGATTTTTTCCCAGTGTCACATGGCCTCGAGTTAAAGGCCTGCTCAGAAAG
GGAGGTTTGCCTGAAAACTTGGCGACCCTGCTCGCCCTTCTGAGTACCGAAGCTC
CACGGGAAGTAGTCCGATTCAGAGGGGAAACGCTGTACGTCGCTAAGGGCCCAC
GCGCTCTGCCACAGGGCGCGCCTACCAGTCCTGCGCTTACTAACGCATTGTGTCT
TAGGTTGGATAAGCGGTTGAGCGCGCTGAGTAAGCGGTTGGGTTTCACATACAC
CAGATACGCTGACGATCTTACATTTAGTTGGAGGCGCGCTAAAAAATCTCGCCAG
AAAGAGTTGCCGCTGGCTGATGCTCCCGTCGCGCTTCTCCTGGCCCGGGTTAAGG
GAGTGTTGGAAGCAGAAGGTTTCACTTTGCATCCTGATAAGACCAGGGTTCAGC
GGAAGGGAAGCCGGCAACGAGTGACCGGCCTCGTGGTTAATGAAGCGCCCGAA
GGAGTGCCAGGTGCCCGAGTTCCCCGAGATGTTGTTCGGAGATTGCGAGCGGCA
ATACACAATAGAGAGCAAGGTAAACCAGGGCCGACTGGAGAAACGTTGGAACA
ACTCAAAGGCTTGGCAGCTTTTCTGCACATGACAGACGCTGAAAAAGGAAGAGC
GTTTCTGAGAAGACTTGAGGCCCTGGAGAAGAGGCAAACTGCGtaggaattcctagagctcg
ctgatcagcctcgactgtgccttctagttgccagccatctgttgtttgcccctcccccgtgccttccttgaccctggaaggtgccactccca
ctgtcctttcctaataaaatgaggaaattgcatcgcattgtctgagtaggtgtcattctattctggggggtggggtggggcaggacagca
agggggaggattgggaagagaatagcaggcatgctggggagcggccgcaggaacccctagtgatggagttggccactccctctct
gcgcgctcgctcgctcactgaggccgggcgaccaaaggtcgcccgacgcccgggctttgcccgggcggcctcagtgagcgagcg
agcgcgcagctgcctgcaggggcgcctgatgcggtattttctccttacgcatctgtgcggtatttcacaccgcatacgtcaaagcaacc
atagtacgcgccctgtagcggcgcattaagcgcggcgggtgtggtggttacgcgcagcgtgaccgctacacttgccagcgccctag
cgcccgctcctttcgctttcttcccttcctttctcgccacgttcgccggctttccccgtcaagctctaaatcgggggctccctttagggttcc
gatttagtgctttacggcacctcgaccccaaaaaacttgatttgggtgatggttcacgtagtgggccatcgccctgatagacggtttttcg
ccctttgacgttggagtccacgttctttaatagtggactcttgttccaaactggaacaacactcaaccctatctcgggctattcttttgatttat
aagggattttgccgatttcggcctattggttaaaaaatgagctgatttaacaaaaatttaacgcgaattttaacaaaatattaacgtttacaat
tttatggtgcactctcagtacaatctgctctgatgccgcatagttaagccagccccgacacccgccaacacccgctgacgcgccctgac
gggcttgtctgctcccggcatccgcttacagacaagctgtgaccgtctccgggagctgcatgtgtcagaggttttcaccgtcatcaccg
aaacgcgcgagacgaaagggcctcgtgatacgcctatttttataggttaatgtcatgataataatggtttcttagacgtcaggtggcacttt
tcggggaaatgtgcgcggaacccctatttgtttatttttctaaatacattcaaatatgtatccgctcatgagacaataaccctgataaatgctt
caataatattgaaaaaggaagagtatgagtattcaacatttccgtgtcgcccttattcccttttttgcggcattttgccttcctgtttttgctcac
ccagaaacgctggtgaaagtaaaagatgctgaagatcagttgggtgcacgagtgggttacatcgaactggatctcaacagcggtaag
atccttgagagttttcgccccgaagaacgttttccaatgatgagcacttttaaagttctgctatgtggcgcggtattatcccgtattgacgcc
gggcaagagcaactcggtcgccgcatacactattctcagaatgacttggttgagtactcaccagtcacagaaaagcatcttacggatgg
catgacagtaagagaattatgcagtgctgccataaccatgagtgataacactgcggccaacttacttctgacaacgatcggaggaccg
aaggagctaaccgcttttttgcacaacatgggggatcatgtaactcgccttgatcgttgggaaccggagctgaatgaagccataccaaa
cgacgagcgtgacaccacgatgcctgtagcaatggcaacaacgttgcgcaaactattaactggcgaactacttactctagcttcccgg
caacaattaatagactggatggaggcggataaagttgcaggaccacttctgcgctcggcccttccggctggctggtttattgctgataaa
tctggagccggtgagcgtggaagccgcggtatcattgcagcactggggccagatggtaagccctcccgtatcgtagttatctacacga
cggggagtcaggcaactatggatgaacgaaatagacagatcgctgagataggtgcctcactgattaagcattggtaactgtcagacca
agtttactcatatatactttagattgatttaaaacttcatttttaatttaaaaggatctaggtgaagatcctttttgataatctcatgaccaaaatc
ccttaacgtgagttttcgttccactgagcgtcagaccccgtagaaaagatcaaaggatcttcttgagatcctttttttctgcgcgtaatctgc
tgcttgcaaacaaaaaaaccaccgctaccagcggtggtttgtttgccggatcaagagctaccaactctttttccgaaggtaactggcttc
agcagagcgcagataccaaatactgtccttctagtgtagccgtagttaggccaccacttcaagaactctgtagcaccgcctacatacctc
gctctgctaatcctgttaccagtggctgctgccagtggcgataagtcgtgtcttaccgggttggactcaagacgatagttaccggataag
gcgcagcggtcgggctgaacggggggttcgtgcacacagcccagcttggagcgaacgacctacaccgaactgagatacctacagc
gtgagctatgagaaagcgccacgcttcccgaagggagaaaggcggacaggtatccggtaagcggcagggtcggaacaggagag
cgcacgagggagcttccagggggaaacgcctggtatctttatagtcctgtcgggtttcgccacctctgacttgagcgtcgatttttgtgat
gctcgtcaggggggcggagcctatggaaaaacgccagcaacgcggcctttttacggttcctggccttttgctggccttttgctcacatgt

**pBZ199-pU6-sgBFP-Ht_CBh-sv40NLS-Cas9-Sa163RT-NLS-T2A-mCherry**

gagggcctatttcccatgattccttcatatttgcatatacgatacaaggctgttagagagataattggaattaatttgactgtaaacacaaag
atattagtacaaaatacgtgacgtagaaagtaataatttcttgggtagtttgcagttttaaaattatgttttaaaatggactatcatatgcttacc
gtaacttgaaagtatttcgatttcttggctttatatatcttgtggaaaggacgaaacaccGCTGAAGCACTGCACGCCAT
gttttagagctagaaatagcaagttaaaataaggctagtccgttatcaacttgaaaaagtggcaccgagtcggtgcCGTACGGT
GCGAGCGACCGAGAGAGGTCCCAAGCCATCAGCCTCAGCGCCTCGAGCGCGAGA
GCGGCGTTGCGCCGCTCTGGTTGAATTGCAGGACACTCTCCGCAAGGTAGCCTGT
TCTTGGCTCTCTTCCCTCCGGTGAGTACCTCTCCGGCCGGGGAGCTGAACCAACG
AACTAGTGCCACCTACGGCAAGCTGACCCTGAAGTTCATCTGCACCACCGGCAA
GCTGCCCGTGCCCTGGCCCACCCTCGTGACCACCCTGACGTACGGCGTGCAGTGC
TTCAGCCGCTACCCCGACCACATGACCTAGGCGCAACCGCCGTTTCCCCGGCCGG
AGAGGTACTCACCGGAGGGGAGAGCCGGTGAGGCTACCGTGCCCCAGGTGAGA
AGGTGGTGCCTTCGGGCCTCCCTCGACCGCTCGCGCtttttttctagaggtacccgttacataacttacg
gtaaatggcccgcctggctgaccgcccaacgacccccgcccattgacgtcaatagtaacgccaatagggactttccattgacgtcaat
gggtggagtatttacggtaaactgcccacttggcagtacatcaagtgtatcatatgccaagtacgccccctattgacgtcaatgacggta
aatggcccgcctggcattgtgcccagtacatgaccttatgggactttcctacttggcagtacatctacgtattagtcatcgctattaccatg
gtcgaggtgagccccacgttctgcttcactctccccatctcccccccctccccacccccaattttgtatttatttattttttaattattttgtgca
gcgatgggggcggggggggggggggggcgcgcgccaggcggggcggggcggggcgaggggcggggcggggcgaggcg
gagaggtgcggcggcagccaatcagagcggcgcgctccgaaagtttccttttatggcgaggcggcggcggcggcggccctataaa
aagcgaagcgcgcggcgggcgggagtcgctgcgacgctgccttcgccccgtgccccgctccgccgccgcctcgcgccgcccgc
cccggctctgactgaccgcgttactcccacaggtgagcgggcgggacggcccttctcctccgggctgtaattagctgagcaagaggt
aagggtttaagggatggttggttggtggggtattaatgtttaattacctggagcacctgcctgaaatcactttttttcaggttggaccggtgc
caccatggactataaggaccacgacggagactacaaggatcatgatattgattacaaagacgatgacgataagatggccccaaagaa
gaagcggaaggtcggtatccacggagtcccagcagccgacaagaagtacagcatcggcctggacatcggcaccaactctgtgggc
tgggccgtgatcaccgacgagtacaaggtgcccagcaagaaattcaaggtgctgggcaacaccgaccggcacagcatcaagaaga
acctgatcggagccctgctgttcgacagcggcgaaacagccgaggccacccggctgaagagaaccgccagaagaagatacacca
gacggaagaaccggatctgctatctgcaagagatcttcagcaacgagatggccaaggtggacgacagcttcttccacagactggaag
agtccttcctggtggaagaggataagaagcacgagcggcaccccatcttcggcaacatcgtggacgaggtggcctaccacgagaag
taccccaccatctaccacctgagaaagaaactggtggacagcaccgacaaggccgacctgcggctgatctatctggccctggcccac
atgatcaagttccggggccacttcctgatcgagggcgacctgaaccccgacaacagcgacgtggacaagctgttcatccagctggtg
cagacctacaaccagctgttcgaggaaaaccccatcaacgccagcggcgtggacgccaaggccatcctgtctgccagactgagcaa
gagcagacggctggaaaatctgatcgcccagctgcccggcgagaagaagaatggcctgttcggaaacctgattgccctgagcctgg
gcctgacccccaacttcaagagcaacttcgacctggccgaggatgccaaactgcagctgagcaaggacacctacgacgacgacctg
gacaacctgctggcccagatcggcgaccagtacgccgacctgtttctggccgccaagaacctgtccgacgccatcctgctgagcgac
atcctgagagtgaacaccgagatcaccaaggcccccctgagcgcctctatgatcaagagatacgacgagcaccaccaggacctgac
cctgctgaaagctctcgtgcggcagcagctgcctgagaagtacaaagagattttcttcgaccagagcaagaacggctacgccggcta
cattgacggcggagccagccaggaagagttctacaagttcatcaagcccatcctggaaaagatggacggcaccgaggaactgctcg
tgaagctgaacagagaggacctgctgcggaagcagcggaccttcgacaacggcagcatcccccaccagatccacctgggagagct
gcacgccattctgcggcggcaggaagatttttacccattcctgaaggacaaccgggaaaagatcgagaagatcctgaccttccgcatc
ccctactacgtgggccctctggccaggggaaacagcagattcgcctggatgaccagaaagagcgaggaaaccatcaccccctgga
acttcgaggaagtggtggacaagggcgcttccgcccagagcttcatcgagcggatgaccaacttcgataagaacctgcccaacgag
aaggtgctgcccaagcacagcctgctgtacgagtacttcaccgtgtataacgagctgaccaaagtgaaatacgtgaccgagggaatg
agaaagcccgccttcctgagcggcgagcagaaaaaggccatcgtggacctgctgttcaagaccaaccggaaagtgaccgtgaagc
agctgaaagaggactacttcaagaaaatcgagtgcttcgactccgtggaaatctccggcgtggaagatcggttcaacgcctccctggg
cacataccacgatctgctgaaaattatcaaggacaaggacttcctggacaatgaggaaaacgaggacattctggaagatatcgtgctg
accctgacactgtttgaggacagagagatgatcgaggaacggctgaaaacctatgcccacctgttcgacgacaaagtgatgaagcag
ctgaagcggcggagatacaccggctggggcaggctgagccggaagctgatcaacggcatccgggacaagcagtccggcaagac
aatcctggatttcctgaagtccgacggcttcgccaacagaaacttcatgcagctgatccacgacgacagcctgacctttaaagaggaca
tccagaaagcccaggtgtccggccagggcgatagcctgcacgagcacattgccaatctggccggcagccccgccattaagaaggg

catcctgcagacagtgaaggtggtggacgagctcgtgaaagtgatgggccggcacaagcccgagaacatcgtgatcgaaatggcc
agagagaaccagaccacccagaagggacagaagaacagccgcgagagaatgaagcggatcgaagagggcatcaaagagctgg
gcagccagatcctgaaagaacaccccgtggaaaacacccagctgcagaacgagaagctgtacctgtactacctgcagaatgggcg
ggatatgtacgtggaccaggaactggacatcaaccggctgtccgactacgatgtggaccatatcgtgcctcagagctttctgaaggac
gactccatcgacaacaaggtgctgaccagaagcgacaagaaccggggcaagagcgacaacgtgccctccgaagaggtcgtgaag
aagatgaagaactactggcggcagctgctgaacgccaagctgattacccagagaaagttcgacaatctgaccaaggccgagagagg
cggcctgagcgaactggataaggccggcttcatcaagagacagctggtggaaacccggcagatcacaaagcacgtggcacagatc
ctggactcccggatgaacactaagtacgacgagaatgacaagctgatccgggaagtgaaagtgatcaccctgaagtccaagctggtg
tccgatttccggaaggatttccagttttacaaagtgcgcgagatcaacaactaccaccacgcccacgacgcctacctgaacgccgtcgt
gggaaccgccctgatcaaaaagtaccctaagctggaaagcgagttcgtgtacggcgactacaaggtgtacgacgtgcggaagatga
tcgccaagagcgagcaggaaatcggcaaggctaccgccaagtacttcttctacagcaacatcatgaactttttcaagaccgagattacc
ctggccaacggcgagatccggaagcggcctctgatcgagacaaacggcgaaaccggggagatcgtgtgggataagggccgggat
tttgccaccgtgcggaaagtgctgagcatgccccaagtgaatatcgtgaaaaagaccgaggtgcagacaggcggcttcagcaaaga
gtctatcctgcccaagaggaacagcgataagctgatcgccagaaagaaggactgggaccctaagaagtacggcggcttcgacagcc
ccaccgtggcctattctgtgctggtggtggccaaagtggaaaagggcaagtccaagaaactgaagagtgtgaaagagctgctgggg
atcaccatcatggaaagaagcagcttcgagaagaatcccatcgactttctggaagccaagggctacaaagaagtgaaaaaggacctg
atcatcaagctgcctaagtactccctgttcgagctggaaaacggccggaagagaatgctggcctctgccggcgaactgcagaaggga
aacgaactggccctgccctccaaatatgtgaacttcctgtacctggccagccactatgagaagctgaagggctcccccgaggataatg
agcagaaacagctgtttgtggaacagcacaagcactacctggacgagatcatcgagcagatcagcgagttctccaagagagtgatcc
tggccgacgctaatctggacaaagtgctgtccgcctacaacaagcaccgggataagcccatcagagagcaggccgagaatatcatc
cacctgtttaccctgaccaatctgggagcccctgccgccttcaagtactttgacaccaccatcgaccggaagaggtacaccagcacca
aagaggtgctggacgccaccctgatccaccagagcatcaccggcctgtacgagacacggatcgacctgtctcagctgggaggcga
ctctggaggatctagcggaggatcctctggcagcgagacaccaggaacaagcgagtcagcaacaccagagagcagtggcggcag
cagcggcggcagcagcATGACGGCAAAACTGGAAAGCCACGTGCCTGCTGCACCACCAG
TCTCAGCGGAAGCGCCGGCACCGACGCGACCTGACGCTGCTAAGCAGGAGGCTC
GACGAGCCCACCATGAAGCCCTGAGACTGCGGTGGAAAGCAATCGAGGAAGCTG
GGGGCACTGACGCCTGGGTACGACAGCAGTTGGTTGCGAAAGGCGTAGCGGCGG
AAGAGGTGGACTTCGAAAGTCTCTCCGACAAGCAAAAAGCGGCTTGGAAAGAGA
AAAAAAAAGCTGAGGCAACCGAGCGGAGGGCTCAAAAACGACTTGCCTGGGAA
GCCTGGAAAGCGACCCATATTCATCATTTGGGCGTTGGAGTTCACTGGGACGAA
GCCGGCGGGCCTGACAAGTTCGATGTGGCAGGGAGGGAAGAAAGGGCAAAGGC
AAACGGGTTGCCCGAAGGTCTTGACAGCGTAGAAGCATTGGCGAAAGCGCTGGG
AATCTCAGTATCCCGATTGCGCTGGTTCTCTTTTCACAGGGAAGTTGATACAGGG
ACACACTACCAAACTTGGGAGATCCCAAAGAGGGATGGGGGCAAACGCACCCTC
ACGGCCCCGAAAAGGGAGCTTAAGGCAGTACAAAGGTGGGTACTCGCCAATGTG
GTGGAAAGACTTCCCGTACACGGAGCTGCTCATGGGTTTGTTGCAGGACGCTCCA
TACTCACTAATGCGCTCGCTCACCAAGGAGCTGACGTGGTTGTTAAAGTTGATAT
GAAAGATTTTTTTCCCAGTGTCACATGGCCTCGAGTTAAAGGCCTGCTCAGAAAG
GGAGGTTTGCCTGAAAACTTGGCGACCCTGCTCGCCCTTCTGAGTACCGAAGCTC
CACGGGAAGTAGTCCGATTCAGAGGGGAAACGCTGTACGTCGCTAAGGGCCCAC
GCGCTCTGCCACAGGGCGCGCCTACCAGTCCTGCGCTTACTAACGCATTGTGTCT
TAGGTTGGATAAGCGGTTGAGCGCGCTGAGTAAGCGGTTGGGTTTCACATACAC
CAGATACGCTGACGATCTTACATTTAGTTGGAGGCGCGCTAAAAAATCTCGCCAG
AAAGAGTTGCCGCTGGCTGATGCTCCCGTCGCGCTTCTCCTGGCCCGGGTTAAGG
GAGTGTTGGAAGCAGAAGGTTTCACTTTGCATCCTGATAAGACCAGGGTTCAGC
GGAAGGGAAGCCGGCAACGAGTGACCGGCCTCGTGGTTAATGAAGCGCCCGAA
GGAGTGCCAGGTGCCCGAGTTCCCCGAGATGTTGTTCGGAGATTGCGAGCGGCA
ATACACAATAGAGAGCAAGGTAAACCAGGGCCGACTGGAGAAACGTTGGAACA
ACTCAAAGGCTTGGCAGCTTTTCTGCACATGACAGACGCTGAAAAAGGAAGAGC
GTTTCTGAGAAGACTTGAGGCCCTGGAGAAGAGGCAAACTGCGaaaaggccggcggccac

gaaaaaggccggccaggcaaaaaagaaaaagcttgagggcagaggaagtctgctaacatgcggtgacgtggaggagaatcccgg
ccctgctagcatggtgagcaagggcgaggaggataacatggccatcatcaaggagttcatgcgcttcaaggtgcacatggagggctc
cgtgaacggccacgagttcgagatcgagggcgagggcgagggccgcccctacgagggcacccagaccgccaagctgaaggtga
ccaagggtggcccccctgcccttcgcctgggacatcctgtcccctcagttcatgtacggctccaaggcctacgtgaagcaccccgccg
acatccccgactacttgaagctgtccttccccgagggcttcaagtgggagcgcgtgatgaacttcgaggacggcggcgtggtgaccg
tgacccaggactcctccctgcaggacggcgagttcatctacaaggtgaagctgcgcggcaccaacttcccctccgacggccccgtaa
tgcagaagaagaccatgggctgggaggcctcctccgagcggatgtaccccgaggacggcgccctgaagggcgagatcaagcaga
ggctgaagctgaaggacggcggccactacgacgctgaggtcaagaccacctacaaggccaagaagcccgtgcagctgcccggcg
cctacaacgtcaacatcaagttggacatcacctcccacaacgaggactacaccatcgtggaacagtacgaacgcgccgagggccgc
cactccaccggcggcatggacgagctgtacaagtgagaattcctagagctcgctgatcagcctcgactgtgccttctagttgccagcc
atctgttgtttgcccctcccccgtgccttccttgaccctggaaggtgccactcccactgtcctttcctaataaaatgaggaaattgcatcgc
attgtctgagtaggtgtcattctattctggggggtggggtggggcaggacagcaagggggaggattgggaagagaatagcaggcat
gctggggagcggccgcaggaacccctagtgatggagttggccactccctctctgcgcgctcgctcgctcactgaggccgggcgacc
aaaggtcgcccgacgcccgggctttgcccgggcggcctcagtgagcgagcgagcgcgcagctgcctgcaggggcgcctgatgcg
gtattttctccttacgcatctgtgcggtatttcacaccgcatacgtcaaagcaaccatagtacgcgccctgtagcggcgcattaagcgcg
gcgggtgtggtggttacgcgcagcgtgaccgctacacttgccagcgccctagcgcccgctcctttcgctttcttcccttcctttctcgcca
cgttcgccggctttccccgtcaagctctaaatcgggggctccctttagggttccgatttagtgctttacggcacctcgaccccaaaaaact
tgatttgggtgatggttcacgtagtgggccatcgccctgatagacggtttttcgccctttgacgttggagtccacgttctttaatagtggact
cttgttccaaactggaacaacactcaaccctatctcgggctattcttttgatttataagggattttgccgatttcggCctattggttaaaaaat
gagctgatttaacaaaaatttaacgcgaattttaacaaaatattaacgtttacaattttatggtgcactctcagtacaatctgctctgatgccg
catagttaagccagccccgacacccgccaacacccgctgacgcgccctgacgggcttgtctgctcccggcatccgcttacagacaag
ctgtgaccgtctccgggagctgcatgtgtcagaggttttcaccgtcatcaccgaaacgcgcgagacgaaagggcctcgtgatacgcct
atttttataggttaatgtcatgataataatggtttcttagacgtcaggtggcacttttcggggaaatgtgcgcggaacccctatttgtttattttt
ctaaatacattcaaatatgtatccgctcatgagacaataaccctgataaatgcttcaataatattgaaaaaggaagagtatgagtattcaac
atttccgtgtcgcccttattcccttttttgcggcattttgccttcctgtttttgctcacccagaaacgctggtgaaagtaaaagatgctgaagat
cagttgggtgcacgagtgggttacatcgaactggatctcaacagcggtaagatccttgagagttttcgccccgaagaacgttttccaatg
atgagcacttttaaagttctgctatgtggcgcggtattatcccgtattgacgccgggcaagagcaactcggtcgccgcatacactattctc
agaatgacttggttgagtactcaccagtcacagaaaagcatcttacggatggcatgacagtaagagaattatgcagtgctgccataacc
atgagtgataacactgcggccaacttacttctgacaacgatcggaggaccgaaggagctaaccgcttttttgcacaacatgggggatc
atgtaactcgccttgatcgttgggaaccggagctgaatgaagccataccaaacgacgagcgtgacaccacgatgcctgtagcaatgg
caacaacgttgcgcaaactattaactggcgaactacttactctagcttcccggcaacaattaatagactggatggaggcggataaagttg
caggaccacttctgcgctcggcccttccggctggctggtttattgctgataaatctggagccggtgagcgtggaagccgcggtatcatt
gcagcactggggccagatggtaagccctcccgtatcgtagttatctacacgacggggagtcaggcaactatggatgaacgaaataga
cagatcgctgagataggtgcctcactgattaagcattggtaactgtcagaccaagtttactcatatatactttagattgatttaaaacttcattt
ttaatttaaaaggatctaggtgaagatcctttttgataatctcatgaccaaaatcccttaacgtgagttttcgttccactgagcgtcagaccc
cgtagaaaagatcaaaggatcttcttgagatcctttttttctgcgcgtaatctgctgcttgcaaacaaaaaaaccaccgctaccagcggtg
gtttgtttgccggatcaagagctaccaactctttttccgaaggtaactggcttcagcagagcgcagataccaaatactgtccttctagtgta
gccgtagttaggccaccacttcaagaactctgtagcaccgcctacatacctcgctctgctaatcctgttaccagtggctgctgccagtgg
cgataagtcgtgtcttaccgggttggactcaagacgatagttaccggataaggcgcagcggtcgggctgaacggggggttcgtgcac
acagcccagcttggagcgaacgacctacaccgaactgagatacctacagcgtgagctatgagaaagcgccacgcttcccgaaggg
agaaaggcggacaggtatccggtaagcggcagggtcggaacaggagagcgcacgagggagcttccagggggaaacgcctggta
tctttatagtcctgtcgggtttcgccacctctgacttgagcgtcgatttttgtgatgctcgtcaggggggcggagcctatggaaaaacgcc
agcaacgcggcctttttacggttcctggccttttgctggccttttgctcacatgt

**pBZ200-pU6-sgBFP-Hn_CBh-sv40NLS-Cas9-Sa163RT-NLS-T2A-mCherry**

gagggcctatttcccatgattccttcatatttgcatatacgatacaaggctgttagagagataattggaattaatttgactgtaaacacaaag
atattagtacaaaatacgtgacgtagaaagtaataatttcttgggtagtttgcagttttaaaattatgttttaaaatggactatcatatgcttacc
gtaacttgaaagtatttcgatttcttggctttatatatcttgtggaaaggacgaaacaccGCTGAAGCACTGCACGCCAT

gttttagagctagaaatagcaagttaaaataaggctagtccgttatcaacttgaaaaagtggcaccgagtcggtgcCGTACGGT
GCGAGCGACCGAGAGAGGTCCCAAGCCATCAGCCTCAGCGCCTCGAGCGCGAGA
GCGGCGTTGCGCCGCTCTGGTTGAATTGCAGGACACTCTCCGCAAGGTAGCCTGT
TCTTGGCTCTCTTCCCTCCGGTGAGTACCTCTCCGGCCGGGGAGCTGAACCAACG
AACTAGTTCATGTGGTCGGGGTAGCGGCTGAAGCACTGCACGCCGTACGTCAGG
GTGGTCACGAGGGTGGGCCAGGGCACGGGCAGCTTGCCGGTGGTGCAGATGAAC
TTCAGGGTCAGCTTGCCGTAGGTGGCCCTAGGCGCAACCGCCGTTTCCCCGGCCG
GAGAGGTACTCACCGGAGGGGAGAGCCGGTGAGGCTACCGTGCCCCAGGTGAG
AAGGTGGTGCCTTCGGGCCTCCCTCGACCGCTCGCGCtttttttctagaggtacccgttacataactta
cggtaaatggcccgcctggctgaccgcccaacgacccccgcccattgacgtcaatagtaacgccaatagggactttccattgacgtca
atgggtggagtatttacggtaaactgcccacttggcagtacatcaagtgtatcatatgccaagtacgccccctattgacgtcaatgacgg
taaatggcccgcctggcattgtgcccagtacatgaccttatgggactttcctacttggcagtacatctacgtattagtcatcgctattaccat
ggtcgaggtgagccccacgttctgcttcactctccccatctcccccccctccccacccccaattttgtatttatttattttttaattattttgtgc
agcgatgggggcgggggggggggggggcgcgcgccaggcggggcggggcggggcgaggggcggggcggggcgaggc
ggagaggtgcggcggcagccaatcagagcggcgcgctccgaaagtttccttttatggcgaggcggcggcggcggcggccctataa
aaagcgaagcgcgcggcgggcgggagtcgctgcgacgctgccttcgccccgtgccccgctccgccgccgcctcgcgccgcccg
ccccggctctgactgaccgcgttactcccacaggtgagcgggcgggacggcccttctcctccgggctgtaattagctgagcaagagg
taagggtttaagggatggttggttggtggggtattaatgtttaattacctggagcacctgcctgaaatcactttttttcaggttggaccggtg
ccaccatggactataaggaccacgacggagactacaaggatcatgatattgattacaaagacgatgacgataagatggccccaaaga
agaagcggaaggtcggtatccacggagtcccagcagccgacaagaagtacagcatcggcctggacatcggcaccaactctgtggg
ctgggccgtgatcaccgacgagtacaaggtgcccagcaagaaattcaaggtgctgggcaacaccgaccggcacagcatcaagaag
aacctgatcggagccctgctgttcgacagcggcgaaacagccgaggccacccggctgaagagaaccgccagaagaagatacacc
agacggaagaaccggatctgctatctgcaagagatcttcagcaacgagatggccaaggtggacgacagcttcttccacagactggaa
gagtccttcctggtggaagaggataagaagcacgagcggcaccccatcttcggcaacatcgtggacgaggtggcctaccacgagaa
gtaccccaccatctaccacctgagaaagaaactggtggacagcaccgacaaggccgacctgcggctgatctatctggccctggccc
acatgatcaagttccggggccacttcctgatcgagggcgacctgaaccccgacaacagcgacgtggacaagctgttcatccagctgg
tgcagacctacaaccagctgttcgaggaaaaccccatcaacgccagcggcgtggacgccaaggccatcctgtctgccagactgagc
aagagcagacggctggaaaatctgatcgcccagctgcccggcgagaagaagaatggcctgttcggaaacctgattgccctgagcct
gggcctgacccccaacttcaagagcaacttcgacctggccgaggatgccaaactgcagctgagcaaggacacctacgacgacgac
ctggacaacctgctggcccagatcggcgaccagtacgccgacctgtttctggccgccaagaacctgtccgacgccatcctgctgagc
gacatcctgagagtgaacaccgagatcaccaaggcccccctgagcgcctctatgatcaagagatacgacgagcaccaccaggacct
gaccctgctgaaagctctcgtgcggcagcagctgcctgagaagtacaaagagattttcttcgaccagagcaagaacggctacgccgg
ctacattgacggcggagccagccaggaagagttctacaagttcatcaagcccatcctggaaaagatggacggcaccgaggaactgc
tcgtgaagctgaacagagaggacctgctgcggaagcagcggaccttcgacaacggcagcatcccccaccagatccacctgggaga
gctgcacgccattctgcggcggcaggaagatttttacccattcctgaaggacaaccgggaaaagatcgagaagatcctgaccttccgc
atcccctactacgtgggccctctggccaggggaaacagcagattcgcctggatgaccagaaagagcgaggaaaccatcaccccctg
gaacttcgaggaagtggtggacaagggcgcttccgcccagagcttcatcgagcggatgaccaacttcgataagaacctgcccaacg
agaaggtgctgcccaagcacagcctgctgtacgagtacttcaccgtgtataacgagctgaccaaagtgaaatacgtgaccgagggaa
tgagaaagcccgccttcctgagcggcgagcagaaaaaggccatcgtggacctgctgttcaagaccaaccggaaagtgaccgtgaa
gcagctgaaagaggactacttcaagaaaatcgagtgcttcgactccgtggaaatctccggcgtggaagatcggttcaacgcctccctg
ggcacataccacgatctgctgaaaattatcaaggacaaggacttcctggacaatgaggaaaacgaggacattctggaagatatcgtgc
tgaccctgacactgtttgaggacagagagatgatcgaggaacggctgaaaacctatgcccacctgttcgacgacaaagtgatgaagc
agctgaagcggcggagatacaccggctggggcaggctgagccggaagctgatcaacggcatccgggacaagcagtccggcaag
acaatcctggatttcctgaagtccgacggcttcgccaacagaaacttcatgcagctgatccacgacgacagcctgacctttaaagagga
catccagaaagcccaggtgtccggccagggcgatagcctgcacgagcacattgccaatctggccggcagccccgccattaagaag
ggcatcctgcagacagtgaaggtggtggacgagctcgtgaaagtgatgggccggcacaagcccgagaacatcgtgatcgaaatgg
ccagagagaaccagaccacccagaagggacagaagaacagccgcgagagaatgaagcggatcgaagagggcatcaaagagct
gggcagccagatcctgaaagaacaccccgtggaaaacacccagctgcagaacgagaagctgtacctgtactacctgcagaatggg
cgggatatgtacgtggaccaggaactggacatcaaccggctgtccgactacgatgtggaccatatcgtgcctcagagctttctgaagg
acgactccatcgacaacaaggtgctgaccagaagcgacaagaaccggggcaagagcgacaacgtgccctccgaagaggtcgtga

agaagatgaagaactactggcggcagctgctgaacgccaagctgattacccagagaaagttcgacaatctgaccaaggccgagaga
ggcggcctgagcgaactggataaggccggcttcatcaagagacagctggtggaaacccggcagatcacaaagcacgtggcacag
atcctggactcccggatgaacactaagtacgacgagaatgacaagctgatccgggaagtgaaagtgatcaccctgaagtccaagctg
gtgtccgatttccggaaggatttccagttttacaaagtgcgcgagatcaacaactaccaccacgcccacgacgcctacctgaacgccgt
cgtgggaaccgccctgatcaaaaagtaccctaagctggaaagcgagttcgtgtacggcgactacaaggtgtacgacgtgcggaaga
tgatcgccaagagcgagcaggaaatcggcaaggctaccgccaagtacttcttctacagcaacatcatgaactttttcaagaccgagatt
accctggccaacggcgagatccggaagcggcctctgatcgagacaaacggcgaaaccggggagatcgtgtgggataagggccgg
gattttgccaccgtgcggaaagtgctgagcatgccccaagtgaatatcgtgaaaaagaccgaggtgcagacaggcggcttcagcaaa
gagtctatcctgcccaagaggaacagcgataagctgatcgccagaaagaaggactgggaccctaagaagtacggcggcttcgaca
gccccaccgtggcctattctgtgctggtggtggccaaagtggaaaagggcaagtccaagaaactgaagagtgtgaaagagctgctg
gggatcaccatcatggaaagaagcagcttcgagaagaatcccatcgactttctggaagccaagggctacaaagaagtgaaaaagga
cctgatcatcaagctgcctaagtactccctgttcgagctggaaaacggccggaagagaatgctggcctctgccggcgaactgcagaa
gggaaacgaactggccctgccctccaaatatgtgaacttcctgtacctggccagccactatgagaagctgaagggctcccccgagga
taatgagcagaaacagctgtttgtggaacagcacaagcactacctggacgagatcatcgagcagatcagcgagttctccaagagagt
gatcctggccgacgctaatctggacaaagtgctgtccgcctacaacaagcaccgggataagcccatcagagagcaggccgagaata
tcatccacctgtttaccctgaccaatctgggagcccctgccgccttcaagtactttgacaccaccatcgaccggaagaggtacaccagc
accaaagaggtgctggacgccaccctgatccaccagagcatcaccggcctgtacgagacacggatcgacctgtctcagctgggagg
cgactctggaggatctagcggaggatcctctggcagcgagacaccaggaacaagcgagtcagcaacaccagagagcagtggcgg
cagcagcggcggcagcagcATGACGGCAAAACTGGAAAGCCACGTGCCTGCTGCACCACC
AGTCTCAGCGGAAGCGCCGGCACCGACGCGACCTGACGCTGCTAAGCAGGAGGC
TCGACGAGCCCACCATGAAGCCCTGAGACTGCGGTGGAAAGCAATCGAGGAAGC
TGGGGGCACTGACGCCTGGGTACGACAGCAGTTGGTTGCGAAAGGCGTAGCGGC
GGAAGAGGTGGACTTCGAAAGTCTCTCCGACAAGCAAAAAGCGGCTTGGAAAGA
GAAAAAAAAAGCTGAGGCAACCGAGCGGAGGGCTCAAAAACGACTTGCCTGGG
AAGCCTGGAAAGCGACCCATATTCATCATTTGGGCGTTGGAGTTCACTGGGACG
AAGCCGGCGGGCCTGACAAGTTCGATGTGGCAGGGAGGGAAGAAAGGGCAAAG
GCAAACGGGTTGCCCGAAGGTCTTGACAGCGTAGAAGCATTGGCGAAAGCGCTG
GGAATCTCAGTATCCCGATTGCGCTGGTTCTCTTTTCACAGGGAAGTTGATACAG
GGACACACTACCAAACTTGGGAGATCCCAAAGAGGGATGGGGGCAAACGCACC
CTCACGGCCCCGAAAAGGGAGCTTAAGGCAGTACAAAGGTGGGTACTCGCCAAT
GTGGTGGAAAGACTTCCCGTACACGGAGCTGCTCATGGGTTTGTTGCAGGACGCT
CCATACTCACTAATGCGCTCGCTCACCAAGGAGCTGACGTGGTTGTTAAAGTTGA
TATGAAAGATTTTTTTCCCAGTGTCACATGGCCTCGAGTTAAAGGCCTGCTCAGA
AAGGGAGGTTTGCCTGAAAACTTGGCGACCCTGCTCGCCCTTCTGAGTACCGAAG
CTCCACGGGAAGTAGTCCGATTCAGAGGGGAAACGCTGTACGTCGCTAAGGGCC
CACGCGCTCTGCCACAGGGCGCGCCTACCAGTCCTGCGCTTACTAACGCATTGTG
TCTTAGGTTGGATAAGCGGTTGAGCGCGCTGAGTAAGCGGTTGGGTTTCACATAC
ACCAGATACGCTGACGATCTTACATTTAGTTGGAGGCGCGCTAAAAAATCTCGCC
AGAAAGAGTTGCCGCTGGCTGATGCTCCCGTCGCGCTTCTCCTGGCCCGGGTTAA
GGGAGTGTTGGAAGCAGAAGGTTTCACTTTGCATCCTGATAAGACCAGGGTTCA
GCGGAAGGGAAGCCGGCAACGAGTGACCGGCCTCGTGGTTAATGAAGCGCCCGA
AGGAGTGCCAGGTGCCCGAGTTCCCCGAGATGTTGTTCGGAGATTGCGAGCGGC
AATACACAATAGAGAGCAAGGTAAACCAGGGCCGACTGGAGAAACGTTGGAAC
AACTCAAAGGCTTGGCAGCTTTTCTGCACATGACAGACGCTGAAAAAGGAAGAG
CGTTTCTGAGAAGACTTGAGGCCCTGGAGAAGAGGCAAACTGCGaaaaggccggcggcc
acgaaaaaggccggccaggcaaaaaagaaaaagcttgagggcagaggaagtctgctaacatgcggtgacgtggaggagaatccc
ggccctgctagcatggtgagcaagggcgaggaggataacatggccatcatcaaggagttcatgcgcttcaaggtgcacatggaggg
ctccgtgaacggccacgagttcgagatcgagggcgagggcgagggccgcccctacgagggcacccagaccgccaagctgaagg
tgaccaagggtggcccctgcccttcgcctgggacatcctgtcccctcagttcatgtacggctccaaggcctacgtgaagcaccccgc
cgacatccccgactacttgaagctgtccttccccgagggcttcaagtgggagcgcgtgatgaacttcgaggacggcggcgtggtgac

cgtgacccaggactcctccctgcaggacggcgagttcatctacaaggtgaagctgcgcggcaccaacttcccctccgacggccccgt
aatgcagaagaagaccatgggctgggaggcctcctccgagcggatgtaccccgaggacggcgccctgaagggcgagatcaagca
gaggctgaagctgaaggacggcggccactacgacgctgaggtcaagaccacctacaaggccaagaagcccgtgcagctgcccgg
cgcctacaacgtcaacatcaagttggacatcacctcccacaacgaggactacaccatcgtggaacagtacgaacgcgccgagggcc
gccactccaccggcggcatggacgagctgtacaagtgagaattcctagagctcgctgatcagcctcgactgtgccttctagttgccag
ccatctgttgtttgcccctcccccgtgccttccttgaccctggaaggtgccactcccactgtcctttcctaataaaatgaggaaattgcatc
gcattgtctgagtaggtgtcattctattctggggggtggggtggggcaggacagcaagggggaggattgggaagagaatagcaggc
atgctggggagcggccgcaggaacccctagtgatggagttggccactccctctctgcgcgctcgctcgctcactgaggccgggcga
ccaaaggtcgcccgacgcccgggctttgcccgggcggcctcagtgagcgagcgagcgcgcagctgcctgcaggggcgcctgatg
cggtattttctccttacgcatctgtgcggtatttcacaccgcatacgtcaaagcaaccatagtacgcgccctgtagcggcgcattaagcg
cggcgggtgtggtggttacgcgcagcgtgaccgctacacttgccagcgccctagcgcccgctcctttcgctttcttcccttcctttctcgc
cacgttcgccggctttccccgtcaagctctaaatcgggggctccctttagggttccgatttagtgctttacggcacctcgaccccaaaaa
acttgatttgggtgatggttcacgtagtgggccatcgccctgatagacggtttttcgccctttgacgttggagtccacgttctttaatagtgg
actcttgttccaaactggaacaacactcaaccctatctcgggctattcttttgatttataagggattttgccgatttcggCctattggttaaaa
aatgagctgatttaacaaaaatttaacgcgaattttaacaaaatattaacgtttacaattttatggtgcactctcagtacaatctgctctgatgc
cgcatagttaagccagccccgacacccgccaacacccgctgacgcgccctgacgggcttgtctgctcccggcatccgcttacagaca
agctgtgaccgtctccgggagctgcatgtgtcagaggttttcaccgtcatcaccgaaacgcgcgagacgaaagggcctcgtgatacg
cctatttttataggttaatgtcatgataataatggtttcttagacgtcaggtggcacttttcggggaaatgtgcgcggaacccctatttgtttat
ttttctaaatacattcaaatatgtatccgctcatgagacaataaccctgataaatgcttcaataatattgaaaaaggaagagtatgagtattc
aacatttccgtgtcgcccttattcccttttttgcggcattttgccttcctgtttttgctcacccagaaacgctggtgaaagtaaaagatgctga
agatcagttgggtgcacgagtgggttacatcgaactggatctcaacagcggtaagatccttgagagttttcgccccgaagaacgttttcc
aatgatgagcacttttaaagttctgctatgtggcgcggtattatcccgtattgacgccgggcaagagcaactcggtcgccgcatacacta
ttctcagaatgacttggttgagtactcaccagtcacagaaaagcatcttacggatggcatgacagtaagagaattatgcagtgctgccat
aaccatgagtgataacactgcggccaacttacttctgacaacgatcggaggaccgaaggagctaaccgcttttttgcacaacatgggg
gatcatgtaactcgccttgatcgttgggaaccggagctgaatgaagccataccaaacgacgagcgtgacaccacgatgcctgtagca
atggcaacaacgttgcgcaaactattaactggcgaactacttactctagcttcccggcaacaattaatagactggatggaggcggataa
agttgcaggaccacttctgcgctcggcccttccggctggctggtttattgctgataaatctggagccggtgagcgtggaagccgcggta
tcattgcagcactggggccagatggtaagccctcccgtatcgtagttatctacacgacggggagtcaggcaactatggatgaacgaaa
tagacagatcgctgagataggtgcctcactgattaagcattggtaactgtcagaccaagtttactcatatatactttagattgatttaaaactt
catttttaatttaaaaggatctaggtgaagatcctttttgataatctcatgaccaaaatcccttaacgtgagttttcgttccactgagcgtcaga
ccccgtagaaaagatcaaaggatcttcttgagatcctttttttctgcgcgtaatctgctgcttgcaaacaaaaaaaccaccgctaccagcg
gtggtttgtttgccggatcaagagctaccaactctttttccgaaggtaactggcttcagcagagcgcagataccaaatactgtccttctagt
gtagccgtagttaggccaccacttcaagaactctgtagcaccgcctacatacctcgctctgctaatcctgttaccagtggctgctgccag
tggcgataagtcgtgtcttaccgggttggactcaagacgatagttaccggataaggcgcagcggtcgggctgaacggggggttcgtg
cacacagcccagcttggagcgaacgacctacaccgaactgagatacctacagcgtgagctatgagaaagcgccacgcttcccgaag
ggagaaaggcggacaggtatccggtaagcggcagggtcggaacaggagagcgcacgagggagcttccagggggaaacgcctg
gtatctttatagtcctgtcgggtttcgccacctctgacttgagcgtcgatttttgtgatgctcgtcaggggggcggagcctatggaaaaac
gccagcaacgcggcctttttacggttcctggccttttgctggccttttgctcacatgt

**pBZ201-pU6-sgBFP-At_CBh-sv40NLS-Cas9-Sa163RT-NLS-T2A-mCherry**

gagggcctatttcccatgattccttcatatttgcatatacgatacaaggctgttagagagataattggaattaatttgactgtaaacacaaag
atattagtacaaaatacgtgacgtagaaagtaataatttcttgggtagtttgcagttttaaaattatgttttaaaatggactatcatatgcttacc
gtaacttgaaagtatttcgatttcttggctttatatatcttgtggaaaggacgaaacaccGCTGAAGCACTGCACGCCAT
gttttagagctagaaatagcaagttaaaataaggctagtccgttatcaacttgaaaaagtggcaccgagtcggtgcCGTACGGT
GCGAGCGACCGAGAGAGGTCCCAAGCCATCAGCCTCAGCGCCTCGAGCGCGAGA
GCGGCGTTGCGCCGCTCTGGTTGAATTGCAGGACACTCTCCGCAAGGTAGCCTGT
TCTTGGCTCTCTTCCCTCCGGTGAGTACCTCTCCGGCCGGGGAGCTGAACCAACG
AACTAGTCCTGAAGTTCATCTGCACCACCGGCAAGCTGCCCGTGCCCTGGCCCAC

CCTCGTGACCACCCTGACGTACGGCGTGCAGTGCTTCAGCCGCTACCCCGACCAC
ATGAAGCAGCACGACTTCCTAGGCGCAACCGCCGTTTCCCCGGCCGGAGAGGTA
CTCACCGGAGGGGAGAGCCGGTGAGGCTACCGTGCCCCAGGTGAGAAGGTGGTG
CCTTCGGGCCTCCCTCGACCGCTCGCGCtttttttctagaggtacccgttacataacttacggtaaatggcccgc
ctggctgaccgcccaacgacccccgcccattgacgtcaatagtaacgccaatagggactttccattgacgtcaatgggtggagtattta
cggtaaactgcccacttggcagtacatcaagtgtatcatatgccaagtacgccccctattgacgtcaatgacggtaaatggcccgcctg
gcattgtgcccagtacatgaccttatgggactttcctacttggcagtacatctacgtattagtcatcgctattaccatggtcgaggtgagcc
ccacgttctgcttcactctccccatctcccccccctccccacccccaattttgtatttatttattttttaattattttgtgcagcgatgggggcgg
gggggggggggggcgcgcgccaggcggggcggggcggggcgaggggcggggcggggcgaggcggagaggtgcggcg
gcagccaatcagagcggcgcgctccgaaagtttccttttatggcgaggcggcggcggcggcggccctataaaaagcgaagcgcgc
ggcgggcgggagtcgctgcgacgctgccttcgccccgtgccccgctccgccgccgcctcgcgccgcccgccccggctctgactga
ccgcgttactcccacaggtgagcgggcgggacggcccttctcctccgggctgtaattagctgagcaagaggtaagggtttaagggat
ggttggttggtggggtattaatgtttaattacctggagcacctgcctgaaatcactttttttcaggttggaccggtgccaccatggactataa
ggaccacgacggagactacaaggatcatgatattgattacaaagacgatgacgataagatggccccaaagaagaagcggaaggtcg
gtatccacggagtcccagcagccgacaagaagtacagcatcggcctggacatcggcaccaactctgtgggctgggccgtgatcacc
gacgagtacaaggtgcccagcaagaaattcaaggtgctgggcaacaccgaccggcacagcatcaagaagaacctgatcggagccc
tgctgttcgacagcggcgaaacagccgaggccacccggctgaagagaaccgccagaagaagatacaccagacggaagaaccgg
atctgctatctgcaagagatcttcagcaacgagatggccaaggtggacgacagcttcttccacagactggaagagtccttcctggtgga
agaggataagaagcacgagcggcaccccatcttcggcaacatcgtggacgaggtggcctaccacgagaagtaccccaccatctacc
acctgagaaagaaactggtggacagcaccgacaaggccgacctgcggctgatctatctggccctggcccacatgatcaagttccgg
ggccacttcctgatcgagggcgacctgaaccccgacaacagcgacgtggacaagctgttcatccagctggtgcagacctacaacca
gctgttcgaggaaaaccccatcaacgccagcggcgtggacgccaaggccatcctgtctgccagactgagcaagagcagacggctg
gaaaatctgatcgcccagctgcccggcgagaagaagaatggcctgttcggaaacctgattgccctgagcctgggcctgacccccaa
cttcaagagcaacttcgacctggccgaggatgccaaactgcagctgagcaaggacacctacgacgacgacctggacaacctgctgg
cccagatcggcgaccagtacgccgacctgtttctggccgccaagaacctgtccgacgccatcctgctgagcgacatcctgagagtga
acaccgagatcaccaaggcccccctgagcgcctctatgatcaagagatacgacgagcaccaccaggacctgaccctgctgaaagct
ctcgtgcggcagcagctgcctgagaagtacaaagagattttcttcgaccagagcaagaacggctacgccggctacattgacggcgga
gccagccaggaagagttctacaagttcatcaagcccatcctggaaaagatggacggcaccgaggaactgctcgtgaagctgaacag
agaggacctgctgcggaagcagcggaccttcgacaacggcagcatcccccaccagatccacctgggagagctgcacgccattctg
cggcggcaggaagatttttacccattcctgaaggacaaccgggaaaagatcgagaagatcctgaccttccgcatcccctactacgtgg
gccctctggccaggggaaacagcagattcgcctggatgaccagaaagagcgaggaaaccatcaccccctggaacttcgaggaagt
ggtggacaagggcgcttccgcccagagcttcatcgagcggatgaccaacttcgataagaacctgcccaacgagaaggtgctgccca
agcacagcctgctgtacgagtacttcaccgtgtataacgagctgaccaaagtgaaatacgtgaccgagggaatgagaaagcccgcct
tcctgagcggcgagcagaaaaaggccatcgtggacctgctgttcaagaccaaccggaaagtgaccgtgaagcagctgaaagagga
ctacttcaagaaaatcgagtgcttcgactccgtggaaatctccggcgtggaagatcggttcaacgcctccctgggcacataccacgatc
tgctgaaaattatcaaggacaaggacttcctggacaatgaggaaaacgaggacattctggaagatatcgtgctgaccctgacactgttt
gaggacagagagatgatcgaggaacggctgaaaacctatgcccacctgttcgacgacaaagtgatgaagcagctgaagcggcgga
gatacaccggctggggcaggctgagccggaagctgatcaacggcatccgggacaagcagtccggcaagacaatcctggatttcctg
aagtccgacggcttcgccaacagaaacttcatgcagctgatccacgacgacagcctgacctttaaagaggacatccagaaagcccag
gtgtccggccagggcgatagcctgcacgagcacattgccaatctggccggcagccccgccattaagaagggcatcctgcagacagt
gaaggtggtggacgagctcgtgaaagtgatgggccggcacaagcccgagaacatcgtgatcgaaatggccagagagaaccagac
cacccagaagggacagaagaacagccgcgagagaatgaagcggatcgaagagggcatcaaagagctgggcagccagatcctga
aagaacaccccgtggaaaacacccagctgcagaacgagaagctgtacctgtactacctgcagaatgggcgggatatgtacgtggac
caggaactggacatcaaccggctgtccgactacgatgtggaccatatcgtgcctcagagctttctgaaggacgactccatcgacaaca
aggtgctgaccagaagcgacaagaaccggggcaagagcgacaacgtgccctccgaagaggtcgtgaagaagatgaagaactact
ggcggcagctgctgaacgccaagctgattacccagagaaagttcgacaatctgaccaaggccgagagaggcggcctgagcgaact
ggataaggccggcttcatcaagagacagctggtggaaacccggcagatcacaaagcacgtggcacagatcctggactcccggatg
aacactaagtacgacgagaatgacaagctgatccgggaagtgaaagtgatcaccctgaagtccaagctggtgtccgatttccggaag
gatttccagttttacaaagtgcgcgagatcaacaactaccaccacgcccacgacgcctacctgaacgccgtcgtgggaaccgccctg
atcaaaaagtaccctaagctggaaagcgagttcgtgtacggcgactacaaggtgtacgacgtgcggaagatgatcgccaagagcga

gcaggaaatcggcaaggctaccgccaagtacttcttctacagcaacatcatgaactttttcaagaccgagattaccctggccaacggcg
agatccggaagcggcctctgatcgagacaaacggcgaaaccggggagatcgtgtgggataagggccgggattttgccaccgtgcg
gaaagtgctgagcatgccccaagtgaatatcgtgaaaaagaccgaggtgcagacaggcggcttcagcaaagagtctatcctgccca
agaggaacagcgataagctgatcgccagaaagaaggactgggaccctaagaagtacggcggcttcgacagccccaccgtggccta
ttctgtgctggtggtggccaaagtggaaaagggcaagtccaagaaactgaagagtgtgaaagagctgctggggatcaccatcatgga
aagaagcagcttcgagaagaatcccatcgactttctggaagccaagggctacaaagaagtgaaaaaggacctgatcatcaagctgcc
taagtactccctgttcgagctggaaaacggccggaagagaatgctggcctctgccggcgaactgcagaagggaaacgaactggccc
tgccctccaaatatgtgaacttcctgtacctggccagccactatgagaagctgaagggctcccccgaggataatgagcagaaacagct
gtttgtggaacagcacaagcactacctggacgagatcatcgagcagatcagcgagttctccaagagagtgatcctggccgacgctaat
ctggacaaagtgctgtccgcctacaacaagcaccgggataagcccatcagagagcaggccgagaatatcatccacctgtttaccctg
accaatctgggagcccctgccgccttcaagtactttgacaccaccatcgaccggaagaggtacaccagcaccaaagaggtgctgga
cgccaccctgatccaccagagcatcaccggcctgtacgagacacggatcgacctgtctcagctgggaggcgactctggaggatcta
gcggaggatcctctggcagcgagacaccaggaacaagcgagtcagcaacaccagagagcagtggcggcagcagcggcggcag
cagcATGACGGCAAAACTGGAAAGCCACGTGCCTGCTGCACCACCAGTCTCAGCG
GAAGCGCCGGCACCGACGCGACCTGACGCTGCTAAGCAGGAGGCTCGACGAGCC
CACCATGAAGCCCTGAGACTGCGGTGGAAAGCAATCGAGGAAGCTGGGGGCACT
GACGCCTGGGTACGACAGCAGTTGGTTGCGAAAGGCGTAGCGGCGGAAGAGGTG
GACTTCGAAAGTCTCTCCGACAAGCAAAAAGCGGCTTGGAAAGAGAAAAAAA
AGCTGAGGCAACCGAGCGGAGGGCTCAAAAACGACTTGCCTGGGAAGCCTGGA
AAGCGACCCATATTCATCATTTGGGCGTTGGAGTTCACTGGGACGAAGCCGGCG
GGCCTGACAAGTTCGATGTGGCAGGGAGGGAAGAAAGGGCAAAGGCAAACGGG
TTGCCCGAAGGTCTTGACAGCGTAGAAGCATTGGCGAAAGCGCTGGGAATCTCA
GTATCCCGATTGCGCTGGTTCTCTTTTCACAGGGAAGTTGATACAGGGACACACT
ACCAAACTTGGGAGATCCCAAAGAGGGATGGGGGCAAACGCACCCTCACGGCCC
CGAAAAGGGAGCTTAAGGCAGTACAAAGGTGGGTACTCGCCAATGTGGTGGAAA
GACTTCCCGTACACGGAGCTGCTCATGGGTTTGTTGCAGGACGCTCCATACTCAC
TAATGCGCTCGCTCACCAAGGAGCTGACGTGGTTGTTAAAGTTGATATGAAAGAT
TTTTTTCCCAGTGTCACATGGCCTCGAGTTAAAGGCCTGCTCAGAAAGGGAGGTT
TGCCTGAAAACTTGGCGACCCTGCTCGCCCTTCTGAGTACCGAAGCTCCACGGGA
AGTAGTCCGATTCAGAGGGGAAACGCTGTACGTCGCTAAGGGCCCACGCGCTCT
GCCACAGGGCGCGCCTACCAGTCCTGCGCTTACTAACGCATTGTGTCTTAGGTTG
GATAAGCGGTTGAGCGCGCTGAGTAAGCGGTTGGGTTTCACATACACCAGATAC
GCTGACGATCTTACATTTAGTTGGAGGCGCGCTAAAAAATCTCGCCAGAAAGAG
TTGCCGCTGGCTGATGCTCCCGTCGCGCTTCTCCTGGCCCGGGTTAAGGGAGTGT
TGGAAGCAGAAGGTTTCACTTTGCATCCTGATAAGACCAGGGTTCAGCGGAAGG
GAAGCCGGCAACGAGTGACCGGCCTCGTGGTTAATGAAGCGCCCGAAGGAGTGC
CAGGTGCCCGAGTTCCCCGAGATGTTGTTCGGAGATTGCGAGCGGCAATACACA
ATAGAGAGCAAGGTAAACCAGGGCCGACTGGAGAAACGTTGGAACAACTCAAA
GGCTTGGCAGCTTTTCTGCACATGACAGACGCTGAAAAAGGAAGAGCGTTTCTG
AGAAGACTTGAGGCCCTGGAGAAGAGGCAAACTGCGaaaaggccggcggccacgaaaaaggc
cggccaggcaaaaaagaaaaagcttgagggcagaggaagtctgctaacatgcggtgacgtggaggagaatcccggccctgctagc
atggtgagcaagggcgaggaggataacatggccatcatcaaggagttcatgcgcttcaaggtgcacatggagggctccgtgaacgg
ccacgagttcgagatcgagggcgagggcgagggccgcccctacgagggcacccagaccgccaagctgaaggtgaccaagggtg
gcccccctgcccttcgcctgggacatcctgtcccctcagttcatgtacggctccaaggcctacgtgaagcaccccgccgacatccccga
ctacttgaagctgtccttccccgagggcttcaagtgggagcgcgtgatgaacttcgaggacggcggcgtggtgaccgtgacccagga
ctcctccctgcaggacggcgagttcatctacaaggtgaagctgcgcggcaccaacttcccctccgacggccccgtaatgcagaagaa
gaccatgggctgggaggcctcctccgagcggatgtaccccgaggacggcgccctgaagggcgagatcaagcagaggctgaagct
gaaggacggcggccactacgacgctgaggtcaagaccacctacaaggccaagaagcccgtgcagctgcccggcgcctacaacgt
caacatcaagttggacatcacctcccacaacgaggactacaccatcgtggaacagtacgaacgcgccgagggccgccactccaccg
gcggcatggacgagctgtacaagtgagaattcctagagctcgctgatcagcctcgactgtgccttctagttgccagccatctgttgtttg

cccctcccccgtgccttccttgaccctggaaggtgccactcccactgtcctttcctaataaaatgaggaaattgcatcgcattgtctgagt
aggtgtcattctattctggggggtggggtggggcaggacagcaagggggaggattgggaagagaatagcaggcatgctggggagc
ggccgcaggaacccctagtgatggagttggccactccctctctgcgcgctcgctcgctcactgaggccgggcgaccaaaggtcgcc
cgacgcccgggctttgcccgggcggcctcagtgagcgagcgagcgcgcagctgcctgcaggggcgcctgatgcggtattttctcctt
acgcatctgtgcggtatttcacaccgcatacgtcaaagcaaccatagtacgcgccctgtagcggcgcattaagcgcggcgggtgtggt
ggttacgcgcagcgtgaccgctacacttgccagcgccctagcgcccgctcctttcgctttcttcccttcctttctcgccacgttcgccggc
tttccccgtcaagctctaaatcgggggctccctttagggttccgatttagtgctttacggcacctcgaccccaaaaaacttgatttgggtga
tggttcacgtagtgggccatcgccctgatagacggtttttcgccctttgacgttggagtccacgttctttaatagtggactcttgttccaaac
tggaacaacactcaaccctatctcgggctattcttttgatttataagggattttgccgatttcggCctattggttaaaaaatgagctgatttaa
caaaaatttaacgcgaattttaacaaaatattaacgtttacaattttatggtgcactctcagtacaatctgctctgatgccgcatagttaagcc
agccccgacacccgccaacacccgctgacgcgccctgacgggcttgtctgctcccggcatccgcttacagacaagctgtgaccgtct
ccgggagctgcatgtgtcagaggttttcaccgtcatcaccgaaacgcgcgagacgaaagggcctcgtgatacgcctatttttataggtt
aatgtcatgataataatggtttcttagacgtcaggtggcacttttcggggaaatgtgcgcggaacccctatttgtttatttttctaaatacattc
aaatatgtatccgctcatgagacaataaccctgataaatgcttcaataatattgaaaaaggaagagtatgagtattcaacatttccgtgtcg
cccttattcccttttttgcggcattttgccttcctgtttttgctcacccagaaacgctggtgaaagtaaaagatgctgaagatcagttgggtgc
acgagtgggttacatcgaactggatctcaacagcggtaagatccttgagagttttcgccccgaagaacgttttccaatgatgagcactttt
aaagttctgctatgtggcgcggtattatcccgtattgacgccgggcaagagcaactcggtcgccgcatacactattctcagaatgacttg
gttgagtactcaccagtcacagaaaagcatcttacggatggcatgacagtaagagaattatgcagtgctgccataaccatgagtgataa
cactgcggccaacttacttctgacaacgatcggaggaccgaaggagctaaccgcttttttgcacaacatgggggatcatgtaactcgcc
ttgatcgttgggaaccggagctgaatgaagccataccaaacgacgagcgtgacaccacgatgcctgtagcaatggcaacaacgttgc
gcaaactattaactggcgaactacttactctagcttcccggcaacaattaatagactggatggaggcggataaagttgcaggaccacttc
tgcgctcggcccttccggctggctggtttattgctgataaatctggagccggtgagcgtggaagccgcggtatcattgcagcactggg
gccagatggtaagccctcccgtatcgtagttatctacacgacggggagtcaggcaactatggatgaacgaaatagacagatcgctga
gataggtgcctcactgattaagcattggtaactgtcagaccaagtttactcatatatactttagattgatttaaaacttcatttttaatttaaaag
gatctaggtgaagatcctttttgataatctcatgaccaaaatcccttaacgtgagttttcgttccactgagcgtcagaccccgtagaaaaga
tcaaaggatcttcttgagatcctttttttctgcgcgtaatctgctgcttgcaaacaaaaaaaccaccgctaccagcggtggtttgtttgccgg
atcaagagctaccaactctttttccgaaggtaactggcttcagcagagcgcagataccaaatactgtccttctagtgtagccgtagttagg
ccaccacttcaagaactctgtagcaccgcctacatacctcgctctgctaatcctgttaccagtggctgctgccagtggcgataagtcgtg
tcttaccgggttggactcaagacgatagttaccggataaggcgcagcggtcgggctgaacggggggttcgtgcacacagcccagctt
ggagcgaacgacctacaccgaactgagatacctacagcgtgagctatgagaaagcgccacgcttcccgaagggagaaaggcgga
caggtatccggtaagcggcagggtcggaacaggagagcgcacgagggagcttccagggggaaacgcctggtatctttatagtcctg
tcgggtttcgccacctctgacttgagcgtcgatttttgtgatgctcgtcaggggggcggagcctatggaaaaacgccagcaacgcggc
ctttttacggttcctggccttttgctggccttttgctcacatgt

**pBZ202-pU6-sgBFP-An_CBh-sv40NLS-Cas9-Sa163RT-NLS-T2A-mCherry**

gagggcctatttcccatgattccttcatatttgcatatacgatacaaggctgttagagagataattggaattaatttgactgtaaacacaaag
atattagtacaaaatacgtgacgtagaaagtaataatttcttgggtagtttgcagttttaaaattatgttttaaaatggactatcatatgcttacc
gtaacttgaaagtatttcgatttcttggctttatatatcttgtggaaaggacgaaacaccGCTGAAGCACTGCACGCCAT
gttttagagctagaaatagcaagttaaaataaggctagtccgttatcaacttgaaaaagtggcaccgagtcggtgcCGTACGGT
GCGAGCGACCGAGAGAGGTCCCAAGCCATCAGCCTCAGCGCCTCGAGCGCGAGA
GCGGCGTTGCGCCGCTCTGGTTGAATTGCAGGACACTCTCCGCAAGGTAGCCTGT
TCTTGGCTCTCTTCCCTCCGGTGAGTACCTCTCCGGCCGGGGAGCTGAACCAACG
AACTAGTAAGTCGTGCTGCTTCATGTGGTCGGGGTAGCGGCTGAAGCACTGCAC
GCCGTACGTCAGGGTGGTCACGAGGGTGGGCCAGGGCACGGGCAGCTTGCCGGT
GGTGCAGATGAACTTCAGGCCTAGGCGCAACCGCCGTTTCCCCGGCCGGAGAGG
TACTCACCGGAGGGGAGAGCCGGTGAGGCTACCGTGCCCCAGGTGAGAAGGTGG
TGCCTTCGGGCCTCCCTCGACCGCTCGCGCtttttttctagaggtacccgttacataacttacggtaaatggcc
cgcctggctgaccgcccaacgacccccgcccattgacgtcaatagtaacgccaatagggactttccattgacgtcaatgggtggagta

tttacggtaaactgcccacttggcagtacatcaagtgtatcatatgccaagtacgccccctattgacgtcaatgacggtaaatggcccgc
ctggcattgtgcccagtacatgaccttatgggactttcctacttggcagtacatctacgtattagtcatcgctattaccatggtcgaggtga
gccccacgttctgcttcactctccccatctcccccccctccccacccccaattttgtatttatttattttttaattattttgtgcagcgatgggg
cgggggggggggggggggcgcgcgccaggcggggcggggcggggcgaggggcggggcggggcgaggcggagaggtgcg
gcggcagccaatcagagcggcgcgctccgaaagtttccttttatggcgaggcggcggcggcggcggccctataaaaagcgaagcg
cgcggcgggcgggagtcgctgcgacgctgccttcgccccgtgccccgctccgccgccgcctcgcgccgcccgccccggctctga
ctgaccgcgttactcccacaggtgagcgggcgggacggcccttctcctccgggctgtaattagctgagcaagaggtaagggtttaag
ggatggttggttggtggggtattaatgtttaattacctggagcacctgcctgaaatcactttttttcaggttggaccggtgccaccatggac
tataaggaccacgacggagactacaaggatcatgatattgattacaaagacgatgacgataagatggccccaaagaagaagcggaa
ggtcggtatccacggagtcccagcagccgacaagaagtacagcatcggcctggacatcggcaccaactctgtgggctgggccgtga
tcaccgacgagtacaaggtgcccagcaagaaattcaaggtgctgggcaacaccgaccggcacagcatcaagaagaacctgatcgg
agccctgctgttcgacagcggcgaaacagccgaggccacccggctgaagagaaccgccagaagaagatacaccagacggaaga
accggatctgctatctgcaagagatcttcagcaacgagatggccaaggtggacgacagcttcttccacagactggaagagtccttcct
ggtggaagaggataagaagcacgagcggcaccccatcttcggcaacatcgtggacgaggtggcctaccacgagaagtaccccacc
atctaccacctgagaaagaaactggtggacagcaccgacaaggccgacctgcggctgatctatctggccctggcccacatgatcaag
ttccggggccacttcctgatcgagggcgacctgaaccccgacaacagcgacgtggacaagctgttcatccagctggtgcagacctac
aaccagctgttcgaggaaaaccccatcaacgccagcggcgtggacgccaaggccatcctgtctgccagactgagcaagagcagac
ggctggaaaatctgatcgcccagctgcccggcgagaagaagaatggcctgttcggaaacctgattgccctgagcctgggcctgacc
cccaacttcaagagcaacttcgacctggccgaggatgccaaactgcagctgagcaaggacacctacgacgacgacctggacaacct
gctggcccagatcggcgaccagtacgccgacctgtttctggccgccaagaacctgtccgacgccatcctgctgagcgacatcctgag
agtgaacaccgagatcaccaaggcccccctgagcgcctctatgatcaagagatacgacgagcaccaccaggacctgaccctgctga
aagctctcgtgcggcagcagctgcctgagaagtacaaagagattttcttcgaccagagcaagaacggctacgccggctacattgacg
gcggagccagccaggaagagttctacaagttcatcaagcccatcctggaaaagatggacggcaccgaggaactgctcgtgaagctg
aacagagaggacctgctgcggaagcagcggaccttcgacaacggcagcatcccccaccagatccacctgggagagctgcacgcc
attctgcggcggcaggaagatttttacccattcctgaaggacaaccgggaaaagatcgagaagatcctgaccttccgcatcccctacta
cgtgggccctctggccaggggaaacagcagattcgcctggatgaccagaaagagcgaggaaaccatcaccccctggaacttcgag
gaagtggtggacaagggcgcttccgcccagagcttcatcgagcggatgaccaacttcgataagaacctgcccaacgagaaggtgct
gcccaagcacagcctgctgtacgagtacttcaccgtgtataacgagctgaccaaagtgaaatacgtgaccgagggaatgagaaagcc
cgccttcctgagcggcgagcagaaaaaggccatcgtggacctgctgttcaagaccaaccggaaagtgaccgtgaagcagctgaaa
gaggactacttcaagaaaatcgagtgcttcgactccgtggaaatctccggcgtggaagatcggttcaacgcctccctgggcacatacc
acgatctgctgaaaattatcaaggacaaggacttcctggacaatgaggaaaacgaggacattctggaagatatcgtgctgaccctgac
actgtttgaggacagagagatgatcgaggaacggctgaaaacctatgcccacctgttcgacgacaaagtgatgaagcagctgaagcg
gcggagatacaccggctggggcaggctgagccggaagctgatcaacggcatccgggacaagcagtccggcaagacaatcctgga
tttcctgaagtccgacggcttcgccaacagaaacttcatgcagctgatccacgacgacagcctgacctttaaagaggacatccagaaa
gcccaggtgtccggccagggcgatagcctgcacgagcacattgccaatctggccggcagccccgccattaagaagggcatcctgc
agacagtgaaggtggtggacgagctcgtgaaagtgatgggccggcacaagcccgagaacatcgtgatcgaaatggccagagaga
accagaccacccagaagggacagaagaacagccgcgagagaatgaagcggatcgaagagggcatcaaagagctgggcagcca
gatcctgaaagaacaccccgtggaaaacacccagctgcagaacgagaagctgtacctgtactacctgcagaatggcgggatatgta
cgtggaccaggaactggacatcaaccggctgtccgactacgatgtggaccatatcgtgcctcagagctttctgaaggacgactccatc
gacaacaaggtgctgaccagaagcgacaagaaccggggcaagagcgacaacgtgccctccgaagaggtcgtgaagaagatgaa
gaactactggcggcagctgctgaacgccaagctgattacccagagaaagttcgacaatctgaccaaggccgagagaggcggcctg
agcgaactggataaggccggcttcatcaagagacagctggtggaaacccggcagatcacaaagcacgtggcacagatcctggactc
ccggatgaacactaagtacgacgagaatgacaagctgatccgggaagtgaaagtgatcaccctgaagtccaagctggtgtccgatttc
cggaaggatttccagttttacaaagtgcgcgagatcaacaactaccaccacgcccacgacgcctacctgaacgccgtcgtgggaacc
gccctgatcaaaaagtaccctaagctggaaagcgagttcgtgtacggcgactacaaggtgtacgacgtgcggaagatgatcgccaag
agcgagcaggaaatcggcaaggctaccgccaagtacttcttctacagcaacatcatgaactttttcaagaccgagattaccctggccaa
cggcgagatccggaagcggcctctgatcgagacaaacggcgaaaccggggagatcgtgtgggataagggccgggattttgccacc
gtgcggaaagtgctgagcatgccccaagtgaatatcgtgaaaaagaccgaggtgcagacaggcggcttcagcaaagagtctatcct
gcccaagaggaacagcgataagctgatcgccagaaagaaggactgggaccctaagaagtacggcggcttcgacagccccaccgt
ggcctattctgtgctggtggtggccaaagtggaaaagggcaagtccaagaaactgaagagtgtgaaagagctgctggggatcaccat

catggaaagaagcagcttcgagaagaatcccatcgactttctggaagccaagggctacaaagaagtgaaaaaggacctgatcatcaa
gctgcctaagtactccctgttcgagctggaaaacggccggaagagaatgctggcctctgccggcgaactgcagaagggaaacgaac
tggccctgccctccaaatatgtgaacttcctgtacctggccagccactatgagaagctgaagggctcccccgaggataatgagcagaa
acagctgtttgtggaacagcacaagcactacctggacgagatcatcgagcagatcagcgagttctccaagagagtgatcctggccga
cgctaatctggacaaagtgctgtccgcctacaacaagcaccgggataagcccatcagagagcaggccgagaatatcatccacctgttt
accctgaccaatctgggagcccctgccgccttcaagtactttgacaccaccatcgaccggaagaggtacaccagcaccaaagaggtg
ctggacgccaccctgatccaccagagcatcaccggcctgtacgagacacggatcgacctgtctcagctgggaggcgactctggagg
atctagcggaggatcctctggcagcgagacaccaggaacaagcgagtcagcaacaccagagagcagtggcggcagcagcggcg
gcagcagcATGACGGCAAAACTGGAAAGCCACGTGCCTGCTGCACCACCAGTCTCAG
CGGAAGCGCCGGCACCGACGCGACCTGACGCTGCTAAGCAGGAGGCTCGACGAG
CCCACCATGAAGCCCTGAGACTGCGGTGGAAAGCAATCGAGGAAGCTGGGGGCA
CTGACGCCTGGGTACGACAGCAGTTGGTTGCGAAAGGCGTAGCGGCGGAAGAGG
TGGACTTCGAAAGTCTCTCCGACAAGCAAAAAGCGGCTTGGAAAGAGAAAAAA
AAGCTGAGGCAACCGAGCGGAGGGCTCAAAAACGACTTGCCTGGGAAGCCTGG
AAAGCGACCCATATTCATCATTTGGGCGTTGGAGTTCACTGGGACGAAGCCGGC
GGGCCTGACAAGTTCGATGTGGCAGGGAGGGAAGAAAGGGCAAAGGCAAACGG
GTTGCCCGAAGGTCTTGACAGCGTAGAAGCATTGGCGAAAGCGCTGGGAATCTC
AGTATCCCGATTGCGCTGGTTCTCTTTTCACAGGGAAGTTGATACAGGGACACAC
TACCAAACTTGGGAGATCCCAAAGAGGGATGGGGGCAAACGCACCCTCACGGCC
CCGAAAAGGGAGCTTAAGGCAGTACAAAGGTGGGTACTCGCCAATGTGGTGGAA
AGACTTCCCGTACACGGAGCTGCTCATGGGTTTGTTGCAGGACGCTCCATACTCA
CTAATGCGCTCGCTCACCAAGGAGCTGACGTGGTTGTTAAAGTTGATATGAAAG
ATTTTTTTCCCAGTGTCACATGGCCTCGAGTTAAAGGCCTGCTCAGAAAGGGAGG
TTTGCCTGAAAACTTGGCGACCCTGCTCGCCCTTCTGAGTACCGAAGCTCCACGG
GAAGTAGTCCGATTCAGAGGGGAAACGCTGTACGTCGCTAAGGGCCCACGCGCT
CTGCCACAGGGCGCGCCTACCAGTCCTGCGCTTACTAACGCATTGTGTCTTAGGT
TGGATAAGCGGTTGAGCGCGCTGAGTAAGCGGTTGGGTTTCACATACACCAGAT
ACGCTGACGATCTTACATTTAGTTGGAGGCGCGCTAAAAAATCTCGCCAGAAAG
AGTTGCCGCTGGCTGATGCTCCCGTCGCGCTTCTCCTGGCCCGGGTTAAGGGAGT
GTTGGAAGCAGAAGGTTTCACTTTGCATCCTGATAAGACCAGGGTTCAGCGGAA
GGGAAGCCGGCAACGAGTGACCGGCCTCGTGGTTAATGAAGCGCCCGAAGGAGT
GCCAGGTGCCCGAGTTCCCCGAGATGTTGTTCGGAGATTGCGAGCGGCAATACA
CAATAGAGAGCAAGGTAAACCAGGGCCGACTGGAGAAACGTTGGAACAACTCA
AAGGCTTGGCAGCTTTTCTGCACATGACAGACGCTGAAAAAGGAAGAGCGTTTC
TGAGAAGACTTGAGGCCCTGGAGAAGAGGCAAACTGCGaaaaggccggcggccacgaaaaag
gccggccaggcaaaaaagaaaaagcttgagggcagaggaagtctgctaacatgcggtgacgtggaggagaatcccggccctgcta
gcatggtgagcaagggcgaggaggataacatggccatcatcaaggagttcatgcgcttcaaggtgcacatggagggctccgtgaac
ggccacgagttcgagatcgagggcgagggcgagggccgcccctacgagggcacccagaccgccaagctgaaggtgaccaagg
gtggcccccctgcccttcgcctgggacatcctgtcccctcagttcatgtacggctccaaggcctacgtgaagcaccccgccgacatccc
cgactacttgaagctgtccttccccgagggcttcaagtgggagcgcgtgatgaacttcgaggacggcggcgtggtgaccgtgaccca
ggactcctccctgcaggacggcgagttcatctacaaggtgaagctgcgcggcaccaacttcccctccgacggccccgtaatgcagaa
gaagaccatgggctgggaggcctcctccgagcggatgtaccccgaggacggcgccctgaagggcgagatcaagcagaggctgaa
gctgaaggacggcggccactacgacgctgaggtcaagaccacctacaaggccaagaagcccgtgcagctgcccggcgcctacaa
cgtcaacatcaagttggacatcacctcccacaacgaggactacaccatcgtggaacagtacgaacgcgccgagggccgccactcca
ccggcggcatggacgagctgtacaagtgagaattcctagagctcgctgatcagcctcgactgtgccttctagttgccagccatctgttgt
ttgcccctcccccgtgccttccttgaccctggaaggtgccactcccactgtcctttcctaataaaatgaggaaattgcatcgcattgtctga
gtaggtgtcattctattctggggggtggggtggggcaggacagcaagggggaggattgggaagagaatagcaggcatgctgggga
gcggccgcaggaacccctagtgatggagttggccactccctctctgcgcgctcgctcgctcactgaggccgggcgaccaaaggtcg
cccgacgcccgggctttgcccgggcggcctcagtgagcgagcgagcgcgcagctgcctgcaggggcgcctgatgcggtattttctc
cttacgcatctgtgcggtatttcacaccgcatacgtcaaagcaaccatagtacgcgccctgtagcggcgcattaagcgcggcgggtgt

ggtggttacgcgcagcgtgaccgctacacttgccagcgccctagcgcccgctcctttcgctttcttcccttcctttctcgccacgttcgcc
ggctttccccgtcaagctctaaatcgggggctccctttagggttccgatttagtgctttacggcacctcgaccccaaaaaacttgatttgg
gtgatggttcacgtagtgggccatcgccctgatagacggtttttcgccctttgacgttggagtccacgttctttaatagtggactcttgttcc
aaactggaacaacactcaaccctatctcgggctattcttttgatttataagggattttgccgatttcggCctattggttaaaaaatgagctga
tttaacaaaaatttaacgcgaattttaacaaaatattaacgtttacaattttatggtgcactctcagtacaatctgctctgatgccgcatagtta
agccagccccgacacccgccaacacccgctgacgcgccctgacgggcttgtctgctcccggcatccgcttacagacaagctgtgac
cgtctccgggagctgcatgtgtcagaggttttcaccgtcatcaccgaaacgcgcgagacgaaagggcctcgtgatacgcctatttttat
aggttaatgtcatgataataatggtttcttagacgtcaggtggcacttttcggggaaatgtgcgcggaacccctatttgtttatttttctaaata
cattcaaatatgtatccgctcatgagacaataaccctgataaatgcttcaataatattgaaaaaggaagagtatgagtattcaacatttccgt
gtcgcccttattcccttttttgcggcattttgccttcctgtttttgctcacccagaaacgctggtgaaagtaaaagatgctgaagatcagttgg
gtgcacgagtgggttacatcgaactggatctcaacagcggtaagatccttgagagttttcgccccgaagaacgttttccaatgatgagc
acttttaaagttctgctatgtggcgcggtattatcccgtattgacgccgggcaagagcaactcggtcgccgcatacactattctcagaatg
acttggttgagtactcaccagtcacagaaaagcatcttacggatggcatgacagtaagagaattatgcagtgctgccataaccatgagt
gataacactgcggccaacttacttctgacaacgatcggaggaccgaaggagctaaccgcttttttgcacaacatgggggatcatgtaac
tcgccttgatcgttgggaaccggagctgaatgaagccataccaaacgacgagcgtgacaccacgatgcctgtagcaatggcaacaac
gttgcgcaaactattaactggcgaactacttactctagcttcccggcaacaattaatagactggatggaggcggataaagttgcaggac
cacttctgcgctcggcccttccggctggctggtttattgctgataaatctggagccggtgagcgtggaagccgcggtatcattgcagca
ctggggccagatggtaagccctcccgtatcgtagttatctacacgacggggagtcaggcaactatggatgaacgaaatagacagatc
gctgagataggtgcctcactgattaagcattggtaactgtcagaccaagtttactcatatatactttagattgatttaaaacttcatttttaattt
aaaaggatctaggtgaagatcctttttgataatctcatgaccaaaatcccttaacgtgagttttcgttccactgagcgtcagaccccgtaga
aaagatcaaaggatcttcttgagatcctttttttctgcgcgtaatctgctgcttgcaaacaaaaaaaccaccgctaccagcggtggtttgttt
gccggatcaagagctaccaactctttttccgaaggtaactggcttcagcagagcgcagataccaaatactgtccttctagtgtagccgta
gttaggccaccacttcaagaactctgtagcaccgcctacatacctcgctctgctaatcctgttaccagtggctgctgccagtggcgataa
gtcgtgtcttaccgggttggactcaagacgatagttaccggataaggcgcagcggtcgggctgaacggggggttcgtgcacacagcc
cagcttggagcgaacgacctacaccgaactgagatacctacagcgtgagctatgagaaagcgccacgcttcccgaagggagaaag
gcggacaggtatccggtaagcggcagggtcggaacaggagagcgcacgagggagcttccagggggaaacgcctggtatctttata
gtcctgtcgggtttcgccacctctgacttgagcgtcgatttttgtgatgctcgtcaggggggcggagcctatggaaaaacgccagcaac
gcggcctttttacggttcctggccttttgctggccttttgctcacatgt

**pBZ203-pU6-sgBFP-Dt_CBh-sv40NLS-Cas9-Sa163RT-NLS-T2A-mCherry**

gagggcctatttcccatgattccttcatatttgcatatacgatacaaggctgttagagagataattggaattaatttgactgtaaacacaaag
atattagtacaaaatacgtgacgtagaaagtaataatttcttgggtagtttgcagttttaaaattatgttttaaaatggactatcatatgcttacc
gtaacttgaaagtatttcgatttcttggctttatatatcttgtggaaaggacgaaacaccGCTGAAGCACTGCACGCCAT
gttttagagctagaaatagcaagttaaaataaggctagtccgttatcaacttgaaaaagtggcaccgagtcggtgcCGTACGGT
GCGAGCGACCGAGAGAGGTCCCAAGCCATCAGCCTCAGCGCCTCGAGCGCGAGA
GCGGCGTTGCGCCGCTCTGGTTGAATTGCAGGACACTCTCCGCAAGGTAGCCTGT
TCTTGGCTCTCTTCCCTCCGGTGAGTACCTCTCCGGCCGGGGAGCTGAACCAACG
AACTAGTCCCTCGTGACCACCCTGACGTACGGCGTGCAGTGCTTCAGCCGCTACC
CCGACCACATGAAGCAGCACGACTTCTTCAAGTCCGCCATGCCCGAAGGCTACG
TCCAGGAGCGCCCTAGGCGCAACCGCCGTTTCCCCGGCCGGAGAGGTACTCACC
GGAGGGGAGAGCCGGTGAGGCTACCGTGCCCCAGGTGAGAAGGTGGTGCCTTCG
GGCCTCCCTCGACCGCTCGCGCtttttttctagaggtacccgttacataacttacggtaaatggcccgcctggctgac
cgcccaacgacccccgcccattgacgtcaatagtaacgccaatagggactttccattgacgtcaatgggtggagtatttacggtaaact
gcccacttggcagtacatcaagtgtatcatatgccaagtacgccccctattgacgtcaatgacggtaaatggcccgcctggcattgtgc
ccagtacatgaccttatgggactttcctacttggcagtacatctacgtattagtcatcgctattaccatggtcgaggtgagccccacgttct
gcttcactctccccatctcccccccctccccacccccaattttgtatttatttattttttaattattttgtgcagcgatgggggcggggggggg
gggggggcgcgcgccaggcggggcggggcggggcgaggggcggggcggggcgaggcggagaggtgcggcggcagccaat
cagagcggcgcgctccgaaagtttccttttatggcgaggcggcggcggcggcggccctataaaaagcgaagcgcgcggcgggcg

ggagtcgctgcgacgctgccttcgccccgtgccccgctccgccgccgcctcgcgccgcccgccccggctctgactgaccgcgttac
tcccacaggtgagcgggcgggacggcccttctcctccgggctgtaattagctgagcaagaggtaagggtttaagggatggttggttg
gtggggtattaatgtttaattacctggagcacctgcctgaaatcactttttttcaggttggaccggtgccaccatggactataaggaccac
gacggagactacaaggatcatgatattgattacaaagacgatgacgataagatggccccaaagaagaagcggaaggtcggtatcca
cggagtcccagcagccgacaagaagtacagcatcggcctggacatcggcaccaactctgtgggctgggccgtgatcaccgacgag
tacaaggtgcccagcaagaaattcaaggtgctgggcaacaccgaccggcacagcatcaagaagaacctgatcggagccctgctgtt
cgacagcggcgaaacagccgaggccacccggctgaagagaaccgccagaagaagatacaccagacggaagaaccggatctgct
atctgcaagagatcttcagcaacgagatggccaaggtggacgacagcttcttccacagactggaagagtccttcctggtggaagagg
ataagaagcacgagcggcaccccatcttcggcaacatcgtggacgaggtggcctaccacgagaagtaccccaccatctaccacctg
agaaagaaactggtggacagcaccgacaaggccgacctgcggctgatctatctggccctggcccacatgatcaagttccggggcca
cttcctgatcgagggcgacctgaaccccgacaacagcgacgtggacaagctgttcatccagctggtgcagacctacaaccagctgtt
cgaggaaaaccccatcaacgccagcggcgtggacgccaaggccatcctgtctgccagactgagcaagagcagacggctggaaaa
tctgatcgcccagctgcccggcgagaagaagaatggcctgttcggaaacctgattgccctgagcctgggcctgacccccaacttcaa
gagcaacttcgacctggccgaggatgccaaactgcagctgagcaaggacacctacgacgacgacctggacaacctgctggcccag
atcggcgaccagtacgccgacctgtttctggccgccaagaacctgtccgacgccatcctgctgagcgacatcctgagagtgaacacc
gagatcaccaaggcccccctgagcgcctctatgatcaagagatacgacgagcaccaccaggacctgaccctgctgaaagctctcgt
gcggcagcagctgcctgagaagtacaaagagattttcttcgaccagagcaagaacggctacgccggctacattgacggcggagcca
gccaggaagagttctacaagttcatcaagcccatcctggaaaagatggacggcaccgaggaactgctcgtgaagctgaacagagag
gacctgctgcggaagcagcggaccttcgacaacggcagcatcccccaccagatccacctgggagagctgcacgccattctgcggc
ggcaggaagatttttacccattcctgaaggacaaccgggaaaagatcgagaagatcctgaccttccgcatcccctactacgtgggccc
tctggccaggggaaacagcagattcgcctggatgaccagaaagagcgaggaaaccatcaccccctggaacttcgaggaagtggtg
gacaagggcgcttccgcccagagcttcatcgagcggatgaccaacttcgataagaacctgcccaacgagaaggtgctgcccaagca
cagcctgctgtacgagtacttcaccgtgtataacgagctgaccaaagtgaaatacgtgaccgagggaatgagaaagcccgccttcctg
agcggcgagcagaaaaaggccatcgtggacctgctgttcaagaccaaccggaaagtgaccgtgaagcagctgaaagaggactact
tcaagaaaatcgagtgcttcgactccgtggaaatctccggcgtggaagatcggttcaacgcctccctgggcacataccacgatctgct
gaaaattatcaaggacaaggacttcctggacaatgaggaaaacgaggacattctggaagatatcgtgctgaccctgacactgtttgag
gacagagagatgatcgaggaacggctgaaaacctatgcccacctgttcgacgacaaagtgatgaagcagctgaagcggcggagat
acaccggctggggcaggctgagccggaagctgatcaacggcatccgggacaagcagtccggcaagacaatcctggatttcctgaa
gtccgacggcttcgccaacagaaacttcatgcagctgatccacgacgacagcctgacctttaaagaggacatccagaaagcccaggt
gtccggccagggcgatagcctgcacgagcacattgccaatctggccggcagccccgccattaagaagggcatcctgcagacagtg
aaggtggtggacgagctcgtgaaagtgatgggccggcacaagcccgagaacatcgtgatcgaaatggccagagagaaccagacc
acccagaagggacagaagaacagccgcgagagaatgaagcggatcgaagagggcatcaaagagctgggcagccagatcctgaa
agaacaccccgtggaaaacacccagctgcagaacgagaagctgtacctgtactacctgcagaatgggcgggatatgtacgtggacc
aggaactggacatcaaccggctgtccgactacgatgtggaccatatcgtgcctcagagctttctgaaggacgactccatcgacaacaa
ggtgctgaccagaagcgacaagaaccggggcaagagcgacaacgtgccctccgaagaggtcgtgaagaagatgaagaactactg
gcggcagctgctgaacgccaagctgattacccagagaaagttcgacaatctgaccaaggccgagagaggcggcctgagcgaactg
gataaggccggcttcatcaagagacagctggtggaaacccggcagatcacaaagcacgtggcacagatcctggactcccggatgaa
cactaagtacgacgagaatgacaagctgatccgggaagtgaaagtgatcaccctgaagtccaagctggtgtccgatttccggaagga
tttccagttttacaaagtgcgcgagatcaacaactaccaccacgcccacgacgcctacctgaacgccgtcgtgggaaccgccctgatc
aaaaagtaccctaagctggaaagcgagttcgtgtacggcgactacaaggtgtacgacgtgcggaagatgatcgccaagagcgagca
ggaaatcggcaaggctaccgccaagtacttcttctacagcaacatcatgaactttttcaagaccgagattaccctggccaacggcgaga
tccggaagcggcctctgatcgagacaaacggcgaaaccggggagatcgtgtgggataagggccgggattttgccaccgtgcggaa
agtgctgagcatgccccaagtgaatatcgtgaaaaagaccgaggtgcagacaggcggcttcagcaaagagtctatcctgcccaaga
ggaacagcgataagctgatcgccagaaagaaggactgggaccctaagaagtacggcggcttcgacagccccaccgtggcctattct
gtgctggtggtggccaaagtggaaaagggcaagtccaagaaactgaagagtgtgaaagagctgctggggatcaccatcatggaaag
aagcagcttcgagaagaatcccatcgactttctggaagccaagggctacaaagaagtgaaaaaggacctgatcatcaagctgcctaa
gtactccctgttcgagctggaaaacggccggaagagaatgctggcctctgccggcgaactgcagaagggaaacgaactggccctgc
cctccaaatatgtgaacttcctgtacctggccagccactatgagaagctgaagggctcccccgaggataatgagcagaaacagctgttt
gtggaacagcacaagcactacctggacgagatcatcgagcagatcagcgagttctccaagagagtgatcctggccgacgctaatctg
gacaaagtgctgtccgcctacaacaagcaccgggataagcccatcagagagcaggccgagaatatcatccacctgtttaccctgacc

aatctgggagcccctgccgccttcaagtactttgacaccaccatcgaccggaagaggtacaccagcaccaaagaggtgctggacgc
caccctgatccaccagagcatcaccggcctgtacgagacacggatcgacctgtctcagctgggaggcgactctggaggatctagcg
gaggatcctctggcagcgagacaccaggaacaagcgagtcagcaacaccagagagcagtggcggcagcagcggcggcagcag
cATGACGGCAAAACTGGAAAGCCACGTGCCTGCTGCACCACCAGTCTCAGCGGA
AGCGCCGGCACCGACGCGACCTGACGCTGCTAAGCAGGAGGCTCGACGAGCCCA
CCATGAAGCCCTGAGACTGCGGTGGAAAGCAATCGAGGAAGCTGGGGGCACTGA
CGCCTGGGTACGACAGCAGTTGGTTGCGAAAGGCGTAGCGGCGGAAGAGGTGGA
CTTCGAAAGTCTCTCCGACAAGCAAAAAGCGGCTTGGAAAGAGAAAAAAAAAG
CTGAGGCAACCGAGCGGAGGGCTCAAAAACGACTTGCCTGGGAAGCCTGGAAA
GCGACCCATATTCATCATTTGGGCGTTGGAGTTCACTGGGACGAAGCCGGCGGG
CCTGACAAGTTCGATGTGGCAGGGAGGGAAGAAAGGGCAAAGGCAAACGGGTT
GCCCGAAGGTCTTGACAGCGTAGAAGCATTGGCGAAAGCGCTGGGAATCTCAGT
ATCCCGATTGCGCTGGTTCTCTTTTCACAGGGAAGTTGATACAGGGACACACTAC
CAAACTTGGGAGATCCCAAAGAGGGATGGGGGCAAACGCACCCTCACGGCCCCG
AAAAGGGAGCTTAAGGCAGTACAAAGGTGGGTACTCGCCAATGTGGTGGAAAG
ACTTCCCGTACACGGAGCTGCTCATGGGTTTGTTGCAGGACGCTCCATACTCACT
AATGCGCTCGCTCACCAAGGAGCTGACGTGGTTGTTAAAGTTGATATGAAAGATT
TTTTTCCCAGTGTCACATGGCCTCGAGTTAAAGGCCTGCTCAGAAAGGGAGGTTT
GCCTGAAAACTTGGCGACCCTGCTCGCCCTTCTGAGTACCGAAGCTCCACGGGAA
GTAGTCCGATTCAGAGGGGAAACGCTGTACGTCGCTAAGGGCCCACGCGCTCTG
CCACAGGGCGCGCCTACCAGTCCTGCGCTTACTAACGCATTGTGTCTTAGGTTGG
ATAAGCGGTTGAGCGCGCTGAGTAAGCGGTTGGGTTTCACATACACCAGATACG
CTGACGATCTTACATTTAGTTGGAGGCGCGCTAAAAAATCTCGCCAGAAAGAGTT
GCCGCTGGCTGATGCTCCCGTCGCGCTTCTCCTGGCCCGGGTTAAGGGAGTGTTG
GAAGCAGAAGGTTTCACTTTGCATCCTGATAAGACCAGGGTTCAGCGGAAGGGA
AGCCGGCAACGAGTGACCGGCCTCGTGGTTAATGAAGCGCCCGAAGGAGTGCCA
GGTGCCCGAGTTCCCCGAGATGTTGTTCGGAGATTGCGAGCGGCAATACACAAT
AGAGAGCAAGGTAAACCAGGGCCGACTGGAGAAACGTTGGAACAACTCAAAGG
CTTGGCAGCTTTTCTGCACATGACAGACGCTGAAAAAGGAAGAGCGTTTCTGAG
AAGACTTGAGGCCCTGGAGAAGAGGCAAACTGCGaaaaggccggcggccacgaaaaaggccgg
ccaggcaaaaaagaaaaagcttgagggcagaggaagtctgctaacatgcggtgacgtggaggagaatcccggccctgctagcatg
gtgagcaagggcgaggaggataacatggccatcatcaaggagttcatgcgcttcaaggtgcacatggagggctccgtgaacggcca
cgagttcgagatcgagggcgagggcgagggccgcccctacgagggcacccagaccgccaagctgaaggtgaccaagggtggcc
ccctgcccttcgcctgggacatcctgtcccctcagttcatgtacggctccaaggcctacgtgaagcaccccgccgacatccccgacta
cttgaagctgtccttccccgagggcttcaagtgggagcgcgtgatgaacttcgaggacggcggcgtggtgaccgtgacccaggactc
ctccctgcaggacggcgagttcatctacaaggtgaagctgcgcggcaccaacttcccctccgacggccccgtaatgcagaagaaga
ccatgggctgggaggcctcctccgagcggatgtaccccgaggacggcgccctgaagggcgagatcaagcagaggctgaagctga
aggacggcggccactacgacgctgaggtcaagaccacctacaaggccaagaagcccgtgcagctgcccggcgcctacaacgtca
acatcaagttggacatcacctcccacaacgaggactacaccatcgtggaacagtacgaacgcgccgagggccgccactccaccgg
cggcatggacgagctgtacaagtgagaattcctagagctcgctgatcagcctcgactgtgccttctagttgccagccatctgttgtttgcc
cctcccccgtgccttccttgaccctggaaggtgccactcccactgtcctttcctaataaaatgaggaaattgcatcgcattgtctgagtag
gtgtcattctattctggggggtggggtggggcaggacagcaagggggaggattgggaagagaatagcaggcatgctggggagcgg
ccgcaggaacccctagtgatggagttggccactccctctctgcgcgctcgctcgctcactgaggccgggcgaccaaaggtcgcccg
acgcccgggctttgcccgggcggcctcagtgagcgagcgagcgcgcagctgcctgcaggggcgcctgatgcggtattttctccttac
gcatctgtgcggtatttcacaccgcatacgtcaaagcaaccatagtacgcgccctgtagcggcgcattaagcgcggcgggtgtggtg
gttacgcgcagcgtgaccgctacacttgccagcgccctagcgcccgctcctttcgctttcttcccttcctttctcgccacgttcgccggctt
tccccgtcaagctctaaatcgggggctccctttagggttccgatttagtgctttacggcacctcgaccccaaaaaacttgatttgggtgat
ggttcacgtagtgggccatcgccctgatagacggtttttcgccctttgacgttggagtccacgttctttaatagtggactcttgttccaaact
ggaacaacactcaaccctatctcgggctattcttttgatttataagggattttgccgatttcggCctattggttaaaaaatgagctgatttaa
caaaaatttaacgcgaattttaacaaaatattaacgtttacaatttatggtgcactctcagtacaatctgctctgatgccgcatagttaagcc

agccccgacacccgccaacacccgctgacgcgccctgacgggcttgtctgctcccggcatccgcttacagacaagctgtgaccgtct
ccgggagctgcatgtgtcagaggttttcaccgtcatcaccgaaacgcgcgagacgaaagggcctcgtgatacgcctatttttataggtt
aatgtcatgataataatggtttcttagacgtcaggtggcacttttcggggaaatgtgcgcggaacccctatttgtttatttttctaaatacattc
aaatatgtatccgctcatgagacaataaccctgataaatgcttcaataatattgaaaaaggaagagtatgagtattcaacatttccgtgtcg
cccttattcccttttttgcggcattttgccttcctgtttttgctcacccagaaacgctggtgaaagtaaaagatgctgaagatcagttgggtgc
acgagtgggttacatcgaactggatctcaacagcggtaagatccttgagagttttcgccccgaagaacgttttccaatgatgagcactttt
aaagttctgctatgtggcgcggtattatcccgtattgacgccgggcaagagcaactcggtcgccgcatacactattctcagaatgacttg
gttgagtactcaccagtcacagaaaagcatcttacggatggcatgacagtaagagaattatgcagtgctgccataaccatgagtgataa
cactgcggccaacttacttctgacaacgatcggaggaccgaaggagctaaccgcttttttgcacaacatgggggatcatgtaactcgcc
ttgatcgttgggaaccggagctgaatgaagccataccaaacgacgagcgtgacaccacgatgcctgtagcaatggcaacaacgttgc
gcaaactattaactggcgaactacttactctagcttcccggcaacaattaatagactggatggaggcggataaagttgcaggaccacttc
tgcgctcggcccttccggctggctggtttattgctgataaatctggagccggtgagcgtggaagccgcggtatcattgcagcactggg
gccagatggtaagccctcccgtatcgtagttatctacacgacggggagtcaggcaactatggatgaacgaaatagacagatcgctga
gataggtgcctcactgattaagcattggtaactgtcagaccaagtttactcatatatactttagattgatttaaaacttcatttttaatttaaaag
gatctaggtgaagatcctttttgataatctcatgaccaaaatcccttaacgtgagttttcgttccactgagcgtcagaccccgtagaaaaga
tcaaaggatcttcttgagatcctttttttctgcgcgtaatctgctgcttgcaaacaaaaaaaccaccgctaccagcggtggtttgtttgccgg
atcaagagctaccaactctttttccgaaggtaactggcttcagcagagcgcagataccaaatactgtccttctagtgtagccgtagttagg
ccaccacttcaagaactctgtagcaccgcctacatacctcgctctgctaatcctgttaccagtggctgctgccagtggcgataagtcgtg
tcttaccgggttggactcaagacgatagttaccggataaggcgcagcggtcgggctgaacggggggttcgtgcacacagcccagctt
ggagcgaacgacctacaccgaactgagatacctacagcgtgagctatgagaaagcgccacgcttcccgaagggagaaaggcgga
caggtatccggtaagcggcagggtcggaacaggagagcgcacgagggagcttccagggggaaacgcctggtatctttatagtcctg
tcgggtttcgccacctctgacttgagcgtcgatttttgtgatgctcgtcaggggggcggagcctatggaaaaacgccagcaacgcggc
ctttttacggttcctggccttttgctggccttttgctcacatgt

#### pBZ204-pU6-sgBFP-Dn_CBh-sv40NLS-Cas9-Sa163RT-NLS-T2A-mCherry

gagggcctatttcccatgattccttcatatttgcatatacgatacaaggctgttagagagataattggaattaatttgactgtaaacacaaag
atattagtacaaaatacgtgacgtagaaagtaataatttcttgggtagtttgcagttttaaaattatgttttaaaatggactatcatatgcttacc
gtaacttgaaagtatttcgatttcttggctttatatatcttgtggaaaggacgaaacaccGCTGAAGCACTGCACGCCAT
gttttagagctagaaatagcaagttaaaataaggctagtccgttatcaacttgaaaaagtggcaccgagtcggtgcCGTACGGT
GCGAGCGACCGAGAGAGGTCCCAAGCCATCAGCCTCAGCGCCTCGAGCGCGAGA
GCGGCGTTGCGCCGCTCTGGTTGAATTGCAGGACACTCTCCGCAAGGTAGCCTGT
TCTTGGCTCTCTTCCCTCCGGTGAGTACCTCTCCGGCCGGGGAGCTGAACCAACG
AACTAGTGCGCTCCTGGACGTAGCCTTCGGGCATGGCGGACTTGAAGAAGTCGT
GCTGCTTCATGTGGTCGGGGTAGCGGCTGAAGCACTGCACGCCGTACGTCAGGG
TGGTCACGAGGGCCTAGGCGCAACCGCCGTTTCCCCGGCCGGAGAGGTACTCAC
CGGAGGGGAGAGCCGGTGAGGCTACCGTGCCCCAGGTGAGAAGGTGGTGCCTTC
GGGCCTCCCTCGACCGCTCGCGCtttttttctagaggtacccgttacataacttacggtaaatggcccgcctggctg
accgcccaacgacccccgcccattgacgtcaatagtaacgccaatagggactttccattgacgtcaatgggtggagtatttacggtaaa
ctgcccacttggcagtacatcaagtgtatcatatgccaagtacgccccctattgacgtcaatgacggtaaatggcccgcctggcattgtg
cccagtacatgaccttatgggactttcctacttggcagtacatctacgtattagtcatcgctattaccatggtcgaggtgagccccacgttc
tgcttcactctccccatctcccccccctccccacccccaattttgtatttatttattttttaattattttgtgcagcgatgggggcggggggg
gggggggcgcgcgccaggcggggcggggcggggcgaggggcggggcggggcgaggcggagaggtgcggcggcagcca
atcagagcggcgcgctccgaaagtttccttttatggcgaggcggcggcggcggcggccctataaaaagcgaagcgcgcggcgggc
gggagtcgctgcgacgctgccttcgccccgtgccccgctccgccgccgcctcgcgccgcccgccccggctctgactgaccgcgtta
ctcccacaggtgagcgggcgggacggcccttctcctccgggctgtaattagctgagcaagaggtaagggtttaagggatggttggttg
gtggggtattaatgtttaattacctggagcacctgcctgaaatcactttttttcaggttggaccggtgccaccatggactataaggaccac
gacggagactacaaggatcatgatattgattacaaagacgatgacgataagatggccccaaagaagaagcggaaggtcggtatcca
cggagtcccagcagccgacaagaagtacagcatcggcctggacatcggcaccaactctgtgggctgggccgtgatcaccgacgag

tacaaggtgcccagcaagaaattcaaggtgctgggcaacaccgaccggcacagcatcaagaagaacctgatcggagccctgctgtt
cgacagcggcgaaacagccgaggccacccggctgaagagaaccgccagaagaagatacaccagacggaagaaccggatctgct
atctgcaagagatcttcagcaacgagatggccaaggtggacgacagcttcttccacagactggaagagtccttcctggtggaagagg
ataagaagcacgagcggcaccccatcttcggcaacatcgtggacgaggtggcctaccacgagaagtaccccaccatctaccacctg
agaaagaaactggtggacagcaccgacaaggccgacctgcggctgatctatctggccctggcccacatgatcaagttccggggcca
cttcctgatcgagggcgacctgaaccccgacaacagcgacgtggacaagctgttcatccagctggtgcagacctacaaccagctgtt
cgaggaaaaccccatcaacgccagcggcgtggacgccaaggccatcctgtctgccagactgagcaagagcagacggctggaaaa
tctgatcgcccagctgcccggcgagaagaagaatggcctgttcggaaacctgattgccctgagcctgggcctgacccccaacttcaa
gagcaacttcgacctggccgaggatgccaaactgcagctgagcaaggacacctacgacgacgacctggacaacctgctggcccag
atcggcgaccagtacgccgacctgtttctggccgccaagaacctgtccgacgccatcctgctgagcgacatcctgagagtgaacacc
gagatcaccaaggcccccctgagcgcctctatgatcaagagatacgacgagcaccaccaggacctgaccctgctgaaagctctcgt
gcggcagcagctgcctgagaagtacaaagagattttcttcgaccagagcaagaacggctacgccggctacattgacggcggagcca
gccaggaagagttctacaagttcatcaagcccatcctggaaaagatggacggcaccgaggaactgctcgtgaagctgaacagagag
gacctgctgcggaagcagcggaccttcgacaacggcagcatcccccaccagatccacctgggagagctgcacgccattctgcggc
ggcaggaagatttttacccattcctgaaggacaaccgggaaaagatcgagaagatcctgaccttccgcatcccctactacgtgggccc
tctggccaggggaaacagcagattcgcctggatgaccagaaagagcgaggaaaccatcaccccctggaacttcgaggaagtggtg
gacaagggcgcttccgcccagagcttcatcgagcggatgaccaacttcgataagaacctgcccaacgagaaggtgctgcccaagca
cagcctgctgtacgagtacttcaccgtgtataacgagctgaccaaagtgaaatacgtgaccgagggaatgagaaagcccgccttcctg
agcggcgagcagaaaaaggccatcgtggacctgctgttcaagaccaaccggaaagtgaccgtgaagcagctgaaagaggactact
tcaagaaaatcgagtgcttcgactccgtggaaatctccggcgtggaagatcggttcaacgcctccctgggcacataccacgatctgct
gaaaattatcaaggacaaggacttcctggacaatgaggaaaacgaggacattctggaagatatcgtgctgaccctgacactgtttgag
gacagagagatgatcgaggaacggctgaaaacctatgcccacctgttcgacgacaaagtgatgaagcagctgaagcggcggagat
acaccggctggggcaggctgagccggaagctgatcaacggcatccgggacaagcagtccggcaagacaatcctggatttcctgaa
gtccgacggcttcgccaacagaaacttcatgcagctgatccacgacgacagcctgacctttaaagaggacatccagaaagcccaggt
gtccggccagggcgatagcctgcacgagcacattgccaatctggccggcagccccgccattaagaagggcatcctgcagacagtg
aaggtggtggacgagctcgtgaaagtgatgggccggcacaagcccgagaacatcgtgatcgaaatggccagagagaaccagacc
acccagaagggacagaagaacagccgcgagagaatgaagcggatcgaagagggcatcaaagagctgggcagccagatcctgaa
agaacaccccgtggaaaacacccagctgcagaacgagaagctgtacctgtactacctgcagaatgggcgggatatgtacgtggacc
aggaactggacatcaaccggctgtccgactacgatgtggaccatatcgtgcctcagagctttctgaaggacgactccatcgacaacaa
ggtgctgaccagaagcgacaagaaccggggcaagagcgacaacgtgccctccgaagaggtcgtgaagaagatgaagaactactg
gcggcagctgctgaacgccaagctgattacccagagaaagttcgacaatctgaccaaggccgagagaggcggcctgagcgaactg
gataaggccggcttcatcaagagacagctggtggaaacccggcagatcacaaagcacgtggcacagatcctggactcccggatgaa
cactaagtacgacgagaatgacaagctgatccgggaagtgaaagtgatcaccctgaagtccaagctggtgtccgatttccggaagga
tttccagttttacaaagtgcgcgagatcaacaactaccaccacgcccacgacgcctacctgaacgccgtcgtgggaaccgccctgatc
aaaaagtaccctaagctggaaagcgagttcgtgtacggcgactacaaggtgtacgacgtgcggaagatgatcgccaagagcgagca
ggaaatcggcaaggctaccgccaagtacttcttctacagcaacatcatgaactttttcaagaccgagattaccctggccaacggcgaga
tccggaagcggcctctgatcgagacaaacggcgaaaccggggagatcgtgtgggataagggccgggattttgccaccgtgcggaa
agtgctgagcatgccccaagtgaatatcgtgaaaaagaccgaggtgcagacaggcggcttcagcaaagagtctatcctgcccaaga
ggaacagcgataagctgatcgccagaaagaaggactgggaccctaagaagtacggcggcttcgacagccccaccgtggcctattct
gtgctggtggtggccaaagtggaaaagggcaagtccaagaaactgaagagtgtgaaagagctgctggggatcaccatcatggaaag
aagcagcttcgagaagaatcccatcgactttctggaagccaagggctacaaagaagtgaaaaaggacctgatcatcaagctgcctaa
gtactccctgttcgagctggaaaacggccggaagagaatgctggcctctgccggcgaactgcagaagggaaacgaactggccctgc
cctccaaatatgtgaacttcctgtacctggccagccactatgagaagctgaagggctcccccgaggataatgagcagaaacagctgttt
gtggaacagcacaagcactacctggacgagatcatcgagcagatcagcgagttctccaagagagtgatcctggccgacgctaatctg
gacaaagtgctgtccgcctacaacaagcaccgggataagcccatcagagagcaggccgagaatatcatccacctgtttaccctgacc
aatctgggagcccctgccgccttcaagtactttgacaccaccatcgaccggaagaggtacaccagcaccaaagaggtgctggacgc
caccctgatccaccagagcatcaccggcctgtacgagacacggatcgacctgtctcagctgggaggcgactctggaggatctagcg
gaggatcctctggcagcgagacaccaggaacaagcgagtcagcaacaccagagagcagtggcggcagcagcggcggcagcag
cATGACGGCAAAACTGGAAAGCCACGTGCCTGCTGCACCACCAGTCTCAGCGGA
AGCGCCGGCACCGACGCGACCTGACGCTGCTAAGCAGGAGGCTCGACGAGCCCA

CCATGAAGCCCTGAGACTGCGGTGGAAAGCAATCGAGGAAGCTGGGGGCACTGA
CGCCTGGGTACGACAGCAGTTGGTTGCGAAAGGCGTAGCGGCGGAAGAGGTGGA
CTTCGAAAGTCTCTCCGACAAGCAAAAAGCGGCTTGGAAAGAGAAAAAAAAAG
CTGAGGCAACCGAGCGGAGGGCTCAAAAACGACTTGCCTGGGAAGCCTGGAAA
GCGACCCATATTCATCATTTGGGCGTTGGAGTTCACTGGGACGAAGCCGGCGGG
CCTGACAAGTTCGATGTGGCAGGGAGGGAAGAAAGGGCAAAGGCAAACGGGTT
GCCCGAAGGTCTTGACAGCGTAGAAGCATTGGCGAAAGCGCTGGGAATCTCAGT
ATCCCGATTGCGCTGGTTCTCTTTTCACAGGGAAGTTGATACAGGGACACACTAC
CAAACTTGGGAGATCCCAAAGAGGGATGGGGGCAAACGCACCCTCACGGCCCCG
AAAAGGGAGCTTAAGGCAGTACAAAGGTGGGTACTCGCCAATGTGGTGGAAAG
ACTTCCCGTACACGGAGCTGCTCATGGGTTTGTTGCAGGACGCTCCATACTCACT
AATGCGCTCGCTCACCAAGGAGCTGACGTGGTTGTTAAAGTTGATATGAAAGATT
TTTTTCCCAGTGTCACATGGCCTCGAGTTAAAGGCCTGCTCAGAAAGGGAGGTTT
GCCTGAAAACTTGGCGACCCTGCTCGCCCTTCTGAGTACCGAAGCTCCACGGGAA
GTAGTCCGATTCAGAGGGGAAACGCTGTACGTCGCTAAGGGCCCACGCGCTCTG
CCACAGGGCGCGCCTACCAGTCCTGCGCTTACTAACGCATTGTGTCTTAGGTTGG
ATAAGCGGTTGAGCGCGCTGAGTAAGCGGTTGGGTTTCACATACACCAGATACG
CTGACGATCTTACATTTAGTTGGAGGCGCGCTAAAAAATCTCGCCAGAAAGAGTT
GCCGCTGGCTGATGCTCCCGTCGCGCTTCTCCTGGCCCGGGTTAAGGGAGTGTTG
GAAGCAGAAGGTTTCACTTTGCATCCTGATAAGACCAGGGTTCAGCGGAAGGGA
AGCCGGCAACGAGTGACCGGCCTCGTGGTTAATGAAGCGCCCGAAGGAGTGCCA
GGTGCCCGAGTTCCCCGAGATGTTGTTCGGAGATTGCGAGCGGCAATACACAAT
AGAGAGCAAGGTAAACCAGGGCCGACTGGAGAAACGTTGGAACAACTCAAAGG
CTTGGCAGCTTTTCTGCACATGACAGACGCTGAAAAAGGAAGAGCGTTTCTGAG
AAGACTTGAGGCCCTGGAGAAGAGGCAAACTGCGaaaaggccggcggccacgaaaaaggccgg
ccaggcaaaaaagaaaaagcttgagggcagaggaagtctgctaacatgcggtgacgtggaggagaatcccggccctgctagcatg
gtgagcaagggcgaggaggataacatggccatcatcaaggagttcatgcgcttcaaggtgcacatggagggctccgtgaacggcca
cgagttcgagatcgagggcgagggcgagggccgcccctacgagggcacccagaccgccaagctgaaggtgaccaagggtggcc
ccctgcccttcgcctgggacatcctgtcccctcagttcatgtacggctccaaggcctacgtgaagcaccccgccgacatccccgacta
cttgaagctgtccttccccgagggcttcaagtgggagcgcgtgatgaacttcgaggacggcggcgtggtgaccgtgacccaggactc
ctccctgcaggacggcgagttcatctacaaggtgaagctgcgcggcaccaacttcccctccgacggccccgtaatgcagaagaaga
ccatgggctgggaggcctcctccgagcggatgtaccccgaggacggcgccctgaagggcgagatcaagcagaggctgaagctga
aggacggcggccactacgacgctgaggtcaagaccacctacaaggccaagaagcccgtgcagctgcccggcgcctacaacgtca
acatcaagttggacatcacctcccacaacgaggactacaccatcgtggaacagtacgaacgcgccgagggccgccactccaccgg
cggcatggacgagctgtacaagtgagaattcctagagctcgctgatcagcctcgactgtgccttctagttgccagccatctgttgtttgcc
cctcccccgtgccttccttgaccctggaaggtgccactcccactgtcctttcctaataaaatgaggaaattgcatcgcattgtctgagtag
gtgtcattctattctggggggtggggtggggcaggacagcaagggggaggattgggaagagaatagcaggcatgctggggagcgg
ccgcaggaacccctagtgatggagttggccactccctctctgcgcgctcgctcgctcactgaggccgggcgaccaaaggtcgcccg
acgcccgggctttgcccgggcggcctcagtgagcgagcgagcgcgcagctgcctgcaggggcgcctgatgcggtattttctccttac
gcatctgtgcggtatttcacaccgcatacgtcaaagcaaccatagtacgcgccctgtagcggcgcattaagcgcggcgggtgtggtg
gttacgcgcagcgtgaccgctacacttgccagcgccctagcgcccgctcctttcgctttcttcccttcctttctcgccacgttcgccggctt
tccccgtcaagctctaaatcgggggctccctttagggttccgatttagtgctttacggcacctcgaccccaaaaaacttgatttgggtgat
ggttcacgtagtgggccatcgccctgatagacggtttttcgccctttgacgttggagtccacgttctttaatagtggactcttgttccaaact
ggaacaacactcaaccctatctcgggctattcttttgatttataagggattttgccgatttcggCctattggttaaaaaatgagctgatttaa
caaaatttaacgcgaattttaacaaaatattaacgtttacaattttatggtgcactctcagtacaatctgctctgatgccgcatagttaagcc
agccccgacacccgccaacacccgctgacgcgccctgacgggcttgtctgctcccggcatccgcttacagacaagctgtgaccgtct
ccgggagctgcatgtgtcagaggttttcaccgtcatcaccgaaacgcgcgagacgaaagggcctcgtgatacgcctatttttataggtt
aatgtcatgataataatggtttcttagacgtcaggtggcacttttcggggaaatgtgcgcggaacccctatttgtttatttttctaaatacattc
aaatatgtatccgctcatgagacaataaccctgataaatgcttcaataatattgaaaaaggaagagtatgagtattcaacatttccgtgtcg
cccttattcccttttttgcggcattttgccttcctgtttttgctcacccagaaacgctggtgaaagtaaaagatgctgaagatcagttgggtgc

acgagtgggttacatcgaactggatctcaacagcggtaagatccttgagagttttcgccccgaagaacgttttccaatgatgagcacttttaaagttctgctatgtggcgcggtattatcccgtattgacgccgggcaagagcaactcggtcgccgcatacactattctcagaatgacttggttgagtactcaccagtcacagaaaagcatcttacggatggcatgacagtaagagaattatgcagtgctgccataaccatgagtgataacactgcggccaacttacttctgacaacgatcggaggaccgaaggagctaaccgcttttttgcacaacatgggggatcatgtaactcgccttgatcgttgggaaccggagctgaatgaagccataccaaacgacgagcgtgacaccacgatgcctgtagcaatggcaacaacgttgcgcaaactattaactggcgaactacttactctagcttcccggcaacaattaatagactggatggaggcggataaagttgcaggaccacttctgcgctcggcccttccggctggctggtttattgctgataaatctggagccggtgagcgtggaagccgcggtatcattgcagcactggggccagatggtaagccctcccgtatcgtagttatctacacgacggggagtcaggcaactatggatgaacgaaatagacagatcgctgagataggtgcctcactgattaagcattggtaactgtcagaccaagtttactcatatatactttagattgatttaaaacttcatttttaatttaaaaggatctaggtgaagatcctttttgataatctcatgaccaaaatcccttaacgtgagttttcgttccactgagcgtcagaccccgtagaaaagatcaaaggatcttcttgagatcctttttttctgcgcgtaatctgctgcttgcaaacaaaaaaaccaccgctaccagcggtggtttgtttgccggatcaagagctaccaactctttttccgaaggtaactggcttcagcagagcgcagataccaaatactgtccttctagtgtagccgtagttaggccaccacttcaagaactctgtagcaccgcctacatacctcgctctgctaatcctgttaccagtggctgctgccagtggcgataagtcgtgtcttaccgggttggactcaagacgatagttaccggataaggcgcagcggtcgggctgaacggggggttcgtgcacacagcccagcttggagcgaacgacctacaccgaactgagatacctacagcgtgagctatgagaaagcgccacgcttcccgaagggagaaaggcggacaggtatccggtaagcggcagggtcggaacaggagagcgcacgagggagcttccagggggaaacgcctggtatctttatagtcctgtcgggtttcgccacctctgacttgagcgtcgatttttgtgatgctcgtcaggggggcggagcctatggaaaaacgccagcaacgcggcctttttacggttcctggccttttgctggccttttgctcacatgt

**pBZ207-pU6-sgBFP-Ht_CBh-sv40NLS-Cas9-Ec86RT-NLS-T2A-mCherry**

catgtgagggcctatttcccatgattccttcatatttgcatatacgatacaaggctgttagagagataattggaattaatttgactgtaaacacaaagatattagtacaaaatacgtgacgtagaaagtaataatttcttgggtagtttgcagttttaaaattatgttttaaaatggactatcatatgcttaccgtaacttgaaagtatttcgatttcttggctttatatatcttgtggaaaggacgaaacaccGCTGAAGCACTGCACGCCATgttttagagctagaaatagcaagttaaaataaggctagtccgttatcaacttgaaaaagtggcaccgagtcggtgcCGTACGATGCGCACCCTTAGCGAGAGGTTTATCATTAAGGTCAACCTCTGGATGTTGTTTCGGCATCCTGCATTGAATCTGAGTTACTGTCTGTTTcCCTACTAGTGCCACCTACGGCAAGCTGACCCTGAAGTTCATCTGCACCACCGGCAAGCTGCCCGTGCCCTGGCCCACCCTCGTGACCACCCTGACGTACGGCGTGCAGTGCTTCAGCCGCTACCCCGACCACATGAcctaggAGGGAACCCGTTTCTTCTGACGTAAGGGTGCGCAttttttctagaggtacccgttacataacttacggtaaatggcccgcctggctgaccgcccaacgacccccgcccattgacgtcaatagtaacgccaatagggactttccattgacgtcaatgggtggagtatttacggtaaactgcccacttggcagtacatcaagtgtatcatatgccaagtacgccccctattgacgtcaatgacggtaaatggcccgcctggcattgtgcccagtacatgaccttatgggactttcctacttggcagtacatctacgtattagtcatcgctattaccatggtcgaggtgagccccacgttctgcttcactctccccatctcccccccctccccacccccaattttgtatttatttattttttaattattttgtgcagcgatgggggcgggggggggggggggcgcgcgccaggcggggcggggcggggcgaggggcggggcggggcgaggcggagaggtgcggcggcagccaatcagagcggcgcgctccgaaagtttccttttatggcgaggcggcggcggcggcggccctataaaaagcgaagcgcgcggcgggcgggagtcgctgcgacgctgccttcgccccgtgccccgctccgccgccgcctcgcgccgcccgccccggctctgactgaccgcgttactcccacaggtgagcgggcgggacggcccttctcctccgggctgtaattagctgagcaagaggtaagggtttaagggatggttggttggtggggtattaatgtttaattacctggagcacctgcctgaaatcactttttttcaggttggaccggtgccaccatggactataaggaccacgacggagactacaaggatcatgatattgattacaaagacgatgacgataagatggccccaaagaagaagcggaaggtcggtatccacggagtcccagcagccgacaagaagtacagcatcggcctggacatcggcaccaactctgtgggctgggccgtgatcaccgacgagtacaaggtgcccagcaagaaattcaaggtgctgggcaacaccgaccggcacagcatcaagaagaacctgatcggagccctgctgttcgacagcggcgaaacagccgaggccacccggctgaagagaaccgccagaagaagatacaccagacggaagaaccggatctgctatctgcaagagatcttcagcaacgagatggccaaggtggacgacagcttcttccacagactggaagagtccttcctggtggaagaggataagaagcacgagcggcaccccatcttcggcaacatcgtggacgaggtggcctaccacgagaagtaccccaccatctaccacctgagaaagaaactggtggacagcaccgacaaggccgacctgcggctgatctatctggccctggcccacatgatcaagttccggggccacttcctgatcgagggcgacctgaaccccgacaacagcgacgtggacaagctgttcatccagctggtgcagacctacaaccagctgttcgaggaaaaccccatcaacgccagcggcgtggacgccaaggccatcctgtctgccagactgagcaagagcagacggctggaaaatctgatcgcccagctgcccggcgagaagaagaatggcctgttcggaaacct

gattgccctgagcctgggcctgacccccaacttcaagagcaacttcgacctggccgaggatgccaaactgcagctgagcaaggaca
cctacgacgacgacctggacaacctgctggcccagatcggcgaccagtacgccgacctgtttctggccgccaagaacctgtccgac
gccatcctgctgagcgacatcctgagagtgaacaccgagatcaccaaggcccccctgagcgcctctatgatcaagagatacgacga
gcaccaccaggacctgaccctgctgaaagctctcgtgcggcagcagctgcctgagaagtacaaagagattttcttcgaccagagcaa
gaacggctacgccggctacattgacggcggagccagccaggaagagttctacaagttcatcaagcccatcctggaaaagatggacg
gcaccgaggaactgctcgtgaagctgaacagagaggacctgctgcggaagcagcggaccttcgacaacggcagcatcccccacc
agatccacctgggagagctgcacgccattctgcggcggcaggaagatttttacccattcctgaaggacaaccgggaaaagatcgaga
agatcctgaccttccgcatcccctactacgtgggccctctggccaggggaaacagcagattcgcctggatgaccagaaagagcgag
gaaaccatcaccccctggaacttcgaggaagtggtggacaagggcgcttccgcccagagcttcatcgagcggatgaccaacttcgat
aagaacctgcccaacgagaaggtgctgcccaagcacagcctgctgtacgagtacttcaccgtgtataacgagctgaccaaagtgaaa
tacgtgaccgagggaatgagaaagcccgccttcctgagcggcgagcagaaaaaggccatcgtggacctgctgttcaagaccaacc
ggaaagtgaccgtgaagcagctgaaagaggactacttcaagaaaatcgagtgcttcgactccgtggaaatctccggcgtggaagatc
ggttcaacgcctccctgggcacataccacgatctgctgaaaattatcaaggacaaggacttcctggacaatgaggaaaacgaggacat
tctggaagatatcgtgctgaccctgacactgtttgaggacagagagatgatcgaggaacggctgaaaacctatgcccacctgttcgac
gacaaagtgatgaagcagctgaagcggcggagatacaccggctggggcaggctgagccggaagctgatcaacggcatccgggac
aagcagtccggcaagacaatcctggatttcctgaagtccgacggcttcgccaacagaaacttcatgcagctgatccacgacgacagc
ctgacctttaaagaggacatccagaaagcccaggtgtccggccagggcgatagcctgcacgagcacattgccaatctggccggcag
ccccgccattaagaagggcatcctgcagacagtgaaggtggtggacgagctcgtgaaagtgatgggccggcacaagcccgagaac
atcgtgatcgaaatggccagagagaaccagaccacccagaagggacagaagaacagccgcgagagaatgaagcggatcgaaga
gggcatcaaagagctgggcagccagatcctgaaagaacaccccgtggaaaacacccagctgcagaacgagaagctgtacctgtac
tacctgcagaatgggcgggatatgtacgtggaccaggaactggacatcaaccggctgtccgactacgatgtggaccatatcgtgcctc
agagctttctgaaggacgactccatcgacaacaaggtgctgaccagaagcgacaagaaccggggcaagagcgacaacgtgccctc
cgaagaggtcgtgaagaagatgaagaactactggcggcagctgctgaacgccaagctgattacccagagaaagttcgacaatctga
ccaaggccgagagaggcggcctgagcgaactggataaggccggcttcatcaagagacagctggtggaaacccggcagatcacaa
agcacgtggcacagatcctggactcccggatgaacactaagtacgacgagaatgacaagctgatccgggaagtgaaagtgatcacc
ctgaagtccaagctggtgtccgatttccggaaggatttccagttttacaaagtgcgcgagatcaacaactaccaccacgcccacgacg
cctacctgaacgccgtcgtgggaaccgccctgatcaaaaagtaccctaagctggaaagcgagttcgtgtacggcgactacaaggtgt
acgacgtgcggaagatgatcgccaagagcgagcaggaaatcggcaaggctaccgccaagtacttcttctacagcaacatcatgaact
tttcaagaccgagattaccctggccaacggcgagatccggaagcggcctctgatcgagacaaacggcgaaaccggggagatcgtg
tgggataagggccgggattttgccaccgtgcggaaagtgctgagcatgccccaagtgaatatcgtgaaaaagaccgaggtgcagac
aggcggcttcagcaaagagtctatcctgcccaagaggaacagcgataagctgatcgccagaaagaaggactgggaccctaagaagt
acggcggcttcgacagccccaccgtggcctattctgtgctggtggtggccaaagtggaaaagggcaagtccaagaaactgaagagt
gtgaaagagctgctggggatcaccatcatggaaagaagcagcttcgagaagaatcccatcgactttctggaagccaagggctacaaa
gaagtgaaaaaggacctgatcatcaagctgcctaagtactccctgttcgagctggaaaacggccggaagagaatgctggcctctgcc
ggcgaactgcagaagggaaacgaactggccctgccctccaaatatgtgaacttcctgtacctggccagccactatgagaagctgaag
ggctcccccgaggataatgagcagaaacagctgtttgtggaacagcacaagcactacctggacgagatcatcgagcagatcagcga
gttctccaagagagtgatcctggccgacgctaatctggacaaagtgctgtccgcctacaacaagcaccgggataagcccatcagaga
gcaggccgagaatatcatccacctgtttaccctgaccaatctgggagcccctgccgccttcaagtactttgacaccaccatcgaccgga
agaggtacaccagcaccaaagaggtgctggacgccaccctgatccaccagagcatcaccggcctgtacgagacacggatcgacct
gtctcagctgggaggcgactctggaggatctagcggaggatcctctggcagcgagacaccaggaacaagcgagtcagcaacacca
gagagcagtggcggcagcagcggcggcagcagcAAGAGTGCGGAATACCTTAACACATTTAGGCT
TAGAAACCTGGGACTCCCCGTTATGAATAACCTGCACGATATGTCTAAGGCAACC
CGAATCAGTGTAGAGACGCTTAGACTCTTGATATATACCGCCGACTTTCGATACA
GAATATATACCGTCGAGAAGAAGGGCCCAGAGAAACGAATGCGGACCATATACC
AACCTAGTAGGGAGCTGAAAGCGCTTCAGGGCTGGGTACTTCGAAATATCCTTG
ACAAACTTTCCAGTTCACCGTTCAGCATTGGTTTCGAGAAACACCAGAGTATTCT
GAATAACGCGACACCTCACATAGGAGCCAATTTCATCCTCAATATTGACCTGGAG
GATTTCTTCCCTAGCCTTACTGCCAATAAAGTGTTCGGTGTATTCCACAGCCTCGG
CTACAACCGACTGATTTCATCTGTACTCACAAAAATTTGTTGCTATAAGAACCTG
CTCCCTCAAGGGGCCCCAAGTAGCCCAAAACTGGCAAACCTCATCTGTTCAAAAT

TGGACTATCGAATTCAGGGCTATGCCGGATCCAGGGGCTTGATCTACACGAGAT
ACGCAGACGACCTGACACTTTCAGCACAATCCATGAAGAAAGTTGTTAAAGCGA
GGGATTTTCTTTTTTCCATTATTCCGTCTGAAGGATTGGTTATTAATTCTAAAAAA
ACTTGCATTAGTGGTCCTCGGTCTCAACGAAAGGTTACAGGCCTGGTAATCTCTC
AGGAGAAGGTTGGGATTGGAAGAGAGAAGTATAAAGAGATTCGCGCCAAAATA
CATCATATTTTTTGCGGTAAATCATCTGAAATCGAGCACGTAAGGGGATGGCTTT
CTTTCATTCTTTCTGTAGACTCCAAATCTCACCGACGACTCATCACATATATAAGT
AAACTGGAGAAAAAATATGGGAAAAATCCGCTTAACAAGGCTAAAACTaaaaggccg
gcggccacgaaaaaggccggccaggcaaaaaagaaaaagcttgagggcagaggaagtctgctaacatgcggtgacgtggagga
gaatcccggccctgctagcatggtgagcaagggcgaggaggataacatggccatcatcaaggagttcatgcgcttcaaggtgcacat
ggagggctccgtgaacggccacgagttcgagatcgagggcgagggcgagggccgcccctacgagggcacccagaccgccaag
ctgaaggtgaccaagggtggcccctgcccttcgcctgggacatcctgtcccctcagttcatgtacggctccaaggcctacgtgaagc
accccgccgacatccccgactacttgaagctgtccttccccgagggcttcaagtgggagcgcgtgatgaacttcgaggacggcggc
gtggtgaccgtgacccaggactcctccctgcaggacggcgagttcatctacaaggtgaagctgcgcggcaccaacttcccctccgac
ggccccgtaatgcagaagaagaccatgggctgggaggcctcctccgagcggatgtaccccgaggacggcgccctgaagggcgag
atcaagcagaggctgaagctgaaggacggcggccactacgacgctgaggtcaagaccacctacaaggccaagaagcccgtgcag
ctgcccggcgcctacaacgtcaacatcaagttggacatcacctcccacaacgaggactacaccatcgtggaacagtacgaacgcgc
cgagggccgccactccaccggcggcatggacgagctgtacaagtgagaattcctagagctcgctgatcagcctcgactgtgccttct
agttgccagccatctgttgtttgcccctcccccgtgccttccttgaccctggaaggtgccactcccactgtcctttcctaataaaatgagga
aattgcatcgcattgtctgagtaggtgtcattctattctggggggtggggtggggcaggacagcaagggggaggattgggaagagaa
tagcaggcatgctggggagcggccgcaggaacccctagtgatggagttggccactccctctctgcgcgctcgctcgctcactgaggc
cgggcgaccaaaggtcgcccgacgcccgggctttgcccgggcggcctcagtgagcgagcgagcgcgcagctgcctgcaggggc
gcctgatgcggtattttctccttacgcatctgtgcggtatttcacaccgcatacgtcaaagcaaccatagtacgcgccctgtagcggcgc
attaagcgcggcgggtgtggtggttacgcgcagcgtgaccgctacacttgccagcgccctagcgcccgctcctttcgctttcttcccttc
ctttctcgccacgttcgccggctttccccgtcaagctctaaatcgggggctccctttagggttccgatttagtgctttacggcacctcgacc
ccaaaaaacttgatttgggtgatggttcacgtagtgggccatcgccctgatagacggtttttcgccctttgacgttggagtccacgttcttt
aatagtggactcttgttccaaactggaacaacactcaaccctatctcgggctattcttttgatttataagggattttgccgatttcggCctatt
ggttaaaaaatgagctgatttaacaaaaatttaacgcgaattttaacaaaatattaacgtttacaattttatggtgcactctcagtacaatctg
ctctgatgccgcatagttaagccagccccgacacccgccaacacccgctgacgcgccctgacgggcttgtctgctcccggcatccgc
ttacagacaagctgtgaccgtctccgggagctgcatgtgtcagaggttttcaccgtcatcaccgaaacgcgcgagacgaaagggcct
cgtgatacgcctatttttataggttaatgtcatgataataatggtttcttagacgtcaggtggcacttttcggggaaatgtgcgcggaaccc
ctatttgtttatttttctaaatacattcaaatatgtatccgctcatgagacaataaccctgataaatgcttcaataatattgaaaaaggaagagt
atgagtattcaacatttccgtgtcgcccttattcccttttttgcggcattttgccttcctgtttttgctcacccagaaacgctggtgaaagtaaa
agatgctgaagatcagttgggtgcacgagtgggttacatcgaactggatctcaacagcggtaagatccttgagagttttcgccccgaag
aacgttttccaatgatgagcacttttaaagttctgctatgtggcgcggtattatcccgtattgacgccgggcaagagcaactcggtcgccg
catacactattctcagaatgacttggttgagtactcaccagtcacagaaaagcatcttacggatggcatgacagtaagagaattatgcag
tgctgccataaccatgagtgataacactgcggccaacttacttctgacaacgatcggaggaccgaaggagctaaccgcttttttgcaca
acatgggggatcatgtaactcgccttgatcgttgggaaccggagctgaatgaagccataccaaacgacgagcgtgacaccacgatgc
ctgtagcaatggcaacaacgttgcgcaaactattaactggcgaactacttactctagcttcccggcaacaattaatagactggatggagg
cggataaagttgcaggaccacttctgcgctcggcccttccggctggctggtttattgctgataaatctggagccggtgagcgtggaagc
cgcggtatcattgcagcactggggccagatggtaagccctcccgtatcgtagttatctacacgacggggagtcaggcaactatggatg
aacgaaatagacagatcgctgagataggtgcctcactgattaagcattggtaactgtcagaccaagtttactcatatatactttagattgat
ttaaaacttcatttttaatttaaaaggatctaggtgaagatcctttttgataatctcatgaccaaaatcccttaacgtgagttttcgttccactga
gcgtcagaccccgtagaaaagatcaaaggatcttcttgagatcctttttttctgcgcgtaatctgctgcttgcaaacaaaaaaaccaccgc
taccagcggtggtttgtttgccggatcaagagctaccaactctttttccgaaggtaactggcttcagcagagcgcagataccaaatactg
ttcttctagtgtagccgtagttaggccaccacttcaagaactctgtagcaccgcctacatacctcgctctgctaatcctgttaccagtggct
gctgccagtggcgataagtcgtgtcttaccgggttggactcaagacgatagttaccggataaggcgcagcggtcgggctgaacggg
gggttcgtgcacacagcccagcttggagcgaacgacctacaccgaactgagatacctacagcgtgagctatgagaaagcgccacgc
ttcccgaagggagaaaggcggacaggtatccggtaagcggcagggtcggaacaggagagcgcacgagggagcttccaggggga

aacgcctggtatctttatagtcctgtcgggtttcgccacctctgacttgagcgtcgatttttgtgatgctcgtcaggggggcggagcctatg
gaaaaacgccagcaacgcggc

**pBZ208-pU6-sgBFP-Hn_CBh-sv40NLS-Cas9-Ec86RT-NLS-T2A-mCherry**

catgtgagggcctatttcccatgattccttcatatttgcatatacgatacaaggctgttagagagataattggaattaatttgactgtaaacac
aaagatattagtacaaaatacgtgacgtagaaagtaataatttcttgggtagtttgcagttttaaaattatgttttaaaatggactatcatatgc
ttaccgtaacttgaaagtatttcgatttcttggctttatatatcttgtggaaaggacgaaacaccGCTGAAGCACTGCACGC
CATgttttagagctagaaatagcaagttaaaataaggctagtccgttatcaacttgaaaaagtggcaccgagtcggtgcCGTAC
GATGCGCACCCTTAGCGAGAGGTTTATCATTAAGGTCAACCTCTGGATGTTGTTT
CGGCATCCTGCATTGAATCTGAGTTACTGTCTGTTTcCCTACTAGTTCATGTGGTC
GGGGTAGCGGCTGAAGCACTGCACGCCGTACGTCAGGGTGGTCACGAGGGTGGG
CCAGGGCACGGGCAGCTTGCCGGTGGTGCAGATGAACTTCAGGGTCAGCTTGCC
GTAGGTGGCcctaggAGGGAACCCGTTTCTTCTGACGTAAGGGTGCGCAttttttctagaggt
acccgttacataacttacggtaaatggcccgcctggctgaccgcccaacgacccccgcccattgacgtcaatagtaacgccaatagg
gactttccattgacgtcaatgggtggagtatttacggtaaactgcccacttggcagtacatcaagtgtatcatatgccaagtacgccccct
attgacgtcaatgacggtaaatggcccgcctggcattgtgcccagtacatgaccttatgggactttcctacttggcagtacatctacgtatt
agtcatcgctattaccatggtcgaggtgagccccacgttctgcttcactctccccatctcccccccctccccacccccaattttgtatttattt
atttttaattattttgtgcagcgatgggggcgggggggggggggggcgcgcgccaggcggggcggggcggggcgaggggcg
gggcggggcgaggcggagaggtgcggcggcagccaatcagagcggcgcgctccgaaagtttccttttatggcgaggcggcggc
ggcggcggccctataaaaagcgaagcgcgcggcgggcgggagtcgctgcgacgctgccttcgccccgtgccccgctccgccgcc
gcctcgcgccgcccgccccggctctgactgaccgcgttactcccacaggtgagcgggcgggacggcccttctcctccgggctgtaa
ttagctgagcaagaggtaagggtttaagggatggttggttggtggggtattaatgtttaattacctggagcacctgcctgaaatcactttttt
tcaggttggaccggtgccaccatggactataaggaccacgacggagactacaaggatcatgatattgattacaaagacgatgacgata
agatggccccaaagaagaagcggaaggtcggtatccacggagtcccagcagccgacaagaagtacagcatcggcctggacatcg
gcaccaactctgtgggctgggccgtgatcaccgacgagtacaaggtgcccagcaagaaattcaaggtgctgggcaacaccgaccg
gcacagcatcaagaagaacctgatcggagccctgctgttcgacagcggcgaaacagccgaggccacccggctgaagagaaccgc
cagaagaagatacaccagacggaagaaccggatctgctatctgcaagagatcttcagcaacgagatggccaaggtggacgacagct
tcttccacagactggaagagtccttcctggtggaagaggataagaagcacgagcggcaccccatcttcggcaacatcgtggacgagg
tggcctaccacgagaagtaccccaccatctaccacctgagaaagaaactggtggacagcaccgacaaggccgacctgcggctgatc
tatctggccctggcccacatgatcaagttccggggccacttcctgatcgagggcgacctgaaccccgacaacagcgacgtggacaa
gctgttcatccagctggtgcagacctacaaccagctgttcgaggaaaaccccatcaacgccagcggcgtggacgccaaggccatcct
gtctgccagactgagcaagagcagacggctggaaaatctgatcgcccagctgcccggcgagaagaagaatggcctgttcggaaac
ctgattgccctgagcctgggcctgacccccaacttcaagagcaacttcgacctggccgaggatgccaaactgcagctgagcaaggac
acctacgacgacgacctggacaacctgctggcccagatcggcgaccagtacgccgacctgtttctggccgccaagaacctgtccga
cgccatcctgctgagcgacatcctgagagtgaacaccgagatcaccaaggcccccctgagcgcctctatgatcaagagatacgacg
agcaccaccaggacctgaccctgctgaaagctctcgtgcggcagcagctgcctgagaagtacaaagagattttcttcgaccagagca
agaacggctacgccggctacattgacggcggagccagccaggaagagttctacaagttcatcaagcccatcctggaaaagatggac
ggcaccgaggaactgctcgtgaagctgaacagagaggacctgctgcggaagcagcggaccttcgacaacggcagcatcccccac
cagatccacctgggagagctgcacgccattctgcggcggcaggaagatttttacccattcctgaaggacaaccgggaaaagatcgag
aagatcctgaccttccgcatcccctactacgtgggccctctggccaggggaaacagcagattcgcctggatgaccagaaagagcga
ggaaaccatcaccccctggaacttcgaggaagtggtggacaagggcgcttccgcccagagcttcatcgagcggatgaccaacttcg
ataagaacctgcccaacgagaaggtgctgcccaagcacagcctgctgtacgagtacttcaccgtgtataacgagctgaccaaagtga
aatacgtgaccgagggaatgagaaagcccgccttcctgagcggcgagcagaaaaaggccatcgtggacctgctgttcaagaccaac
cggaaagtgaccgtgaagcagctgaaagaggactacttcaagaaaatcgagtgcttcgactccgtggaaatctccggcgtggaagat
cggttcaacgcctccctgggcacataccacgatctgctgaaaattatcaaggacaaggacttcctggacaatgaggaaaacgaggac
attctggaagatatcgtgctgaccctgacactgtttgaggacagagagatgatcgaggaacggctgaaaacctatgcccacctgttcga
cgacaaagtgatgaagcagctgaagcggcggagatacaccggctggggcaggctgagccggaagctgatcaacggcatccggga
caagcagtccggcaagacaatcctggatttcctgaagtccgacggcttcgccaacagaaacttcatgcagctgatccacgacgacag

cctgacctttaaagaggacatccagaaagcccaggtgtccggccagggcgatagcctgcacgagcacattgccaatctggccggca
gccccgccattaagaagggcatcctgcagacagtgaaggtggtggacgagctcgtgaaagtgatgggccggcacaagcccgagaa
catcgtgatcgaaatggccagagagaaccagaccacccagaagggacagaagaacagccgcgagagaatgaagcggatcgaag
agggcatcaaagagctgggcagccagatcctgaaagaacaccccgtggaaaacacccagctgcagaacgagaagctgtacctgta
ctacctgcagaatgggcgggatatgtacgtggaccaggaactggacatcaaccggctgtccgactacgatgtggaccatatcgtgcct
cagagctttctgaaggacgactccatcgacaacaaggtgctgaccagaagcgacaagaaccggggcaagagcgacaacgtgccct
ccgaagaggtcgtgaagaagatgaagaactactggcggcagctgctgaacgccaagctgattacccagagaaagttcgacaatctg
accaaggccgagagaggcggcctgagcgaactggataaggccggcttcatcaagagacagctggtggaaacccggcagatcaca
aagcacgtggcacagatcctggactcccggatgaacactaagtacgacgagaatgacaagctgatccgggaagtgaaagtgatcac
cctgaagtccaagctggtgtccgatttccggaaggatttccagttttacaaagtgcgcgagatcaacaactaccaccacgcccacgac
gcctacctgaacgccgtcgtgggaaccgccctgatcaaaaagtaccctaagctggaaagcgagttcgtgtacggcgactacaaggtg
tacgacgtgcggaagatgatcgccaagagcgagcaggaaatcggcaaggctaccgccaagtacttcttctacagcaacatcatgaac
tttttcaagaccgagattaccctggccaacggcgagatccggaagcggcctctgatcgagacaaacggcgaaaccggggagatcgt
gtgggataagggccgggattttgccaccgtgcggaaagtgctgagcatgccccaagtgaatatcgtgaaaaagaccgaggtgcaga
caggcggcttcagcaaagagtctatcctgcccaagaggaacagcgataagctgatcgccagaaagaaggactgggaccctaagaa
gtacggcggcttcgacagccccaccgtggcctattctgtgctggtggtggccaaagtggaaaagggcaagtccaagaaactgaaga
gtgtgaaagagctgctggggatcaccatcatggaaagaagcagcttcgagaagaatcccatcgactttctggaagccaagggctaca
aagaagtgaaaaaggacctgatcatcaagctgcctaagtactccctgttcgagctggaaaacggccggaagagaatgctggcctctg
ccggcgaactgcagaagggaaacgaactggccctgccctccaaatatgtgaacttcctgtacctggccagccactatgagaagctga
agggctcccccgaggataatgagcagaaacagctgtttgtggaacagcacaagcactacctggacgagatcatcgagcagatcagc
gagttctccaagagagtgatcctggccgacgctaatctggacaaagtgctgtccgcctacaacaagcaccgggataagcccatcaga
gagcaggccgagaatatcatccacctgtttaccctgaccaatctgggagcccctgccgccttcaagtactttgacaccaccatcgaccg
gaagaggtacaccagcaccaaagaggtgctggacgccaccctgatccaccagagcatcaccggcctgtacgagacacggatcgac
ctgtctcagctgggaggcgactctggaggatctagcggaggatcctctggcagcgagacaccaggaacaagcgagtcagcaacac
cagagagcagtggcggcagcagcggcggcagcagcAAGAGTGCGGAATACCTTAACACATTTAGGC
TTAGAAACCTGGGACTCCCCGTTATGAATAACCTGCACGATATGTCTAAGGCAAC
CCGAATCAGTGTAGAGACGCTTAGACTCTTGATATATACCGCCGACTTTCGATAC
AGAATATATACCGTCGAGAAGAAGGGCCCAGAGAAACGAATGCGGACCATATA
CCAACCTAGTAGGGAGCTGAAAGCGCTTCAGGGCTGGGTACTTCGAAATATCCTT
GACAAACTTTCCAGTTCACCGTTCAGCATTGGTTTCGAGAAACACCAGAGTATTC
TGAATAACGCGACACCTCACATAGGAGCCAATTTCATCCTCAATATTGACCTGGA
GGATTTCTTCCCTAGCCTTACTGCCAATAAAGTGTTCGGTGTATTCCACAGCCTCG
GCTACAACCGACTGATTTCATCTGTACTCACAAAAATTTGTTGCTATAAGAACCT
GCTCCCTCAAGGGGCCCCAAGTAGCCCAAAACTGGCAAACCTCATCTGTTCAAA
ATTGGACTATCGAATTCAGGGCTATGCCGGATCCAGGGGCTTGATCTACACGAG
ATACGCAGACGACCTGACACTTTCAGCACAATCCATGAAGAAAGTTGTTAAAGC
GAGGGATTTTCTTTTTTCCATTATTCCGTCTGAAGGATTGGTTATTAATTCTAAAA
AAACTTGCATTAGTGGTCCTCGGTCTCAACGAAAGGTTACAGGCCTGGTAATCTC
TCAGGAGAAGGTTGGGATTGGAAGAGAGAAGTATAAAGAGATTCGCGCCAAAAT
ACATCATATTTTTTGCGGTAAATCATCTGAAATCGAGCACGTAAGGGGATGGCTT
TCTTTCATTCTTTCTGTAGACTCCAAATCTCACCGACGACTCATCACATATATAAG
TAAACTGGAGAAAAAATATGGGAAAAATCCGCTTAACAAGGCTAAAACTaaaaggc
cggcggccacgaaaaaggccggccaggcaaaaaagaaaaagcttgagggcagaggaagtctgctaacatgcggtgacgtggag
gagaatcccggccctgctagcatggtgagcaagggcgaggaggataacatggccatcatcaaggagttcatgcgcttcaaggtgca
catggagggctccgtgaacggccacgagttcgagatcgagggcgagggcgagggccgcccctacgagggcacccagaccgcca
agctgaaggtgaccaagggtggcccccctgcccttcgcctgggacatcctgtcccctcagttcatgtacggctccaaggcctacgtgaa
gcaccccgccgacatccccgactacttgaagctgtccttccccgagggcttcaagtgggagcgcgtgatgaacttcgaggacggcg
gcgtggtgaccgtgacccaggactcctccctgcaggacggcgagttcatctacaaggtgaagctgcgcggcaccaacttcccctccg
acggccccgtaatgcagaagaagaccatgggctgggaggcctcctccgagcggatgtaccccgaggacggcgccctgaagggcg
agatcaagcagaggctgaagctgaaggacggcggccactacgacgctgaggtcaagaccacctacaaggccaagaagcccgtgc

agctgcccggcgcctacaacgtcaacatcaagttggacatcacctcccacaacgaggactacaccatcgtggaacagtacgaacgc
gccgagggccgccactccaccggcggcatggacgagctgtacaagtgagaattcctagagctcgctgatcagcctcgactgtgcctt
ctagttgccagccatctgttgtttgcccctcccccgtgccttccttgaccctggaaggtgccactcccactgtcctttcctaataaaatgag
gaaattgcatcgcattgtctgagtaggtgtcattctattctggggggtggggtggggcaggacagcaagggggaggattgggaagag
aatagcaggcatgctggggagcggccgcaggaacccctagtgatggagttggccactccctctctgcgcgctcgctcgctcactgag
gccgggcgaccaaaggtcgcccgacgcccgggctttgcccgggcggcctcagtgagcgagcgagcgcgcagctgcctgcaggg
gcgcctgatgcggtattttctccttacgcatctgtgcggtatttcacaccgcatacgtcaaagcaaccatagtacgcgccctgtagcggc
gcattaagcgcggcgggtgtggtggttacgcgcagcgtgaccgctacacttgccagcgccctagcgcccgctcctttcgctttcttccc
ttcctttctcgccacgttcgccggctttccccgtcaagctctaaatcgggggctccctttagggttccgatttagtgctttacggcacctcg
accccaaaaaacttgatttgggtgatggttcacgtagtgggccatcgccctgatagacggtttttcgccctttgacgttggagtccacgtt
ctttaatagtggactcttgttccaaactggaacaacactcaaccctatctcgggctattcttttgatttataagggattttgccgatttcggCc
tattggttaaaaaatgagctgatttaacaaaaatttaacgcgaattttaacaaaatattaacgtttacaattttatggtgcactctcagtacaat
ctgctctgatgccgcatagttaagccagccccgacacccgccaacacccgctgacgcgccctgacgggcttgtctgctcccggcatc
cgcttacagacaagctgtgaccgtctccgggagctgcatgtgtcagaggttttcaccgtcatcaccgaaacgcgcgagacgaaaggg
cctcgtgatacgcctatttttataggttaatgtcatgataataatggtttcttagacgtcaggtggcacttttcggggaaatgtgcgcggaac
ccctatttgtttatttttctaaatacattcaaatatgtatccgctcatgagacaataaccctgataaatgcttcaataatattgaaaaaggaaga
gtatgagtattcaacatttccgtgtcgcccttattcccttttttgcggcattttgccttcctgtttttgctcacccagaaacgctggtgaaagta
aaagatgctgaagatcagttgggtgcacgagtgggttacatcgaactggatctcaacagcggtaagatccttgagagttttcgccccga
agaacgttttccaatgatgagcacttttaaagttctgctatgtggcgcggtattatcccgtattgacgccgggcaagagcaactcggtcgc
cgcatacactattctcagaatgacttggttgagtactcaccagtcacagaaaagcatcttacggatggcatgacagtaagagaattatgc
agtgctgccataaccatgagtgataacactgcggccaacttacttctgacaacgatcggaggaccgaaggagctaaccgcttttttgca
caacatgggggatcatgtaactcgccttgatcgttgggaaccggagctgaatgaagccataccaaacgacgagcgtgacaccacgat
gcctgtagcaatggcaacaacgttgcgcaaactattaactggcgaactacttactctagcttcccggcaacaattaatagactggatgga
ggcggataaagttgcaggaccacttctgcgctcggcccttccggctggctggtttattgctgataaatctggagccggtgagcgtggaa
gccgcggtatcattgcagcactggggccagatggtaagccctcccgtatcgtagttatctacacgacggggagtcaggcaactatgga
tgaacgaaatagacagatcgctgagataggtgcctcactgattaagcattggtaactgtcagaccaagtttactcatatatactttagattg
atttaaaacttcatttttaatttaaaaggatctaggtgaagatcctttttgataatctcatgaccaaaatcccttaacgtgagttttcgttccact
gagcgtcagaccccgtagaaaagatcaaaggatcttcttgagatcctttttttctgcgcgtaatctgctgcttgcaaacaaaaaaaccacc
gctaccagcggtggtttgtttgccggatcaagagctaccaactctttttccgaaggtaactggcttcagcagagcgcagataccaaatac
tgttcttctagtgtagccgtagttaggccaccacttcaagaactctgtagcaccgcctacatacctcgctctgctaatcctgttaccagtgg
ctgctgccagtggcgataagtcgtgtcttaccgggttggactcaagacgatagttaccggataaggcgcagcggtcgggctgaacgg
ggggttcgtgcacacagcccagcttggagcgaacgacctacaccgaactgagatacctacagcgtgagctatgagaaagcgccacg
cttcccgaagggagaaaggcggacaggtatccggtaagcggcagggtcggaacaggagagcgcacgagggagcttccaggggg
aaacgcctggtatctttatagtcctgtcgggtttcgccacctctgacttgagcgtcgatttttgtgatgctcgtcaggggggcggagcctat
ggaaaaacgccagcaacgcggc

**pBZ209-pU6-sgBFP-At_CBh-sv40NLS-Cas9-Ec86RT-NLS-T2A-mCherry**

catgtgagggcctatttcccatgattccttcatatttgcatatacgatacaaggctgttagagagataattggaattaatttgactgtaaacac
aaagatattagtacaaaatacgtgacgtagaaagtaataatttcttgggtagtttgcagttttaaaattatgttttaaaatggactatcatatgc
ttaccgtaacttgaaagtatttcgatttcttggctttatatatcttgtggaaaggacgaaacaccGCTGAAGCACTGCACGC
CATgttttagagctagaaatagcaagttaaaataaggctagtccgttatcaacttgaaaaagtggcaccgagtcggtgcCGTAC
GATGCGCACCCTTAGCGAGAGGTTTATCATTAAGGTCAACCTCTGGATGTTGTTT
CGGCATCCTGCATTGAATCTGAGTTACTGTCTGTTTcCCTACTAGTCCTGAAGTTC
ATCTGCACCACCGGCAAGCTGCCCGTGCCCTGGCCCACCCTCGTGACCACCCTGA
CGTACGGCGTGCAGTGCTTCAGCCGCTACCCCGACCACATGAAGCAGCACGACT
TcctaggAGGGAACCCGTTTCTTCTGACGTAAGGGTGCGCAtttttttctagaggtacccgttacataac
ttacggtaaatggcccgcctggctgaccgcccaacgacccccgcccattgacgtcaataatgacgtatgttcccatagtaacgccaatagggactttccattgacg
tcaatgggtggagtatttacggtaaactgcccacttggcagtacatcaagtgtatcatatgccaagtacgccccctattgacgtcaatgac

ggtaaatggcccgcctggcattgtgcccagtacatgaccttatgggactttcctacttggcagtacatctacgtattagtcatcgctattac
catggtcgaggtgagccccacgttctgcttcactctccccatctcccccccctccccacccccaattttgtatttatttattttttaattattttgt
gcagcgatgggggcggggggggggggggggcgcgcgccaggcggggcggggcggggcgaggggcggggcggggcgag
gcggagaggtgcggcggcagccaatcagagcggcgcgctccgaaagtttccttttatggcgaggcggcggcggcggcggccctat
aaaaagcgaagcgcgcggcgggcgggagtcgctgcgacgctgccttcgccccgtgccccgctccgccgccgcctcgcgccgccc
gccccggctctgactgaccgcgttactcccacaggtgagcgggcgggacggcccttctcctccgggctgtaattagctgagcaagag
gtaagggtttaagggatggttggttggtggggtattaatgtttaattacctggagcacctgcctgaaatcacttttttttcaggttggaccggt
gccaccatggactataaggaccacgacggagactacaaggatcatgatattgattacaaagacgatgacgataagatggccccaaag
aagaagcggaaggtcggtatccacggagtcccagcagccgacaagaagtacagcatcggcctggacatcggcaccaactctgtgg
gctgggccgtgatcaccgacgagtacaaggtgcccagcaagaaattcaaggtgctgggcaacaccgaccggcacagcatcaagaa
gaacctgatcggagccctgctgttcgacagcggcgaaacagccgaggccacccggctgaagagaaccgccagaagaagatacac
cagacggaagaaccggatctgctatctgcaagagatcttcagcaacgagatggccaaggtggacgacagcttcttccacagactgga
agagtccttcctggtggaagaggataagaagcacgagcggcaccccatcttcggcaacatcgtggacgaggtggcctaccacgaga
agtaccccaccatctaccacctgagaaagaaactggtggacagcaccgacaaggccgacctgcggctgatctatctggccctggcc
cacatgatcaagttccggggccacttcctgatcgagggcgacctgaaccccgacaacagcgacgtggacaagctgttcatccagctg
gtgcagacctacaaccagctgttcgaggaaaaccccatcaacgccagcggcgtggacgccaaggccatcctgtctgccagactgag
caagagcagacggctggaaaatctgatcgcccagctgcccggcgagaagaagaatggcctgttcggaaacctgattgccctgagcc
tgggcctgacccccaacttcaagagcaacttcgacctggccgaggatgccaaactgcagctgagcaaggacacctacgacgacgac
ctggacaacctgctggcccagatcggcgaccagtacgccgacctgtttctggccgccaagaacctgtccgacgccatcctgctgagc
gacatcctgagagtgaacaccgagatcaccaaggcccccctgagcgcctctatgatcaagagatacgacgagcaccaccaggacct
gaccctgctgaaagctctcgtgcggcagcagctgcctgagaagtacaaagagattttcttcgaccagagcaagaacggctacgccgg
ctacattgacggcggagccagccaggaagagttctacaagttcatcaagcccatcctggaaaagatggacggcaccgaggaactgc
tcgtgaagctgaacagagaggacctgctgcggaagcagcggaccttcgacaacggcagcatcccccaccagatccacctgggaga
gctgcacgccattctgcggcggcaggaagatttttacccattcctgaaggacaaccgggaaaagatcgagaagatcctgaccttccgc
atcccctactacgtgggccctctggccaggggaaacagcagattcgcctggatgaccagaaagagcgaggaaaccatcaccccctg
gaacttcgaggaagtggtggacaagggcgcttccgcccagagcttcatcgagcggatgaccaacttcgataagaacctgcccaacg
agaaggtgctgcccaagcacagcctgctgtacgagtacttcaccgtgtataacgagctgaccaaagtgaaatacgtgaccgagggaa
tgagaaagcccgccttcctgagcggcgagcagaaaaaggccatcgtggacctgctgttcaagaccaaccggaaagtgaccgtgaa
gcagctgaaagaggactacttcaagaaaatcgagtgcttcgactccgtggaaatctccggcgtggaagatcggttcaacgcctccctg
ggcacataccacgatctgctgaaaattatcaaggacaaggacttcctggacaatgaggaaaacgaggacattctggaagatatcgtgc
tgaccctgacactgtttgaggacagagagatgatcgaggaacggctgaaaacctatgcccacctgttcgacgacaaagtgatgaagc
agctgaagcggcggagatacaccggctggggcaggctgagccggaagctgatcaacggcatccgggacaagcagtccggcaag
acaatcctggatttcctgaagtccgacggcttcgccaacagaaacttcatgcagctgatccacgacgacagcctgacctttaaagagga
catccagaaagcccaggtgtccggccagggcgatagcctgcacgagcacattgccaatctggccggcagccccgccattaagaag
ggcatcctgcagacagtgaaggtggtggacgagctcgtgaaagtgatgggccggcacaagcccgagaacatcgtgatcgaaatgg
ccagagagaaccagaccacccagaagggacagaagaacagccgcgagagaatgaagcggatcgaagagggcatcaaagagct
gggcagccagatcctgaaagaacaccccgtggaaaacacccagctgcagaacgagaagctgtacctgtactacctgcagaatggg
cgggatatgtacgtggaccaggaactggacatcaaccggctgtccgactacgatgtggaccatatcgtgcctcagagctttctgaagg
acgactccatcgacaacaaggtgctgaccagaagcgacaagaaccggggcaagagcgacaacgtgccctccgaagaggtcgtga
agaagatgaagaactactggcggcagctgctgaacgccaagctgattacccagagaaagttcgacaatctgaccaaggccgagaga
ggcggcctgagcgaactggataaggccggcttcatcaagagacagctggtggaaacccggcagatcacaaagcacgtggcacag
atcctggactcccggatgaacactaagtacgacgagaatgacaagctgatccgggaagtgaaagtgatcaccctgaagtccaagctg
gtgtccgatttccggaaggatttccagttttacaaagtgcgcgagatcaacaactaccaccacgcccacgacgcctacctgaacgccgt
cgtgggaaccgccctgatcaaaaagtaccctaagctggaaagcgagttcgtgtacggcgactacaaggtgtacgacgtgcggaaga
tgatcgccaagagcgagcaggaaatcggcaaggctaccgccaagtacttcttctacagcaacatcatgaactttttcaagaccgagatt
accctggccaacggcgagatccggaagcggcctctgatcgagacaaacggcgaaaccggggagatcgtgtgggataagggccgg
gattttgccaccgtgcggaaagtgctgagcatgccccaagtgaatatcgtgaaaaagaccgaggtgcagacaggcggcttcagcaaa
gagtctatcctgcccaagaggaacagcgataagctgatcgccagaaagaaggactgggaccctaagaagtacggcggcttcgaca
gccccaccgtggcctattctgtgctggtggtggccaaagtggaaaagggcaagtccaagaaactgaagagtgtgaaagagctgctg
gggatcaccatcatggaaagaagcagcttcgagaagaatcccatcgactttctggaagccaagggctacaaagaagtgaaaaagga

cctgatcatcaagctgcctaagtactccctgttcgagctggaaaacggccggaagagaatgctggcctctgccggcgaactgcagaa
gggaaacgaactggccctgccctccaaatatgtgaacttcctgtacctggccagccactatgagaagctgaagggctcccccgagga
taatgagcagaaacagctgtttgtggaacagcacaagcactacctggacgagatcatcgagcagatcagcgagttctccaagagagt
gatcctggccgacgctaatctggacaaagtgctgtccgcctacaacaagcaccgggataagcccatcagagagcaggccgagaata
tcatccacctgtttaccctgaccaatctgggagcccctgccgccttcaagtactttgacaccaccatcgaccggaagaggtacaccagc
accaaagaggtgctggacgccaccctgatccaccagagcatcaccggcctgtacgagacacggatcgacctgtctcagctgggagg
cgactctggaggatctagcggaggatcctctggcagcgagacaccaggaacaagcgagtcagcaacaccagagagcagtggcgg
cagcagcggcggcagcagcAAGAGTGCGGAATACCTTAACACATTTAGGCTTAGAAACCT
GGGACTCCCCGTTATGAATAACCTGCACGATATGTCTAAGGCAACCCGAATCAGT
GTAGAGACGCTTAGACTCTTGATATATACCGCCGACTTTCGATACAGAATATATA
CCGTCGAGAAGAAGGGCCCAGAGAAACGAATGCGGACCATATACCAACCTAGTA
GGGAGCTGAAAGCGCTTCAGGGCTGGGTACTTCGAAATATCCTTGACAAACTTTC
CAGTTCACCGTTCAGCATTGGTTTCGAGAAACACCAGAGTATTCTGAATAACGCG
ACACCTCACATAGGAGCCAATTTCATCCTCAATATTGACCTGGAGGATTTCTTCC
CTAGCCTTACTGCCAATAAAGTGTTCGGTGTATTCCACAGCCTCGGCTACAACCG
ACTGATTTCATCTGTACTCACAAAAATTGTTGCTATAAGAACCTGCTCCCTCAA
GGGGCCCCAAGTAGCCCAAAACTGGCAAACCTCATCTGTTCAAAATTGGACTAT
CGAATTCAGGGCTATGCCGGATCCAGGGGCTTGATCTACACGAGATACGCAGAC
GACCTGACACTTTCAGCACAATCCATGAAGAAAGTTGTTAAAGCGAGGGATTTTC
TTTTTTCCATTATTCCGTCTGAAGGATTGGTTATTAATTCTAAAAAAACTTGCATT
AGTGGTCCTCGGTCTCAACGAAAGGTTACAGGCCTGGTAATCTCTCAGGAGAAG
GTTGGGATTGGAAGAGAGAAGTATAAAGAGATTCGCGCCAAAATACATCATATT
TTTTGCGGTAAATCATCTGAAATCGAGCACGTAAGGGGATGGCTTTCTTTCATTC
TTTCTGTAGACTCCAAATCTCACCGACGACTCATCACATATATAAGTAAACTGGA
GAAAAAATATGGAAAAATCCGCTTAACAAGGCTAAAACTaaaaggccggcggccacgaaa
aaggccggccaggcaaaaaagaaaaagcttgagggcagaggaagtctgctaacatgcggtgacgtggaggagaatcccggccct
gctagcatggtgagcaagggcgaggaggataacatggccatcatcaaggagttcatgcgcttcaaggtgcacatggagggctccgt
gaacggccacgagttcgagatcgagggcgagggcgagggccgcccctacgagggcacccagaccgccaagctgaaggtgacca
agggtggccccctgcccttcgcctgggacatcctgtcccctcagttcatgtacggctccaaggcctacgtgaagcaccccgccgacat
ccccgactacttgaagctgtccttccccgagggcttcaagtgggagcgcgtgatgaacttcgaggacggcggcgtggtgaccgtgac
ccaggactcctccctgcaggacggcgagttcatctacaaggtgaagctgcgcggcaccaacttcccctccgacggccccgtaatgca
gaagaagaccatgggctgggaggcctcctccgagcggatgtaccccgaggacggcgccctgaagggcgagatcaagcagaggct
gaagctgaaggacggcggccactacgacgctgaggtcaagaccacctacaaggccaagaagcccgtgcagctgcccggcgccta
caacgtcaacatcaagttggacatcacctcccacaacgaggactacaccatcgtggaacagtacgaacgcgccgagggccgccact
ccaccggcggcatggacgagctgtacaagtgagaattcctagagctcgctgatcagcctcgactgtgccttctagttgccagccatctg
ttgtttgcccctcccccgtgccttccttgaccctggaaggtgccactcccactgtcctttcctaataaaatgaggaaattgcatcgcattgtc
tgagtaggtgtcattctattctggggggtggggtggggcaggacagcaagggggaggattgggaagagaatagcaggcatgctgg
ggagcggccgcaggaacccctagtgatggagttggccactccctctctgcgcgctcgctcgctcactgaggccgggcgaccaaag
gtcgcccgacgcccgggctttgcccgggcggcctcagtgagcgagcgagcgcgcagctgcctgcaggggcgcctgatgcggtatt
ttctccttacgcatctgtgcggtatttcacaccgcatacgtcaaagcaaccatagtacgcgccctgtagcggcgcattaagcgcggcgg
gtgtggtggttacgcgcagcgtgaccgctacacttgccagcgccctagcgcccgctcctttcgctttcttcccttcctttctcgccacgttc
gccggctttccccgtcaagctctaaatcgggggctccctttagggttccgatttagtgctttacggcacctcgaccccaaaaaacttgatt
tgggtgatggttcacgtagtgggccatcgccctgatagacggtttttcgccctttgacgttggagtccacgttctttaatagtggactcttgt
tccaaactggaacaacactcaaccctatctcgggctattcttttgatttataagggattttgccgatttcggCctattggttaaaaaatgagc
tgatttaacaaaaatttaacgcgaattttaacaaaatattaacgtttacaattttatggtgcactctcagtacaatctgctctgatgccgcatag
ttaagccagccccgacacccgccaacacccgctgacgcgccctgacgggcttgtctgctcccggcatccgcttacagacaagctgtg
accgtctccgggagctgcatgtgtcagaggttttcaccgtcatcaccgaaacgcgcgagacgaaagggcctcgtgatacgcctatttt
ataggttaatgtcatgataataatggtttcttagacgtcaggtggcacttttcggggaaatgtgcgcggaacccctatttgtttatttttctaaa
tacattcaaatatgtatccgctcatgagacaataaccctgataaatgcttcaataatattgaaaaaggaagagtatgagtattcaacatttcc
gtgtcgcccttattcccttttttgcggcattttgccttcctgtttttgctcacccagaaacgctggtgaaagtaaaagatgctgaagatcagtt

gggtgcacgagtgggttacatcgaactggatctcaacagcggtaagatccttgagagttttcgccccgaagaacgttttccaatgatga
gcacttttaaagttctgctatgtggcgcggtattatcccgtattgacgccgggcaagagcaactcggtcgccgcatacactattctcaga
atgacttggttgagtactcaccagtcacagaaaagcatcttacggatggcatgacagtaagagaattatgcagtgctgccataaccatg
agtgataacactgcggccaacttacttctgacaacgatcggaggaccgaaggagctaaccgcttttttgcacaacatgggggatcatgt
aactcgccttgatcgttgggaaccggagctgaatgaagccataccaaacgacgagcgtgacaccacgatgcctgtagcaatggcaac
aacgttgcgcaaactattaactggcgaactacttactctagcttcccggcaacaattaatagactggatggaggcggataaagttgcag
gaccacttctgcgctcggcccttccggctggctggtttattgctgataaatctggagccggtgagcgtggaagccgcggtatcattgca
gcactggggccagatggtaagccctcccgtatcgtagttatctacacgacggggagtcaggcaactatggatgaacgaaatagacag
atcgctgagataggtgcctcactgattaagcattggtaactgtcagaccaagtttactcatatatactttagattgatttaaaacttcattttta
atttaaaaggatctaggtgaagatcctttttgataatctcatgaccaaaatcccttaacgtgagttttcgttccactgagcgtcagaccccgt
agaaaagatcaaaggatcttcttgagatcctttttttctgcgcgtaatctgctgcttgcaaacaaaaaaaccaccgctaccagcggtggttt
gtttgccggatcaagagctaccaactctttttccgaaggtaactggcttcagcagagcgcagataccaaatactgttcttctagtgtagcc
gtagttaggccaccacttcaagaactctgtagcaccgcctacatacctcgctctgctaatcctgttaccagtggctgctgccagtggcga
taagtcgtgtcttaccgggttggactcaagacgatagttaccggataaggcgcagcggtcgggctgaacggggggttcgtgcacaca
gcccagcttggagcgaacgacctacaccgaactgagatacctacagcgtgagctatgagaaagcgccacgcttcccgaagggaga
aaggcggacaggtatccggtaagcggcagggtcggaacaggagagcgcacgagggagcttccagggggaaacgcctggtatctt
tatagtcctgtcgggtttcgccacctctgacttgagcgtcgatttttgtgatgctcgtcaggggggcggagcctatggaaaaacgccagc
aacgcggc

**pBZ210-pU6-sgBFP-An_CBh-sv40NLS-Cas9-Ec86RT-NLS-T2A-mCherry**

catgtgagggcctatttcccatgattccttcatatttgcatatacgatacaaggctgttagagagataattggaattaatttgactgtaaacac
aaagatattagtacaaaatacgtgacgtagaaagtaataatttcttgggtagtttgcagttttaaaattatgttttaaaatggactatcatatgc
ttaccgtaacttgaaagtatttcgatttcttggctttatatatcttgtggaaaggacgaaacaccGCTGAAGCACTGCACGC
CATgttttagagctagaaatagcaagttaaaataaggctagtccgttatcaacttgaaaaagtggcaccgagtcggtgcCGTAC
GATGCGCACCCTTAGCGAGAGGTTTATCATTAAGGTCAACCTCTGGATGTTGTTT
CGGCATCCTGCATTGAATCTGAGTTACTGTCTGTTTcCCTACTAGTAAGTCGTGCT
GCTTCATGTGGTCGGGGTAGCGGCTGAAGCACTGCACGCCGTACGTCAGGGTGG
TCACGAGGGTGGGCCAGGGCACGGGCAGCTTGCCGGTGGTGCAGATGAACTTCA
GGcctaggAGGGAACCCGTTTCTTCTGACGTAAGGGTGCGCAtttttttctagaggtacccgttacata
acttacggtaaatggcccgcctggctgaccgcccaacgacccccgcccattgacgtcaatagtaacgccaatagggactttccattga
cgtcaatgggtggagtatttacggtaaactgcccacttggcagtacatcaagtgtatcatatgccaagtacgccccctattgacgtcaatg
acggtaaatggcccgcctggcattgtgcccagtacatgaccttatgggactttcctacttggcagtacatctacgtattagtcatcgctatt
accatggtcgaggtgagccccacgttctgcttcactctccccatctcccccccctccccacccccaattttgtatttatttattttttaattattt
tgtgcagcgatgggggcgggggggggggggggcgcgcgccaggcggggcggggcggggcgaggggcggggcggggcg
aggcggagaggtgcggcggcagccaatcagagcggcgcgctccgaaagtttccttttatggcgaggcggcggcggcggcggccc
tataaaaagcgaagcgcgcggcgggcgggagtcgctgcgacgctgccttcgccccgtgccccgctccgccgccgcctcgcgccg
cccgccccggctctgactgaccgcgttactcccacaggtgagcgggcgggacggcccttctcctccgggctgtaattagctgagcaa
gaggtaagggtttaagggatggttggttggtggggtattaatgtttaattacctggagcacctgcctgaaatcactttttttcaggttggacc
ggtgccaccatggactataaggaccacgacggagactacaaggatcatgatattgattacaaagacgatgacgataagatggcccca
aagaagaagcggaaggtcggtatccacggagtcccagcagccgacaagaagtacagcatcggcctggacatcggcaccaactctg
tgggctgggccgtgatcaccgacgagtacaaggtgcccagcaagaaattcaaggtgctgggcaacaccgaccggcacagcatcaa
gaagaacctgatcggagccctgctgttcgacagcggcgaaacagccgaggccacccggctgaagagaaccgccagaagaagata
caccagacggaagaaccggatctgctatctgcaagagatcttcagcaacgagatggccaaggtggacgacagcttcttccacagact
ggaagagtccttcctggtggaagaggataagaagcacgagcggcaccccatcttcggcaacatcgtggacgaggtggcctaccacg
agaagtaccccaccatctaccacctgagaaagaaactggtggacagcaccgacaaggccgacctgcggctgatctatctggccctg
gcccacatgatcaagttccggggccacttcctgatcgagggcgacctgaaccccgacaacagcgacgtggacaagctgttcatccag
ctggtgcagacctacaaccagctgttcgaggaaaaccccatcaacgccagcggcgtggacgccaaggccatcctgtctgccagact
gagcaagagcagacggctggaaaatctgatcgcccagctgcccggcgagaagaagaatggcctgttcggaaacctgattgccctga

gcctgggcctgacccccaacttcaagagcaacttcgacctggccgaggatgccaaactgcagctgagcaaggacacctacgacgac
gacctggacaacctgctggcccagatcggcgaccagtacgccgacctgtttctggccgccaagaacctgtccgacgccatcctgctg
agcgacatcctgagagtgaacaccgagatcaccaaggcccccctgagcgcctctatgatcaagagatacgacgagcaccaccagg
acctgaccctgctgaaagctctcgtgcggcagcagctgcctgagaagtacaaagagattttcttcgaccagagcaagaacggctacg
ccggctacattgacggcggagccagccaggaagagttctacaagttcatcaagcccatcctggaaaagatggacggcaccgaggaa
ctgctcgtgaagctgaacagagaggacctgctgcggaagcagcggaccttcgacaacggcagcatccccccaccagatccacctgg
gagagctgcacgccattctgcggcggcaggaagatttttacccattcctgaaggacaaccgggaaaagatcgagaagatcctgacctt
ccgcatcccctactacgtgggccctctggccaggggaaacagcagattcgcctggatgaccagaaagagcgaggaaaccatcacc
ccctggaacttcgaggaagtggtggacaagggcgcttccgcccagagcttcatcgagcggatgaccaacttcgataagaacctgccc
aacgagaaggtgctgcccaagcacagcctgctgtacgagtacttcaccgtgtataacgagctgaccaaagtgaaatacgtgaccgag
ggaatgagaaagcccgccttcctgagcggcgagcagaaaaaggccatcgtggacctgctgttcaagaccaaccggaaagtgaccg
tgaagcagctgaaagaggactacttcaagaaaatcgagtgcttcgactccgtggaaatctccggcgtggaagatcggttcaacgcctc
cctgggcacataccacgatctgctgaaaattatcaaggacaaggacttcctggacaatgaggaaaacgaggacattctggaagatatc
gtgctgaccctgacactgtttgaggacagagagatgatcgaggaacggctgaaaacctatgcccacctgttcgacgacaaagtgatg
aagcagctgaagcggcggagatacaccggctggggcaggctgagccggaagctgatcaacggcatccgggacaagcagtccgg
caagacaatcctggatttcctgaagtccgacggcttcgccaacagaaacttcatgcagctgatccacgacgacagcctgacctttaaag
aggacatccagaaagcccaggtgtccggccagggcgatagcctgcacgagcacattgccaatctggccggcagccccgccattaa
gaagggcatcctgcagacagtgaaggtggtggacgagctcgtgaaagtgatgggccggcacaagcccgagaacatcgtgatcgaa
atggccagagagaaccagaccacccagaagggacagaagaacagccgcgagagaatgaagcggatcgaagagggcatcaaag
agctgggcagccagatcctgaaagaacaccccgtggaaaacacccagctgcagaacgagaagctgtacctgtactacctgcagaat
gggcgggatatgtacgtggaccaggaactggacatcaaccggctgtccgactacgatgtggaccatatcgtgcctcagagctttctga
aggacgactccatcgacaacaaggtgctgaccagaagcgacaagaaccggggcaagagcgacaacgtgccctccgaagaggtcg
tgaagaagatgaagaactactggcggcagctgctgaacgccaagctgattacccagagaaagttcgacaatctgaccaaggccgag
agaggcggcctgagcgaactggataaggccggcttcatcaagagacagctggtggaaacccggcagatcacaaagcacgtggca
cagatcctggactcccggatgaacactaagtacgacgagaatgacaagctgatccgggaagtgaaagtgatcaccctgaagtccaag
ctggtgtccgatttccggaaggatttccagttttacaaagtgcgcgagatcaacaactaccaccacgcccacgacgcctacctgaacgc
cgtcgtgggaaccgccctgatcaaaaagtaccctaagctggaaagcgagttcgtgtacggcgactacaaggtgtacgacgtgcgga
agatgatcgccaagagcgagcaggaaatcggcaaggctaccgccaagtacttcttctacagcaacatcatgaactttttcaagaccga
gattaccctggccaacggcgagatccggaagcggcctctgatcgagacaaacggcgaaaccggggagatcgtgtgggataaggg
ccgggattttgccaccgtgcggaaagtgctgagcatgccccaagtgaatatcgtgaaaaagaccgaggtgcagacaggcggcttca
gcaaagagtctatcctgcccaagaggaacagcgataagctgatcgccagaaagaaggactgggaccctaagaagtacggcggcttc
gacagccccaccgtggcctattctgtgctggtggtggccaaagtggaaaagggcaagtccaagaaactgaagagtgtgaaagagct
gctggggatcaccatcatggaaagaagcagcttcgagaagaatcccatcgactttctggaagccaagggctacaaagaagtgaaaaa
ggacctgatcatcaagctgcctaagtactccctgttcgagctggaaaacggccggaagagaatgctggcctctgccggcgaactgca
gaagggaaacgaactggccctgccctccaaatatgtgaacttcctgtacctggccagccactatgagaagctgaagggctcccccga
ggataatgagcagaaacagctgtttgtggaacagcacaagcactacctggacgagatcatcgagcagatcagcgagttctccaagag
agtgatcctggccgacgctaatctggacaaagtgctgtccgcctacaacaagcaccgggataagcccatcagagagcaggccgaga
atatcatccacctgtttaccctgaccaatctgggagcccctgccgccttcaagtactttgacaccaccatcgaccggaagaggtacacc
agcaccaaagaggtgctggacgccaccctgatccaccagagcatcaccggcctgtacgagacacggatcgacctgtctcagctggg
aggcgactctggaggatctagcggaggatcctctggcagcgagacaccaggaacaagcgagtcagcaacaccagagagcagtgg
cggcagcagcggcggcagcagcAAGAGTGCGGAATACCTTAACACATTTAGGCTTAGAAAC
CTGGGACTCCCCGTTATGAATAACCTGCACGATATGTCTAAGGCAACCCGAATCA
GTGTAGAGACGCTTAGACTCTTGATATATACCGCCGACTTTCGATACAGAATATA
TACCGTCGAGAAGAAGGGCCCAGAGAAACGAATGCGGACCATATACCAACCTAG
TAGGGAGCTGAAAGCGCTTCAGGGCTGGGTACTTCGAAATATCCTTGACAAACTT
TCCAGTTCACCGTTCAGCATTGGTTTCGAGAAACACCAGAGTATTCTGAATAACG
CGACACCTCACATAGGAGCCAATTTCATCCTCAATATTGACCTGGAGGATTTCTT
CCCTAGCCTTACTGCCAATAAAGTGTTCGGTGTATTCCACAGCCTCGGCTACAAC
CGACTGATTTCATCTGTACTCACAAAAATTTGTTGCTATAAGAACCTGCTCCCTC
AAGGGGCCCCAAGTAGCCCAAAACTGGCAAACCTCATCTGTTCAAAATTGGACT

ATCGAATTCAGGGCTATGCCGGATCCAGGGGCTTGATCTACACGAGATACGCAG
ACGACCTGACACTTTCAGCACAATCCATGAAGAAAGTTGTTAAAGCGAGGGATT
TTCTTTTTCCATTATTCCGTCTGAAGGATTGGTTATTAATTCTAAAAAAACTTGC
ATTAGTGGTCCTCGGTCTCAACGAAAGGTTACAGGCCTGGTAATCTCTCAGGAGA
AGGTTGGGATTGGAAGAGAGAAGTATAAAGAGATTCGCGCCAAAATACATCATA
TTTTTTGCGGTAAATCATCTGAAATCGAGCACGTAAGGGGATGGCTTTCTTTCATT
CTTTCTGTAGACTCCAAATCTCACCGACGACTCATCACATATATAAGTAAACTGG
AGAAAAAATATGGAAAAATCCGCTTAACAAGGCTAAAACTaaaaggccggcggccacga
aaaaggccggccaggcaaaaaagaaaaagcttgagggcagaggaagtctgctaacatgcggtgacgtggaggagaatcccggcc
ctgctagcatggtgagcaagggcgaggaggataacatggccatcatcaaggagttcatgcgcttcaaggtgcacatggagggctcc
gtgaacggccacgagttcgagatcgagggcgagggcgagggccgcccctacgagggcacccagaccgccaagctgaaggtgac
caagggtggcccccctgcccttcgcctgggacatcctgtcccctcagttcatgtacggctccaaggcctacgtgaagcaccccgccga
catccccgactacttgaagctgtccttccccgagggcttcaagtgggagcgcgtgatgaacttcgaggacggcggcgtggtgaccgt
gacccaggactcctccctgcaggacggcgagttcatctacaaggtgaagctgcgcggcaccaacttcccctccgacggccccgtaat
gcagaagaagaccatgggctgggaggcctcctccgagcggatgtaccccgaggacggcgccctgaagggcgagatcaagcaga
ggctgaagctgaaggacggcggccactacgacgctgaggtcaagaccacctacaaggccaagaagcccgtgcagctgcccggcg
cctacaacgtcaacatcaagttggacatcacctcccacaacgaggactacaccatcgtggaacagtacgaacgcgccgagggccgc
cactccaccggcggcatggacgagctgtacaagtgagaattcctagagctcgctgatcagcctcgactgtgccttctagttgccagcc
atctgttgtttgcccctcccccgtgccttccttgaccctggaaggtgccactcccactgtcctttcctaataaaatgaggaaattgcatcgc
attgtctgagtaggtgtcattctattctggggggtggggtggggcaggacagcaagggggaggattgggaagagaatagcaggcat
gctggggagcggccgcaggaacccctagtgatggagttggccactccctctctgcgcgctcgctcgctcactgaggccgggcgacc
aaaggtcgcccgacgcccgggctttgcccgggcggcctcagtgagcgagcgagcgcgcagctgcctgcaggggcgcctgatgcg
gtattttctccttacgcatctgtgcggtatttcacaccgcatacgtcaaagcaaccatagtacgcgccctgtagcggcgcattaagcgcg
gcgggtgtggtggttacgcgcagcgtgaccgctacacttgccagcgccctagcgcccgctcctttcgctttcttcccttcctttctcgcca
cgttcgccggctttccccgtcaagctctaaatcgggggctccctttagggttccgatttagtgctttacggcacctcgaccccaaaaaact
tgatttgggtgatggttcacgtagtgggccatcgccctgatagacggtttttcgccctttgacgttggagtccacgttctttaatagtggact
cttgttccaaactggaacaacactcaaccctatctcgggctattcttttgatttataagggattttgccgatttcggCctattggttaaaaaat
gagctgatttaacaaaaatttaacgcgaattttaacaaaatattaacgtttacaattttatggtgcactctcagtacaatctgctctgatgccg
catagttaagccagccccgacacccgccaacacccgctgacgcgccctgacgggcttgtctgctcccggcatccgcttacagacaag
ctgtgaccgtctccgggagctgcatgtgtcagaggttttcaccgtcatcaccgaaacgcgcgagacgaaagggcctcgtgatacgcct
atttttataggttaatgtcatgataataatggtttcttagacgtcaggtggcacttttcggggaaatgtgcgcggaacccctatttgtttattttt
ctaaatacattcaaatatgtatccgctcatgagacaataaccctgataaatgcttcaataatattgaaaaaggaagagtatgagtattcaac
atttccgtgtcgcccttattcccttttttgcggcattttgccttcctgtttttgctcacccagaaacgctggtgaaagtaaaagatgctgaagat
cagttgggtgcacgagtgggttacatcgaactggatctcaacagcggtaagatccttgagagttttcgccccgaagaacgttttccaatg
atgagcacttttaaagttctgctatgtggcgcggtattatcccgtattgacgccgggcaagagcaactcggtcgccgcatacactattctc
agaatgacttggttgagtactcaccagtcacagaaaagcatcttacggatggcatgacagtaagagaattatgcagtgctgccataacc
atgagtgataacactgcggccaacttacttctgacaacgatcggaggaccgaaggagctaaccgcttttttgcacaacatgggggatc
atgtaactcgccttgatcgttgggaaccggagctgaatgaagccataccaaacgacgagcgtgacaccacgatgcctgtagcaatgg
caacaacgttgcgcaaactattaactggcgaactacttactctagcttcccggcaacaattaatagactggatggaggcggataaagttg
caggaccacttctgcgctcggcccttccggctggctggtttattgctgataaatctggagccggtgagcgtggaagccgcggtatcatt
gcagcactggggccagatggtaagccctcccgtatcgtagttatctacacgacggggagtcaggcaactatggatgaacgaaataga
cagatcgctgagataggtgcctcactgattaagcattggtaactgtcagaccaagtttactcatatatactttagattgatttaaaacttcattt
ttaatttaaaaggatctaggtgaagatcctttttgataatctcatgaccaaaatcccttaacgtgagttttcgttccactgagcgtcagaccc
cgtagaaaagatcaaaggatcttcttgagatcctttttttctgcgcgtaatctgctgcttgcaaacaaaaaaaccaccgctaccagcggtg
gtttgtttgccggatcaagagctaccaactctttttccgaaggtaactggcttcagcagagcgcagataccaaatactgttcttctagtgta
gccgtagttaggccaccacttcaagaactctgtagcaccgcctacatacctcgctctgctaatcctgttaccagtggctgctgccagtgg
cgataagtcgtgtcttaccgggttggactcaagacgatagttaccggataaggcgcagcggtcgggctgaacggggggttcgtgcac
acagcccagcttggagcgaacgacctacaccgaactgagatacctacagcgtgagctatgagaaagcgccacgcttcccgaaggg
agaaaggcggacaggtatccggtaagcggcagggtcggaacaggagagcgcacgagggagcttccagggggaaacgcctggta

tctttatagtcctgtcgggtttcgccacctctgacttgagcgtcgatttttgtgatgctcgtcaggggggcggagcctatggaaaaacgcc
agcaacgcggc

**pBZ211-pU6-sgBFP-Dt_CBh-sv40NLS-Cas9-Ec86RT-NLS-T2A-mCherry**

catgtgagggcctatttcccatgattccttcatatttgcatatacgatacaaggctgttagagagataattggaattaatttgactgtaaacac
aaagatattagtacaaaatacgtgacgtagaaagtaataatttcttgggtagtttgcagttttaaaattatgttttaaaatggactatcatatgc
ttaccgtaacttgaaagtatttcgatttcttggctttatatatcttgtggaaaggacgaaacaccGCTGAAGCACTGCACGC
CATgttttagagctagaaatagcaagttaaaataaggctagtccgttatcaacttgaaaaagtggcaccgagtcggtgcCGTAC
GATGCGCACCCTTAGCGAGAGGTTTATCATTAAGGTCAACCTCTGGATGTTGTTT
CGGCATCCTGCATTGAATCTGAGTTACTGTCTGTTTcCCTACTAGTCCCTCGTGAC
CACCCTGACGTACGGCGTGCAGTGCTTCAGCCGCTACCCCGACCACATGAAGCA
GCACGACTTCTTCAAGTCCGCCATGCCCGAAGGCTACGTCCAGGAGCGCcctaggAG
GGAACCCGTTTCTTCTGACGTAAGGGTGCGCAtttttttctagaggtacccgttacataacttacggtaaat
ggcccgcctggctgaccgcccaacgacccccgcccattgacgtcaatagtaacgccaatagggactttccattgacgtcaatgggtg
gagtatttacggtaaactgcccacttggcagtacatcaagtgtatcatatgccaagtacgccccctattgacgtcaatgacggtaaatgg
cccgcctggcattgtgcccagtacatgaccttatgggactttcctacttggcagtacatctacgtattagtcatcgctattaccatggtcga
ggtgagccccacgttctgcttcactctccccatctcccccccctccccacccccaattttgtatttatttattttttaattattttgtgcagcgat
gggggcgggggggggggggggcgcgcgccaggcggggcggggcggggcgaggggcggggcggggcgaggcggagag
gtgcggcggcagccaatcagagcggcgcgctccgaaagtttccttttatggcgaggcggcggcggcggcggccctataaaaagcg
aagcgcgcggcgggcgggagtcgctgcgacgctgccttcgccccgtgccccgctccgccgccgcctcgcgccgcccgccccgg
ctctgactgaccgcgttactcccacaggtgagcgggcgggacggcccttctcctccgggctgtaattagctgagcaagaggtaaggg
tttaagggatggttggttggtggggtattaatgtttaattacctggagcacctgcctgaaatcactttttttcaggttggaccggtgccacca
tggactataaggaccacgacggagactacaaggatcatgatattgattacaaagacgatgacgataagatggccccaaagaagaagc
ggaaggtcggtatccacggagtcccagcagccgacaagaagtacagcatcggcctggacatcggcaccaactctgtgggctgggc
cgtgatcaccgacgagtacaaggtgcccagcaagaaattcaaggtgctgggcaacaccgaccggcacagcatcaagaagaacctg
atcggagccctgctgttcgacagcggcgaaacagccgaggccacccggctgaagagaaccgccagaagaagatacaccagacgg
aagaaccggatctgctatctgcaagagatcttcagcaacgagatggccaaggtggacgacagcttcttccacagactggaagagtcct
tcctggtggaagaggataagaagcacgagcggcaccccatcttcggcaacatcgtggacgaggtggcctaccacgagaagtaccc
caccatctaccacctgagaaagaaactggtggacagcaccgacaaggccgacctgcggctgatctatctggccctggcccacatgat
caagttccggggccacttcctgatcgagggcgacctgaaccccgacaacagcgacgtggacaagctgttcatccagctggtgcaga
cctacaaccagctgttcgaggaaaaccccatcaacgccagcggcgtggacgccaaggccatcctgtctgccagactgagcaagagc
agacggctggaaaatctgatcgcccagctgcccggcgagaagaagaatggcctgttcggaaacctgattgccctgagcctgggcct
gacccccaacttcaagagcaacttcgacctggccgaggatgccaaactgcagctgagcaaggacacctacgacgacgacctggac
aacctgctggcccagatcggcgaccagtacgccgacctgtttctggccgccaagaacctgtccgacgccatcctgctgagcgacatc
ctgagagtgaacaccgagatcaccaaggcccccctgagcgcctctatgatcaagagatacgacgagcaccaccaggacctgaccct
gctgaaagctctcgtgcggcagcagctgcctgagaagtacaaagagattttcttcgaccagagcaagaacggctacgccggctacatt
gacggcggagccagccaggaagagttctacaagttcatcaagcccatcctggaaaagatggacggcaccgaggaactgctcgtga
agctgaacagagaggacctgctgcggaagcagcggaccttcgacaacggcagcatcccccaccagatccacctgggagagctgc
acgccattctgcggcggcaggaagatttttacccattcctgaaggacaaccgggaaaagatcgagaagatcctgaccttccgcatccc
ctactacgtgggccctctggccaggggaaacagcagattcgcctggatgaccagaaagagcgaggaaaccatcaccccctggaact
tcgaggaagtggtggacaagggcgcttccgcccagagcttcatcgagcggatgaccaacttcgataagaacctgcccaacgagaag
gtgctgcccaagcacagcctgctgtacgagtacttcaccgtgtataacgagctgaccaaagtgaaatacgtgaccgagggaatgaga
aagcccgccttcctgagcggcgagcagaaaaaggccatcgtggacctgctgttcaagaccaaccggaaagtgaccgtgaagcagct
gaaagaggactacttcaagaaaatcgagtgcttcgactccgtggaaatctccggcgtggaagatcggttcaacgcctccctgggcaca
taccacgatctgctgaaaattatcaaggacaaggacttcctggacaatgaggaaaacgaggacattctggaagatatcgtgctgaccct
gacactgtttgaggacagagagatgatcgaggaacggctgaaaacctatgcccacctgttcgacgacaaagtgatgaagcagctgaa
gcggcggagatacaccggctggggcaggctgagccggaagctgatcaacggcatccgggacaagcagtccggcaagacaatcct
ggatttcctgaagtccgacggcttcgccaacagaaacttcatgcagctgatccacgacgacagcctgacctttaaagaggacatccag

aaagcccaggtgtccggccagggcgatagcctgcacgagcacattgccaatctggccggcagccccgccattaagaagggcatcct
gcagacagtgaaggtggtggacgagctcgtgaaagtgatgggccggcacaagcccgagaacatcgtgatcgaaatggccagaga
gaaccagaccacccagaagggacagaagaacagccgcgagagaatgaagcggatcgaagagggcatcaaagagctgggcagc
cagatcctgaaagaacaccccgtggaaaacacccagctgcagaacgagaagctgtacctgtactacctgcagaatgggcgggatat
gtacgtggaccaggaactggacatcaaccggctgtccgactacgatgtggaccatatcgtgcctcagagctttctgaaggacgactcc
atcgacaacaaggtgctgaccagaagcgacaagaaccggggcaagagcgacaacgtgccctccgaagaggtcgtgaagaagatg
aagaactactggcggcagctgctgaacgccaagctgattacccagagaaagttcgacaatctgaccaaggccgagagaggcggcct
gagcgaactggataaggccggcttcatcaagagacagctggtggaaacccggcagatcacaaagcacgtggcacagatcctggac
tcccggatgaacactaagtacgacgagaatgacaagctgatccgggaagtgaaagtgatcaccctgaagtccaagctggtgtccgatt
tccggaaggatttccagttttacaaagtgcgcgagatcaacaactaccaccacgcccacgacgcctacctgaacgccgtcgtgggaa
ccgccctgatcaaaaagtaccctaagctggaaagcgagttcgtgtacggcgactacaaggtgtacgacgtgcggaagatgatcgcca
agagcgagcaggaaatcggcaaggctaccgccaagtacttcttctacagcaacatcatgaactttttcaagaccgagattaccctggcc
aacggcgagatccggaagcggcctctgatcgagacaaacggcgaaaccggggagatcgtgtgggataagggccgggattttgcca
ccgtgcggaaagtgctgagcatgccccaagtgaatatcgtgaaaaagaccgaggtgcagacaggcggcttcagcaaagagtctatc
ctgcccaagaggaacagcgataagctgatcgccagaaagaaggactgggaccctaagaagtacggcggcttcgacagccccacc
gtggcctattctgtgctggtggtggccaaagtggaaaagggcaagtccaagaaactgaagagtgtgaaagagctgctggggatcacc
atcatggaaagaagcagcttcgagaagaatcccatcgactttctggaagccaagggctacaaagaagtgaaaaaggacctgatcatc
aagctgcctaagtactccctgttcgagctggaaaacggccggaagagaatgctggcctctgccggcgaactgcagaagggaaacga
actggccctgccctccaaatatgtgaacttcctgtacctggccagccactatgagaagctgaagggctcccccgaggataatgagcag
aaacagctgtttgtggaacagcacaagcactacctggacgagatcatcgagcagatcagcgagttctccaagagagtgatcctggcc
gacgctaatctggacaaagtgctgtccgcctacaacaagcaccgggataagcccatcagagagcaggccgagaatatcatccacct
gtttaccctgaccaatctgggagcccctgccgccttcaagtactttgacaccaccatcgaccggaagaggtacaccagcaccaaaga
ggtgctggacgccaccctgatccaccagagcatcaccggcctgtacgagacacggatcgacctgtctcagctgggaggcgactctg
gaggatctagcggaggatcctctggcagcgagacaccaggaacaagcgagtcagcaacaccagagagcagtggcggcagcagc
ggcggcagcagcAAGAGTGCGGAATACCTTAACACATTTAGGCTTAGAAACCTGGGAC
TCCCCGTTATGAATAACCTGCACGATATGTCTAAGGCAACCCGAATCAGTGTAGA
GACGCTTAGACTCTTGATATATACCGCCGACTTTCGATACAGAATATATACCGTC
GAGAAGAAGGGCCCAGAGAAACGAATGCGGACCATATACCAACCTAGTAGGGA
GCTGAAAGCGCTTCAGGGCTGGGTACTTCGAAATATCCTTGACAAACTTTCCAGT
TCACCGTTCAGCATTGGTTTCGAGAAACACCAGAGTATTCTGAATAACGCGACAC
CTCACATAGGAGCCAATTTCATCCTCAATATTGACCTGGAGGATTTCTTCCCTAG
CCTTACTGCCAATAAAGTGTTCGGTGTATTCCACAGCCTCGGCTACAACCGACTG
ATTTCATCTGTACTCACAAAAATTTGTTGCTATAAGAACCTGCTCCCTCAAGGGG
CCCCAAGTAGCCCAAAACTGGCAAACCTCATCTGTTCAAAATTGGACTATCGAAT
TCAGGGCTATGCCGGATCCAGGGGCTTGATCTACACGAGATACGCAGACGACCT
GACACTTTCAGCACAATCCATGAAGAAAGTTGTTAAAGCGAGGGATTTTCTTTTT
TCCATTATTCCGTCTGAAGGATTGGTTATTAATTCTAAAAAAACTTGCATTAGTG
GTCCTCGGTCTCAACGAAAGGTTACAGGCCTGGTAATCTCTCAGGAGAAGGTTG
GGATTGGAAGAGAGAAGTATAAAGAGATTCGCGCCAAAATACATCATATTTTTT
GCGGTAAATCATCTGAAATCGAGCACGTAAGGGGATGGCTTTCTTTCATTCTTTC
TGTAGACTCCAAATCTCACCGACGACTCATCACATATATAAGTAAACTGGAGAA
AAAATATGGAAAAATCCGCTTAACAAGGCTAAAACTaaaaggccggcggccacgaaaaagg
ccggccaggcaaaaaagaaaaagcttgagggcagaggaagtctgctaacatgcggtgacgtggaggagaatcccggccctgctag
catggtgagcaagggcgaggaggataacatggccatcatcaaggagttcatgcgcttcaaggtgcacatggagggctccgtgaacg
gccacgagttcgagatcgagggcgagggcgagggccgcccctacgagggcacccagaccgccaagctgaaggtgaccaagggt
ggcccccctgcccttcgcctgggacatcctgtcccctcagttcatgtacggctccaaggcctacgtgaagcaccccgccgacatccccg
actacttgaagctgtccttccccgagggcttcaagtgggagcgcgtgatgaacttcgaggacggcggcgtggtgaccgtgacccagg
actccctcctgcaggacggcgagttcatctacaaggtgaagctgcgcggcaccaacttcccctccgacggccccgtaatgcagaaga
agaccatgggctgggaggcctcctccgagcggatgtaccccgaggacggcgccctgaagggcgagatcaagcagaggctgaagc
tgaaggacggcggccactacgacgctgaggtcaagaccacctacaaggccaagaagcccgtgcagctgcccggcgcctacaacg

tcaacatcaagttggacatcacctcccacaacgaggactacaccatcgtggaacagtacgaacgcgccgagggccgccactccacc
ggcggcatggacgagctgtacaagtgagaattcctagagctcgctgatcagcctcgactgtgccttctagttgccagccatctgttgttt
gcccctcccccgtgccttccttgaccctggaaggtgccactcccactgtcctttcctaataaaatgaggaaattgcatcgcattgtctgag
taggtgtcattctattctggggggtggggtggggcaggacagcaagggggaggattgggaagagaatagcaggcatgctggggag
cggccgcaggaacccctagtgatggagttggccactccctctctgcgcgctcgctcgctcactgaggccgggcgaccaaaggtcgc
ccgacgcccgggctttgcccgggcggcctcagtgagcgagcgagcgcgcagctgcctgcaggggcgcctgatgcggtattttctcc
ttacgcatctgtgcggtatttcacaccgcatacgtcaaagcaaccatagtacgcgccctgtagcggcgcattaagcgcggcgggtgtg
gtggttacgcgcagcgtgaccgctacacttgccagcgccctagcgcccgctcctttcgctttcttcccttcctttctcgccacgttcgccg
gctttccccgtcaagctctaaatcgggggctccctttagggttccgatttagtgctttacggcacctcgaccccaaaaaacttgatttgggt
gatggttcacgtagtgggccatcgccctgatagacggtttttcgccctttgacgttggagtccacgttctttaatagtggactcttgttccaa
actggaacaacactcaaccctatctcgggctattcttttgatttataagggattttgccgatttcggCctattggttaaaaaatgagctgattt
aacaaaaatttaacgcgaattttaacaaaatattaacgtttacaattttatggtgcactctcagtacaatctgctctgatgccgcatagttaag
ccagccccgacacccgccaacacccgctgacgcgccctgacgggcttgtctgctcccggcatccgcttacagacaagctgtgaccg
tctccgggagctgcatgtgtcagaggttttcaccgtcatcaccgaaacgcgcgagacgaaagggcctcgtgatacgcctatttttatag
gttaatgtcatgataataatggtttcttagacgtcaggtggcacttttcggggaaatgtgcgcggaacccctatttgtttatttttctaaataca
ttcaaatatgtatccgctcatgagacaataaccctgataaatgcttcaataatattgaaaaaggaagagtatgagtattcaacatttccgtgt
cgcccttattcccttttttgcggcattttgccttcctgtttttgctcacccagaaacgctggtgaaagtaaaagatgctgaagatcagttgggt
gcacgagtgggttacatcgaactggatctcaacagcggtaagatccttgagagttttcgccccgaagaacgttttccaatgatgagcact
tttaaagttctgctatgtggcgcggtattatcccgtattgacgccgggcaagagcaactcggtcgccgcatacactattctcagaatgact
tggttgagtactcaccagtcacagaaaagcatcttacggatggcatgacagtaagagaattatgcagtgctgccataaccatgagtgat
aacactgcggccaacttacttctgacaacgatcggaggaccgaaggagctaaccgcttttttgcacaacatgggggatcatgtaactcg
ccttgatcgttgggaaccggagctgaatgaagccataccaaacgacgagcgtgacaccacgatgcctgtagcaatggcaacaacgtt
gcgcaaactattaactggcgaactacttactctagcttcccggcaacaattaatagactggatggaggcggataaagttgcaggaccac
ttctgcgctcggcccttccggctggctggtttattgctgataaatctggagccggtgagcgtggaagccgcggtatcattgcagcactg
gggccagatggtaagccctcccgtatcgtagttatctacacgacggggagtcaggcaactatggatgaacgaaatagacagatcgct
gagataggtgcctcactgattaagcattggtaactgtcagaccaagtttactcatatatactttagattgatttaaaacttcatttttaatttaaa
aggatctaggtgaagatcctttttgataatctcatgaccaaaatcccttaacgtgagttttcgttccactgagcgtcagaccccgtagaaaa
gatcaaaggatcttcttgagatcctttttttctgcgcgtaatctgctgcttgcaaacaaaaaaaccaccgctaccagcggtggtttgtttgcc
ggatcaagagctaccaactctttttccgaaggtaactggcttcagcagagcgcagataccaaatactgttcttctagtgtagccgtagtta
ggccaccacttcaagaactctgtagcaccgcctacatacctcgctctgctaatcctgttaccagtggctgctgccagtggcgataagtcg
tgtcttaccgggttggactcaagacgatagttaccggataaggcgcagcggtcgggctgaacggggggttcgtgcacacagcccag
cttggagcgaacgacctacaccgaactgagatacctacagcgtgagctatgagaaagcgccacgcttcccgaagggagaaaggcg
gacaggtatccggtaagcggcagggtcggaacaggagagcgcacgagggagcttccagggggaaacgcctggtatctttatagtc
ctgtcgggtttcgccacctctgacttgagcgtcgatttttgtgatgctcgtcaggggggcggagcctatggaaaaacgccagcaacgc
ggc

**pBZ212-pU6-sgBFP-Dn_CBh-sv40NLS-Cas9-Ec86RT-NLS-T2A-mCherry**

catgtgagggcctatttcccatgattccttcatatttgcatatacgatacaaggctgttagagagataattggaattaatttgactgtaaacac
aaagatattagtacaaaatacgtgacgtagaaagtaataatttcttgggtagtttgcagttttaaaattatgttttaaaatggactatcatatgc
ttaccgtaacttgaaagtatttcgatttcttggctttatatatcttgtggaaaggacgaaacaccGCTGAAGCACTGCACGC
CATgttttagagctagaaatagcaagttaaaataaggctagtccgttatcaacttgaaaaagtggcaccgagtcggtgcCGTAC
GATGCGCACCCTTAGCGAGAGGTTTATCATTAAGGTCAACCTCTGGATGTTGTTT
CGGCATCCTGCATTGAATCTGAGTTACTGTCTGTTTcCCTACTAGTGCGCTCCTGG
ACGTAGCCTTCGGGCATGGCGGACTTGAAGAAGTCGTGCTGCTTCATGTGGTCGG
GGTAGCGGCTGAAGCACTGCACGCCGTACGTCAGGGTGGTCACGAGGGcctaggAG
GGAACCCGTTTCTTCTGACGTAAGGGTGCGCAtttttttctagaggtacccgttacataacttacggtaaat
ggcccgcctggctgaccgcccaacgacccccgcccattgacgtcaatagtaacgccaatagggactttccattgacgtcaatgggtg
gagtatttacggtaaactgcccacttggcagtacatcaagtgtatcatatgccaagtacgccccctattgacgtcaatgacggtaaatgg

cccgcctggcattgtgcccagtacatgaccttatgggactttcctacttggcagtacatctacgtattagtcatcgctattaccatggtcga
ggtgagccccacgttctgcttcactctccccatctcccccccctccccacccccaattttgtatttatttattttttaattattttgtgcagcgat
gggggcgggggggggggggggcgcgcgccaggcggggcggggcggggcgaggggcggggcggggcgaggcggagag
gtgcggcggcagccaatcagagcggcgcgctccgaaagtttccttttatggcgaggcggcggcggcggcggccctataaaaagcg
aagcgcgcggcgggcgggagtcgctgcgacgctgccttcgccccgtgccccgctccgccgccgcctcgcgccgcccgccccgg
ctctgactgaccgcgttactcccacaggtgagcgggcgggacggcccttctcctccgggctgtaattagctgagcaagaggtaaggg
tttaagggatggttggttggtggggtattaatgtttaattacctggagcacctgcctgaaatcactttttttcaggttggaccggtgccacca
tggactataaggaccacgacggagactacaaggatcatgatattgattacaaagacgatgacgataagatggccccaaagaagaagc
ggaaggtcggtatccacggagtcccagcagccgacaagaagtacagcatcggcctggacatcggcaccaactctgtgggctgggc
cgtgatcaccgacgagtacaaggtgcccagcaagaaattcaaggtgctgggcaacaccgaccggcacagcatcaagaagaacctg
atcggagccctgctgttcgacagcggcgaaacagccgaggccacccggctgaagagaaccgccagaagaagatacaccagacgg
aagaaccggatctgctatctgcaagagatcttcagcaacgagatggccaaggtggacgacagcttcttccacagactggaagagtcct
tcctggtggaagaggataagaagcacgagcggcaccccatcttcggcaacatcgtggacgaggtggcctaccacgagaagtaccc
caccatctaccacctgagaaagaaactggtggacagcaccgacaaggccgacctgcggctgatctatctggccctggcccacatgat
caagttccggggccacttcctgatcgagggcgacctgaaccccgacaacagcgacgtggacaagctgttcatccagctggtgcaga
cctacaaccagctgttcgaggaaaaccccatcaacgccagcggcgtggacgccaaggccatcctgtctgccagactgagcaagagc
agacggctggaaaatctgatcgcccagctgcccggcgagaagaagaatggcctgttcggaaacctgattgccctgagcctgggcct
gacccccaacttcaagagcaacttcgacctggccgaggatgccaaactgcagctgagcaaggacacctacgacgacgacctggac
aacctgctggcccagatcggcgaccagtacgccgacctgtttctggccgccaagaacctgtccgacgccatcctgctgagcgacatc
ctgagagtgaacaccgagatcaccaaggcccccctgagcgcctctatgatcaagagatacgacgagcaccaccaggacctgaccct
gctgaaagctctcgtgcggcagcagctgcctgagaagtacaaagagattttcttcgaccagagcaagaacggctacgccggctacatt
gacggcggagccagccaggaagagttctacaagttcatcaagcccatcctggaaaagatggacggcaccgaggaactgctcgtga
agctgaacagagaggacctgctgcggaagcagcggaccttcgacaacggcagcatcccccaccagatccacctgggagagctgc
acgccattctgcggcggcaggaagatttttacccattcctgaaggacaaccgggaaaagatcgagaagatcctgaccttccgcatccc
ctactacgtgggccctctggccaggggaaacagcagattcgcctggatgaccagaaagagcgaggaaaccatcaccccctggaact
tcgaggaagtggtggacaagggcgcttccgcccagagcttcatcgagcggatgaccaacttcgataagaacctgcccaacgagaag
gtgctgcccaagcacagcctgctgtacgagtacttcaccgtgtataacgagctgaccaaagtgaaatacgtgaccgagggaatgaga
aagcccgccttcctgagcggcgagcagaaaaaggccatcgtggacctgctgttcaagaccaaccggaaagtgaccgtgaagcagct
gaaagaggactacttcaagaaaatcgagtgcttcgactccgtggaaatctccggcgtggaagatcggttcaacgcctccctgggcaca
taccacgatctgctgaaaattatcaaggacaaggacttcctggacaatgaggaaaacgaggacattctggaagatatcgtgctgaccct
gacactgtttgaggacagagagatgatcgaggaacggctgaaaacctatgcccacctgttcgacgacaaagtgatgaagcagctgaa
gcggcggagatacaccggctggggcaggctgagccggaagctgatcaacggcatccgggacaagcagtccggcaagacaatcct
ggatttcctgaagtccgacggcttcgccaacagaaacttcatgcagctgatccacgacgacagcctgacctttaaagaggacatccag
aaagcccaggtgtccggccagggcgatagcctgcacgagcacattgccaatctggccggcagccccgccattaagaagggcatcct
gcagacagtgaaggtggtggacgagctcgtgaaagtgatgggccggcacaagcccgagaacatcgtgatcgaaatggccagaga
gaaccagaccacccagaagggacagaagaacagccgcgagagaatgaagcggatcgaagagggcatcaaagagctgggcagc
cagatcctgaaagaacaccccgtggaaaacacccagctgcagaacgagaagctgtacctgtactacctgcagaatgggcgggatat
gtacgtggaccaggaactggacatcaaccggctgtccgactacgatgtggaccatatcgtgcctcagagctttctgaaggacgactcc
atcgacaacaaggtgctgaccagaagcgacaagaaccggggcaagagcgacaacgtgccctccgaagaggtcgtgaagaagatg
aagaactactggcggcagctgctgaacgccaagctgattacccagagaaagttcgacaatctgaccaaggccgagagaggcggcct
gagcgaactggataaggccggcttcatcaagagacagctggtggaaacccggcagatcacaaagcacgtggcacagatcctggac
tcccggatgaacactaagtacgacgagaatgacaagctgatccgggaagtgaaagtgatcaccctgaagtccaagctggtgtccgatt
tccggaaggatttccagttttacaaagtgcgcgagatcaacaactaccaccacgcccacgacgcctacctgaacgccgtcgtgggaa
ccgccctgatcaaaaagtaccctaagctggaaagcgagttcgtgtacggcgactacaaggtgtacgacgtgcggaagatgatcgcca
agagcgagcaggaaatcggcaaggctaccgccaagtacttcttctacagcaacatcatgaactttttcaagaccgagattaccctggcc
aacggcgagatccggaagcggcctctgatcgagacaaacggcgaaaccggggagatcgtgtgggataagggccgggattttgcca
ccgtgcggaaagtgctgagcatgccccaagtgaatatcgtgaaaaagaccgaggtgcagacaggcggcttcagcaaagagtctatc
ctgcccaagaggaacagcgataagctgatcgccagaaagaaggactgggaccctaagaagtacggcggcttcgacagccccacc
gtggcctattctgtgctggtggtggccaaagtggaaaagggcaagtccaagaaactgaagagtgtgaaagagctgctggggatcacc
atcatggaaagaagcagcttcgagaagaatcccatcgactttctggaagccaagggctacaaagaagtgaaaaaggacctgatcatc

aagctgcctaagtactccctgttcgagctggaaaacggccggaagagaatgctggcctctgccggcgaactgcagaagggaaacga
actggccctgccctccaaatatgtgaacttcctgtacctggccagccactatgagaagctgaagggctcccccgaggataatgagcag
aaacagctgtttgtggaacagcacaagcactacctggacgagatcatcgagcagatcagcgagttctccaagagagtgatcctggcc
gacgctaatctggacaaagtgctgtccgcctacaacaagcaccgggataagcccatcagagagcaggccgagaatatcatccacct
gtttaccctgaccaatctgggagcccctgccgccttcaagtactttgacaccaccatcgaccggaagaggtacaccagcaccaaaga
ggtgctggacgccaccctgatccaccagagcatcaccggcctgtacgagacacggatcgacctgtctcagctgggaggcgactctg
gaggatctagcggaggatcctctggcagcgagacaccaggaacaagcgagtcagcaacaccagagagcagtggcggcagcagc
ggcggcagcagcAAGAGTGCGGAATACCTTAACACATTTAGGCTTAGAAACCTGGGAC
TCCCCGTTATGAATAACCTGCACGATATGTCTAAGGCAACCCGAATCAGTGTAGA
GACGCTTAGACTCTTGATATATACCGCCGACTTTCGATACAGAATATATACCGTC
GAGAAGAAGGGCCCAGAGAAACGAATGCGGACCATATACCAACCTAGTAGGGA
GCTGAAAGCGCTTCAGGGCTGGGTACTTCGAAATATCCTTGACAAACTTTCCAGT
TCACCGTTCAGCATTGGTTTCGAGAAACACCAGAGTATTCTGAATAACGCGACAC
CTCACATAGGAGCCAATTTCATCCTCAATATTGACCTGGAGGATTTCTTCCCTAG
CCTTACTGCCAATAAAGTGTTCGGTGTATTCCACAGCCTCGGCTACAACCGACTG
ATTTCATCTGTACTCACAAAAATTTGTTGCTATAAGAACCTGCTCCCTCAAGGGG
CCCCAAGTAGCCCAAAACTGGCAAACCTCATCTGTTCAAAATTGGACTATCGAAT
TCAGGGCTATGCCGGATCCAGGGGCTTGATCTACACGAGATACGCAGACGACCT
GACACTTTCAGCACAATCCATGAAGAAAGTTGTTAAAGCGAGGGATTTTCTTTTT
TCCATTATTCCGTCTGAAGGATTGGTTATTAATTCTAAAAAAACTTGCATTAGTG
GTCCTCGGTCTCAACGAAAGGTTACAGGCCTGGTAATCTCTCAGGAGAAGGTTG
GGATTGGAAGAGAGAAGTATAAAGAGATTCGCGCCAAAATACATCATATTTTTT
GCGGTAAATCATCTGAAATCGAGCACGTAAGGGGATGGCTTTCTTTCATTCTTTC
TGTAGACTCCAAATCTCACCGACGACTCATCACATATATAAGTAAACTGGAGAA
AAAATATGGGAAAAATCCGCTTAACAAGGCTAAAACTaaaaggccggcggccacgaaaaagg
ccggccaggcaaaaaagaaaaagcttgagggcagaggaagtctgctaacatgcggtgacgtggaggagaatcccggccctgctag
catggtgagcaagggcgaggaggataacatggccatcatcaaggagttcatgcgcttcaaggtgcacatggagggctccgtgaacg
gccacgagttcgagatcgagggcgagggcgagggccgcccctacgagggcacccagaccgccaagctgaaggtgaccaagggt
ggccccctgcccttcgcctgggacatcctgtcccctcagttcatgtacggctccaaggcctacgtgaagcaccccgccgacatccccg
actacttgaagctgtccttccccgagggcttcaagtgggagcgcgtgatgaacttcgaggacggcggcgtggtgaccgtgacccagg
actcctccctgcaggacggcgagttcatctacaaggtgaagctgcgcggcaccaacttcccctccgacggccccgtaatgcagaaga
agaccatgggctgggaggcctcctccgagcggatgtaccccgaggacggcgccctgaagggcgagatcaagcagaggctgaagc
tgaaggacggcggccactacgacgctgaggtcaagaccacctacaaggccaagaagcccgtgcagctgcccggcgcctacaacg
tcaacatcaagttggacatcacctcccacaacgaggactacaccatcgtggaacagtacgaacgcgccgagggccgccactccacc
ggcggcatggacgagctgtacaagtgagaattcctagagctcgctgatcagcctcgactgtgccttctagttgccagccatctgttgttt
gcccctcccccgtgccttccttgaccctggaaggtgccactcccactgtcctttcctaataaaatgaggaaattgcatcgcattgtctgag
taggtgtcattctattctggggggtggggtggggcaggacagcaagggggaggattgggaagagaatagcaggcatgctggggag
cggccgcaggaacccctagtgatggagttggccactccctctctgcgcgctcgctcgctcactgaggccgggcgaccaaaggtcgc
ccgacgcccgggctttgcccgggcggcctcagtgagcgagcgagcgcgcagctgcctgcaggggcgcctgatgcggtattttctcc
ttacgcatctgtgcggtatttcacaccgcatacgtcaaagcaaccatagtacgcgccctgtagcggcgcattaagcgcggcgggtgtg
gtggttacgcgcagcgtgaccgctacacttgccagcgccctagcgcccgctcctttcgctttcttcccttcctttctcgccacgttcgccg
gctttccccgtcaagctctaaatcgggggctccctttagggttccgatttagtgctttacggcacctcgaccccaaaaaacttgatttgggt
gatggttcacgtagtgggccatcgccctgatagacggtttttcgccctttgacgttggagtccacgttctttaatagtggactcttgttccaa
actggaacaacactcaaccctatctcgggctattcttttgatttataagggattttgccgatttcggCctattggttaaaaaatgagctgattt
aacaaaaatttaacgcgaattttaacaaaatattaacgtttacaattttatggtgcactctcagtacaatctgctctgatgccgcatagttaag
ccagccccgacacccgccaacacccgctgacgcgccctgacgggcttgtctgctcccggcatccgcttacagacaagctgtgaccg
tctccgggagctgcatgtgtcagaggttttcaccgtcatcaccgaaacgcgcgagacgaaagggcctcgtgatacgcctatttttatag
gttaatgtcatgataataatggtttcttagacgtcaggtggcacttttcggggaaatgtgcgcggaacccctatttgtttatttttctaaataca
ttcaaatatgtatccgctcatgagacaataaccctgataaatgcttcaataatattgaaaaaggaagagtatgagtattcaacatttccgtgt
cgcccttattcccttttttgcggcattttgccttcctgtttttgctcacccagaaacgctggtgaaagtaaaagatgctgaagatcagttgggt

gcacgagtgggttacatcgaactggatctcaacagcggtaagatccttgagagttttcgccccgaagaacgttttccaatgatgagcact
tttaaagttctgctatgtggcgcggtattatcccgtattgacgccgggcaagagcaactcggtcgccgcatacactattctcagaatgact
tggttgagtactcaccagtcacagaaaagcatcttacggatggcatgacagtaagagaattatgcagtgctgccataaccatgagtgat
aacactgcggccaacttacttctgacaacgatcggaggaccgaaggagctaaccgcttttttgcacaacatgggggatcatgtaactcg
ccttgatcgttgggaaccggagctgaatgaagccataccaaacgacgagcgtgacaccacgatgcctgtagcaatggcaacaacgtt
gcgcaaactattaactggcgaactacttactctagcttcccggcaacaattaatagactggatggaggcggataaagttgcaggaccac
ttctgcgctcggcccttccggctggctggtttattgctgataaatctggagccggtgagcgtggaagccgcggtatcattgcagcactg
gggccagatggtaagccctcccgtatcgtagttatctacacgacggggagtcaggcaactatggatgaacgaaatagacagatcgct
gagataggtgcctcactgattaagcattggtaactgtcagaccaagtttactcatatatactttagattgatttaaaacttcatttttaatttaaa
aggatctaggtgaagatcctttttgataatctcatgaccaaaatcccttaacgtgagttttcgttccactgagcgtcagaccccgtagaaaa
gatcaaaggatcttcttgagatcctttttttctgcgcgtaatctgctgcttgcaaacaaaaaaaccaccgctaccagcggtggtttgtttgcc
ggatcaagagctaccaactctttttccgaaggtaactggcttcagcagagcgcagataccaaatactgttcttctagtgtagccgtagtta
ggccaccacttcaagaactctgtagcaccgcctacatacctcgctctgctaatcctgttaccagtggctgctgccagtggcgataagtcg
tgtcttaccgggttggactcaagacgatagttaccggataaggcgcagcggtcgggctgaacggggggttcgtgcacacagcccag
cttggagcgaacgacctacaccgaactgagatacctacagcgtgagctatgagaaagcgccacgcttcccgaagggagaaaggcg
gacaggtatccggtaagcggcagggtcggaacaggagagcgcacgagggagcttccagggggaaacgcctggtatctttatagtc
ctgtcgggtttcgccacctctgacttgagcgtcgatttttgtgatgctcgtcaggggggcggagcctatggaaaaacgccagcaacgc
ggc

### Supplemental References


1. Richardson CD, Ray GJ, DeWitt MA, Curie GL, Corn JE. Enhancing homology-directed genome editing by catalytically active and inactive CRISPR-Cas9 using asymmetric donor DNA. *Nat Biotechnol*. Mar 2016;34(3):339-44. doi:10.1038/nbt.3481
2. Chu VT, Weber T, Wefers B, et al. Increasing the efficiency of homology-directed repair for CRISPR-Cas9-induced precise gene editing in mammalian cells. *Nat Biotechnol*. May 2015;33(5):543-8. doi:10.1038/nbt.3198
